# Third-generation in situ hybridization chain reaction: multiplexed, quantitative, sensitive, versatile, robust

**DOI:** 10.1101/285213

**Authors:** Harry M.T. Choi, Maayan Schwarzkopf, Mark E. Fornace, Aneesh Acharya, Georgios Artavanis, Johannes Stegmaier, Alexandre Cunha, Niles A. Pierce

**Affiliations:** Division of Biology & Biological Engineering, California Institute of Technology, Pasadena, CA 91125, USA; Division of Chemistry & Chemical Engineering, California Institute of Technology, Pasadena, CA 91125, USA; Center for Advanced Methods in Biological Image Analysis, Beckman Institute, California Institute of Technology, Pasadena, CA 91125, USA; Institute for Automation & Applied Informatics, Karlsruhe Institute of Technology, Karlsruhe, Germany; Institute of Imaging & Computer Vision, RWTH Aachen University, Aachen, Germany; Center for Data-Driven Discovery, California Institute of Technology, Pasadena, CA 91125, USA; Division of Engineering & Applied Science, California Institute of Technology, Pasadena, CA 91125, USA; Weatherall Institute of Molecular Medicine, University of Oxford, Oxford OX3 9DS, UK

**Keywords:** in situ HCR v3.0, qHCR imaging, qHCR flow cytometry, dHCR imaging, multiplexed in situ hybridization, quantitative in situ hybridization, single-molecule mRNA imaging, mRNA flow cytometry, whole-mount vertebrate embryos, mammalian cells, bacterial cells, split-initiator probes, automatic background suppression

## Abstract

In situ hybridization based on the mechanism of hybridization chain reaction (HCR) has addressed multi-decade challenges to imaging mRNA expression in diverse organisms, offering a unique combination of multiplexing, quantitation, sensitivity, resolution, and versatility. Here, with third-generation in situ HCR, we augment these capabilities using probes and amplifiers that combine to provide automatic background suppression throughout the protocol, ensuring that even if reagents bind non-specifically within the sample they will not generate amplified background. Automatic background suppression dramatically enhances performance and robustness, combining the benefits of higher signal-to-background with the convenience of using unoptimized probe sets for new targets and organisms. In situ HCR v3.0 enables multiplexed quantitative mRNA imaging with subcellular resolution in the anatomical context of whole-mount vertebrate embryos, multiplexed quantitative mRNA flow cytometry for high-throughput single-cell expression profiling, and multiplexed quantitative single-molecule mRNA imaging in thick autofluorescent samples.

**SUMMARY:** In situ hybridization chain reaction (HCR) v3.0 exploits automatic background suppression to enable multiplexed quantitative mRNA imaging and flow cytometry with dramatically enhanced ease-of-use and performance.

---

HCR provides isothermal enzyme-free signal amplification in diverse technological settings in vitro, in situ, and in vivo (Ikbal *et al.*, 2015; Bi *et al.*, 2017). Each HCR amplifier consists of two species of kinetically trapped DNA hairpins (H1 and H2; Figure 1A) that co-exist metastably on lab time scales, storing the energy to drive a conditional self-assembly cascade upon exposure to a cognate DNA initiator sequence (I1) (Dirks *et al.*, 2004; Choi *et al.*, 2014). Initiator I1 hybridizes to the input domain of hairpin H1, opening the hairpin to expose its output domain, which in turn hybridizes to the input domain of hairpin H2, exposing its output domain which is identical in sequence to initiator I1, thus providing the basis for a chain reaction of alternating H1 and H2 polymerization steps.

**Fig. 1:**
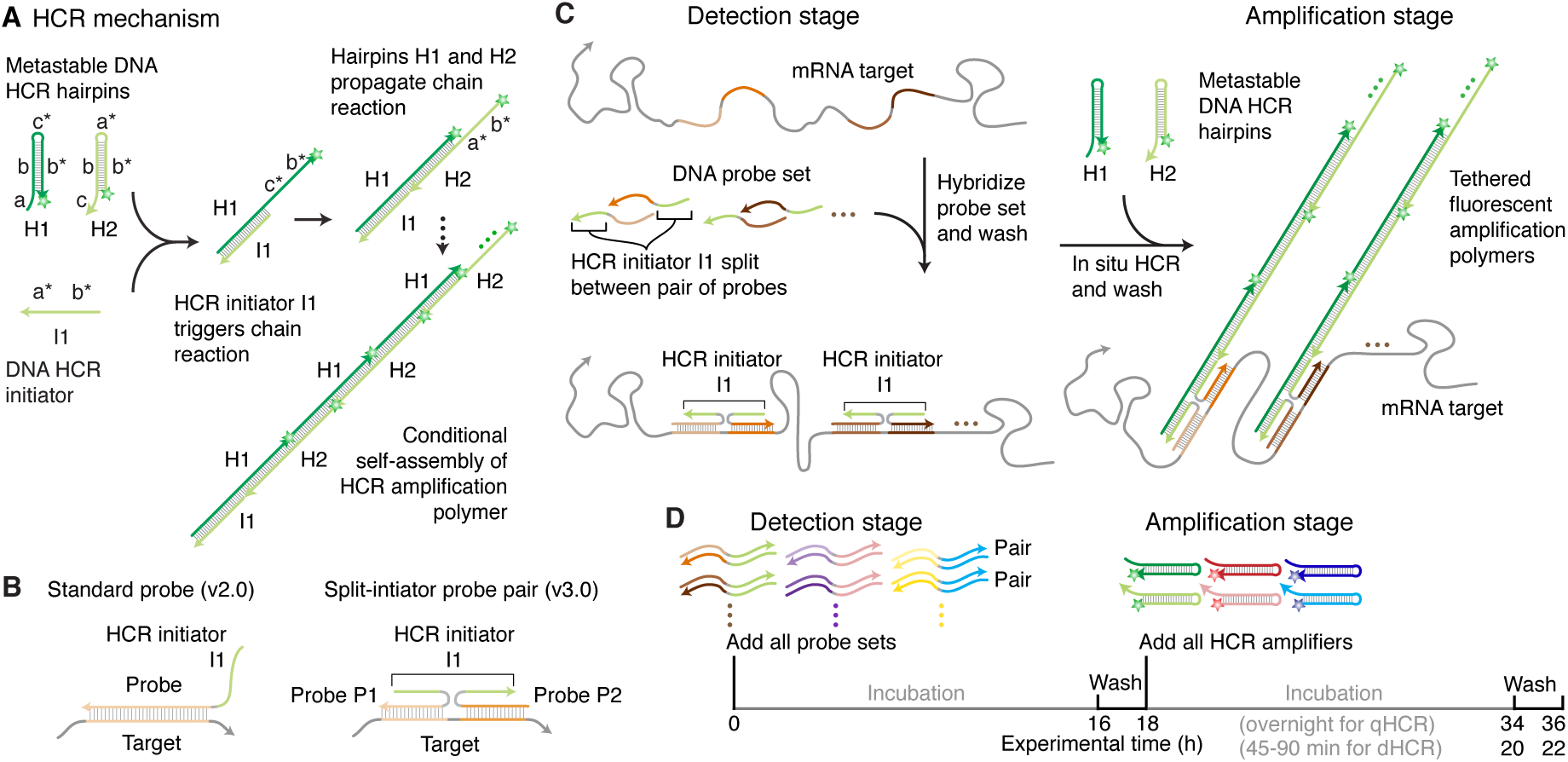
In situ HCR v3.0 using split-initiator probes. (A) HCR mechanism. Green stars denote fluorophores. Arrowhead denotes 3’ end of each strand. (B) Comparison of standard probes (v2.0) and split-initiator probes (v3.0). Standard probes carry full HCR initiator I1 and generate amplified background if they bind non-specifically. Split-initiator probes P1 and P2 each carry half of HCR initiator I1, and do not generate amplified background if they bind non-specifically. (C) 2-stage in situ HCR protocol. Detection stage: probe sets hybridize to mRNA targets, unused probes are washed from the sample. Amplification stage: specifically bound probe pairs trigger self-assembly of a tethered fluorescent amplification polymer, unused hairpins are washed from the sample. Automatic background suppression throughout the protocol: any reagents that bind non-specifically do not lead to generation of amplified background. (D) Multiplexing timeline. The same 2-stage protocol is used independent of the number of target mRNAs. HCR amplification is performed overnight for qHCR imaging and qHCR flow cytometry experiments (to maximize signal-to-background) and for 45-90 min for dHCR imaging experiments (to resolve individual molecules as diffraction-limited dots).

In the context of fluorescence in situ hybridization experiments, where the objective is to image mRNA expression patterns within fixed biological specimens, the role of HCR in situ amplification is to boost the signal above background autofluorescence inherent to the sample. Using in situ HCR v2.0, the initiator I1 is appended to DNA probes complementary to a target mRNA of interest, triggering the self-assembly of fluorophore-labeled H1 and H2 hairpins into tethered fluorescent amplification polymers (Choi *et al.*, 2014; Shah *et al.*, 2016b; Choi *et al.*, 2016). In situ HCR v2.0 enables state-of-the-art mRNA imaging in challenging imaging settings (Choi *et al.*, 2016) including whole-mount vertebrate embryos and thick tissue sections, offering three unique capabilities: straightforward multiplexing with simultaneous 1-stage signal amplification for up to 5 targets (Choi *et al.*, 2014), analog mRNA relative quantitation in an anatomical context (qHCR imaging) (Trivedi *et al.*, 2018), digital mRNA absolute quantitation in an anatomical context (dHCR imaging) (Shah *et al.*, 2016b).

Using in situ HCR v2.0, each target mRNA is detected using multiple probes each carrying a full HCR initiator I1 (Figure 1B; left). If a probe binds non-specifically within the sample, initiator I1 will nonetheless trigger HCR, generating amplified background that decreases the signal-to-background ratio of the image. As a result, using in situ HCR v2.0, it is critical to use probe sets that exclude probes that bind non-specifically, sometimes necessitating probe set optimization in which probes are tested individually to remove “bad probes”. To enhance robustness and eliminate the potential need for probe set optimization when exploring new targets, in situ HCRv3.0 employs probe and amplifier concepts that combine to achieve automatic background suppression throughout the protocol, ensuring that even if a reagent binds non-specifically within the sample, it will not lead to generation of amplified background.

Automatic background suppression is inherent to HCR hairpins because polymerization is conditional on the presence of the initiator I1; individual H1 or H2 hairpins that bind non-specifically in the sample do not trigger formation of an amplification polymer. Hence, the needed innovation is a probe concept that will generate initiator I1 conditionally upon detection of the target mRNA. In situ HCR v3.0 achieves this goal by replacing each *standard probe* carrying the full HCR initiator I1 (Figure 1B; left) with a pair of cooperative *split-initiator probes* that each carry half of HCR initiator I1 (Figure 1B; right). Probe pairs that hybridize specifically to their adjacent binding sites on the target mRNA colocalize the two halves of initiator I1, enabling cooperative initiation of HCR signal amplification. Meanwhile, any individual probes that bind non-specifically in the sample do not colocalize the two halves of initiator I1, do not trigger HCR, and thus suppress generation of amplified background.

## RESULTS

### Validation of split-initiator HCR suppression in vitro and in situ

We first tested split-initiator HCR suppression in solution using gel studies to quantify conversion of HCR hairpins into HCR amplification polymers (Figure 2). There is minimal leakage of hairpins H1 and H2 out of their kinetically trapped states in the absence of HCR initiator I1 (Lane 1). This result demonstrates the automatic background suppression that HCR provides during the amplification stage of an in situ hybridization protocol: if a hairpin binds non-specifically in the sample, it does not trigger HCR, and hence does not generate amplified background. As a positive control, we then verified that HCR initiator I1 triggers full conversion of HCR hairpins into amplification polymers (Lane 2). If initiator I1 is carried by a standard probe, amplification polymers would represent either amplified signal or amplified background depending on whether or not the probe is bound specifically to the target. It is this conceptual weakness that split-initiator probes seek to eliminate. Using a pair of split-initiator probes (P1 and P2) that each carry half of HCR initiator I1, we expect HCR to be triggered if and only if both P1 and P2 bind specifically to their adjacent binding sites on the target. Consistent with this expectation, we observe strong conversion of hairpins H1 and H2 into amplification polymer if P1 and P2 are both introduced with the target (Lane 3), but minimal conversion into polymer if either P1 or P2 is introduced alone (Lanes 4 and 5), reflecting the HCR suppression capabilities of split-initiator probes. Indeed, if the target is absent, even if P1 and P2 are present in solution together, we observe minimal conversion of hairpins into polymer (Lane 6). These results indicate that replacement of a standard probe (v2.0) with a pair of split-initiator probes (v3.0) is expected to modestly decrease amplified signal (lane 2 vs lane 3) but to dramatically decrease amplified background (lane 2 vs lanes 4 and 5). Gel studies of five HCR amplifiers demonstrate typical HCR suppression of *≈*60-fold (Figures S3 and S4; lane 3 vs lanes 4 and 5) using split-initiator probes.

**Fig. 2:**
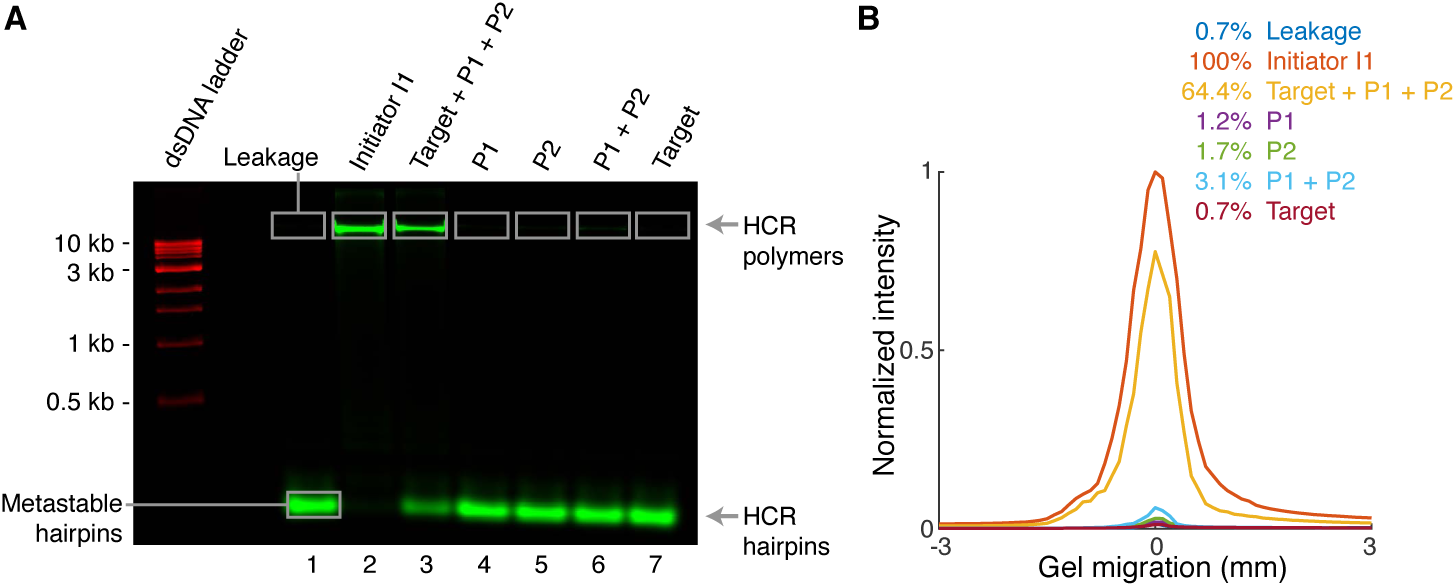
Test tube validation of split-initiator HCR suppression. (A) Agarose gel electrophoresis. Reaction conditions: hairpins H1 and H2 at 0.5 *µ*M each (Lanes 1-7); initiator I1, probes P1 and P2 (each carrying half of initiator I1; Figure 1B), and/or DNA target at 5 nM each (lanes noted on the gel); 5 *×* SSCT buffer; overnight reaction at room temperature. Hairpins H1 and H2 labeled with Alexa 647 fluorophore (green channel). dsDNA 1 kb ladder pre-stained with SYBR Gold (red channel). (B) Quantification of polymer band in panel A. See Figures S3 and S4 for additional data.

We then measured split-initiator HCR suppression in situ by comparing the signal using full probe sets (i.e., both odd and even probes) vs partial probe sets that eliminate one probe from each pair (i.e., only odd probes or only even probes). For five HCR amplifiers, we observe typical HCR suppression of *≈*50-fold (Table S9) using split-initiator probes in situ.

### In situ validation of automatic background suppression with split-initiator probes in whole-mount chicken embryos

We next compared the performance of standard probes (v2.0) and split-initiator probes (v3.0) in the challenging imaging environment of whole-mount chicken embryos (Figure 3). Using standard probes, as the probe set size is increased from 5 to 10 to 20 probes by adding untested probes to a previously validated set of 5 probes (Choi *et al.*, 2016), the background increases dramatically (panel A; magenta) and the signal-to-background ratio *decreases* monotonically (panel B; magenta). Using split-initiator probe pairs that address nearly identical target subsequences, increasing the probe set size causes no measurable change in the background (panel A; orange) and the signal-to-background ratio *increases* monotonically (panel B; orange). Representative images using the largest of these unoptimized probe sets (20 standard probes or 20 split-initiator probe pairs) exhibit high background using standard probes and no visible background using split-initiator probes (panel C); corresponding pixel intensity histograms for regions of high expression (Signal + Background) and no or low expression (Background) are overlapping using standard probes and non-overlapping using split-initiator probes (panel D). These data illustrate the significant benefit of automatic background suppression using split-initiator probes: even if there are non-specific probes in the probe set, they do not generate amplified background, so it is straightforward to increase the signal-to-background ratio simply by increasing the probe set size without probe set optimization.

**Fig. 3:**
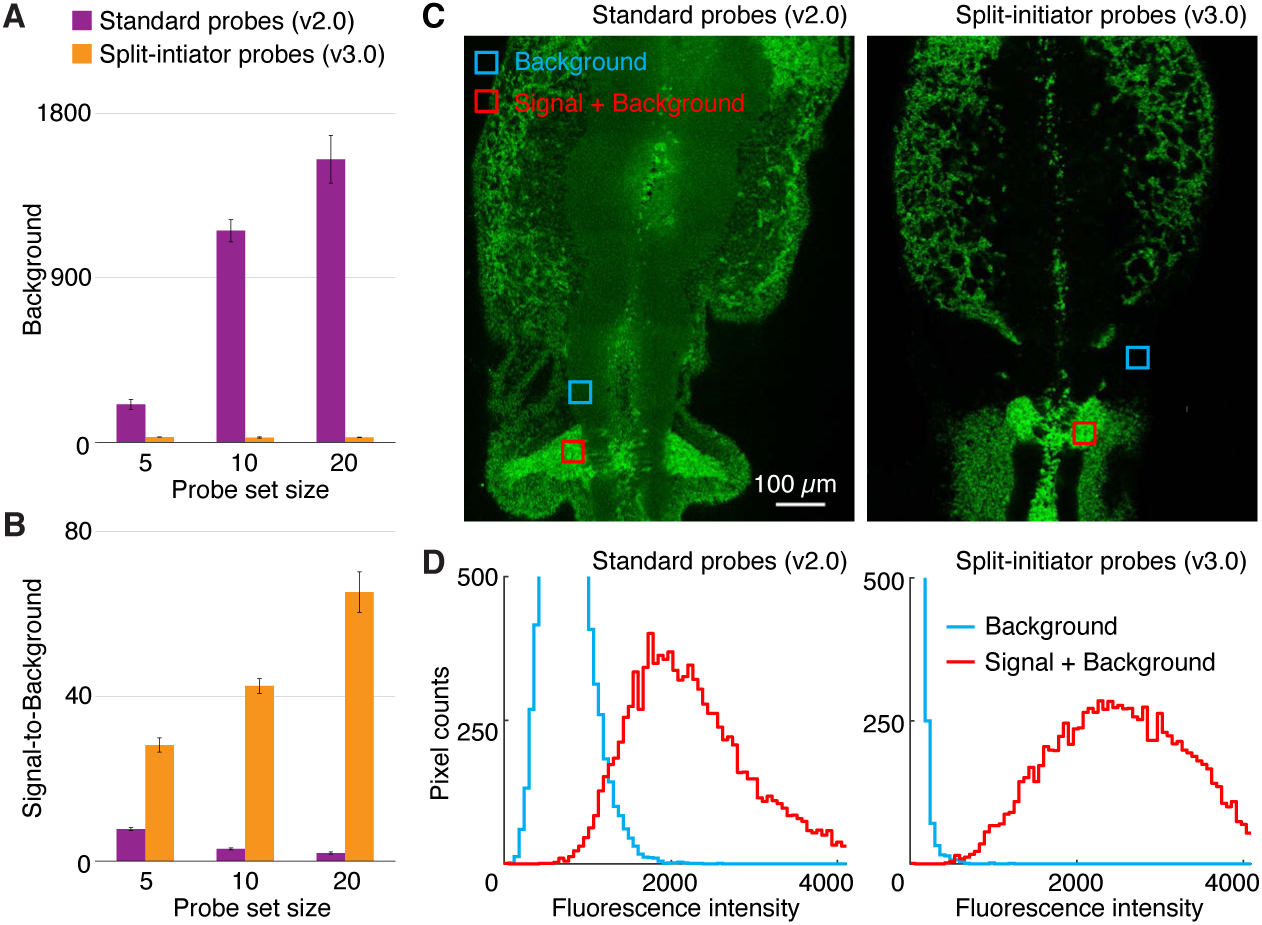
In situ validation of automatic background suppression with split-initiator probes in whole-mount chicken embryos. (A) Fluorescent background and (B) signal-to-background as probe set size is increased by adding unoptimized probes: total of 5, 10, or 20 standard probes (v2.0) vs 5, 10, or 20 split-initiator probe pairs (v3.0). Any standard probes that bind non-specifically will generate amplified background, necessitating probe set optimization; split-initiator probes eliminate the potential need for probe set optimization by providing automatic background suppression. (C) Confocal micrographs in the neural crest of fixed whole-mount chicken embryos. Probe set: 20 standard probes (left) or 20 split-initiator probe pairs (right). (D) Pixel intensity histograms for background and signal plus background (pixels in the depicted regions of panel C): overlapping distributions using unoptimized standard probes, non-overlapping distributions using unoptimized split-initiator probes. Embryos fixed: stage HH 11. Target mRNA: *Sox10*. See Figures S5–S11 and Tables S10–S14 for additional data.

This improved performance is not simply an increase in selectivity resulting from use of probes with a shorter target-binding site (50 nt for standard probes vs 25 nt for each split-initiator probe within a pair): if the split-initiator probe set with 20 probe pairs is modified so that one probe within each pair carries the full initiator I1 (with its partner carrying no initiator), the background increases by an order of magnitude (Figure S9 and Table S12) and the signal-to-background ratio decreases by 1–2 orders of magnitude (Figure S10 and Table S13).

### Multiplexed mRNA imaging in whole-mount chicken embryos with large unoptimized split-initiator probe sets

To test the robustness of automatic background suppression, we performed a 4-channel multiplexed experiment using large unoptimized split-initiator probe sets (v3.0) in the neural crest of whole-mount chicken embryos (Figure 4). Three target mRNAs (*EphA4, Sox10, Dmbx1*) were each detected with 20 split-initiator probe pairs and one shorter target mRNA (*FoxD3*) was detected with 12 split-initiator probe pairs. We observed signal-to-background for each channel ranging from approximately 27 to 59 without probe set optimization. This level of performance was achieved for all targets simultaneously in 4-channel images using fluorophores that compete with lower autofluorescence (Alexa 647) as well as with higher autofluorescence (Alexa 488).

**Fig. 4:**
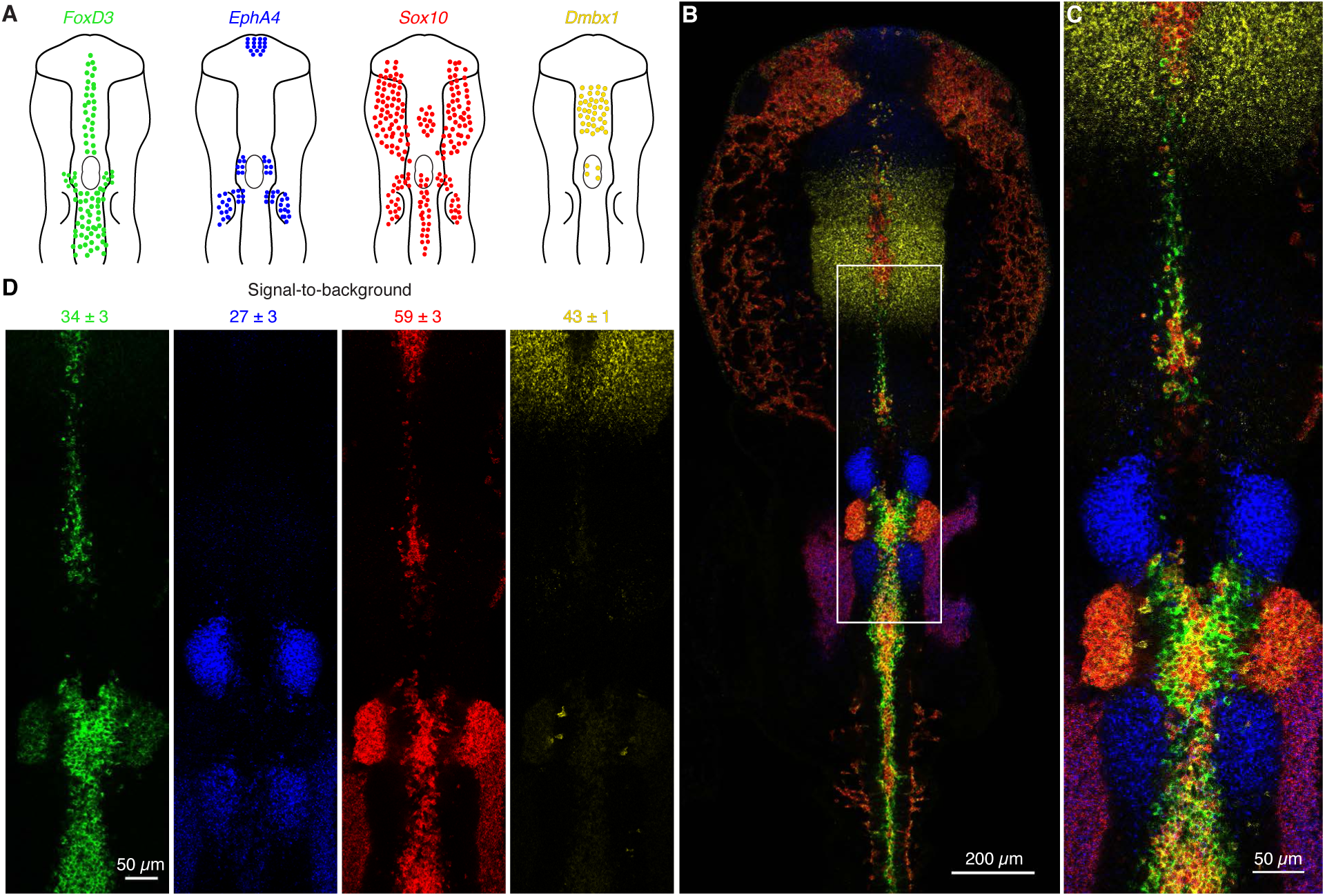
Multiplexed mRNA imaging in whole-mount chicken embryos with large unoptimized probe sets using in situ HCR v3.0. (A) Expression schematics for four target mRNAs in the head and neural crest: *FoxD3, EphA4, Sox10, Dmbx1*. (B) Four-channel confocal micrograph. (C) Zoom of depicted region of panel B. (D) Four individual channels from panel C with signal-to-background measurements (mean *±* standard error, *N* = 3 embryos). Probe sets: 12-20 split-initiator probe pairs per target. Amplifiers: four orthogonal HCR amplifiers carrying spectrally distinct fluorophores. Embryo fixed: stage HH 10. See Figure S12 and Table S15 for additional data.

By comparison, we previously optimized standard probe sets (v2.0) for three target mRNAs in the neural crest of whole-mount chicken embryos (Choi *et al.*, 2016). Starting with 13 to 16 standard probes (each carrying 2 HCR initiators), we arrived at optimized probe sets of 5 to 9 probes, achieving signal-to-background ratios of approximately 5 to 8 (Choi *et al.*, 2016). This represents good performance after an initial investment of labor to perform probe set optimization, but even optimized standard probe sets do not perform as well as unoptimized split-initiator probe sets. Split-initiator probes not only dramatically improve ease-of-use by removing the need for probe set optimization, they also dramatically increase signal-to-background, offering a win/win proposition over standard probes.

### qHCR imaging: analog mRNA relative quantitation with sub-cellular resolution in an anatomical context

We previously demonstrated that in situ HCR v2.0 overcomes the longstanding tradeoff between RNA quantitation and anatomical context, using *optimized* standard HCR probe sets to perform analog mRNA relative quantitation (qHCR imaging) with subcellular resolution within whole-mount vertebrate embryos (Trivedi *et al.*, 2018). Precision increases with probe set size (Trivedi *et al.*, 2018), so the prospect of using large *unoptimized* split-initiator probe sets is highly appealing. To test mRNA relative quantitation with automatic background suppression, we redundantly detected target mRNAs using two split-initiator probe sets each triggering a different spectrally-distinct HCR amplifier (Figure 5AB). If HCR signal scales approximately linearly with the number of target mRNAs per voxel, a 2-channel scatter plot of normalized voxel intensities will yield a tight linear distribution with approximately zero intercept. Conversely, observing a tight linear distribution with approximately zero intercept (Figure 5C), we conclude that the HCR signal scales approximately linearly with the number of target mRNAs per imaging voxel, after first ruling out potential systematic crowding effects that could permit pairwise voxel intensities to slide undetected along the line (Supplementary Figures S13 and S23). Using 20 unoptimized split-initiator probe pairs (v3.0) per channel, the observed accuracy (linearity with zero intercept) and precision (scatter around the line) are both excellent for subcellular 2.1*×*2.1*×*2.7 *µ*m voxels within a whole-mount chicken embryo. Just as quantitative PCR (qPCR) enables analog mRNA relative quantitation in vitro (Gibson *et al.*, 1996; Heid *et al.*, 1996), qHCR imaging enables analog mRNA relative quantitation in situ.

**Fig. 5:**
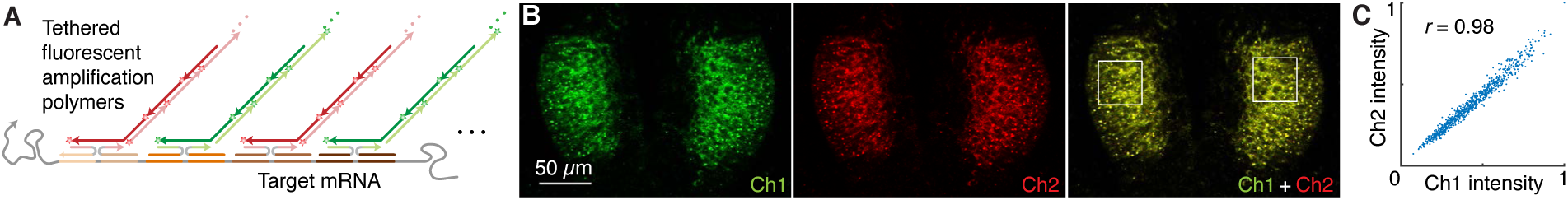
qHCR imaging: analog mRNA relative quantitation with subcellular resolution in an anatomical context. (A) 2-channel redundant detection of target mRNA *EphA4* in a whole-mount chicken embryo. The target is detected using two probe sets, each initiating an orthogonal spectrally distinct HCR amplifier (Ch1: Alexa 546, Ch2: Alexa 647). Confocal microscopy: 0.2*×*0.2 *µ*m pixels. Probe sets: 20 split-initiator probe pairs per channel; no probe set optimization. Embryo fixed: stage HH 10. (B) High accuracy and precision for mRNA relative quantitation in an anatomical context. Highly correlated normalized signal (Pearson correlation coefficient, *r*) for subcellular 2.1 *×* 2.1 *×* 2.7 *µ*m voxels in the selected regions of panel B. Accuracy: linearity with zero intercept. Precision: scatter around the line. See Figures S18 and S19 and Table S16 for additional data.

### qHCR flow cytometry: analog mRNA relative quantitation for high-throughput analysis of human and bacterial cells

The accuracy, precision, and resolution achieved using qHCR imaging suggests the potential for mRNA analog relative quantitation in high-throughput flow cytometry and cell sorting studies. In this case, the instrument treats each cell as a voxel, with both signal and background integrated over the volume of the cell. Using qHCR flow cytometry with 10-20 split-initiator probe pairs per channel (v3.0), we observe high signal-to-background (Figure 6A) and excellent accuracy and precision (Figure 6B) for both human and bacterial cells. Multiplexed qHCR flow cytometry (Figure S27 and S28) will enable high-throughput expression profiling without the need for engineering reporter lines (e.g., for profiling stem cell heterogeneity or sorting bacterial species in heterogeneous environmental samples).

**Fig. 6:**
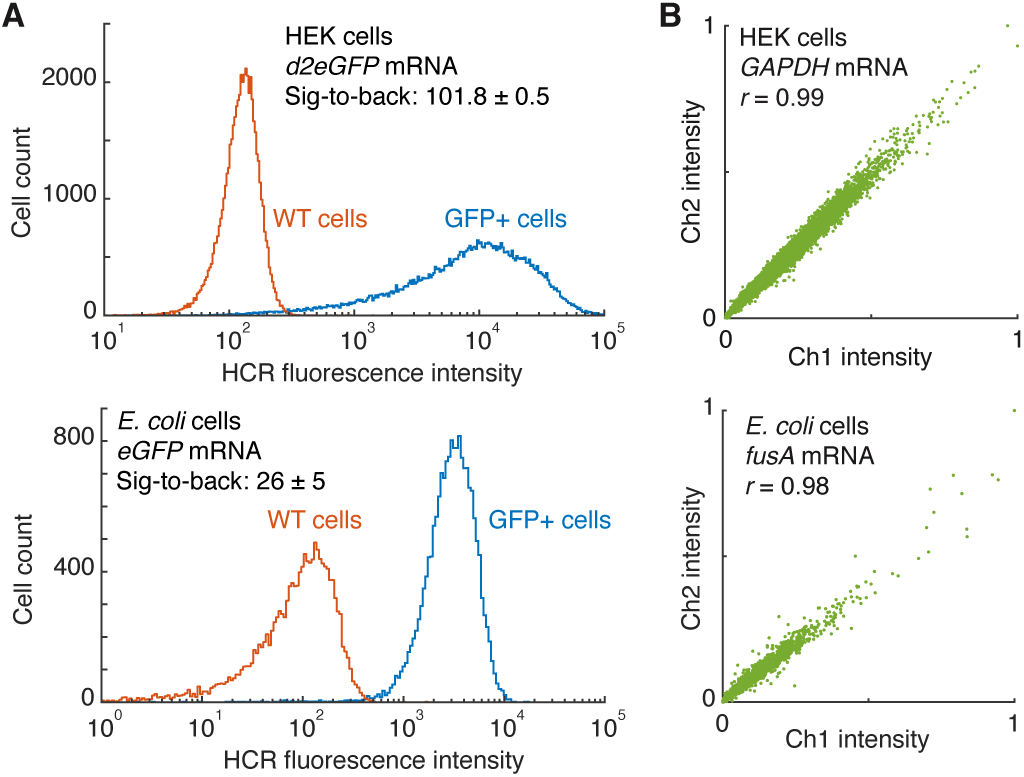
qHCR flow cytometry: analog mRNA relative quantitation for high-throughput analysis of human and bacterial cells. (A) High signal-to-background for transgenic target mRNAs. Mean *±* standard error, *N* = 55,000 HEK cells (top), *N* = 18,000 *E. coli* cells (bottom). Probe sets: 12 split-initiator probe pairs; no probe set optimization. (B) High accuracy and precision for high-throughput mRNA relative quantitation. 2-channel redundant detection of endogenous target mRNAs. Each target mRNA is detected using two probe sets, each initiating an orthogonal spectrally distinct HCR amplifier (Ch1: Alexa 488, Ch2: Alexa 594). Highly correlated normalized signal (Pearson correlation coefficient, *r*), *N* = 20,000 HEK cells (top), *N* = 3,400 *E. coli* cells (bottom). Accuracy: linearity with zero intercept. Precision: scatter around the line. Probe sets: 10 split-initiator probe pairs per channel for *GAPDH*, 18 split-initiator probe pairs per channel for *fusA*; no probe set optimization. See Figures S20–S27 and Tables S17–S23 for additional data.

### dHCR imaging: digital mRNA absolute quantitation in an anatomical context

We have previously shown that in situ HCR v2.0 achieves single-molecule sensitivity and resolution even in thick autofluorescent samples (e.g., 0.5 mm cleared adult mouse brain sections) (Shah *et al.*, 2016b), providing a basis for digital mRNA absolute quantitation (dHCR imaging). For dHCR imaging, we employ large probe sets (to distinguish mRNAs bound by multiple probes from background) and short amplification times (to grow short amplification polymers and resolve individual mRNAs as diffraction-limited dots). Because it is impractical to optimize large probe sets, it is especially appealing to use split-initiator probe sets that offer automatic background suppression and require no optimization.

To validate dHCR imaging using split-initiator probes, we redundantly detected individual mRNA targets using two independent probes sets and HCR amplifiers. We then used dot detection methods from the computer vision community to automatically identify dots in each channel (Supplementary Section S1.6.6). As mRNA false-positive and false-negative rates for each channel go to zero, the colocalization fraction for each channel (fraction of dots in a given channel that are in both channels) will approach one from below. Using large un-optimized split-initiator probe sets (23–25 split-initiator probe pairs per channel), we observe colocalization fractions of *≈*0.84 in cultured human cells and whole-mount chicken embryos (Figure 7). These results improve significantly on the colocalization fractions of *≈*0.50 observed in our previous dHCR imaging studies using unoptimized standard probe sets (39 standard probes per channel) in whole-mount zebrafish embryos (Figure S31)(Shah *et al.*, 2016b). Just as digital PCR (dPCR) enables digital mRNA absolute quantitation in vitro (Vogelstein & Kinzler, 1999; Sanders *et al.*, 2013), dHCR imaging enables digital mRNA absolute quantitation in situ.

**Fig. 7:**
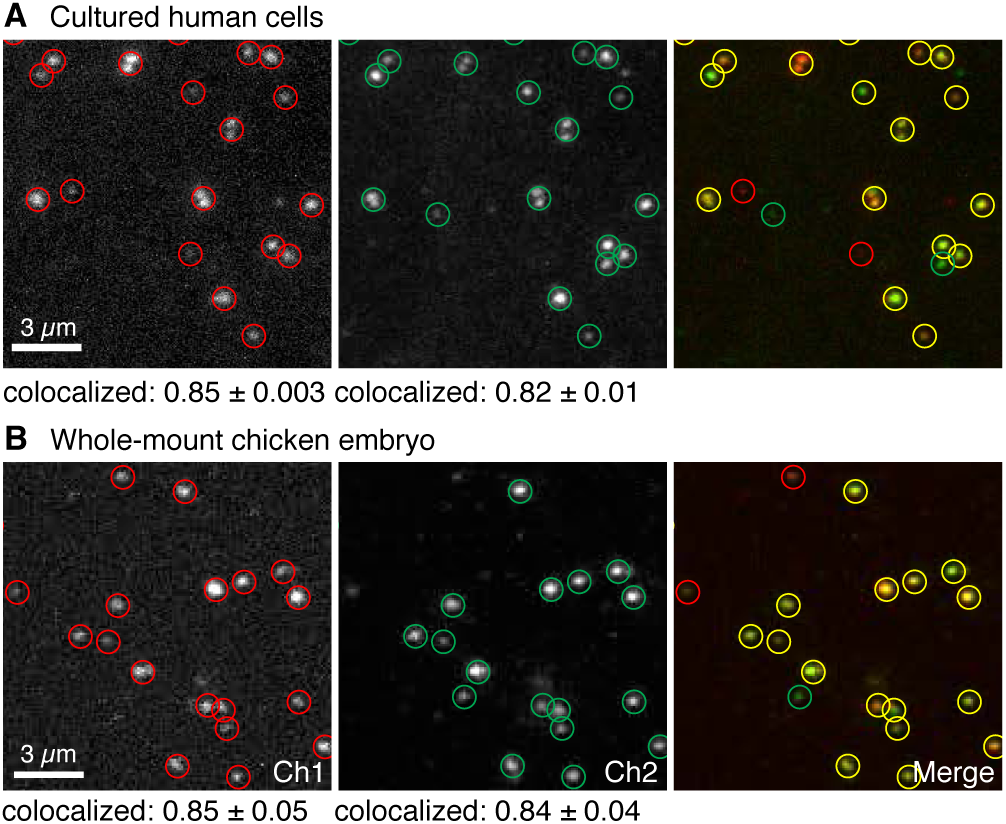
dHCR imaging: digital mRNA absolute quantitation in cultured human cells and whole-mount chicken embryos. (A) Redundant detection of target mRNA *BRAF* in HEK cells. Probe sets: 23 split-initiator probe pairs per channel; no probe set optimization. Pixel size: 0.06 *×*0.06 *µ*m. (B) Redundant detection of target mRNA *Dmbx1* in whole-mount chicken embryos. 25 split-initiator probe pairs per channel; no probe set optimization. Pixel size: 0.1*×* 0.1 *µ*m. Embryos fixed: stage HH 8. (A,B) Each target mRNA is detected using two probe sets, each initiating an orthogonal spectrally distinct HCR amplifier (Ch1: Alexa 647, Ch2: Alexa 546 for panel A; Ch1: Alexa 647, Ch2: Alexa 594 for panel B). Representative field-of-view from confocal micrographs. Red circles: dots detected in Ch1. Green circles: dots detected in Ch2. Yellow circles: dots detected in both channels. Colocalization represents fraction of dots in one channel that are detected in both channels (mean *±* standard error, *N* = 3 slides for panel A, *N* = 3 embryos for panel B). See Figure S29–S31 and Tables S25–S27 for additional data.

## DISCUSSION

### Comparison of alternative probe schemes

To fully appreciate the automatic background suppression properties of split-initiator probes combined with HCR amplifiers, it is helpful to compare alternative concepts. Figure 8 depicts five related in situ hybridization schemes. In a multistage scheme, we say that a method provides automatic background suppression during a given stage if non-specific binding of a reagent during that stage predominantly does not lead to generation of amplified background during subsequent stages. As the final stage of each scheme, signal amplification is performed using HCR. Because HCR hairpins are kinetically trapped and execute a conditional self-assembly cascade that is triggered by the HCR initiator, hairpins that bind non-specifically within the sample predominantly do not trigger growth of HCR amplification polymers. Hence, HCR provides automatic background suppression during the final stage of all five schemes. The challenge then, is to devise a probe concept that maintains automatic background suppression during the earlier stages of the protocol.

**Fig. 8:**
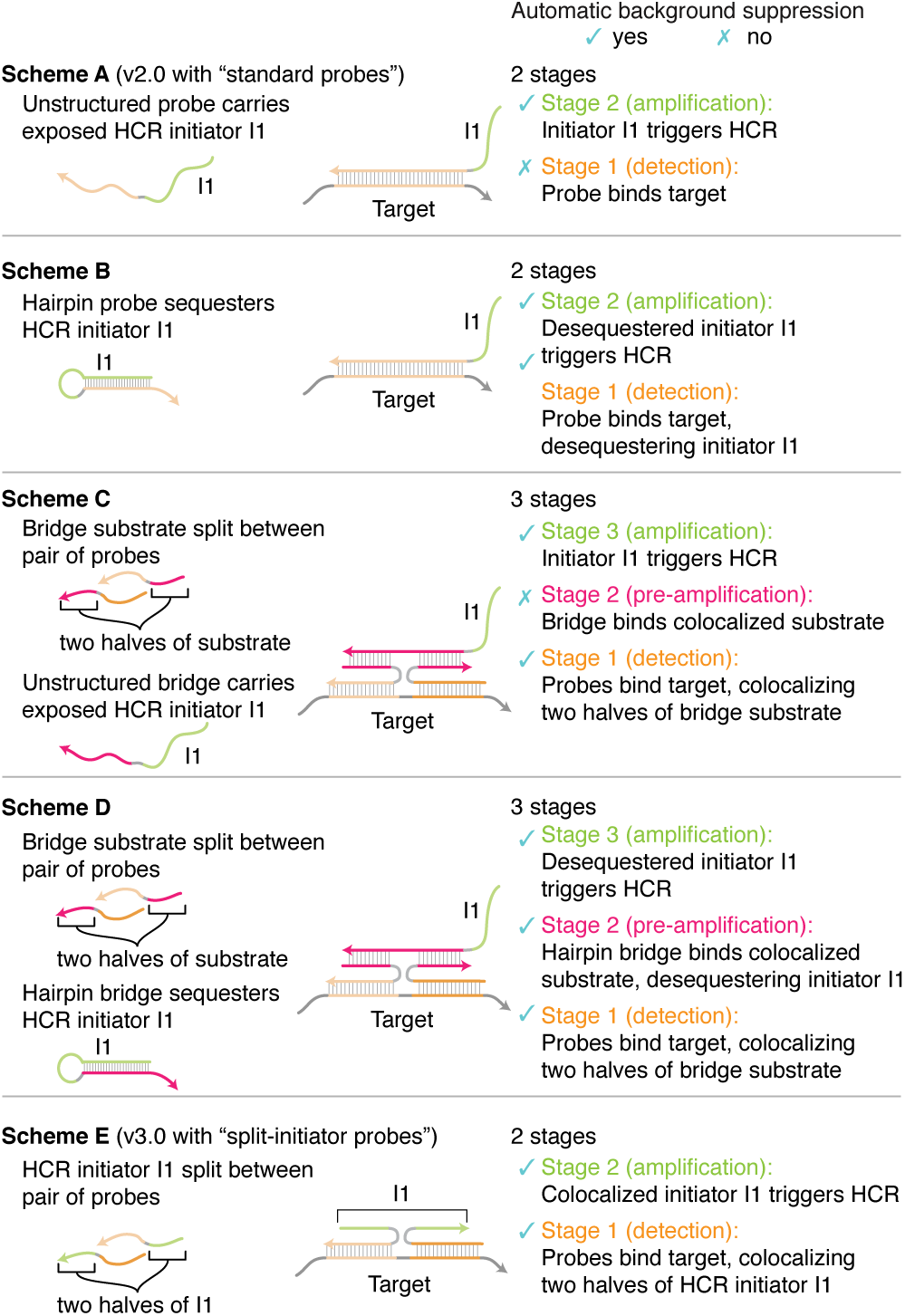
Comparison of probe concepts. Scheme A corresponds to in situ HCR v2.0 with standard probes. Scheme E corresponds to in situ HCR v3.0 with split-initiator probes. Scheme A is vulnerable to non-specific probe binding in Stage 1 leading to amplified background in Stage 2. Scheme B provides automatic background suppression throughout the protocol at the cost of introducing sequence dependence between the target and the HCR amplifier. Scheme C provides automatic background suppression in Stage 1 but is vulnerable to non-specific bridge binding in Stage 2 leading to amplified background in Stage 3 (a weakness shared by the pre-amplification and amplification stages (Stages 2 and 3) of 4-stage bDNA methods (Wang *et al.*, 2012)). Scheme D provides automatic background suppression throughout the protocol at the cost of using a 3-stage protocol. Scheme E offers all of the advantages and none of the disadvantages of Schemes A, B, C, and D, providing automatic background suppression throughout the protocol, avoiding sequence dependence between the HCR amplifier and the target mRNA, and employing a 2-stage protocol. Arrowhead denotes 3’ end of each strand.

To provide a starting point for discussion, Scheme A depicts the standard probes used for in situ HCR v2.0 (Choi *et al.*, 2014; Shah *et al.*, 2016b; Choi *et al.*, 2016). As previously noted, because each probe carries an exposed HCR initiator I1, this scheme has the drawback that non-specific probe binding in Stage 1 will lead to generation of amplified background during Stage 2.

Scheme B resolves this issue by using a hairpin probe that sequesters HCR initiatior I1, exposing the initiator only upon hybridization to the target. As a result, probes that bind non-specifically during Stage 1 predominantly do not generate amplified background during Stage 2, ensuring automatic background suppression throughout the protocol. Unfortunately, suppressing background via conformation change of a hairpin probe imposes sequence dependence between the target and the HCR amplifier, which would necessitate use of a custom HCR amplifier for each new target.

To sidestep this sequence dependence issue, Scheme C uses colocalization instead of conformation change as an alternative principle for achieving automatic background suppression. During Stage 1, the target is detected using a pair of probes that each carry half of a bridge substrate. Specific hybridization of the probes to the target molecule colocalizes the two halves of the bridge substrate. During Stage 2, an unstructured bridge strand that carries exposed HCR initiator I1 is designed to bind stably to the colocalized substrate, but not to either half alone. Thus non-specific binding of either probe during Stage 1 predominantly will not generate amplified background during Stage 2. The drawback to Scheme C is that non-specific binding of the bridge strand during Stage 2 will lead to generation of amplified background during Stage 3. In essence, the unstructured bridge strand in Scheme C has the same conceptual weakness as the unstructured probe in Scheme A.

The principles, strengths, and weaknesses underlying Stages 1 and 2 of Scheme C are similar to those of branched DNA methods (bDNA), which use a 4-stage protocol (Wang *et al.*, 2012): Stage 1) target detection with a pair of probes each carrying half of a bridge substrate; Stage 2) pre-amplification with an unstructured bridge strand that binds to a colocalized bridge substrate and carries multiple exposed amplifier substrates; Stage 3) amplification with an unstructured amplifier strand that binds to an exposed amplifier substrate and carries multiple exposed label substrates; Stage 4) signal generation with an unstructured label strand that binds to an exposed label substrate. This approach has the conceptual strength that non-specific binding of individual probes during Stage 1 will predominantly not lead to generation of amplified background (as only bridge substrates colocalized by the target will mediate amplification), but also the conceptual weakness that non-specific binding of reagents in Stages 2 or 3 will lead to generation of amplified background (as unstructured bridge strands carry exposed amplifier substrates and unstructured amplifier strands carry exposed label substrates). Hence, automatic background suppression is achieved in Stage 1 based on the principle of colocalization, but then not maintained during Stages 2 and 3 as a result of reliance on unstructured strands that carry exposed substrates for downstream reagents.

To achieve automatic background suppression throughout the protocol, Scheme D improves on Scheme C by replacing the unstructured bridge strand with a hairpin bridge that initially sequesters HCR initiator I1, exposing I1 only upon hybridizing to the colocalized bridge substrate. Automatic background suppression is achieved in Stage 1 based on the principle of colocalization and then maintained during Stage 2 based on the principle of conformation change. The drawback of Scheme D is the increase in number of stages from 2 to 3.

As the final step in the derivation of split-initiator probes, Scheme E simplifies Scheme D by noting that the conformation-change property of the hairpin bridge is also a property of the HCR hairpins used for amplification. Therefore with Scheme E, we stipulate that the bridge substrate is an HCR initiator sequence, enabling HCR hairpins to bridge between colocalized probes and amplify the signal in a single stage. As a result, Scheme E becomes a 2-stage protocol.

Scheme E, which provides the basis for in situ HCR v3.0 in the current work, provides all of the benefits and none of the drawbacks of the other four schemes. First, we have the simplicity of a 2-stage protocol (Stage 1: detection, Stage 2: amplification). Second, we have the flexibility of sequence independence between the target and the HCR amplifier, enabling use of a validated library of HCR amplifiers for new targets of interest. Third, we have the robustness of automatic background suppression throughout the protocol: at every stage during the protocol, non-specific binding of reagents will predominantly not lead to generation of amplified background.

### Enhanced robustness and signal-to-background

Automatic background suppression using split-initiator probes has important consequences for both robustness and signal-to-background. Using standard probes, increasing the size of the probe set will reliably increase amplified signal but might increase amplified background even more, so use of a large v2.0 probe set can be a double-edged sword; probe set optimization is sometimes required to ensure that increasing probe set size does more good than harm. By contrast, using split-initiator probe sets, the signal-to-background ratio increases reliably with probe set size, so it is advantageous to use large v3.0 probe sets without optimization and achieve high signal-to-background on the first try.

### qHCR and dHCR quantitative imaging modes

In situ HCR enables two quantitative imaging modes in thick autofluorescent samples:

- qHCR imaging: analog mRNA relative quantitation with subcellular resolution; HCR signal is analog in the form of fluorescence voxel intensities that scale approximately linearly with the number of target molecules per voxel.
- dHCR imaging: digital mRNA absolute quantitation; HCR signal is digital in the form of diffraction-limited dots representing individual target molecules.

For qHCR imaging, we recommend using 20 split-initiator probe pairs per target and amplifying overnight. For dHCR imaging, we recommend maximizing the number of probe pairs per target (at least 25 probe pairs is preferred) and amplifying for 45-90 minutes. Note that because the qHCR signal per imaging voxel is quantitative, it will naturally decrease to zero as the number of targets per voxel decreases to zero; for sufficiently low expression, the signal will not be observable above autofluorescence. However, the dHCR signal per target molecule does not decrease with expression level. Hence, the qHCR and dHCR quantitative imaging modes are complementary, with qHCR suitable for medium-and high-copy targets (where the quantitative signal dominates autofluorescent background), and dHCR suitable for low-copy targets (where the signal from individual target molecules can be spatially separated). The same probe set can be used for either imaging mode, so imaging can be performed in qHCR mode (longer amplification time, lower magnification) or dHCR imaging mode (shorter amplification time, higher magnification) depending on the expression level observed in situ.

### Quantitative read-out and read-in

The quantitative properties of in situ HCR enable gene expression queries in two directions (Trivedi *et al.*, 2018): *read-out* from anatomical space to expression space to discover co-expression relationships in selected regions of the specimen; conversely, *read-in* from multidimensional expression space to anatomical space to discover those anatomical locations in which selected gene co-expression relationships occur. Quantitative read-out and read-in analyses provide the strengths of flow cytometry expression analyses, but by preserving anatomical context, they enable bi-directional queries that open a new era for in situ hybridization (Trivedi *et al.*, 2018). In situ HCR v3.0 using large split-initiator probe sets enhances accuracy and precision for read-out/read-in using either qHCR relative quantitation (Trivedi *et al.*, 2018) or dHCR absolute quantitation (Shah *et al.*, 2016a).

### In situ HCR resolves longstanding shortcomings of traditional CARD in situ amplification methods

Fluorescent in situ hybridization methods are used across the life sciences to image mRNA expression within fixed cells, tissues, and organisms. In challenging imaging settings, including whole-mount vertebrate embryos and thick tissue sections, autofluorescence within the sample necessitates the use of in situ amplification to boost the signal-to-background ratio (Tautz & Pfeifle, 1989; Harland, 1991; Lehmann & Tautz, 1994; Kerstens *et al.*, 1995; Nieto *et al.*, 1996; Wiedorn *et al.*, 1999; Player *et al.*, 2001; Pernthaler *et al.*, 2002; Thisse *et al.*, 2004; Denkers *et al.*, 2004; Kosman *et al.*, 2004; Zhou *et al.*, 2004; Larsson *et al.*, 2004; Clay & Ramakrishnan, 2005; Barroso-Chinea *et al.*, 2007; Acloque *et al.*, 2008; Piette *et al.*, 2008; Thisse & Thisse, 2008; Weiszmann *et al.*, 2009; Larsson *et al.*, 2010; Wang *et al.*, 2012). For decades, traditional in situ amplification approaches based on catalytic reporter deposition (CARD) have been the dominant approach for generating high signal-to-background in samples with high autofluorescence (Tautz & Pfeifle, 1989; Harland, 1991; Lehmann & Tautz, 1994; Kerstens *et al.*, 1995; Nieto *et al.*, 1996; Pernthaler *et al.*, 2002; Kosman *et al.*, 2004; Thisse *et al.*, 2004; Denkers *et al.*, 2004; Clay & Ramakrishnan, 2005; Barroso-Chinea *et al.*, 2007; Acloque *et al.*, 2008; Piette *et al.*, 2008; Thisse & Thisse, 2008; Weiszmann *et al.*, 2009) despite three significant drawbacks: multiplexing is cumbersome due to the need to perform signal amplification for one target mRNA at a time (Lehmann & Tautz, 1994; Nieto *et al.*, 1996; Thisse *et al.*, 2004; Denkers *et al.*, 2004; Kosman *et al.*, 2004; Clay & Ramakrishnan, 2005; Barroso-Chinea *et al.*, 2007; Acloque *et al.*, 2008; Piette *et al.*, 2008), staining is qualitative rather than quantitative due to the nonlinear effects of the CARD amplification cascade, and spatial resolution is routinely compromised by diffusion of reporter molecules prior to deposition (Tautz & Pfeifle, 1989; Thisse *et al.*, 2004; Thisse & Thisse, 2008; Acloque *et al.*, 2008; Piette *et al.*, 2008; Weiszmann *et al.*, 2009).

In situ HCR v2.0 overcame these longstanding difficulties, enabling multiplexed, quantitative, high-resolution imaging of mRNA expression with high signal-to-background in diverse organisms including whole-mount vertebrate embryos (Choi *et al.*, 2014; Choi *et al.*, 2016; Trivedi *et al.*, 2018). Orthogonal HCR amplifiers operate independently within the sample so the experimental timeline for multiplexed experiments is independent of the number of target mRNAs (Choi *et al.*, 2010; Choi *et al.*, 2014). The amplified HCR signal scales approximately linearly with the number of target molecules, enabling accurate and precise mRNA relative quantitation with subcellular resolution in the anatomical context of whole-mount vertebrate embryos (Trivedi *et al.*, 2018). Amplification polymers remain tethered to their initiating probes, enabling imaging of mRNA expression with subcellular or single-molecule resolution as desired (Choi *et al.*, 2014; Shah *et al.*, 2016b; Choi *et al.*, 2016). With split-initiator probes, in situ HCR v3.0 adds the performance and robustness benefits of automatic background suppression, providing biologists with an enhanced state-of-the-art research tool for the study of mRNA expression (Table 1).

**Table 1:** mRNA imaging using in situ HCR

|  |  |
| --- | --- |
| simple | 2-stage protocol independent of number of targets |
| amplified | boost signal above autofluorescence |
| multiplexed | amplification for up to 5 targets simultaneously |
| quantitative | signal scales linearly with target abundance |
| penetrating | whole-mount vertebrate embryos and thick tissue sections |
| resolved | subcellular or single-molecule resolution as desired |
| sensitive | single molecules detected even in thick autofluorescent samples |
| versatile | suitable for use in diverse organisms |
| robust | automatic background suppression throughout protocol |

## Acknowledgments

We thank C. Calvert for assistance with test tube validation of split-initiator probes, J. Tan-Cabugao and M. Simons Costa in the M. Bronner Lab for preparation of chicken embryos, A. Collazo and S. Wilbert of the Caltech Biological Imaging Facility (BIF) for assistance with imaging, R. Diamond and D. Perez of the Caltech Flow Cytometry Facility (FCF) for assistance with flow cytometry, and M. Mann and the D. Baltimore Lab for generously providing access to their flow cytometer for preliminary studies.

## Author contributions

Conceptualization: H.M.T.C., N.A.P.; Methodology: H.M.T.C., M.S., M.F., N.A.P.; Software: M.F., J.S., A.C.; Validation: H.M.T.C., M.S.; Investigation: H.M.T.C., M.S., A.A., G.A.; Writing-original draft: H.M.T.C., M.S., N.A.P.; Writing -review & editing: H.M.T.C., M.S., M.F., A.A., G.A., J.S., A.C., N.A.P.; Visualization: H.M.T.C., M.S., N.A.P.; Supervision: N.A.P.; Project administration: N.A.P.; Funding acquisition: N.A.P.

## Supplementary information

Supplementary information available online.

