## Supplementary Materials for "Third-generation in situ hybridization chain reaction: multiplexed, quantitative, sensitive, versatile, robust"

### S1 Materials and methods

#### S1.1 Probe sets, amplifiers, and buffers

For each target mRNA, a kit containing a DNA probe set, a DNA HCR amplifier, and hybridization, wash, and amplification buffers was purchased from Molecular Technologies (moleculartechnologies.org), a non-profit academic resource within the Beckman Institute at Caltech. For gel studies, see Table S1 for sequence information. For in situ HCR studies, see Table S2 for a summary of sample, probe set, and amplifier details and Section S4 for probe sequences. Sequences for HCR amplifiers B1, B2, B3, B4, and B5 are given in Section S6 of Choi *et al.* (2014).

| HCR Amplifier | Oligo | Length (nt) | Sequence (5′ to 3′) | Figures |
|---|---|---|---|---|
| B1-Alexa647 | I1 | 36 | gAggAgggCAgCAAACgggAAgAgTCTTCCTTTACg | S3 |
| | P1 | 45 | gAggAgggCAgCAAACggAAAgACgTTgTggCTgTTgTAgTTgTA | S3 |
| | P2 | 45 | CgTTCTTCTgCTTgTCggCCATgATTAgAAgAgTCTTCCTTTACg | S3 |
| B2-Alexa647 | I1 | 36 | CCTCgTAAATCCTCATCAATCATCCAgTAAACCgCC | S3 |
| | P1 | 45 | CCTCgTAAATCCTCATCAAAAgACgTTgTggCTgTTgTAgTTgTA | S3 |
| | P2 | 45 | CgTTCTTCTgCTTgTCggCCATgATAAATCATCCAgTAAACCgCC | S3 |
| B3-Alexa647 | I1 | 36 | gTCCCTgCCTCTATATCTCCACTCAACTTTAACCCg | 2, S4 |
| | P1 | 45 | gTCCCTgCCTCTATATCTTTAgACgTTgTggCTgTTgTAgTTgTA | 2, S4 |
| | P2 | 45 | CgTTCTTCTgCTTgTCggCCATgATTTCCACTCAACTTTAACCCg | 2, S4 |
| B4-Alexa647 | I1 | 36 | CCTCAACCTACCTCCAACTCTCACCATATTCgCTTC | S3 |
| | P1 | 45 | CCTCAACCTACCTCCAACAAAgACgTTgTggCTgTTgTAgTTgTA | S3 |
| | P2 | 45 | CgTTCTTCTgCTTgTCggCCATgATATTCTCACCATATTCgCTTC | S3 |
| B5-Alexa647 | I1 | 36 | CTCACTCCCAATCTCTATCTACCCTACAAATCCAAT | S3 |
| | P1 | 45 | CTCACTCCCAATCTCTATAAAgACgTTgTggCTgTTgTAgTTgTA | S3 |
| | P2 | 45 | CgTTCTTCTgCTTgTCggCCATgATAACTACCCTACAAATCCAAT | S3 |

**Table S1. Sequences for gel studies.** Initiator sequence in green, spacer sequence in blue, target-binding sequence in black. In all cases, the Target oligo is 5′-TACAACTACAACAgCCACAACgTCTATATCATggCCgACAAgCAgAAgAACg-3′.

| Organism | Target | Standard probes (v2.0) | Split-initiator probe pairs (v3.0) | HCR amplifier | Amplification time | Figures |
|---|---|---|---|---|---|---|
| *G. gallus domesticus* | *Sox10* | 5, 10, 20 | | B3-Alexa647 | overnight | 3, S5, S7 |
| | *Sox10* | | 5, 10, 20 | B3-Alexa647 | overnight | 3, S6, S8 |
| | *Sox10* | | 20 | B3-Alexa647 | overnight | S9, S10 |
| | *Sox10* | 20 | | B3-Alexa647 | overnight | S9, S10 |
| | *EphA4* | | 20 | B2-Alexa647 | overnight | S11 |
| | *FoxD3* | | 12 | B4-Alexa488 | overnight | 4, S12 |
| | *Dmbx1* | | 20 | B1-Alexa514 | overnight | 4, S12 |
| | *Sox10* | | 20 | B3-Alexa546 | overnight | 4, S12 |
| | *EphA4* | | 20 | B2-Alexa647 | overnight | 4, S12 |
| | *Dmbx1* | | 20 | B1-Alexa546 | overnight | 5, S18 |
| | *Dmbx1* | | 20 | B2-Alexa647 | overnight | 5, S18 |
| | *EphA4* | | 20 | B1-Alexa546 | overnight | 5, S19 |
| | *EphA4* | | 20 | B2-Alexa647 | overnight | 5, S19 |
| *H. sapiens sapiens* | *d2eGFP* | | 12 | B3-Alexa594 | overnight | 6A, S20 |
| | *GAPDH* | | 10 | B5-Alexa488 | overnight | 6B, S24, S23 |
| | *GAPDH* | | 10 | B4-Alexa594 | overnight | 6B, S21, S24 |
| | *ACTB* | | 10 | B2-Alexa594 | overnight | S23 |
| | *PGK1* | | 18 | B1-Alexa488 | overnight | S25 |
| | *PGK1* | | 18 | B2-Alexa594 | overnight | S25, S27 |
| | *GAPDH* | | 10 | B4-Alexa488 | overnight | S27 |
| *E. coli* | *eGFP* | | 12 | B3-Alexa594 | overnight | 6A, S22 |
| | *fusA* | | 18 | B3-Alexa488 | overnight | 6B, S26, S28 |
| | *fusA* | | 18 | B2-Alexa594 | overnight | 6B, S26 |
| | *icd* | | 20 | B1-Alexa594 | overnight | S28 |
| *H. sapiens sapiens* | *BRAF* | | 23 | B3-Alexa647 | 45 min | 7A, S29 |
| | *BRAF* | | 23 | B4-Alexa546 | 45 min | 7A, S29 |
| *G. gallus domesticus* | *Dmbx1* | | 25 | B1-Alexa594 | 90 min | 7B, S30 |
| | *Dmbx1* | | 25 | B2-Alexa647 | 90 min | 7B, S30 |
| *G. gallus domesticus* | *EphA4* | | 20 | B1-Alexa546 | overnight | S13, S14, S15 |
| | *Egr2* | | 20 | B3-Alexa647 | overnight | S13, S14, S16 |

**Table S2. Organisms, target mRNAs, probe sets, amplifiers, and figure numbers for in situ HCR experiments.**

#### S1.2 Gel electrophoresis

DNA HCR reactions for Figures 2, S3, and S4 were performed in 5× SSC with 0.1% Tween 20. All hairpins were labeled with Alexa 647 (green channel). DNA hairpins were snap-cooled separately at 3 $\mu$M in hairpin storage buffer (Molecular Technologies). DNA initiators (I1), split-initiator probes (P1, P2), and target (Target) were diluted to 0.03 $\mu$M in 5× SSC. Each lane was prepared by mixing 0.8 $\mu$L of 5× SSC, 1.2 $\mu$L of 5× SSC with 1% Tween 20, and 2 $\mu$L of each hairpin. For the lanes with DNA oligos (I1, P1, P2, or Target), 2 $\mu$L of each oligo was added. An appropriate amount of 5× SSC was added to each lane to bring the reaction volume to 12 $\mu$L. The reactions were incubated at room temperature overnight. The samples were supplemented with 3 $\mu$L of 5× gel loading buffer (50% glycerol with bromophenol blue and xylene cyanol tracking dyes) and loaded into a native 1% agarose gel, prepared with 1× LB buffer (Faster Better Media). The gel was run at 150 V for 60 min at room temperature and imaged using an FLA-5100 fluorescent scanner (Fujifilm Life Science) with a 635 nm laser and a 665 nm long-pass filter. The 1 kb DNA ladder (red channel) was prestained with SYBR Gold (Invitrogen) and imaged using a 488 nm laser and a 575 nm long-pass filter.

Multi Gauge software (Fuji Photo Film) was used to calculate the Alexa 647 intensity profile surrounding the polymer band for each lane (lanes 1-7 in Figures 2, S3, and S4). Each intensity profile is displayed for ± 3 mm of gel migration distance with the peak value centered at 0; the intensity values are normalized so that the highest peak value for each gel is set to 1. Signal for each band was calculated using Multi Gauge with auto-detection of signal and background; the calculated percentages were normalized to the measured value with the full initiator (lane 2). Based on repeated analysis using Multi Gauge, the uncertainty in quantifying the bands in any given gel is estimated to be less than 0.1% of the band signal used for normalization.

#### S1.3 In situ hybridization

In situ HCR v2.0 with standard probes was performed using the protocols detailed in Section S8 of Choi *et al.* (2016). In situ HCR v3.0 with split-initiator probes was performed using the protocols detailed in Section S2. For signal amplification in analog qHCR mode (high signal-to-background with quantitative voxel intensities for imaging with subcellular resolution or high-throughput flow cytometry; e.g., Figures 3–6), amplification was performed overnight to generate long HCR amplification polymers. For signal amplification in digital dHCR mode (single-molecule sensitivity and resolution with individual target molecules resolved as diffraction-limited dots; e.g., Figure 7), amplification was performed for 45-90 min to generate short HCR amplification polymers. See Table S2 for the amplification time for each experiment.

#### S1.4 Confocal microscopy

A Zeiss LSM 710 inverted confocal microscope equipped with an LD C-Apochromat 40×/1.1 W Korr M27 objective was used to image whole-mount chicken embryos in Figures 3, 5, S5–S11, S14–S17, S18, and S19. The same microscope equipped with an LD LCI Plan-Apochromat 25×/0.8 Imm Korr DIC M27 was used to image 4-color whole-mount chicken embryos in Figures 4 and S12. A Zeiss LSM 800 inverted confocal microscope equipped with a Plan-Apochromat 63×/1.4 Oil DIC M27 objective was used to image whole-mount chicken embryos in Figures 7 and S30. The same microscope equipped with an alpha Plan-Apochromat 100×/1.46 Oil DIC (UV) M27 objective was used to image mammalian cells in Figures 7 and S29. See Table S3 for a summary of excitation laser sources, beam splitters, and tuned emission bandpass filters used for each experiment. All images are displayed without background subtraction.

For dHCR imaging studies (e.g., Figure 7), TetraSpeck Microspheres (0.2 $\mu$m, fluorescent blue/green/orange/dark red; Thermo Fisher Scientific, Cat. # T7280) were used as references for channel alignment. Images from two channels were registered using the Channel Alignment feature in ZEN Black software (Zeiss) and registration parameters were recorded for alignment of data imaged in dHCR 2-channel redundant detection experiments using identical imaging settings.

| Target | Fluorophore | Laser (nm) | Beam Splitter | Filter (nm) | Pixel size ($x \times y \times z\ \mu$m) | Voxel size ($x \times y \times z\ \mu$m) | Focal planes | Figures |
|---|---|---|---|---|---|---|---|---|
| *EphA4* | Alexa647 | 633 | MBS 488/561/633 | 650–689 | $0.208 \times 0.208 \times 2.7$ | | 1 | S11 |
| *Sox10* | Alexa647 | 633 | MBS 488/561/633 | 650–699 | $0.415 \times 0.415 \times 2.7$ | | 1 | 3, S5–S10 |
| *FoxD3* | Alexa488 | 488 | MBS 488/561/633 | 491–525 | $0.664 \times 0.664 \times 4$ | | 1 | 4, S12 |
| *Dmbx1* | Alexa514 | 514 | MBS 458/514 | 546–564 | $0.664 \times 0.664 \times 4$ | | 1 | 4, S12 |
| *Sox10* | Alexa546 | 561 | MBS 488/561/633 | 573–612 | $0.664 \times 0.664 \times 4$ | | 1 | 4, S12 |
| *EphA4* | Alexa647 | 633 | MBS 488/561/633 | 654–687 | $0.664 \times 0.664 \times 4$ | | 1 | 4, S12 |
| *Dmbx1* | Alexa546 | 561 | MBS 488/561/633 | 563–592 | $0.208 \times 0.208 \times 2.7$ | $2.1 \times 2.1 \times 2.7$ | 1 | 5, S18 |
| *Dmbx1* | Alexa647 | 633 | MBS 488/561/633 | 650–689 | $0.208 \times 0.208 \times 2.7$ | $2.1 \times 2.1 \times 2.7$ | 1 | 5, S18 |
| *EphA4* | Alexa546 | 561 | MBS 488/561/633 | 563–592 | $0.208 \times 0.208 \times 2.7$ | $2.1 \times 2.1 \times 2.7$ | 1 | 5, S19 |
| *EphA4* | Alexa647 | 633 | MBS 488/561/633 | 650–689 | $0.208 \times 0.208 \times 2.7$ | $2.1 \times 2.1 \times 2.7$ | 1 | 5, S19 |
| *BRAF* | Alexa546 | 561 | MBS 405/488/561/640 (T10/R90) | 566–623 | $0.0624 \times 0.0624 \times 0.42$ | | 17 | 7A, S29 |
| *BRAF* | Alexa647 | 640 | MBS 405/488/561/640 (T10/R90) | 656–700 | $0.0624 \times 0.0624 \times 0.42$ | | 17 | 7A, S29 |
| — | DAPI | 405 | MBS 405/488/561/640 (T10/R90) | 410–470 | $0.0624 \times 0.0624 \times 0.42$ | | 17 | S29 |
| *Dmbx1* | Alexa594 | 561 | MBS 405/488/561/640 (T10/R90) | 580–647 | $0.099 \times 0.099 \times 0.420$ | | 22 | 7B, S30 |
| *Dmbx1* | Alexa647 | 640 | MBS 405/488/561/640 (T10/R90) | 645–700 | $0.099 \times 0.099 \times 0.420$ | | 22 | 7B, S30 |
| *EphA4* | Alexa546 | 561 | MBS 488/561/633 | 563–592 | $0.415 \times 0.415 \times 2.7$ | $2.1 \times 2.1 \times 2.7$ | 1 | S14–S17 |
| *Egr2* | Alexa647 | 633 | MBS 488/561/633 | 650–689 | $0.415 \times 0.415 \times 2.7$ | $2.1 \times 2.1 \times 2.7$ | 1 | S14–S17 |

**Table S3. Confocal microscope settings.**

#### S1.5 Flow cytometry

Prior to flow cytometry, cells were filtered through a 35 $\mu$m or a 40 $\mu$m mesh. Flow cytometry studies were performed using a MACSQuant VYB (Miltenyi Biotec). See Table S4 for a summary of excitation laser sources and filters used for each experiment. Flow cytometry data were gated using EasyFlow (Antebi *et al.*, 2017) and plotted using MATLAB (Mathworks). For HEK cells, two gates were applied to data (e.g., Figure S1): a first gate of forward scatter area (FSC-A) vs side scatter area (SSC-A) to select cells, and a second gate of FSC-A vs forward scatter height (FSC-H) to select single cells. Only cells satisfying both gates were used for the analysis. For *E. coli* cells, one gate of FSC-A vs SSC-A was applied to select cells (e.g., Figure S2).

| Target | Fluorophore | Laser (nm) | Filter | Emission (nm) | Figures |
|---|---|---|---|---|---|
| — | d2eGFP | 488 | B1 | 525/50 | S20 |
| *d2eGFP* | Alexa594 | 561 | Y2 | 615/20 | 6A, S20 |
| *GAPDH* | Alexa488 | 488 | B1 | 525/50 | 6B, S23, S24, S27 |
| *GAPDH* | Alexa594 | 635 | Y2 | 615/20 | 6B, S21, S24 |
| *ACTB* | Alexa594 | 635 | Y2 | 615/20 | S23 |
| *PGK1* | Alexa488 | 488 | B1 | 525/50 | S25 |
| *PGK1* | Alexa594 | 561 | Y2 | 615/20 | S25, S27 |
| — | eGFP | 488 | B1 | 525/50 | S22 |
| *eGFP* | Alexa594 | 561 | Y2 | 615/20 | 6A, S22 |
| *fusA* | Alexa488 | 488 | B1 | 525/50 | 6B, S26, S28 |
| *fusA* | Alexa594 | 635 | Y2 | 615/20 | 6B, S26 |
| *icd* | Alexa594 | 635 | Y2 | 615/20 | S28 |

**Table S4. Flow cytometer settings.**

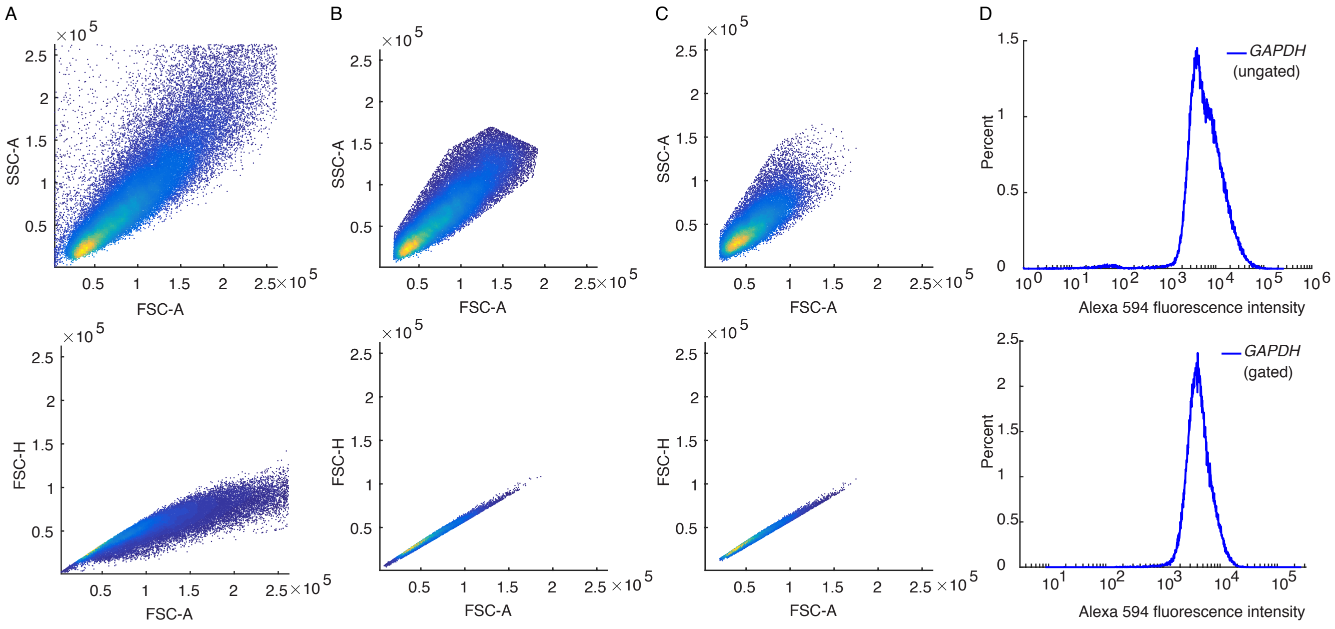

**Figure S1. Illustration of gates used for flow cytometry analysis of HEK cells.** (A) Scatter plots for ungated data. Top: side scatter area (SSC-A) vs forward scatter area (FSC-A). Bottom: forward scatter height (FSC-H) vs forward scatter area (FSC-A). (B) Scatter plots after applying one gate. Top: gate on FSC-A vs SSC-A to remove debris and select cells. Bottom: gate on FSC-A vs FSC-H to remove clumps of cells and select single cells. (C) Scatter plots after applying both gates. (D) Signal plus background for ungated (top) and gated (bottom) samples. Target: *GAPDH*. Probe set: 10 split-initiator probe pairs. Amplifier: B4-Alexa594. The depicted gates were used for the SIG+BACK data in Figure S21.

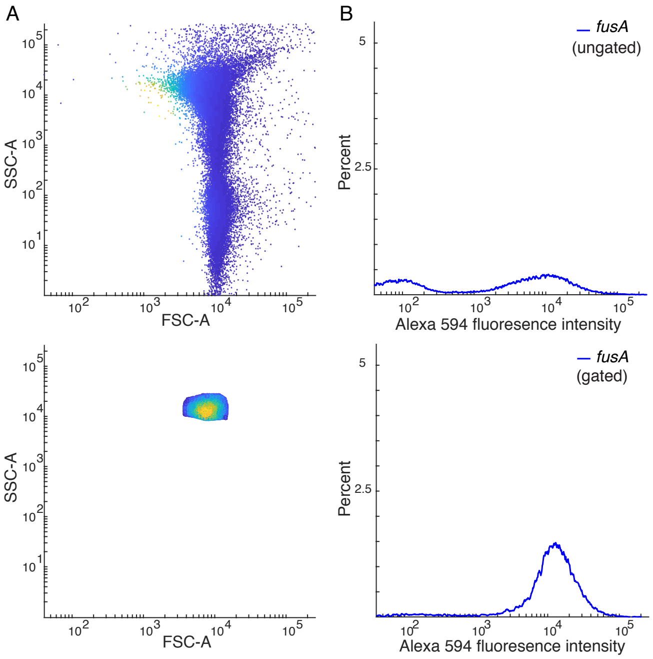

**Figure S2. Illustration of gates used for flow cytometry analysis of *E. coli*.** (A) Scatter plots for ungated sample (top) and gated sample (bottom): side scatter area (SSC-A) vs forward scatter area (FSC-A). (B) Signal plus background for ungated sample (top) and gated sample (bottom). Target: *fusA*. Probe set: 18 split-initiator probe pairs. Amplifier: B4-Alexa594. The depicted gate was used for the SIG+BACK data in Figure S26.

##### S1.6 Image analysis

We build on an image analysis framework developed over a series of publications (Choi *et al.*, 2010, 2014, 2016; Trivedi *et al.*, 2018). For convenience, here we provide a self-contained description of the details relevant to the present work.

###### S1.6.1 Raw pixel intensities

The total fluorescence within a pixel is a combination of signal and background. Fluorescent background (BACK) arises from three sources in each channel:

- autofluorescence (AF): fluorescence inherent to the sample,
- non-specific detection (NSD): probes that bind non-specifically in the sample and subsequently trigger HCR amplification,
- non-specific amplification (NSA): HCR hairpins that bind non-specifically in the sample.

Fluorescent signal (SIG) in each channel corresponds to:

- signal (SIG): probes that hybridize specifically to the target mRNA and subsequently trigger HCR amplification.

For pixel $i$ of replicate embryo $n$, we denote the background

$$X_{n,i}^{\rm BACK} = X_{n,i}^{\rm NSD} + X_{n,i}^{\rm NSA} + X_{n,i}^{\rm AF}, \tag{S1}$$

the signal:

$$X_{n,i}^{\rm SIG}, \tag{S2}$$

and the total fluorescence (SIG+BACK):

$$X_{n,i}^{\rm SIG+BACK} = X_{n,i}^{\rm SIG} + X_{n,i}^{\rm BACK}. \tag{S3}$$

###### S1.6.2 Measurement of signal, background, and signal-to-background

For each target mRNA, background (BACK) is characterized for pixels in a representative rectangular region of no- or low-expression and the combination of signal plus background (SIG+BACK) is characterized for pixels in a representative rectangular region of high expression (e.g., Figures 3C, S7A, S8A, S10A, and S12A). For the pixels in these regions, we characterize the distribution by plotting an intensity histogram (e.g., Figures 3D, S7B, S8B, S10B, and S12B) and characterize average performance by calculating the mean pixel intensity ($\bar{X}_n^{\rm BACK}$ or $\bar{X}_n^{\rm SIG+BACK}$ for replicate embryo $n$). Performance across replicate embryos is characterized by calculating the sample means ($\bar{X}^{\rm BACK}$ and $\bar{X}^{\rm SIG+BACK}$) and standard errors ($s_{\bar{X}^{\rm BACK}}$ and $s_{\bar{X}^{\rm SIG+BACK}}$). The mean signal is then estimated as

$$\bar{X}^{\rm SIG} = \bar{X}^{\rm SIG+BACK} - \bar{X}^{\rm BACK} \tag{S4}$$

with the standard error estimated via uncertainty propagation as

$$s_{\bar{X}^{\rm SIG}} \leq \sqrt{(s_{\bar{X}^{\rm SIG+BACK}})^2 + (s_{\bar{X}^{\rm BACK}})^2}. \tag{S5}$$

The signal-to-background ratio is estimated as:

$$\bar{X}^{\rm SIG/BACK} = \bar{X}^{\rm SIG} / \bar{X}^{\rm BACK} \tag{S6}$$

with standard error estimated via uncertainty propagation as

$$s^{\rm SIG/BACK} \leq \bar{X}^{\rm SIG/BACK} \sqrt{\left(\frac{s_{\bar{X}^{\rm SIG}}}{\bar{X}^{\rm SIG}}\right)^2 + \left(\frac{s_{\bar{X}^{\rm BACK}}}{\bar{X}^{\rm BACK}}\right)^2}. \tag{S7}$$

These upper bounds on estimated standard errors hold under the assumption that the correlation between SIG and BACK is non-negative. Tables S10–S16 display signal, background, and/or signal-to-background values to characterize the performance of in situ HCR v3.0 within whole-mount chicken embryos.

| | Experiment type | Quantity | Reagents: Probes | Reagents: Hairpins | Expression region |
|---|---|---|---|---|---|
| **A** | 1 | SIG+NSD+NSA+AF = SIG+BACK | odd + even | ✓ | high |
| | 1 | NSD+NSA+AF = BACK | odd + even | ✓ | no/low |
| **B** | 2 | NSA+AF | | ✓ | high |
| | 3 | AF | | | high |
| **C** | 4 | $\text{SIG}^{\text{odd}}$+$\text{NSD}^{\text{odd}}$+NSA+AF = $\text{SIG}^{\text{odd}}$+$\text{BACK}^{\text{odd}}$ | odd | ✓ | high |
| | 4 | $\text{NSD}^{\text{odd}}$+NSA+AF = $\text{BACK}^{\text{odd}}$ | odd | ✓ | no/low |
| | 5 | $\text{SIG}^{\text{even}}$+$\text{NSD}^{\text{even}}$+NSA+AF = $\text{SIG}^{\text{even}}$+$\text{BACK}^{\text{even}}$ | even | ✓ | high |
| | 5 | $\text{NSD}^{\text{even}}$+NSA+AF = $\text{BACK}^{\text{even}}$ | even | ✓ | no/low |

**Table S5. Experiment types for qHCR imaging using in situ HCR v3.0.** (A) Characterize signal, background, and signal-to-background. (B) Characterize components of background (AF, NSA, NSD). (C) Characterize split-initiator HCR suppression.

##### S1.6.3 Measurement of background components

Calculation of the signal-to-background ratio (Section S1.6.2) requires only a Type 1 experiment (using the terminology of Table S5A), yielding the values $\bar{X}^{\text{SIG+BACK}}$ and $\bar{X}^{\text{BACK}}$ that are needed to calculate SIG/BACK. If desired, additional control experiments that omit certain reagents can be used to characterize the individual components of background (AF, NSA, NSD). A Type 2 experiment (no probes, hairpins only) yields $\bar{X}^{\text{NSA+AF}}$ and a Type 3 experiment (no probes, no hairpins) yields $\bar{X}^{\text{AF}}$.* The background components can then be estimated via calculations analogous to (S4) and (S5). The estimated means are:

$$\bar{X}^{\text{NSD}} = \bar{X}^{\text{BACK}} - \bar{X}^{\text{NSA+AF}} \tag{S8}$$

$$\bar{X}^{\text{NSA}} = \bar{X}^{\text{NSA+AF}} - \bar{X}^{\text{AF}} \tag{S9}$$

with estimated standard errors:

$$s_{\bar{X}^{\text{NSD}}} \leq \sqrt{(s_{\bar{X}^{\text{BACK}}})^2 + (s_{\bar{X}^{\text{NSA+AF}}})^2} \tag{S10}$$

$$s_{\bar{X}^{\text{NSA}}} \leq \sqrt{(s_{\bar{X}^{\text{NSA+AF}}})^2 + (s_{\bar{X}^{\text{AF}}})^2}. \tag{S11}$$

These upper bounds on estimated standard errors hold under the assumption that the correlations are non-negative for the different components of background. For a given quantity, if the estimated mean is less than the estimated standard error, we report the standard error as an upper bound, and use this bound for uncertainty propagation.

Table S14B provides estimates for AF, NSA, and NSD when imaging *EphA4* in whole-mount chicken embryos using in situ HCR v3.0, all of which are small compared to SIG. Kinetically trapped HCR hairpins automatically suppress NSA and the combination of split-initiator probes and HCR hairpins automatically suppress NSD. Furthermore, HCR generates amplified SIG. All of these factors contribute to achieving a high signal-to-background ratio. If a Type 1 experiment demonstrates SIG $\gg$ BACK, as is typically the case using in situ HCR v3.0, then there is little motivation to perform Type 2 and Type 3 experiments to characterize the individual background components (AF, NSA, NSD) as these are all bounded above by BACK.

##### S1.6.4 Measurement of split-initiator HCR suppression

To characterize performance of in situ HCR v3.0 in suppressing triggering of HCR by individual split-initiator probes, we augment experiments of Type 1 with additional control experiments that omit certain reagents. First, let us define the NSD and SIG observed using odd probes only or even probes only:

*If a microscope generates non-negligible fluorescence intensities in the absence of sample, this so-called instrument noise (NOISE) should be taken into consideration when calculating background and signal contributions, leading to four Experiment Types (1. SIG+BACK+NOISE, 1. BACK+NOISE, 2. NSA+AF+NOISE, 3. AF+NOISE, 4. NOISE; cf. Table S5AB).

- odd non-specific detection (NSD$^{\text{odd}}$): odd probes that bind non-specifically in the sample and subsequently trigger HCR amplification.
- even non-specific detection (NSD$^{\text{even}}$): even probes that bind non-specifically in the sample and subsequently trigger HCR amplification.
- odd signal (SIG$^{\text{odd}}$): odd probes that hybridize specifically to the target mRNA and subsequently trigger HCR amplification.
- even signal (SIG$^{\text{even}}$): even probes that hybridize specifically to the target mRNA in the sample and subsequently trigger HCR amplification.

A Type 4 experiment (odd probes only, with hairpins) yields $\bar{X}^{\text{BACK}^{\text{odd}}+\text{SIG}^{\text{odd}}}$ and $\bar{X}^{\text{BACK}^{\text{odd}}}$ and a Type 5 experiment (even probes only, with hairpins) yields $\bar{X}^{\text{BACK}^{\text{even}}+\text{SIG}^{\text{even}}}$ and $\bar{X}^{\text{BACK}^{\text{even}}}$. These quantities in turn can be used to calculate SIG$^{\text{odd}}$ and SIG$^{\text{even}}$ via calculations analogous to (S4) and (S5). The estimated means are:

$$\bar{X}^{\text{SIG}^{\text{odd}}} = \bar{X}^{\text{SIG}^{\text{odd}}+\text{BACK}^{\text{odd}}} - \bar{X}^{\text{BACK}^{\text{odd}}} \tag{S12}$$

$$\bar{X}^{\text{SIG}^{\text{even}}} = \bar{X}^{\text{SIG}^{\text{even}}+\text{BACK}^{\text{even}}} - \bar{X}^{\text{BACK}^{\text{even}}} \tag{S13}$$

with estimated standard errors:

$$s_{\bar{X}^{\text{SIG}^{\text{odd}}}} \leq \sqrt{(s_{\bar{X}^{\text{SIG}^{\text{odd}}+\text{BACK}^{\text{odd}}}})^2 + (s_{\bar{X}^{\text{BACK}^{\text{odd}}}})^2} \tag{S14}$$

$$s_{\bar{X}^{\text{SIG}^{\text{even}}}} \leq \sqrt{(s_{\bar{X}^{\text{SIG}^{\text{even}}+\text{BACK}^{\text{even}}}})^2 + (s_{\bar{X}^{\text{BACK}^{\text{even}}}})^2} \tag{S15}$$

These upper bounds on estimated standard errors hold under the assumption that the correlations are non-negative for SIG$^{\text{odd}}$ and BACK$^{\text{odd}}$ and for SIG$^{\text{even}}$ and BACK$^{\text{even}}$. For a given quantity, if the estimated mean is less than the estimated standard error, we report the standard error as an upper bound, and use this bound for uncertainty propagation.

Split-initiator HCR suppression can then be characterized by calculating SIG/SIG$^{\text{odd}}$ and SIG/SIG$^{\text{even}}$, with higher values corresponding to more effective suppression. This in situ characterization is akin to the gel studies (Figures 2, S3, and S4) that compare test tube triggering of HCR by odd/even probe pairs colocalized by the target (lane 3) or by either probe alone (lanes 4 or 5). For the in situ data, signal-to-signal ratios are obtained via calculations analogous to (S6) and (S7). The estimated means are:

$$\bar{X}^{\text{SIG}/\text{SIG}^{\text{odd}}} = \bar{X}^{\text{SIG}} / \bar{X}^{\text{SIG}^{\text{odd}}} \tag{S16}$$

$$\bar{X}^{\text{SIG}/\text{SIG}^{\text{even}}} = \bar{X}^{\text{SIG}} / \bar{X}^{\text{SIG}^{\text{even}}} \tag{S17}$$

with estimated standard errors:

$$s^{\text{SIG}/\text{SIG}^{\text{odd}}} \leq \bar{X}^{\text{SIG}/\text{SIG}^{\text{odd}}} \sqrt{\left(\frac{s_{\bar{X}^{\text{SIG}}}}{\bar{X}^{\text{SIG}}}\right)^2 + \left(\frac{s_{\bar{X}^{\text{SIG}^{\text{odd}}}}}{\bar{X}^{\text{SIG}^{\text{odd}}}}\right)^2} \tag{S18}$$

$$s^{\text{SIG}/\text{SIG}^{\text{even}}} \leq \bar{X}^{\text{SIG}/\text{SIG}^{\text{even}}} \sqrt{\left(\frac{s_{\bar{X}^{\text{SIG}}}}{\bar{X}^{\text{SIG}}}\right)^2 + \left(\frac{s_{\bar{X}^{\text{SIG}^{\text{even}}}}}{\bar{X}^{\text{SIG}^{\text{even}}}}\right)^2}. \tag{S19}$$

These upper bounds on estimated standard errors hold under the assumption that the correlations are non-negative for SIG and SIG$^{\text{odd}}$, and for SIG and SIG$^{\text{even}}$.

Table S14C displays the signal-to-signal ratios SIG/SIG$^{\text{odd}}$ and SIG/SIG$^{\text{even}}$ when imaging *EphA4* in whole-mount chicken embryos using in situ HCR v3.0. Split-initiator probes colocalized by the target are more than an order of magnitude more effective at triggering HCR than odd or even probes alone. Interestingly, with this assay, we are quantifying the automatic *background* suppression capabilities of the split-initiator probes by measuring automatic *signal* suppression, taking advantage of the fact that the target molecules in the embryo will colocalize odd/even probe pairs for a Type 1 experiment and will localize odd or even probes to the same expression region for a Type 4 or 5 experiment.

##### S1.6.5 Normalized subcellular voxel intensities for mRNA relative quantitation

For quantitative mRNA imaging using in situ HCR, precision increases with voxel size as long as the imaging voxels remain smaller than the features in the expression pattern (see Section S2.2 of Trivedi *et al.* (2018)). To increase precision, we calculate raw voxel intensities by averaging neighboring pixel intensities while still maintaining a subcellcular voxel size. To facilitate relative quantitation between voxels, we estimate the normalized HCR signal of voxel $j$ in replicate $n$ as:

$$x_{n,j} \equiv \frac{X_{n,j}^{\text{SIG+BACK}} - X^{\text{BOT}}}{X^{\text{TOP}} - X^{\text{BOT}}}, \tag{S20}$$

which translates and rescales the data so that the voxel intensities in each channel fall in the interval [0,1]. Here,

$$X^{\text{BOT}} \equiv \bar{X}^{\text{BACK}} \tag{S21}$$

is the mean background across replicates (see Section S1.6.2) and

$$X^{\text{TOP}} \equiv \max_{n,j} X_{n,j}^{\text{SIG+BACK}} \tag{S22}$$

is the maximum total fluorescence for a voxel across replicates.

Pairwise expression scatter plots that each display normalized voxel intensities for two channels (e.g., Figures 4 and 5 of Trivedi *et al.* (2018)) provide a powerful quantitative framework for performing multidimensional read-out/read-in analyses (Figure 6 of Trivedi *et al.* (2018)). Read-out from anatomical space to expression space enables discovery of expression clusters of voxels with quantitatively related expression levels and ratios (amplitudes and slopes in the expression scatter plots), while read-in from expression space to anatomical space enables discovery of the corresponding anatomical locations of these expression clusters within the embryo. The simple and practical normalization approach of (S20)–(S22) translates and rescales all voxels identically within a given channel (enabling comparison of amplitudes and slopes in scatter plots between replicates), and does not attempt to remove scatter in the normalized signal estimate that is caused by scatter in the background.

To validate relative mRNA quantitation with subcellular resolution (2×2×2.7 $\mu$m voxels) in whole-mount chicken embryos, Figures 5C, S18C, and S19C display highly correlated normalized voxel intensities for 2-channel redundant detection of *Dmbx1* and *EphA4*. In this setting, accuracy corresponds to linearity with zero intercept, and precision corresponds to scatter around the line (Trivedi *et al.*, 2018).

##### S1.6.6 Dot detection and colocalization for dHCR single-molecule imaging

To validate the performance of in situ HCR (v3.0) for single-molecule imaging, we perform a 2-channel redundant detection experiment in which a target mRNA is detected using two independent probe sets and HCR amplifiers. Let $N_1$ denote the number of dots detected in channel 1, $N_2$ the number of dots detected in channel 2, and $N_{12}$ the number of colocalized dots appearing in both channels. We define the colocalization fraction for each channel:

$$C_1 = N_{12}/N_1, \tag{S23}$$

$$C_2 = N_{12}/N_2. \tag{S24}$$

As the false-positive and false-negative rates for single-molecule detection got to zero, $C_1$ and $C_2$ will both approach 1 from below, providing a quantitative basis for evaluating performance. Colocalization results using in situ HCR v3.0 with split-initiator probes (23-25 probe pairs per channel) in cultured human cells (Table S25) and whole-mount chicken embryos (Table S26) are ≈84%, compared with ≈50% using in situ HCR v2.0 (39 standard probe pairs per channel) in a previous study in whole-mount zebrafish embryos (Shah *et al.*, 2016).

Single molecules were identified in each channel using the following dot detection algorithm applied to a three-dimensional confocal image stack:

- **Step 1: Blur noise.** To remove noise smaller than the dots of interest, the image was convolved with an isotropic Gaussian blur (standard deviation $\sigma_{\text{blur}}$).

- **Step 2: Local background subtraction.** To eliminate variations in pixel intensity arising from background variations that occur on a length scale larger than the dots of interest, local background subtraction was performed by subtracting the mean pixel intensity of a cube (edge length $d_{\text{back}}$) from the intensity of the pixel at the center of the cube.

- **Step 3: Global threshold on pixel intensity.** To eliminate dim features, the resulting pixel intensities were subjected to a global threshold ($t_{\text{pixel}}$), and the range [$t_{\text{pixel}}$,1] was renormalized to a [0,1] scale.

- **Step 4: Watershed dot detection.** To identify single mRNA molecules as dots within the image, regional image maxima were segmented using the minimum saliency watershed method (Couprie & Bertrand, 1997; Yoo *et al.*, 2002). In this method, two maxima are labeled as the same dot if the minimum boundary height between the maxima is less than a given threshold ($t_{\text{watershed}}$). Dot coordinates were estimated as the intensity-weighted centroid of each watershed basin, and dot intensities were estimated as the integrated pixel intensity within each basin.

- **Step 5: Global threshold on dot intensity.** To eliminate dim dots, the resulting dot intensities were normalized on a [0,1] scale and a global threshold ($t_{\text{dot}}$) was applied. The resulting number of dots $N_i$ was recorded for channel $i$.

After identifying the dots in each channel of a 2-channel redundant detection image, a dot $i$ in Channel 1 and a dot $j$ in Channel 2 were considered colocalized if all of the following four statements were true:

- **Test 1:** The $xy$ centroids differed by less than a lateral distance threshold ($r_{xy} = 0.22$ $\mu$m).

- **Test 2:** Dot $i$ is the closest dot in Channel 1 to dot $j$ in Channel 2.

- **Test 3:** Dot $j$ is the closest dot in Channel 2 to dot $i$ in Channel 1.

- **Test 4:** The $z$ centroids differed by less than the axial distance threshold ($r_z = 0.42$ $\mu$m).

Note that due to the lower axial resolution, dot colocalization was tested separately for $xy$ and $z$. The same distance thresholds were used across all sample types and replicates.†

| | | HEK cells | | Chicken embryos | | Zebrafish embryos | |
|---|---|---|---|---|---|---|---|
| lateral resolution $d_{xy}$ ($\mu$m) | | 0.0624 | | 0.099 | | 0.2167 | |
| axial resolution $d_z$ ($\mu$m) | | 0.42 | | 0.42 | | 0.3369 | |
| Step 1: Gaussian blur radius $\sigma_{\text{blur}}$ ($\mu$m) | | 0.2 | | 0.2 | | 0.1 | |
| Step 2: Mean subtraction cube length $d_{\text{back}}$ ($\mu$m) | | 2.5 | | 3.0 | | 5.5 | |
| | Replicate | Ch1 | Ch2 | Ch1 | Ch2 | Ch1 | Ch2 |
| Step 3: Global pixel threshold ($t_{\text{pixel}}$) | 1 | 0.003 | 0.012 | 0.04 | 0.02 | 0.02 | 0.02 |
| | 2 | 0.005 | 0.012 | 0.04 | 0.02 | 0.01 | 0.003 |
| | 3 | 0.004 | 0.01 | 0.04 | 0.02 | 0.01 | 0.005 |
| Step 4: Watershed saliency minimum ($t_{\text{watershed}}$) | 1 | 0.15 | 0.15 | 0.15 | 0.15 | 0.15 | 0.15 |
| | 2 | 0.15 | 0.15 | 0.15 | 0.15 | 0.30 | 0.20 |
| | 3 | 0.15 | 0.15 | 0.15 | 0.15 | 0.35 | 0.30 |
| Step 5: Global dot intensity threshold ($t_{\text{dot}}$) | 1 | 0.01 | 0.005 | 0.00 | 0.01 | 0.01 | 0.01 |
| | 2 | 0.02 | 0.02 | 0.006 | 0.013 | 0.025 | 0.025 |
| | 3 | 0.03 | 0.012 | 0.01 | 0.025 | 0.04 | 0.04 |

**Table S6.** Parameters used for dot detection in dHCR images.

†For chicken embryo replicate 3, Channels 1 and 2 were manually aligned by applying a constant offset of $+0.35\mu$m to the $z$-coordinates of Channel 1 after the detected dots showed a clear bias in $z$ coordinates between the two channels. No other images were manually aligned.

#### S1.7 Flow cytometry data analysis

Data analysis for flow cytometry experiments on cultured cells closely follows the image analysis of Section S1.6 as detailed below.

##### S1.7.1 Raw cell intensities

The components of background (AF, NSA, NSD) and signal (SIG) are defined as before (Section S1.6.1), where $n$ is treated as an index over cells, and $i = 1$ for each cell since the flow cytometer returns one value per cell.

##### S1.7.2 Measurement of signal, background, signal-to-background, background components, and split-initiator HCR suppression for transgenic targets

For a transgenic target mRNA, signal and background are characterized based on flow cytometry experiments of Types 1a and 1b (Table S7A) with SIG+BACK measured in transgenic cells containing the target and BACK measured in wildtype (WT) cells lacking the target. This approach parallels that for characterizing signal and background in images (Section S1.6.2) with transgenic cells taking the place of a region of high expression and WT cells taking the place of a region of no/low expression. If desired, additional control experiments of Types 2 and 3 (Table S7B) can be performed to characterize the components of background (AF, NSA, NSD) using the calculations of Section S1.6.3. Likewise, additional control experiments of Types 4a, 4b, 5a, and 5b (Table S7C) can be performed to characterize split-initiator HCR suppression using the calculations of Section S1.6.4. For transgenic target mRNAs in human and bacterial cells, Figures S20 and S22 display distributions of cell intensities characterizing signal and background (panel A) and split-initiator HCR suppression (panel B). Tables S17 and S19 display corresponding values for SIG, BACK, and SIG/BACK (panel A), background components AF, NSA, and NSD (panel B), and SIG/SIG$^{\text{odd}}$ and SIG/SIG$^{\text{even}}$ (panel C).

| | Experiment type | Quantity | Reagents | | Cell type |
|---|---|---|---|---|---|
| | | | Probes | Hairpins | |
| **A** | 1a | SIG+NSD+NSA+AF = SIG+BACK | odd + even | ✓ | transgenic |
| | 1b | NSD+NSA+AF = BACK | odd + even | ✓ | WT |
| **B** | 2 | NSA+AF | | ✓ | transgenic |
| | 3 | AF | | | transgenic |
| **C** | 4a | SIG$^{\text{odd}}$+NSD$^{\text{odd}}$+NSA+AF = SIG$^{\text{odd}}$+BACK$^{\text{odd}}$ | odd | ✓ | transgenic |
| | 4b | NSD$^{\text{odd}}$+NSA+AF = BACK$^{\text{odd}}$ | odd | ✓ | WT |
| | 5a | SIG$^{\text{even}}$+NSD$^{\text{even}}$+NSA+AF = SIG$^{\text{even}}$+BACK$^{\text{even}}$ | even | ✓ | transgenic |
| | 5b | NSD$^{\text{even}}$+NSA+AF = BACK$^{\text{even}}$ | even | ✓ | WT |

**Table S7. Experiment types for flow cytometry using in situ HCR v3.0 with a transgenic target mRNA.** (A) Characterize signal, background, and signal-to-background. (B) Characterize components of background (AF, NSA, NSD). (C) Characterize split-initiator HCR suppression.

##### S1.7.3 Measurement of signal, background, signal-to-background, background components, and split-initiator HCR suppression for endogenous targets

For an endogenous target mRNA, signal and background are characterized based on flow cytometry experiments of Types 1a and 1b (Table S8A) with SIG+BACK measured using a probe set (odd + even) that address the target in WT cells and BACK measured using a probe set (Tg(odd) and Tg(even)) that addresses a different transgenic target absent from WT cells. Use of a previously validated transgenic probe set to measure background in WT cells ensures that a low measured fluorescence value does not simply indicate a dysfunctional probe set, but indeed represents low background generated by a probe set that is known to be functional if the target is present in the sample. If desired,

| | Experiment type | Quantity | Reagents: Probes | Reagents: Hairpins | Cell type |
|---|---|---|---|---|---|
| **A** | 1a | SIG+NSD+NSA+AF = SIG+BACK | odd + even | ✓ | WT |
| | 1b | NSD+NSA+AF = BACK | Tg(odd) + Tg(even) | ✓ | WT |
| **B** | 2 | NSA+AF | | ✓ | WT |
| | 3 | AF | | | WT |
| **C** | 4a | $SIG^{odd}$+$NSD^{odd}$+NSA+AF = $SIG^{odd}$+$BACK^{odd}$ | odd | ✓ | WT |
| | 4b | $NSD^{odd}$+NSA+AF = $BACK^{odd}$ | Tg(odd) | ✓ | WT |
| | 5a | $SIG^{even}$+$NSD^{even}$+NSA+AF = $SIG^{even}$+$BACK^{even}$ | even | ✓ | WT |
| | 5b | $NSD^{even}$+NSA+AF = $BACK^{even}$ | Tg(even) | ✓ | WT |

**Table S8. Experiment types for flow cytometry using in situ HCR v3.0 with an endogenous target mRNA.** (A) Characterize signal, background, and signal-to-background. (B) Characterize components of background (AF, NSA, NSD). (C) Characterize split-initiator HCR suppression. Here, Tg(odd) and Tg(even) denote odd and even probes from a probe set targeting a transgenic mRNA that is absent from WT cells.

additional control experiments of Types 2 and 3 (Table S8B) can be performed to characterize the components of background (AF, NSA, NSD) using the calculations of Section S1.6.3. Likewise, additional control experiments of Types 4a, 4b, 5a, and 5b (Table S8C) can be performed to characterize split-initiator HCR suppression using the calculations of Section S1.6.4. For endogenous target mRNA *GAPDH*, Figure S21 displays distributions of cell intensities characterizing signal and background (panel A) and split-initiator HCR suppression (panel B). Table S18 displays corresponding values for SIG, BACK, and SIG/BACK (panel A), background components AF, NSA, and NSD (panel B), and SIG/$SIG^{odd}$ and SIG/$SIG^{even}$ (panel C).

##### S1.7.4 Normalized single-cell intensities for mRNA relative quantitation

For relative mRNA quantitation between cells, the single-cell intensities within a channel are normalized using equation (S20) with BOT (mean BACK intensity across cells) and TOP (maximum SIG+BACK intensity for a single cell) defined by equations (S21) and (S22). Redundant detection experiments validating mRNA single-cell relative quantitation are displayed for endogenous targets *GAPDH* (Figure S24 and Table S20) and *PGK1* (Figure S25 and Table S21) in HEK cells, and for endogenous target *fusA* in *E. coli* (Figure S26 and Table S22).

### S2 Protocols for in situ HCR v3.0

#### S2.1 Protocols for whole-mount chicken embryos

##### S2.1.1 Preparation of fixed whole-mount chicken embryos

1. Collect chicken embryos on 3M filter paper and place in a petri dish containing Ringer's solution.

2. Transfer embryos into a new petri dish with fresh Ringer's solution.
   NOTE: *This is to rinse away egg yolk before fixation.*

3. Transfer into a petri dish containing 4% paraformaldehyde (PFA).
   CAUTION: *Use PFA with extreme care as it is a hazardous material.*
   NOTE: *Use fresh PFA and cool to 4 °C before use to avoid increased autofluorescence.*

4. Fix the samples at room temperature for 1 h.

5. Transfer embryos into a petri dish containing PBST.

6. Dissect the embryos off the filter paper. Remove the fixed vitelline membrane (opaque) and cut the squares around area pellucida, without leaving excess of extra-embryonic tissue.

7. Transfer embryos into a 2 mL Eppendorf tube containing PBST.

8. Nutate for 5 mins with the tube positioned horizontally in a small ice bucket.

9. Wash embryos two additional times with 2 mL of PBST, each time nutating for 5 min on ice.

10. Dehydrate embryos with 2 $\times$ 5 min washes of 2 mL methanol (MeOH) on ice.

11. Store embryos at -20 °C overnight before use.
    NOTE: *Embryos can be stored for six months at -20 °C.*

12. Transfer the required number of embryos for an experiment to a 2 mL Eppendorf tube.
    NOTE: *Do not place more than 4 embryos in each 2 mL Eppendorf tube.*

13. Rehydrate with a series of graded 2 mL MeOH/PBST washes for 5 min each on ice:
    (a) 75% MeOH / 25% PBST
    (b) 50% MeOH / 50% PBST
    (c) 25% MeOH / 75% PBST
    (d) 100% PBST
    (e) 100% PBST.

14. Treat embryos with 2 mL of 10 $\mu$g/mL proteinase K solution for 2 min (stage HH 8) or 2.5 min (stages HH 10–11) at room temperature.
    NOTE: *Proteinase K concentration and treatment time should be reoptimized for each batch of proteinase K, or for samples at a different developmental stage.*

15. Postfix with 2 mL of 4% PFA for 20 min at room temperature.

16. Wash embryos 2 $\times$ 5 min with 2 mL of PBST on ice.

17. Wash embryos with 2 mL of 50% PBST / 50% 5$\times$ SSCT for 5 min on ice.

18. Wash embryos with 2 mL of 5$\times$ SSCT for 5 min on ice.

##### S2.1.2 Buffer recipes for sample preparation

| **Ringer's solution** | For 2 L of solution |
| --- | --- |
| 123 mM NaCl | 14.4 g of NaCl |
| 1.53 mM $CaCl_2$ | 340 mg of $CaCl_2$ |
| 4.96 mM $KCl_2$ | 740 mg of KCl |
| 0.81 mM $Na_2HPO_4$ | 230 mg of $Na_2HPO_4$ |
| 0.15 mM $KH_2PO_4$ | 40 mg of $KH_2PO_4$ |
| | Bring volume up to 1.5 L with ultrapure $H_2O$ |
| | Adjust pH to 7.4 and fill up to 2 L with ultrapure $H_2O$ |
| | Filter sterilize with 0.22 $\mu$m bottle top filter |

| **4% Paraformaldehyde (PFA)** | For 25 mL of solution |
| --- | --- |
| 4% PFA | 1 g of PFA powder |
| 1× PBS | 25 mL of 1× PBS |
| | Heat solution at 50–60 °C to dissolve powder |

| **PBST** | For 50 mL of solution |
| --- | --- |
| 1× PBS | 5 mL of 10× PBS |
| 0.1% Tween 20 | 500 $\mu$L of 10% Tween 20 |
| | Fill up to 50 mL with ultrapure $H_2O$ |

| **Proteinase K solution** | For 1 mL of solution |
| --- | --- |
| 20 $\mu$g/mL proteinase K | 1 $\mu$L of 20 mg/mL proteinase K |
| | Fill up to 1 mL with PBST |

NOTE: *Avoid using calcium chloride and magnesium chloride in PBS as this leads to increased autofluorescence in the embryos.*

##### S2.1.3 Multiplexed in situ HCR v3.0 using split-initiator probes

###### Detection stage

1. For each sample, transfer 1-4 embryos to a 2 mL Eppendorf tube.
   NOTE: *Do not place more than 4 embryos in each 2 mL Eppendorf tube.*

2. Incubate embryos in 1 mL of 30% probe hybridization buffer on ice for 5 min.
   CAUTION: *probe hybridization buffer contains formamide, a hazardous material.*
   NOTE: *Pre-heat probe hybridization buffer to 37 °C before use.*

3. Remove the buffer and pre-hybridize with 1 mL of 30% probe hybridization buffer for 30 min at 37 °C.

4. Prepare probe solution by adding 4 pmol of each probe mixture (odd & even: 2 $\mu$L of 2 $\mu$M stock per probe mixture) to 1 mL of 30% probe hybridization buffer at 37 °C.
   NOTE: *For Figures 7B and S30 (smHCR), 10 pmol of each probe was used to improve probe hybridization efficiency.*

5. Remove the pre-hybridization solution and add the probe solution.

6. Incubate embryos overnight (12–16 h) at 37 °C.

7. Remove excess probes by washing embryos 4 $\times$ 15 min with 1 mL of 30% probe wash buffer at 37 °C:
   CAUTION: *probe wash buffer contains formamide, a hazardous material.*
   NOTE: *Pre-heat probe wash buffer to 37 °C before use.*

8. Wash samples 2 $\times$ 5 min with 5$\times$ SSCT at room temperature.

###### Amplification stage

1. Pre-amplify embryos with 500 mL of amplification buffer for 5 min at room temperature.
   NOTE: *Equilibrate amplification buffer to room temperature before use.*

2. Prepare 30 pmol of each fluorescently labeled hairpin by snap cooling 10 $\mu$L of 3 $\mu$M stock in hairpin storage buffer (heat at 95 °C for 90 seconds and cool to room temperature in a dark drawer for 30 min).

3. Prepare hairpin solution by adding all snap-cooled hairpins to 500 $\mu$L of amplification buffer at room temperature.

4. Remove the pre-amplification solution and add the hairpin solution.

5. Incubate the embryos overnight (12–16 h) in the dark at room temperature.
   NOTE: *For Figures 7B and S30 (smHCR), a 90 min amplification time was used to ensure single-molecule dots are diffraction limited.*

6. Remove excess hairpins by washing with 1 mL of 5$\times$ SSCT at room temperature:

   (a) 2 $\times$ 5 min
   (b) 2 $\times$ 30 min
   (c) 1 $\times$ 5 min

##### S2.1.4 Buffer recipes for in situ HCR v3.0

Probes, amplifiers, probe hybridization buffer, and probe wash buffer should be stored at -20 °C. Amplification buffer should be stored at 4 °C. Keep these reagents on ice at all times during probe and amplifier preparation. Make sure all solutions are well mixed before use.

| **30% probe hybridization buffer** | For 40 mL of solution |
|---|---|
| 30% formamide | 12 mL formamide |
| 5× sodium chloride sodium citrate (SSC) | 10 mL of 20× SSC |
| 9 mM citric acid (pH 6.0) | 360 $\mu$L 1 M citric acid, pH 6.0 |
| 0.1% Tween 20 | 400 $\mu$L of 10% Tween 20 |
| 50 $\mu$g/mL heparin | 200 $\mu$L of 10 mg/mL heparin |
| 1× Denhardt's solution | 800 $\mu$L of 50× Denhardt's solution |
| 10% dextran sulfate | 8 mL of 50% dextran sulfate |
| | Fill up to 40 mL with ultrapure $H_2O$ |

| **30% probe wash buffer** | For 40 mL of solution |
|---|---|
| 30% formamide | 12 mL formamide |
| 5× sodium chloride sodium citrate (SSC) | 10 mL of 20× SSC |
| 9 mM citric acid (pH 6.0) | 360 $\mu$L 1 M citric acid, pH 6.0 |
| 0.1% Tween 20 | 400 $\mu$L of 10% Tween 20 |
| 50 $\mu$g/mL heparin | 200 $\mu$L of 10 mg/mL heparin |
| | Fill up to 40 mL with ultrapure $H_2O$ |

| **Amplification buffer** | For 40 mL of solution |
|---|---|
| 5× sodium chloride sodium citrate (SSC) | 10 mL of 20× SSC |
| 0.1% Tween 20 | 400 $\mu$L of 10% Tween 20 |
| 10% dextran sulfate | 8 mL of 50% dextran sulfate |
| | Fill up to 40 mL with ultrapure $H_2O$ |

| **5× SSCT** | For 40 mL of solution |
|---|---|
| 5× sodium chloride sodium citrate (SSC) | 10 mL of 20× SSC |
| 0.1% Tween 20 | 400 $\mu$L of 10% Tween 20 |
| | Fill up to 40 mL with ultrapure $H_2O$ |

| **50% dextran sulfate** | For 40 mL of solution |
|---|---|
| 50% dextran sulfate | 20 g of dextran sulfate powder |
| | Fill up to 40 mL with ultrapure $H_2O$ |

###### S2.1.5 Sample mounting for microscopy

1. A chamber for mounting each embryo was made by aligning two stacks of double-sided tape (2 pieces per stack) 1 cm apart on a 25 mm $\times$ 75 mm glass slide.

2. Place an embryo between the tape stacks on the slide and remove as much solution as possible.

3. Align the embryo for dorsal imaging and carefully touch the slide with a kimwipe to further dry the area around the embryo.

4. Add two drops of SlowFade Gold antifade mountant on top of the embryo.

5. Place a 22 mm $\times$ 30 mm No. 1 coverslip on top of the stacks to close the chamber.
   NOTE: *See Section S1.4 for details of confocal microscopes used to image whole-mount chicken embryos.*

##### S2.1.6 Reagents and supplies

Paraformaldehyde (PFA) (Sigma Cat. # P6148)
Methanol (Mallinckrodt Chemicals Cat. # 3016-16)
Proteinase K, molecular biology grade (NEB Cat. # P8107S)
Formamide (Deionized) (Ambion Cat. # AM9342)
20× sodium chloride sodium citrate (SSC) (Life Technologies Cat. # 15557-044)
Heparin (Sigma Cat. # H3393)
50% Tween 20 (Life Technologies Cat. # 00-3005)
50× Denhardt's solution (Life Technologies Cat. # 750018)
Dextran sulfate, mol. wt. > 500,000 (Sigma Cat. # D6001)
25 mm × 75 mm glass slide (VWR Cat. # 48300-025)
22 mm × 30 mm No. 1 coverslip (VWR Cat. # 48393-026)
SlowFade Gold antifade mountant (Life Technologies Cat. # S36937)

#### S2.2 Protocols for mammalian cells on a chambered slide

##### S2.2.1 Preparation of fixed mammalian cells on a chambered slide

1. Coat bottom of each chamber by applying 300 $\mu$L of 0.01% poly-D-lysine prepared in cell culture grade $H_2O$.
   NOTE: *A volume of 300 $\mu$L is sufficient per chamber on an 8-chamber slide.*

2. Incubate for at least 30 min at room temperature.

3. Aspirate the coating solution and wash each chamber twice with molecular biology grade $H_2O$.

4. Plate desired number of cells in each chamber.

5. Grow cells to desired confluency for 24–48 h.

6. Aspirate growth media and wash each chamber with 300 $\mu$L of DPBS.
   NOTE: *Avoid using calcium chloride and magnesium chloride in DPBS as this leads to increased autofluorescence.*

7. Add 300 $\mu$L of 4% formaldehyde to each chamber.
   CAUTION: *Use formaldehyde with extreme care as it is a hazardous material.*

8. Incubate for 10 min at room temperature.

9. Aspirate fixative and wash each chamber 2 $\times$ 300 $\mu$L of DPBS.

10. Aspirate DPBS and add 300 $\mu$L of ice-cold 70% ethanol.

11. Permeabilize cells overnight at -20 °C.

12. Cells can be stored at -20 °C or 4 °C until use.

##### S2.2.2 Buffer recipes for sample preparation

| **4% formaldehyde in PBS** | For 10 mL of solution |
|---|---|
| 4% formaldehyde | 2.5 mL of 16% formaldehyde |
| 1× PBS | 1 mL of 10× PBS |
| | Fill up to 10 mL with molecular biology grade $H_2O$ |

NOTE: *Avoid using calcium chloride and magnesium chloride in PBS as this leads to increased autofluorescence in the samples.*

##### S2.2.3 Multiplexed in situ HCR v3.0 using split-initiator probes

**Detection stage**

1. Aspirate EtOH and air dry samples at room temperature.
   NOTE: *Drying of sample is optional.*

2. Wash samples two times with 300 $\mu$L of 2$\times$ SSC.

3. Pre-hybridize samples in 300 $\mu$L of 30% probe hybridization buffer for 30 min at 37 °C.
   CAUTION: *probe hybridization buffer contains formamide, a hazardous material.*
   NOTE: *Pre-heat probe hybridization buffer to 37 °C before use.*

4. Prepare probe solution by adding 1.2 pmol of each probe mixture (odd & even: 0.6 $\mu$L of 2 $\mu$M stock per probe mixture) to 300 $\mu$L of 30% probe hybridization buffer at 37 °C.
   NOTE: *For Figures 7A and S29 (smHCR), 3 pmol of each probe was used to improve probe hybridization efficiency.*

5. Remove the pre-hybridization solution and add the probe solution.

6. Incubate samples overnight (12–16 h) at 37 °C.

7. Remove excess probes by washing 4 $\times$ 5 min with 300 $\mu$L of 30% probe wash buffer at 37 °C.
   CAUTION: *probe wash buffer contains formamide, a hazardous material.*
   NOTE: *Pre-heat probe wash buffer to 37 °C before use.*

8. Wash samples 2 $\times$ 5 min with 5$\times$ SSCT at room temperature.

**Amplification stage**

1. Pre-amplify samples in 300 $\mu$L of amplification buffer for 30 min at room temperature.
   NOTE: *Equilibrate amplification buffer to room temperature before use.*

2. Prepare 18 pmol of each fluorescently labeled hairpin by snap cooling 6 $\mu$L of 3 $\mu$M stock in hairpin storage buffer (heat at 95 °C for 90 seconds and cool to room temperature in a dark drawer for 30 min).

3. Prepare hairpin solution by adding all snap-cooled hairpins to 300 $\mu$L of amplification buffer at room temperature.

4. Remove the pre-amplification solution and add the hairpin solution.

5. Incubate samples overnight (12–16 h) in the dark at room temperature.
   NOTE: *For Figures 7A and S29 (smHCR), a 45 min amplification time was used to ensure single-molecule dots are diffraction-limited.*

6. Remove excess hairpins by washing 5 $\times$ 5 min with 300 $\mu$L of 5$\times$ SSCT at room temperature.

7. Aspirate 5$\times$ SSCT and add $\approx$100 $\mu$L of SlowFade Gold antifade mountant with DAPI.

8. Samples can be stored at 4 °C protected from light prior to imaging.

##### S2.2.4 Buffer recipes for in situ HCR v3.0

Probes, amplifiers, probe hybridization buffer, and probe wash buffer should be stored at -20 °C. Amplification buffer should be stored at 4 °C. Keep these reagents on ice at all times during probe and amplifier preparation. Make sure all solutions are well mixed before use.

| **30% probe hybridization buffer** | For 40 mL of solution |
|---|---|
| 30% formamide | 12 mL formamide |
| 5× sodium chloride sodium citrate (SSC) | 10 mL of 20× SSC |
| 9 mM citric acid (pH 6.0) | 360 $\mu$L 1 M citric acid, pH 6.0 |
| 0.1% Tween 20 | 400 $\mu$L of 10% Tween 20 |
| 50 $\mu$g/mL heparin | 200 $\mu$L of 10 mg/mL heparin |
| 1× Denhardt's solution | 800 $\mu$L of 50× Denhardt's solution |
| 10% dextran sulfate | 8 mL of 50% dextran sulfate |
| | Fill up to 40 mL with ultrapure $H_2O$ |

| **30% probe wash buffer** | For 40 mL of solution |
|---|---|
| 30% formamide | 12 mL formamide |
| 5× sodium chloride sodium citrate (SSC) | 10 mL of 20× SSC |
| 9 mM citric acid (pH 6.0) | 360 $\mu$L 1 M citric acid, pH 6.0 |
| 0.1% Tween 20 | 400 $\mu$L of 10% Tween 20 |
| 50 $\mu$g/mL heparin | 200 $\mu$L of 10 mg/mL heparin |
| | Fill up to 40 mL with ultrapure $H_2O$ |

| **Amplification buffer** | For 40 mL of solution |
|---|---|
| 5× sodium chloride sodium citrate (SSC) | 10 mL of 20× SSC |
| 0.1% Tween 20 | 400 $\mu$L of 10% Tween 20 |
| 10% dextran sulfate | 8 mL of 50% dextran sulfate |
| | Fill up to 40 mL with ultrapure $H_2O$ |

| **5× SSCT** | For 40 mL of solution |
|---|---|
| 5× sodium chloride sodium citrate (SSC) | 10 mL of 20× SSC |
| 0.1% Tween 20 | 400 $\mu$L of 10% Tween 20 |
| | Fill up to 40 mL with ultrapure $H_2O$ |

| **50% dextran sulfate** | For 40 mL of solution |
|---|---|
| 50% dextran sulfate | 20 g of dextran sulfate powder |
| | Fill up to 40 mL with ultrapure $H_2O$ |

##### S2.2.5 Reagents and supplies

Molecular biology grade $H_2O$ (Corning Cat. # 46-000-CV)
16% Formaldehyde (w/v), Methanol-free (Life Technologies Cat. # 28906)
DPBS, no calcium, no magnesium (Life Technologies Cat. # 14190144)
10× PBS (Ambion Cat. # AM9624)
Formamide (Deionized) (Ambion Cat. # AM9342)
20× sodium chloride sodium citrate (SSC) (Life Technologies Cat. # 15557-044)
Heparin (Sigma Cat. # H3393)
10% Tween 20 (BioRad Cat. # 161-0781)
50× Denhardt's solution (Life Technologies Cat. # 750018)
Dextran sulfate, mol. wt. $>$ 500,000 (Sigma Cat. # D6001)
ibidi $\mu$-slide ibitreat (ibidi Cat. # 80826)
SlowFade Gold antifade mountant with DAPI (Life Technologies Cat. # S36938)

#### S2.3 Protocols for mammalian cells in suspension

##### S2.3.1 Preparation of fixed mammalian cells in suspension

1. Aspirate growth media from culture plate and wash cells with DPBS.
   NOTE: *Avoid using calcium chloride and magnesium chloride in DPBS as this leads to increased autofluorescence.*

2. Add 3 mL of trypsin per 10 cm plate and incubate in a 5% $CO_2$ incubator for 5 min at 37 °C.

3. Quench trypsin by adding 3 mL of growth media.

4. Transfer cells to a conical tube and centrifuge for 5 min at 180 $\times$ g.

5. Aspirate supernatant and re-suspend cells in 4% formaldehyde to reach $\approx 10^6$ cells/mL.
   CAUTION: *Use formaldehyde with extreme care as it is a hazardous material.*

6. Fix cells for 1 hr at room temperature.

7. Centrifuge for 5 min at 180 $\times$ g and remove supernatant.

8. Wash cells 4 times with PBST (use the same volume as formaldehyde solution).
   NOTE: *Centrifuge for 5 min at 180 $\times$ g and aspirate supernatant between washes.*

9. Re-suspend cells in ice-cold 70% ethanol (use the same volume as formaldehyde and PBST solutions).

10. Permeabilize cells overnight at 4 °C.

11. Cells can be stored at 4 °C until use.

##### S2.3.2 Buffer recipes for sample preparation

| **4% formaldehyde in PBST** | For 36 mL of solution |
| --- | --- |
| 4% formaldehyde | 9 mL of 16% formaldehyde |
| 1× PBS, 0.1% Tween 20 | 3.6 mL of 10× PBST |
| | 180 $\mu$L of 10% Tween 20 |
| | Fill up to 36 mL with ultrapure $H_2O$ |

NOTE: *Avoid using calcium chloride and magnesium chloride in PBS as this leads to increased autofluorescence in the samples.*

##### S2.3.3 Multiplexed in situ HCR v3.0 using split-initiator probes

**Detection stage**

1. Transfer desired amount (0.5-1 $\times$ $10^6$) of fixed cells into a 1.5 mL Eppendorf tube.
2. Centrifuge for 5 min to remove EtOH.
   NOTE: *All centrifugation steps are done at 180 $\times$ g.*
3. Wash cells twice with 500 $\mu$L of PBST. Centrifuge for 5 min to remove supernatant.
4. Re-suspend the pellet with 400 $\mu$L of 30% probe hybridization buffer and pre-hybridize for 30 min at 37 °C.
   CAUTION: *probe hybridization buffer contains formamide, a hazardous material.*
   NOTE: *Pre-heat probe hybridization buffer to 37 °C before use.*
5. In the meantime, prepare probe solution by adding 2 pmol of each probe mixture (odd & even: 1 $\mu$L of 2 $\mu$M stock per probe mixture) to 100 $\mu$L of 30% probe hybridization buffer at 37 °C.
6. Add the probe solution directly to the sample to reach a final probe concentration of 4 nM.
7. Incubate the sample overnight at 37 °C.
8. Centrifuge for 5 min to remove probe solution.
9. Re-suspend the cell pellet with 500 $\mu$L of 30% probe wash buffer.
   CAUTION: *probe wash buffer contains formamide, a hazardous material.*
   NOTE: *Pre-heat probe wash buffer to 37 °C before use.*
10. Incubate for 10 min at 37 °C and remove the wash solution by centrifugation for 5 min.
11. Repeat steps 9 and 10 for three additional times.
12. Re-suspend the cell pellet with 500 $\mu$L of 5$\times$ SSCT.
13. Incubate for 5 min at room temperature.
14. Proceed to hairpin amplification.

**Amplification stage**

1. Centrifuge for 5 min to pellet the cells.
2. Re-suspend the cell pellet with 150 $\mu$L of amplification buffer and pre-amplify for 30 min at room temperature.
   NOTE: *Equilibrate amplification buffer to room temperature before use.*
3. Prepare 15 pmol of each fluorescently labeled hairpin by snap cooling 5 $\mu$L of 3 $\mu$M stock in hairpin storage buffer (heat at 95 °C for 90 seconds and cool to room temperature in a dark drawer for 30 min).
4. Prepare hairpin mixture by adding all snap-cooled hairpins to 100 $\mu$L of amplification buffer at room temperature.
5. Add the hairpin mixture directly to the sample to reach a final hairpin concentration of 60 nM.
6. Incubate the sample overnight ($>$12 h) in the dark at room temperature.
7. Centrifuge for 5 min and remove the hairpin solution.
8. Re-suspend the cell pellet with 500 $\mu$L of 5$\times$ SSCT.

9. Without incubation, remove the wash solution by centrifugation for 5 min.

10. Repeat steps 8 and 9 for five additional times.

11. Re-suspend the cell pellet in desired buffer and volume.
    NOTE: *Samples can be stored at 4 °C protected from light before flow cytometry or imaging.*

12. Filter cells before flow cytometry.

##### S2.3.4 Buffer recipes for in situ HCR v3.0

Probes, amplifiers, probe hybridization buffer, and probe wash buffer should be stored at -20 °C. Amplification buffer should be stored at 4 °C. Keep these reagents on ice at all times during probe and amplifier preparation. Make sure all solutions are well mixed before use.

| **30% probe hybridization buffer (low MW D. S.)** | For 40 mL of solution |
|---|---|
| 30% formamide | 12 mL formamide |
| 5× sodium chloride sodium citrate (SSC) | 10 mL of 20× SSC |
| 9 mM citric acid (pH 6.0) | 360 $\mu$L 1 M citric acid, pH 6.0 |
| 0.1% Tween 20 | 400 $\mu$L of 10% Tween 20 |
| 50 $\mu$g/mL heparin | 200 $\mu$L of 10 mg/mL heparin |
| 1× Denhardt's solution | 800 $\mu$L of 50× Denhardt's solution |
| 10% low MW dextran sulfate | 8 mL of 50% low MW dextran sulfate |
| | Fill up to 40 mL with ultrapure $H_2O$ |

| **30% probe wash buffer** | For 40 mL of solution |
|---|---|
| 30% formamide | 12 mL formamide |
| 5× sodium chloride sodium citrate (SSC) | 10 mL of 20× SSC |
| 9 mM citric acid (pH 6.0) | 360 $\mu$L 1 M citric acid, pH 6.0 |
| 0.1% Tween 20 | 400 $\mu$L of 10% Tween 20 |
| 50 $\mu$g/mL heparin | 200 $\mu$L of 10 mg/mL heparin |
| | Fill up to 40 mL with ultrapure $H_2O$ |

| **Amplification buffer (low MW D. S.)** | For 40 mL of solution |
|---|---|
| 5× sodium chloride sodium citrate (SSC) | 10 mL of 20× SSC |
| 0.1% Tween 20 | 400 $\mu$L of 10% Tween 20 |
| 10% low MW dextran sulfate | 8 mL of 50% low MW dextran sulfate |
| | Fill up to 40 mL with ultrapure $H_2O$ |

| **5× SSCT** | For 40 mL of solution |
|---|---|
| 5× sodium chloride sodium citrate (SSC) | 10 mL of 20× SSC |
| 0.1% Tween 20 | 400 $\mu$L of 10% Tween 20 |
| | fill up to 40 mL with ultrapure $H_2O$ |

| **1× PBST** | For 40 mL of solution |
|---|---|
| 1× PBS | 10 mL of 10× PBST (0.5% Tween 20) |
| 0.1% Tween 20 | 200 $\mu$L of 10% Tween 20 |
| | Fill up to 40 mL with ultrapure $H_2O$ |

| **50% dextran sulfate** | For 40 mL of solution |
|---|---|
| 50% dextran sulfate | 20 g of low MW dextran sulfate powder |
| | Fill up to 40 mL with ultrapure $H_2O$ |

##### S2.3.5 Reagents and supplies

DPBS, no calcium, no magnesium (Life Technologies Cat. # 14190144)
Trypsin-EDTA (0.25%), phenol red (Life Technologies Cat. # 25200072)
16% Formaldehyde (w/v), methanol-free (Life Technologies Cat. # 28908)
10× PBST (Rockland Cat. # MB-075-1000)
10% Tween 20 solution (Bio-Rad Cat. # 161-0781)
Formamide (Deionized) (Ambion Cat. # AM9342)
20× sodium chloride sodium citrate (SSC) (Life Technologies Cat. # 15557-044)
Heparin (Sigma Cat. # H3393)
50% Tween 20 (Life Technologies Cat. # 00-3005)
50× Denhardt's solution (Life Technologies Cat. # 750018)
Dextran sulfate, mol. wt. 6,500-10,000 (Sigma Cat. # D4911)

#### S2.4 Protocols for bacteria in suspension

##### S2.4.1 Preparation of fixed bacteria in suspension

1. Grow *E. coli* from streaked plate or frozen glycerol stocks in 2–3 mL of LB media overnight in a 37°C shaker.

2. Dilute to make a 5 mL liquid culture with $OD_{600}$ = 0.05.

3. Incubate in a 37°C shaker until $OD_{600} \approx 0.5$ (exponential phase).

4. Aliquot 1 mL of cells and centrifuge for 10 min.
   NOTE: *Centrifugation should be a gentle as possible to pellet cells. For E. coli all centrifugation steps are done at 4000 × g.*

5. Remove supernatant and re-suspend cells in 750 μL 1 × phosphate-buffered saline (PBS).
   NOTE: *Remove all solutions via pipetting throughout the protcol.*

6. Add 250 μL of 4% formaldehyde to and incubate overnight at 4°C.
   CAUTION: *Use formaldehyde with extreme care as it is a hazardous material.*

7. Centrifuge for 10 min and remove supernatant.

8. Re-suspend cells in 150 μL 1 × PBS.

9. Add 850 μL of 100% MeOH and store at -20°C before use.

##### S2.4.2 Buffer recipes for sample preparation

**LB media**
5 g of Novagen LB Broth Miller powder
Fill up to 200 mL with ultrapure $H_2O$
Autoclave at 121°Cfor 20 min

**4% formaldehyde in PBS**
4% formaldehyde
1× PBS

For 4 mL of solution
1 mL of 16% formaldehyde
0.4 mL of 10× PBS
Fill up to 4 mL with ultrapure $H_2O$

NOTE: *Avoid using calcium chloride and magnesium chloride in PBS as this leads to increased autofluorescence in the samples.*

##### S2.4.3 Multiplexed in situ HCR v3.0 using split-initiator probes

**Detection stage**

1. Transfer 150 $\mu$L of fixed cells into a 1.5 mL Eppendorf tube.
2. Centrifuge for 5 min and remove supernatant.
3. Wash cells with 500 $\mu$L of 1× PBST. Centrifuge for 5 min to remove supernatant.
4. Re-suspend the pellet with 400 $\mu$L of 30% LMW probe hybridization buffer and pre-hybridize for 1 hr at 37 °C.
   CAUTION: *probe hybridization buffer contains formamide, a hazardous material.*
   NOTE: *Pre-heat probe hybridization buffer to 37 °C before use.*
5. In the meantime, prepare probe solution by adding 2 pmol of each probe mixture (odd & even: 1 $\mu$L of 2 $\mu$M stock per probe mixture) to 100 $\mu$L of LMW 30% probe hybridization buffer at 37 °C.
6. Add the probe solution directly to the sample to reach a final probe concentration of 4 nM.
7. Incubate the sample overnight at 37 °C.
8. Add 1mL of probe wash buffer to the sample.
   CAUTION: *probe wash buffer contains formamide, a hazardous material.*
   NOTE: *Pre-heat probe wash buffer to 37 °C before use.*
9. Centrifuge for 5 min and remove the wash solution.
10. Re-suspend the cell pellet with 500 $\mu$L wash solution.
11. Incubate for 5 min at 37 °C and remove the wash solution by centrifugation for 5 min.
12. Repeat steps 10 and 11 for two additional times but with 10 min incubation.
13. Proceed to hairpin amplification.

**Amplification stage**

1. Re-suspend the cell pellet with 150 $\mu$L of LMW amplification buffer and pre-amplify for 30 min at room temperature.
   NOTE: *Equilibrate amplification buffer to room temperature before use.*
2. Prepare 15 pmol of each fluorescently labeled hairpin by snap cooling 5 $\mu$L of 3 $\mu$M stock in hairpin storage buffer (heat at 95 °C for 90 seconds and cool to room temperature in a dark drawer for 30 min).
3. Prepare hairpin mixture by adding all snap-cooled hairpins to 100 $\mu$L of LMW amplification buffer at room temperature.
4. Add the hairpin mixture directly to the sample to reach a final hairpin concentration of 60 nM.
5. Incubate the sample overnight (>12 h) in the dark at room temperature.
6. Add 1 mL of 5× SSCT at room temperature to the sample to dilute the solution.
7. Centrifuge for 5 min and remove the hairpin solution.
8. Re-suspend the cell pellet with 500 $\mu$L of 5× SSCT and incubate 5 min at room temperature.
9. Centrifuge for 5 min and remove the wash solution.

10. Repeat steps 8 and 9 for two additional times but with a 10 min incubation.
11. Re-suspend the cell pellet in desired buffer and volume.
    NOTE: *Samples can be stored at 4 °C protected from light before flow cytometry.*
12. Filter cells before flow cytometry.

##### S2.4.4 Buffer recipes for in situ HCR v3.0

Probes, amplifiers, probe hybridization buffer, and probe wash buffer should be stored at -20 °C. Amplification buffer should be stored at 4 °C. Make sure all solutions are well mixed before use.

| **30% probe hybridization buffer (low MW D. S.)** | For 40 mL of solution |
|---|---|
| 30% formamide | 12 mL formamide |
| 5× sodium chloride sodium citrate (SSC) | 10 mL of 20× SSC |
| 9 mM citric acid (pH 6.0) | 360 $\mu$L 1 M citric acid, pH 6.0 |
| 0.1% Tween 20 | 400 $\mu$L of 10% Tween 20 |
| 50 $\mu$g/mL heparin | 200 $\mu$L of 10 mg/mL heparin |
| 1× Denhardt's solution | 800 $\mu$L of 50× Denhardt's solution |
| 10% low MW dextran sulfate | 8 mL of 50% low MW dextran sulfate |
| | Fill up to 40 mL with ultrapure $H_2O$ |

| **30% probe wash buffer** | For 40 mL of solution |
|---|---|
| 30% formamide | 12 mL formamide |
| 5× sodium chloride sodium citrate (SSC) | 10 mL of 20× SSC |
| 9 mM citric acid (pH 6.0) | 360 $\mu$L 1 M citric acid, pH 6.0 |
| 0.1% Tween 20 | 400 $\mu$L of 10% Tween 20 |
| 50 $\mu$g/mL heparin | 200 $\mu$L of 10 mg/mL heparin |
| | Fill up to 40 mL with ultrapure $H_2O$ |

| **Amplification buffer (low MW D. S.)** | For 40 mL of solution |
|---|---|
| 5× sodium chloride sodium citrate (SSC) | 10 mL of 20× SSC |
| 0.1% Tween 20 | 400 $\mu$L of 10% Tween 20 |
| 10% low MW dextran sulfate | 8 mL of 50% low MW dextran sulfate |
| | Fill up to 40 mL with ultrapure $H_2O$ |

| **5× SSCT** | For 40 mL of solution |
|---|---|
| 5× sodium chloride sodium citrate (SSC) | 10 mL of 20× SSC |
| 0.1% Tween 20 | 400 $\mu$L of 10% Tween 20 |
| | fill up to 40 mL with ultrapure $H_2O$ |

| **1× PBST** | For 40 mL of solution |
|---|---|
| 1× PBS | 10 mL of 10× PBST (0.5% Tween 20) |
| 0.1% Tween 20 | 200 $\mu$L of 10% Tween 20 |
| | Fill up to 40 mL with ultrapure $H_2O$ |

| **50% dextran sulfate** | For 40 mL of solution |
|---|---|
| 50% dextran sulfate | 20 g of low MW dextran sulfate powder |
| | Fill up to 40 mL with ultrapure $H_2O$ |

##### S2.4.5 Reagents and supplies

LB Broth Miller (Novagen Cat. # 71753-5)
16% Formaldehyde (w/v), methanol-free (Life Technologies Cat. # 28908)
10× PBS (Ambion Cat. # AM9624)
10× PBST (Rockland Cat. # MB-075-1000)
10% Tween 20 solution (Bio-Rad Cat. # 161-0781)
Formamide (Deionized) (Ambion Cat. # AM9342)
20× sodium chloride sodium citrate (SSC) (Life Technologies Cat. # 15557-044)
Heparin (Sigma Cat. # H3393)
50% Tween 20 (Life Technologies Cat. # 00-3005)
50× Denhardt's solution (Life Technologies Cat. # 750018)
Dextran sulfate, mol. wt. 6,500-10,000 (Sigma Cat. # D4911)

### S3 Additional studies

#### S3.1 Validation of split-initiator HCR suppression in vitro and in situ (cf. Figure 2)

Figures S3 and S4 display gel studies measuring split-initiator HCR suppression for amplifiers B1–B5, revealing typical ≈60-fold suppression. Table S9 displays signal-to-signal ratios for the same amplifiers used in situ within whole-mount chicken embryos (imaging) and/or within cultured human or bacterial cells (flow cytometry), revealing typical ≈50-fold suppression.

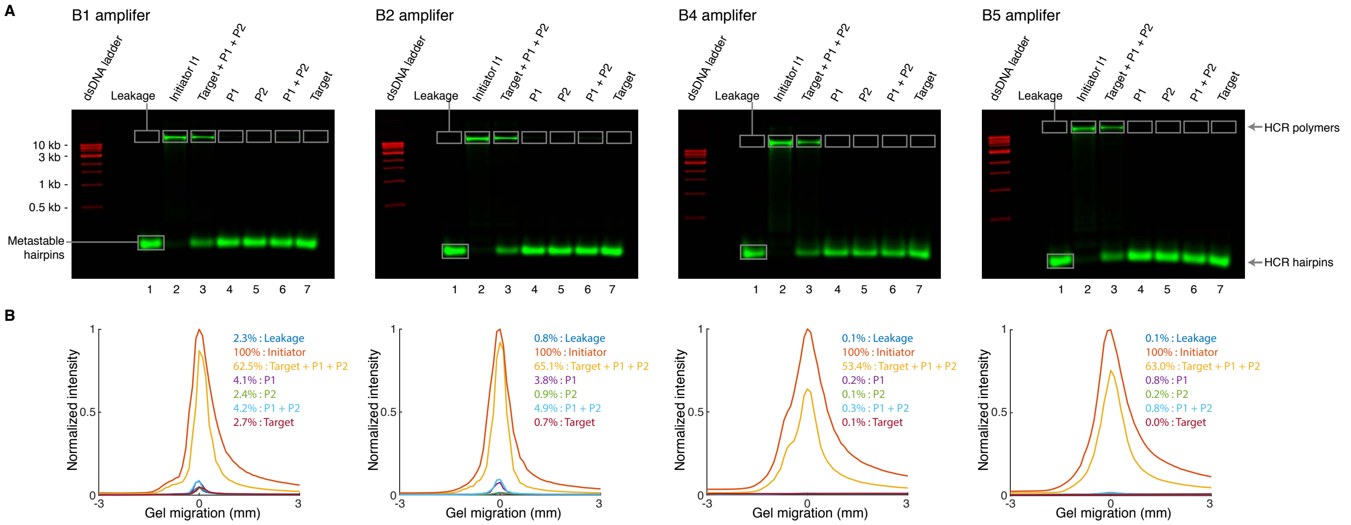

**Figure S3. Test tube validation of split-initiator HCR suppression for amplifiers B1, B2, B4, and B5 (cf. Figure 2).** (A) Agarose gel electrophoresis. Reaction conditions: hairpins H1 and H2 at 0.5 $\mu$M each (Lanes 1-7); DNA oligos I1, P1, P2, and/or Target at 5 nM each (lanes noted on the gel); 5× SSCT buffer; overnight reaction at room temperature. Hairpins H1 and H2 labeled with Alexa 647 fluorophore (green channel). dsDNA 1 kb ladder pre-stained with SYBR Gold (red channel). (B) Quantification of the polymer band in panel (A).

| Organism | Target | Samples | Channel | SIG/SIG$^{\text{odd}}$ | SIG/SIG$^{\text{even}}$ | Table |
|---|---|---|---|---|---|---|
| *G. gallus domesticus* | *EphA4* | 3 embryos | B2-Alexa647 | >800 | 57 ± 5 | S14 |
| *H. sapiens sapiens* | *Tg(d2eGFP)* | 55,000 cells | B3-Alexa594 | 461 ± 9 | 22.2 ± 0.1 | S17 |
| *H. sapiens sapiens* | *GAPDH* | 30,000 cells | B4-Alexa594 | 55 ± 5 | 42.9 ± 0.6 | S18 |
| *H. sapiens sapiens* | *GAPDH* | 20,000 cells | B5-Alexa488 | 40.4 ± 0.8 | 3229 ± 4 | S20 |
| *H. sapiens sapiens* | *GAPDH* | 20,000 cells | B4-Alexa594 | 67 ± 2 | 52 ± 2 | S20 |
| *H. sapiens sapiens* | *PGK1* | 54,000 cells | B1-Alexa488 | >5000 | 49.0 ± 0.5 | S21 |
| *H. sapiens sapiens* | *PGK1* | 54,000 cells | B2-Alexa594 | 42 ± 1 | 13 ± 1 | S21 |
| *H. sapiens sapiens* | *GAPDH* | 18,000 cells | B4-Alexa488 | 93 ± 6 | 91 ± 17 | S20 |
| *H. sapiens sapiens* | *PGK1* | 18,000 cells | B2-Alexa594 | 21 ± 2 | 18.8 ± 0.4 | S21 |
| *E. coli* | *Tg(rfp)* | 18,000 cells | B3-Alexa546 | >3000 | 9.9 ± 0.5 | S19 |
| *E. coli* | *fusA* | 3,400 cells | B3-Alexa488 | 40 ± 20 | 14 ± 3 | S22 |
| *E. coli* | *fusA)* | 3,400 cells | B2-Alexa594 | 50 ± 20 | 6.2 ± 0.6 | S22 |
| *E. coli* | *fusA* | 35,000 cells | B3-Alexa488 | 600 ± 300 | 17 ± 8 | S24 |
| *E. coli* | *icd)* | 35,000 cells | B1-Alexa594 | 800 ± 300 | 85 ± 1 | S24 |

**Table S9. In situ validation of split-initiator HCR suppression.**

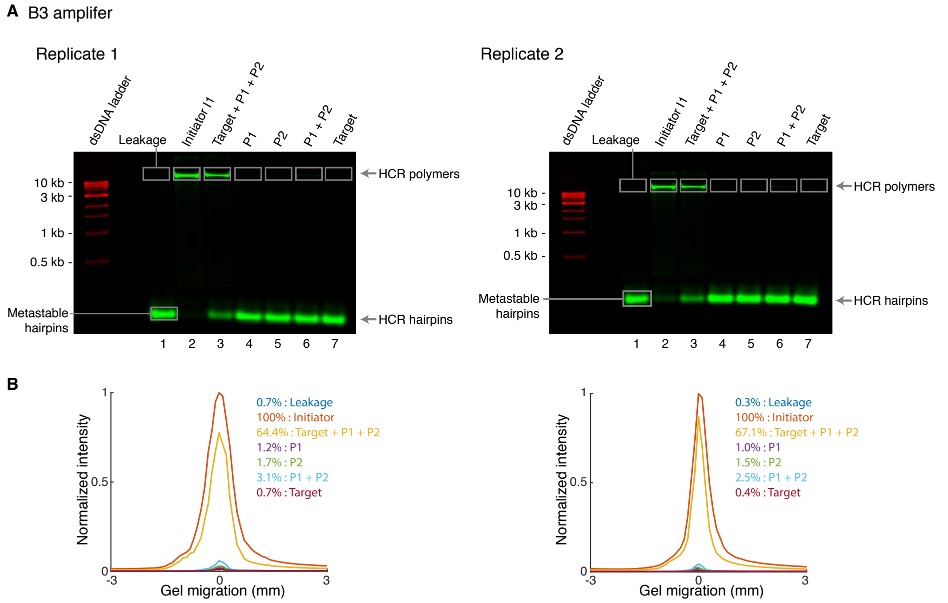

**Figure S4. Test tube validation of split-initiator HCR suppression for amplifier B3 (cf. Figure 2).** (A) Agarose gel electrophoresis (Replicate 1 is displayed in Figure 2). Reaction conditions: hairpins H1 and H2 at 0.5 $\mu$M each (Lanes 1-7); DNA oligos I1, P1, P2, and/or Target at 5 nM each (lanes noted on the gel); 5$\times$ SSCT buffer; overnight reaction at room temperature. Hairpins H1 and H2 labeled with Alexa 647 fluorophore (green channel). dsDNA 1 kb ladder pre-stained with SYBR Gold (red channel). (B) Quantification of the polymer band in panel (A).

#### S3.2 In situ validation of automatic background suppression with split-initiator probes in whole-mount chicken embryos (cf. Figure 3)

The following studies are included:

- **Measurement of background and signal-to-background for unoptimized standard and split-initiator probe sets as a function of probe set size.** Probe sets with 5, 10, or 20 standard probes or split-initiator probe pairs. For background comparisons (Figures S5 and S6 and Table S10), the PMT gain is held constant for all experiments to enable comparison of background intensities between experiments. For signal-to-background comparisons (Figures S7 and S8 and Table S11), the PMT gain is adjusted to use the full dynamic range for each probe set.
- **Measurement of background and signal-to-background for standard and split-initiator probes with identical target-binding domains.** For these studies, standard probes are constructed from split-initiator probe pairs as follows. For each split-initiator probe pair in a probe set, the full initiator is shifted onto either the odd probe (full-initiator odd probe + initiator-free even probe) or onto the even probe (initiator-free odd probe + full-initiator even probe). Within each pair, one probe is then a full-initiator standard probe and one probe is a helper probe that contains no initiator. The helper probes are employed to ensure that each standard probe pair (full-initiator probe + helper probe) has the same target-binding capabilities as its analogous split-initiator probe pair. Probe sets with 20 standard probe pairs or split-initiator probe pairs. For background comparisons (Figure S9 and Table S12), the PMT gain is held constant for all experiments to enable comparison of background intensities between experiments. For signal-to-background comparisons (Figure S10 and Table S13), the PMT gain is adjusted to use the full dynamic range for each probe set.
- **Measurement of signal, background, and signal-to-background, background components, and split-initiator HCR suppression.** Measurement of signal, background, and signal-to-background (Figure S11A and Table S14A) using the methods of Section S1.6.2. Measurement of background components (AF, NSA, NSD; Figure S11B and Table S14B) using the methods of Section S1.6.3. Measurement of split-initiator HCR suppression (Figure S11C and Table S14C) using the methods of Section S1.6.4.

##### S3.2.1 Measurement of background and signal-to-background for unoptimized standard and split-initiator probe sets as a function of probe set size

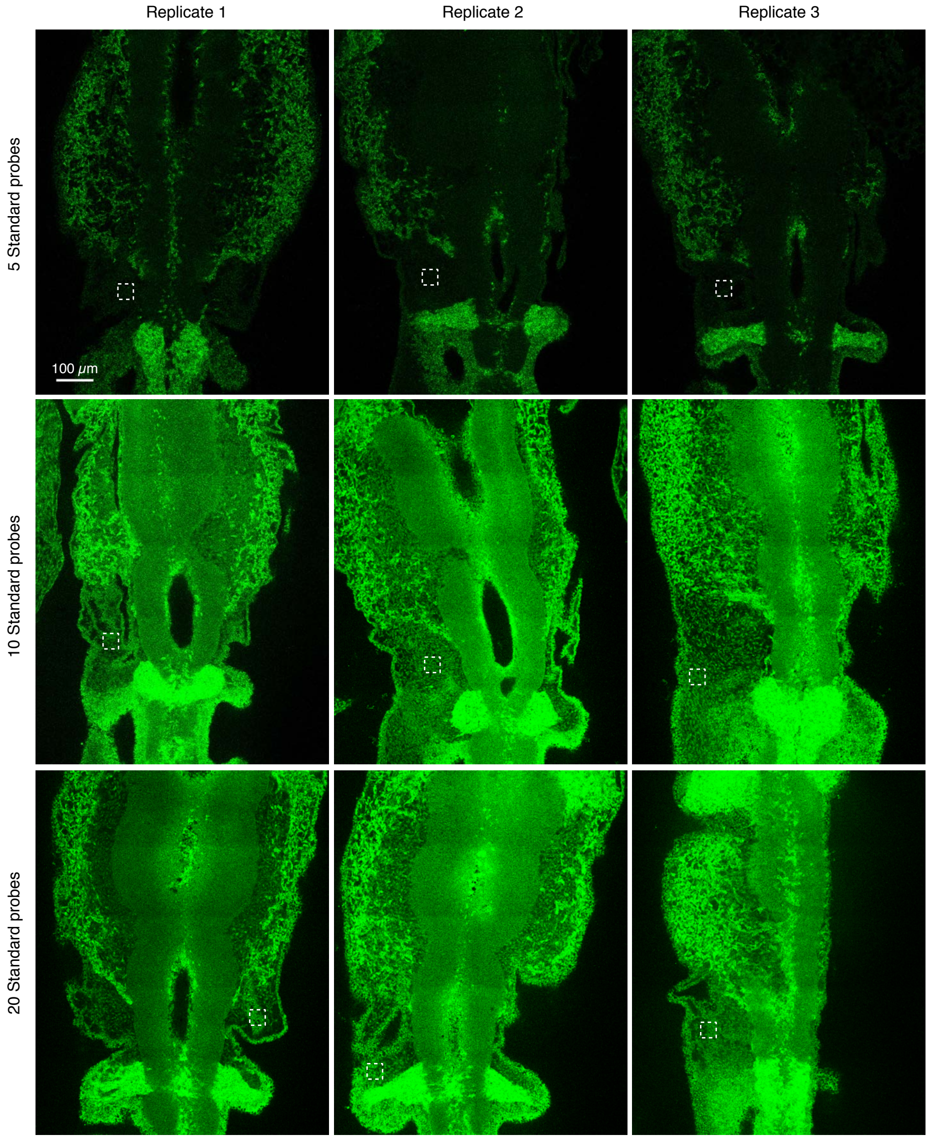

**Figure S5. Measurement of background for unoptimized standard probe sets as a function of probe set size (cf. Figure 3A).** Confocal images collected with the microscope PMT gain adjusted to avoid saturating pixels using 20 standard probes (row 3). Protocol: in situ HCR v2.0 (Section S8 of Choi *et al.* (2016)). Target mRNA: *Sox10*. Probe set: 5, 10, or 20 standard probes. Amplifier: B3-Alexa647. Whole-mount chicken embryos fixed stage HH 11.

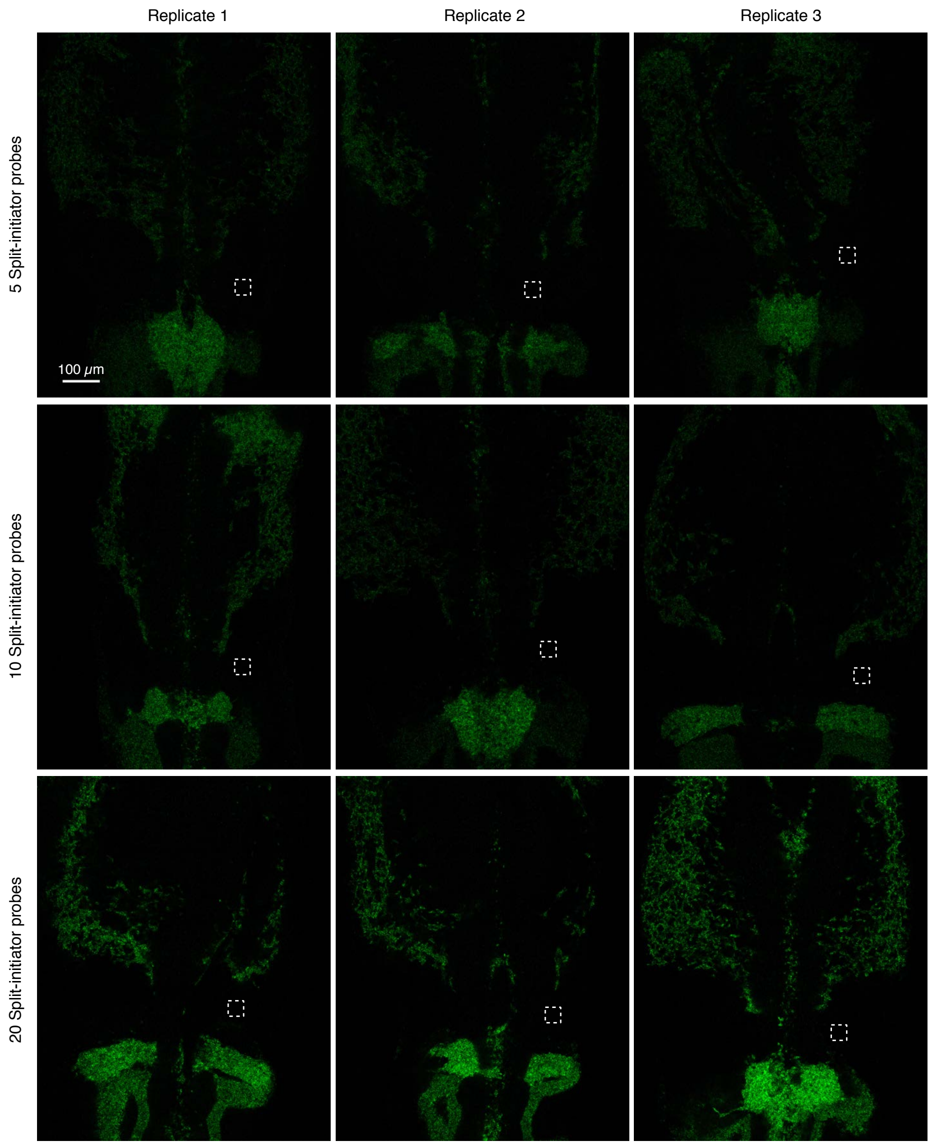

**Figure S6. Measurement of background for unoptimized split-initiator probe sets as a function of probe set size (cf. Figure 3A).** Confocal images collected with the microscope PMT gain adjusted to avoid saturating pixels using 20 standard probes (Figure S5, row 3). Protocol: in situ HCR v3.0 (Section S2.1). Target mRNA: *Sox10*. Probe set: 5, 10, or 20 split-initiator probe pairs. Amplifier: B3-Alexa647. Whole-mount chicken embryos fixed stage HH 11.

| Probe type | Probe set size | BACK |
|---|---|---|
| Standard (v2.0) | 5 | $210 \pm 30$ |
| | 10 | $1160 \pm 60$ |
| | 20 | $1500 \pm 100$ |
| Split-initiator (v3.0) | 5 | $29 \pm 1$ |
| | 10 | $26 \pm 3$ |
| | 20 | $28 \pm 1$ |

**Table S10. Estimated background for standard and split-initiator probes sets as a function of probe set size (cf. Figure 3A).** Mean $\pm$ standard error, $N = 3$ replicate embryos. Analysis based on rectangular regions depicted in Figures S5 and S6 using methods of Section S1.6.2.

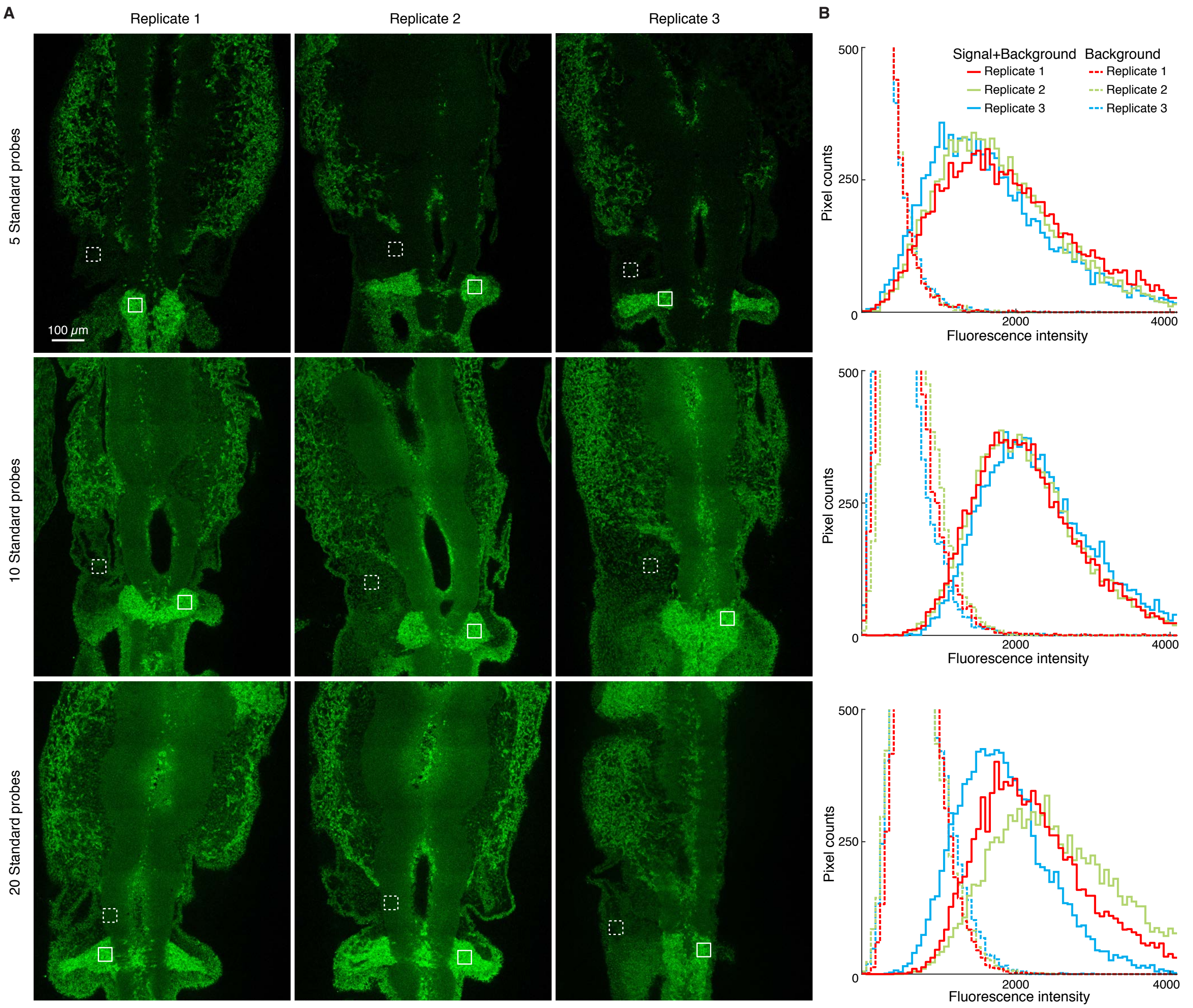

**Figure S7. Measurement of signal and background for unoptimized standard probe sets as a function of probe set size (cf. Figure 3B).** (A) Confocal images collected with the microscope PMT gain optimized for each probe set. (B) Pixel intensity histograms for Signal + Background (pixels within solid boundary) and Background (pixels within dashed boundary) per embryo in panel A. Protocol: in situ HCR v2.0 (Section S8 of Choi *et al.* (2016)). Target mRNA: *Sox10*. Probe set: 5, 10, or 20 standard probes. Amplifier: B3-Alexa647. Whole-mount chicken embryos fixed stage HH 11.

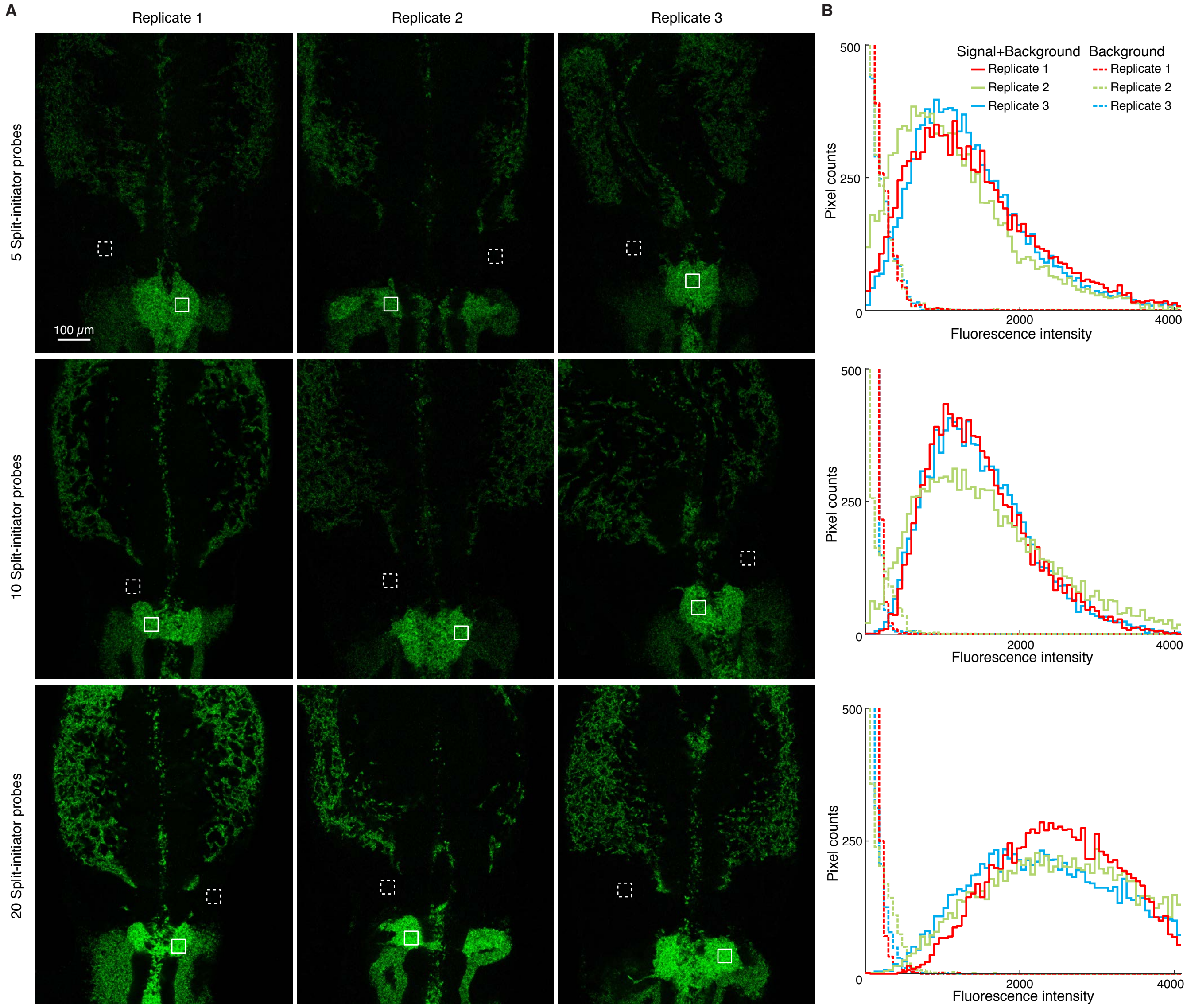

**Figure S8. Measurement of signal and background for unoptimized split-initiator probe sets as a function of probe set size (cf. Figure 3B).** (A) Confocal images collected with the microscope PMT gain optimized for each probe set. (B) Pixel intensity histograms for Signal + Background (pixels within solid boundary) and Background (pixels within dashed boundary) per embryo in panel A. Protocol: in situ HCR v3.0 (Section S2.1). Target mRNA: *Sox10*. Probe set: 5, 10, or 20 split-initiator probe pairs. Amplifier: B3-Alexa647. Whole-mount chicken embryos fixed stage HH 11.

| Probe type | Probe set size | BACK | SIG+BACK | SIG | SIG/BACK |
|---|---|---|---|---|---|
| Standard (v2.0) | 5 | 200 ± 1 | 1770 ± 70 | 1570 ± 70 | 7.8 ± 0.3 |
| | 10 | 540 ± 40 | 2170 ± 40 | 1630 ± 60 | 3.0 ± 0.2 |
| | 20 | 720 ± 10 | 2200 ± 200 | 1400 ± 200 | 2.0 ± 0.3 |
| Split-initiator (v3.0) | 5 | 43.7 ± 0.6 | 1270 ± 70 | 1230 ± 70 | 28 ± 2 |
| | 10 | 34 ± 1 | 1480 ± 30 | 1450 ± 30 | 43 ± 2 |
| | 20 | 37 ± 3 | 2460 ± 40 | 2420 ± 40 | 65 ± 5 |

**Table S11. Estimated signal-to-background for standard and split-initiator probes sets as a function of probe set size (cf. Figure 3B).** Mean ± standard error, $N = 3$ replicate embryos. Analysis based on rectangular regions depicted in Figures S7 and S8 using methods of Section S1.6.2.

##### S3.2.2 Measurement of background and signal-to-background for standard and split-initiator probes with identical target-binding domains

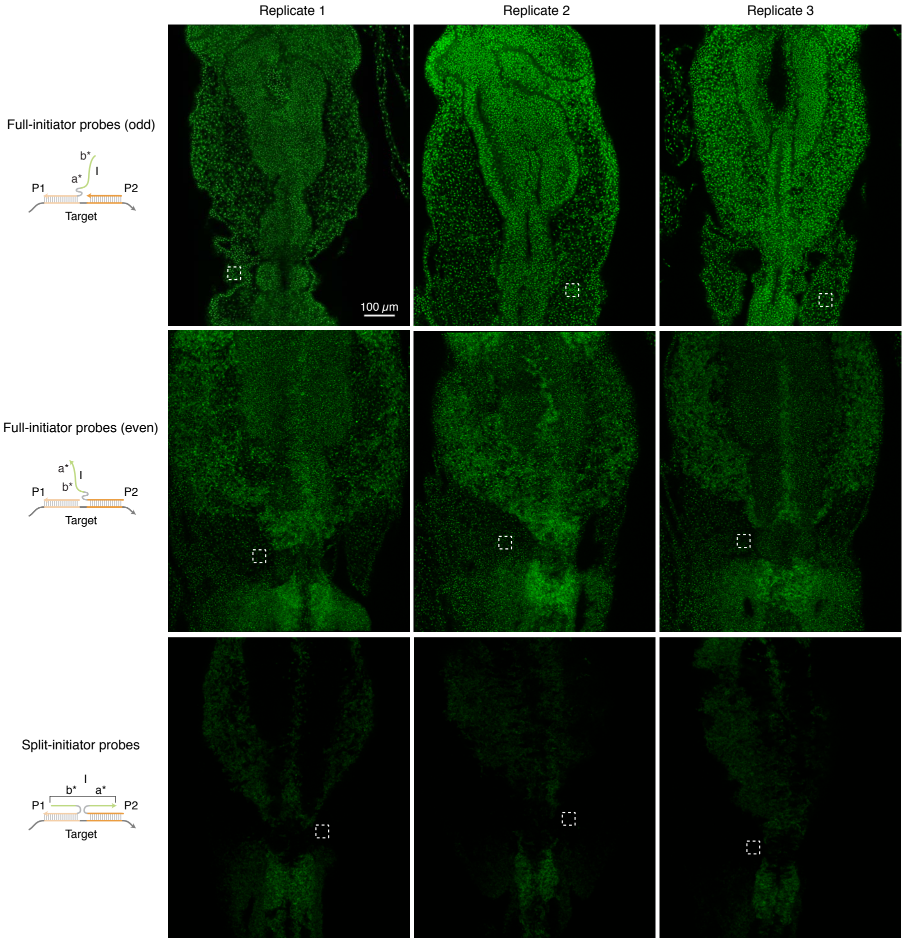

**Figure S9. Measurement of background for standard and split-initiator probes with identical target-binding domains.** (A) Confocal images collected with the microscope PMT gain adjusted to avoid saturating pixels using full-initiator odd probe sets (top row). Protocol: in situ HCR v3.0 (Section S2.1). Target mRNA: *Sox10*. Probe sets (20 probe pairs per probe set): full-initiator odd probes + initiator-free even probes (top row), initiator-free odd probes + full-initiator even probes (middle row), split-initiator odd and even probes (bottom row). Amplifier: B3-Alexa647. Whole-mount chicken embryos fixed stage HH 10.

| Sample | BACK |
|---|---|
| Full-initiator probes (odd) | 910 $\pm$ 80 |
| Full-initiator probes (even) | 330 $\pm$ 40 |
| Split-initiator probes | 19.9 $\pm$ 0.9 |

**Table S12. Estimated background for standard and split-initiator probes with identical target-binding domains.** Mean $\pm$ standard error, $N = 3$ replicate embryos. Analysis based on rectangular regions depicted in Figure S9 using methods of Section S1.6.2.

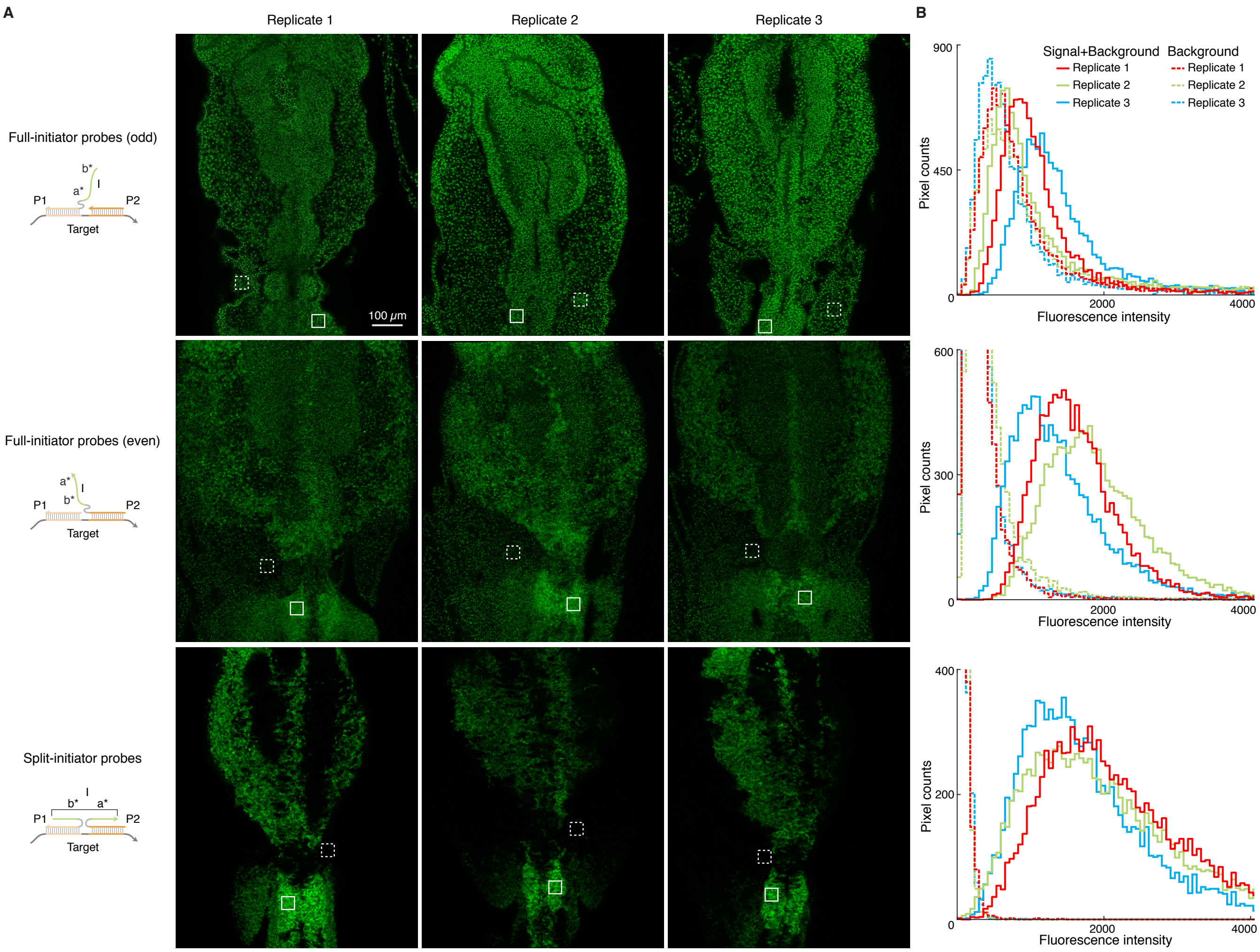

**Figure S10. Measurement of signal and background for standard and split-initiator probes with identical target-binding domains.** (A) Confocal images collected with the microscope PMT gain optimized for each probe set. (B) Pixel intensity histograms for Signal + Background (pixels within solid boundary) and Background (pixels within dashed boundary) per embryo in panel A. Protocol: in situ HCR v3.0 (Section S2.1). Target mRNA: *Sox10*. Probe sets (20 probe pairs per probe set): full-initiator odd probes + initiator-free even probes (top row), initiator-free odd probes + full-initiator even probes (middle row), split-initiator odd and even probes (bottom row). Amplifier: B3-Alexa647. Whole-mount chicken embryos fixed stage HH 10.

| Sample | BACK | SIG+BACK | SIG | SIG/BACK |
|---|---|---|---|---|
| Full-initiator probes (odd) | 910 ± 80 | 1200 ± 100 | 300 ± 100 | 0.4 ± 0.2 |
| Full-initiator probes (even) | 330 ± 40 | 1600 ± 200 | 1300 ± 200 | 3.9 ± 0.6 |
| Split-initiator probes | 26.4 ± 0.7 | 1900 ± 100 | 1800 ± 100 | 70 ± 5 |

**Table S13. Estimated signal-to-background for standard and split-initiator probes with identical target-binding domains.** Mean ± standard error, $N = 3$ replicate embryos. Analysis based on rectangular regions depicted in Figure S10 using methods of Section S1.6.2.

##### S3.2.3 Measurement of signal, background, signal-to-background, background components, and split-initiator HCR suppression

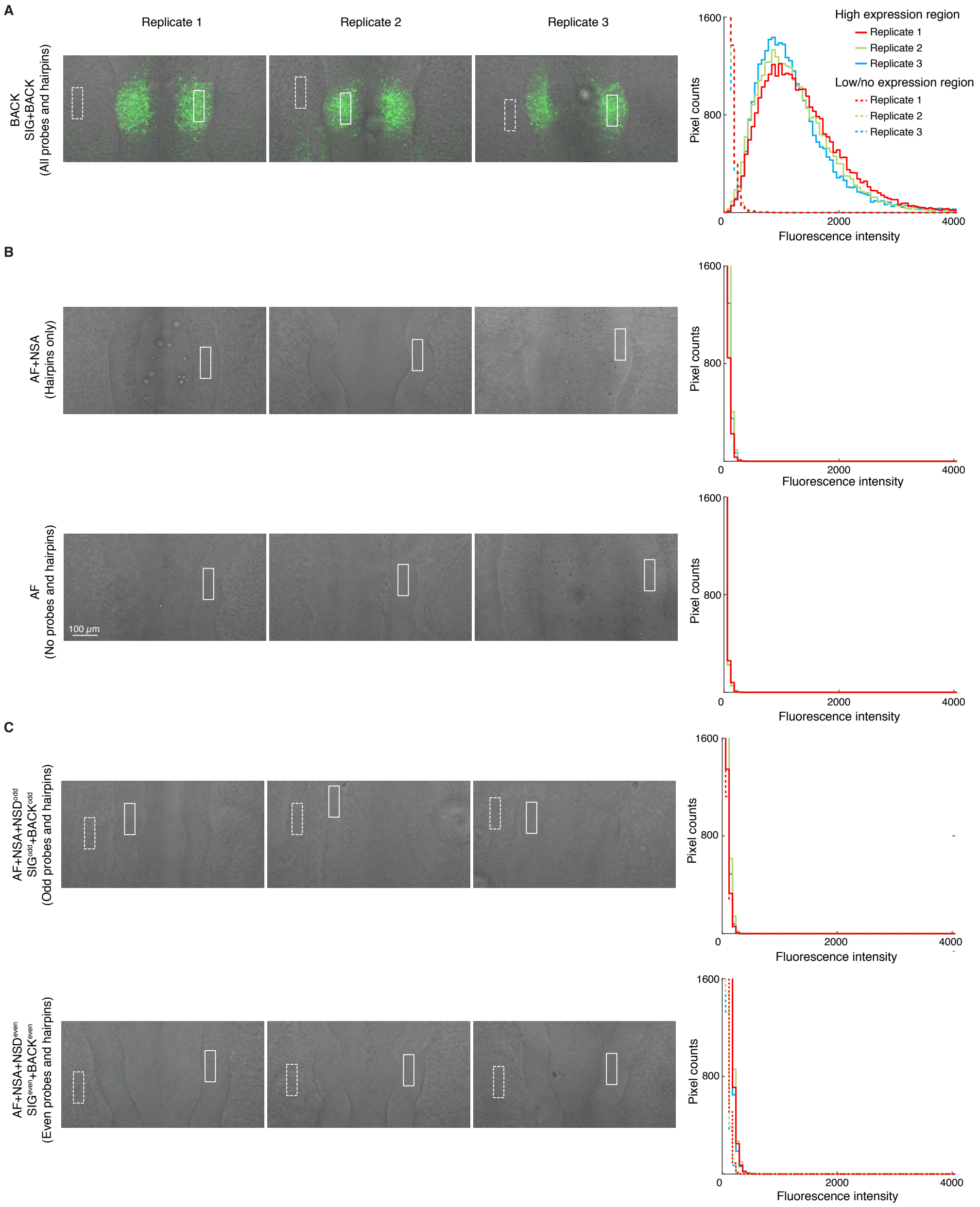

**Figure S11. Measurement of signal and background, background components, and split-initiator HCR suppression.** Confocal image: fluorescence merged with bright field. (A) Signal and background: use experiment of Type 1 in Table S5A (odd probes + even probes + hairpins) to measure SIG+BACK (region of high expression) and BACK (region of no/low expression). (B) Background components: use experiment of Type 2 in Table S5B (no probes, hairpins only) to measure NSA+AF (region of high expression); use experiment of Type 3 (no probes, no hairpins) to measure AF (region of high expression). (C) Split-initiator HCR suppression: use experiment of Type 4 in Table S5C (odd probes, hairpins) to measure $\text{SIG}^{\text{odd}}+\text{BACK}^{\text{odd}}$ (region of high expression) and $\text{BACK}^{\text{odd}}$ (region of no/low expression); use experiment of Type 5 in Table S5C (even probes, hairpins) to measure $\text{SIG}^{\text{even}}+\text{BACK}^{\text{even}}$ (region of high expression) and $\text{BACK}^{\text{even}}$ (region of no/low expression). Left: confocal images collected with the microscope PMT gain optimized to avoid saturating pixels using the full method (top row). Right: pixel intensity histograms for a region of high expression (pixels within solid boundary) and/or low/no expression (pixels within dashed boundary) per embryo. Protocol: in situ HCR v3.0 (Section S2.1). Target mRNA: *EphA4*. Probe set: 20 split-initiator probe pairs. Amplifier: B2-Alexa647. Whole-mount chicken embryos fixed stage HH 11.

| | Quantity | Channel B2-Alexa647 | | Reagents Probes | Reagents Hairpins | Expression region | Figure |
|---|---|---|---|---|---|---|---|
| **A** | SIG+NSD+NSA+AF = SIG+BACK | 1260 | $\pm$ 30 | odd + even | ✓ | high | S11C |
| | NSD+NSA+AF = BACK | 19 | $\pm$ 1 | odd + even | ✓ | low/no | S11C |
| | SIG | 1240 | $\pm$ 30 | | | | |
| | SIG/BACK | 67 | $\pm$ 4 | | | | |
| **B** | NSA+AF | 8.1 | $\pm$ 0.9 | | ✓ | high | S11B |
| | AF | 6.97 | $\pm$ 0.08 | | | high | S11A |
| | NSA | 1.2 | $\pm$ 0.9 | | | | |
| | NSD | 10 | $\pm$ 1 | | | | |
| **C** | $NSD^{odd}$+NSA+AF = $BACK^{odd}$ | 11 | $\pm$ 1 | odd | ✓ | low/no | S11D |
| | $NSD^{even}$+NSA+AF = $BACK^{even}$ | 10.0 | $\pm$ 0.6 | even | ✓ | low/no | S11E |
| | $SIG^{odd}$+$BACK^{odd}$ | 11 | $\pm$ 1 | odd | ✓ | high | S11D |
| | $SIG^{even}$+$BACK^{even}$ | 32 | $\pm$ 2 | even | ✓ | high | S11E |
| | $NSD^{odd}$ | 3 | $\pm$ 1 | | | | |
| | $NSD^{even}$ | 2 | $\pm$ 1 | | | | |
| | $SIG^{odd}$ | $<1.5$ | | | | | |
| | $SIG^{even}$ | 22 | $\pm$ 2 | | | | |
| | $SIG/SIG^{odd}$ | $>800$ | | | | | |
| | $SIG/SIG^{even}$ | 57 | $\pm$ 5 | | | | |

**Table S14. Estimated signal-to-background, background components, and split-initiator HCR suppression.** (A) Signal-to-background (SIG/BACK) based on methods of Section S1.6.2. (B) Background components (AF, NSA, NSD) based on methods of Section S1.6.3. (C) Split-initiator HCR suppression ($SIG/SIG^{odd}$, $SIG/SIG^{even}$) based on methods of Section S1.6.4. Mean $\pm$ standard error, $N = 3$ replicate embryos. Analysis based on rectangular regions depicted in Figure S11.

#### S3.3 Multiplexed 4-channel mRNA imaging with high signal-to-background in whole-mount chicken embryos (cf. Figure 4)

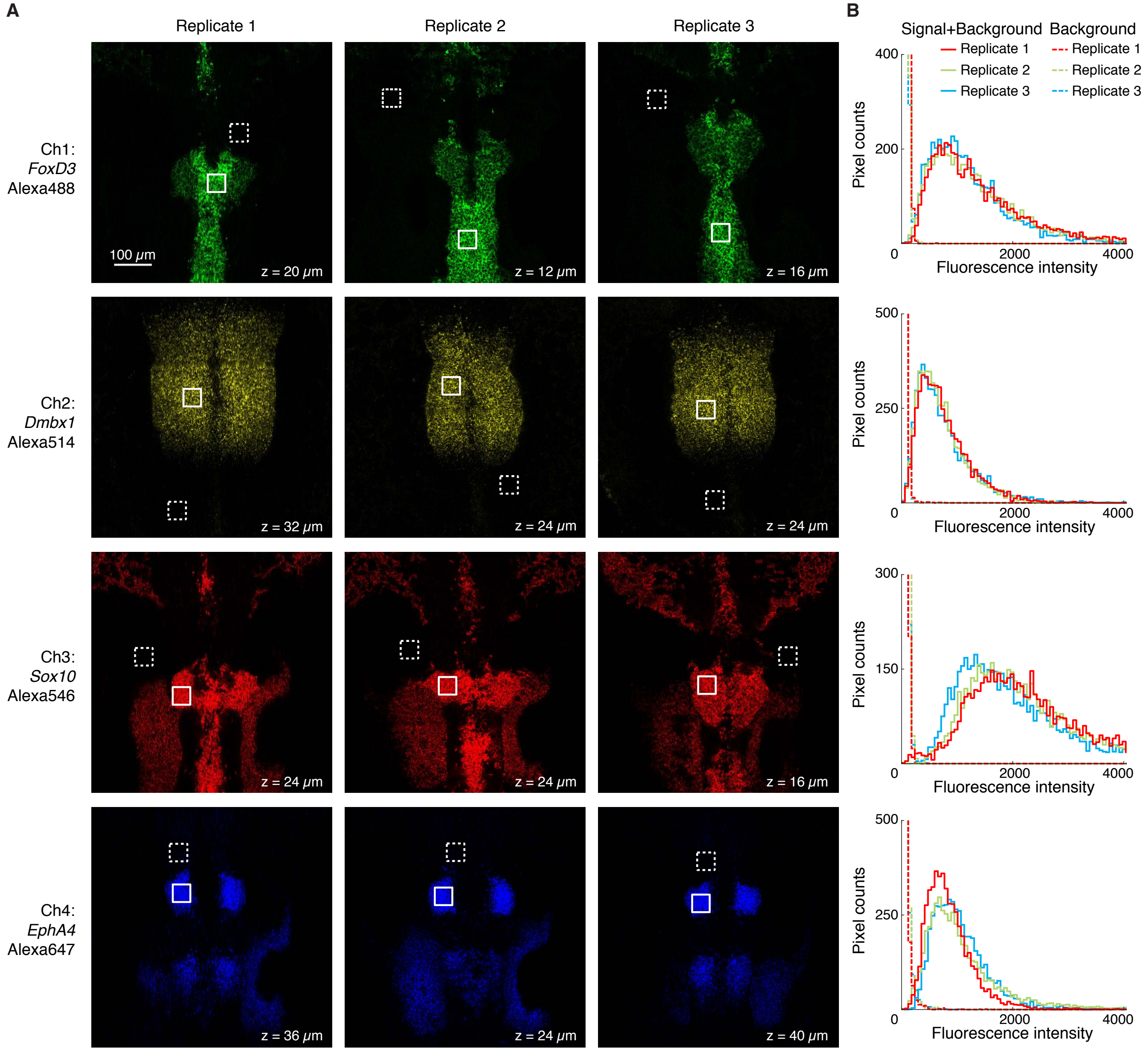

**Figure S12. Measurement of signal and background for multiplexed 4-channel mRNA imaging (cf. Figure 4).** (A) Individual channels of 4-channel confocal images. For each of three replicate embryos, a representative optical section was selected for each channel based on the expression depth of the corresponding target mRNA. (B) Pixel intensity histograms for Signal + Background (pixels within solid boundary) and Background (pixels within dashed boundary). Ch1: target mRNA *FoxD3*, probe set with 12 split-initiator probe pairs, amplifier B4-Alexa488. Ch2: target mRNA *Dmbx1*, probe set with 20 split-initiator probe pairs, amplifier B1-Alexa514. Ch 3: target mRNA *Sox10*, probe set with 20 split-initiator probe pairs, amplifier B3-Alexa546. Ch4: *EphA4*, probe set with 20 split-initiator probe pairs, amplifier B2-Alexa647. Whole-mount chicken embryos fixed stage HH 10.

| Target mRNA | BACK | SIG+BACK | SIG | SIG/BACK |
|---|---|---|---|---|
| *FoxD3* | $36 \pm 3$ | $1270 \pm 50$ | $1230 \pm 50$ | $34 \pm 3$ |
| *Dmbx1* | $16.4 \pm 0.2$ | $730 \pm 20$ | $710 \pm 20$ | $43 \pm 1$ |
| *Sox10* | $34.0 \pm 0.2$ | $2030 \pm 80$ | $2000 \pm 80$ | $59 \pm 3$ |
| *EphA4* | $35 \pm 3$ | $980 \pm 70$ | $950 \pm 70$ | $27 \pm 3$ |

**Table S15. Estimated signal-to-background for multiplexed 4-channel mRNA imaging.** Mean $\pm$ standard error, $N = 3$ replicate embryos. Analysis based on rectangular regions depicted in Figure S12 using methods of Section S1.6.2.

#### S3.4 qHCR imaging: analog mRNA relative quantitation with subcellular resolution in whole-mount chicken embryos (cf. Figure 5)

##### S3.4.1 Testing for a crowding effect

In order to perform multiplexed quantitative imaging using HCR, it is important that there is not a crowding effect in which amplification polymers tethered to one target molecule affect the signal intensity for a different target molecule. To test for a possible crowding effect, we imaged two target mRNAs that are highly expressed in the same cells (*EphA4* and *Egr2*) individually (1-target studies) and also simultaneously (2-target studies) within whole-mount chicken embryos. Figure S13 compares the signal intensity distributions for 1-target and 2-target studies, revealing similar intensity distributions whether targets were detected alone or together, suggesting that there is not a significant crowding effect (either antagonistic or synergistic).

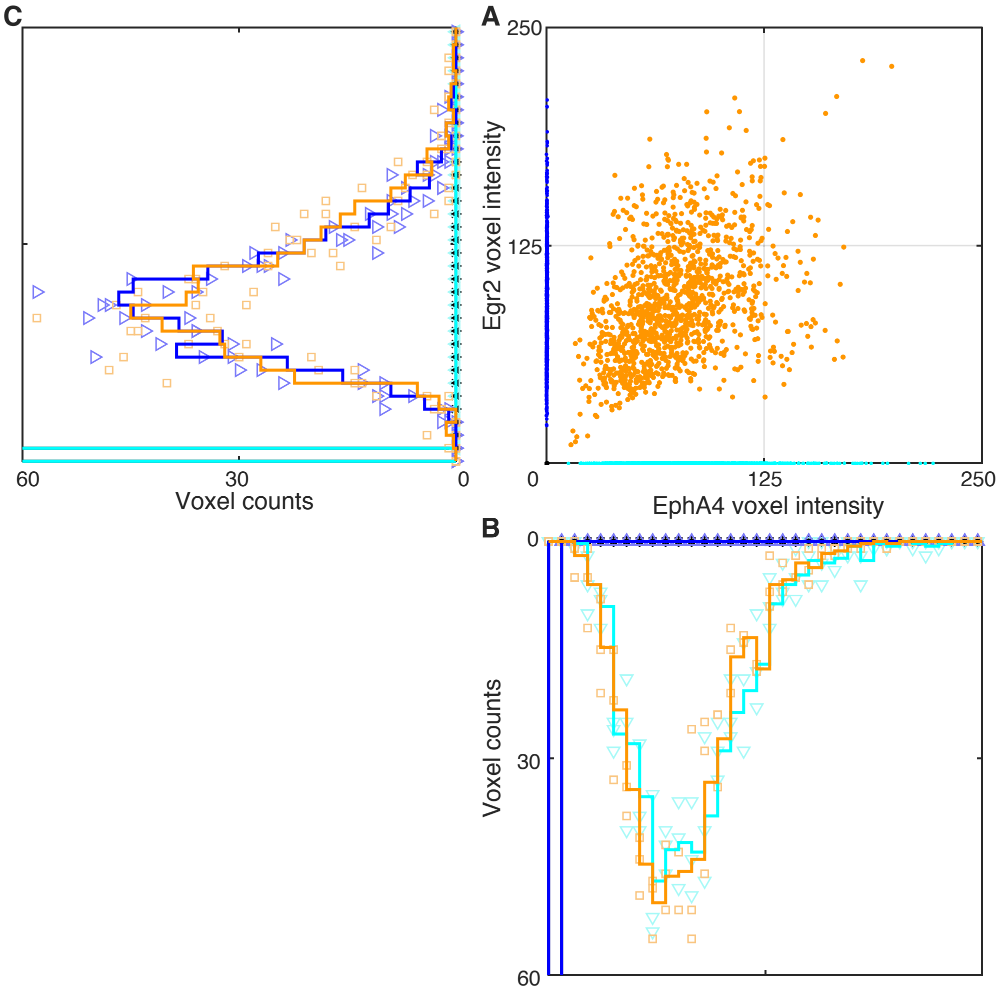

**Figure S13.** Comparison of signal intensity distributions for individual and simultaneous imaging of *EphA4* and *Egr2*. (A) Raw voxel intensity scatter plot: *Egr2* channel vs *EphA4* channel. (B) Raw voxel intensity histogram for *EphA4* channel. (C) Raw voxel intensity histogram for *Egr2* channel. In panels B and C, solid lines denote average histograms over 3 replicate embryos while symbols denote individual histograms (1 histogram per replicate). Orange data: signal plus background for *EphA4* and *Egr2* (Figure S14). Cyan data: signal plus background for *EphA4* and background for *Egr2* (Figure S15). Blue data: background for *EphA4* and signal plus background for *Egr2* (Figure S16). Black data: background for *EphA4* and *Egr2* (Figure S17). Voxel size: 2.1×2.1×2.7 $\mu$m. Whole-mount wildtype chicken embryos fixed stage HH 10.

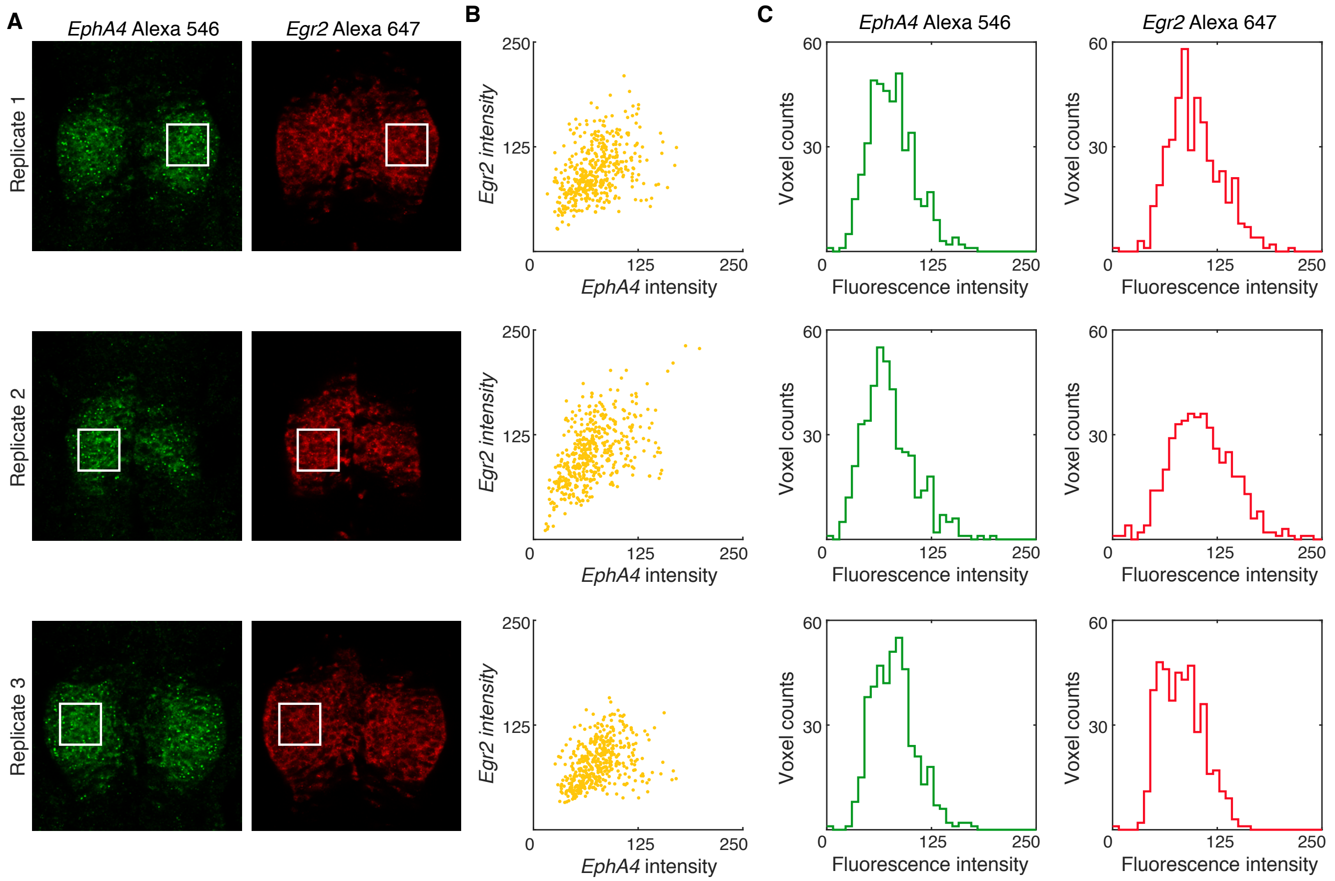

**Figure S14.** Characterizing signal plus background for *EphA4* and *Egr2* in a 2-target experiment. (A) Individual channels from 2-channel confocal images depicting regions used to estimate signal plus background. *EphA4* channel: 20 probe pairs and amplifier B1-Alexa546. *Egr2* channel: 20 probe pairs and amplifier B3-Alexa647. For each of 3 replicate embryos, a representative optical section was selected based on the expression depth of the target mRNAs. Pixel size: $0.4\times0.4$ $\mu$m. (B) Raw voxel intensity scatter plots for the selected regions of panel A representing signal plus background for *EphA4* and *Egr2*. (C) Raw voxel intensity histograms for the selected regions of panel A representing signal plus background for *EphA4* and *Egr2*. Same microscope settings used for all replicates in Figures S14–S17. Voxel size: $2.1\times2.1\times2.7$ $\mu$m. Whole-mount wildtype chicken embryos fixed stage HH 10.

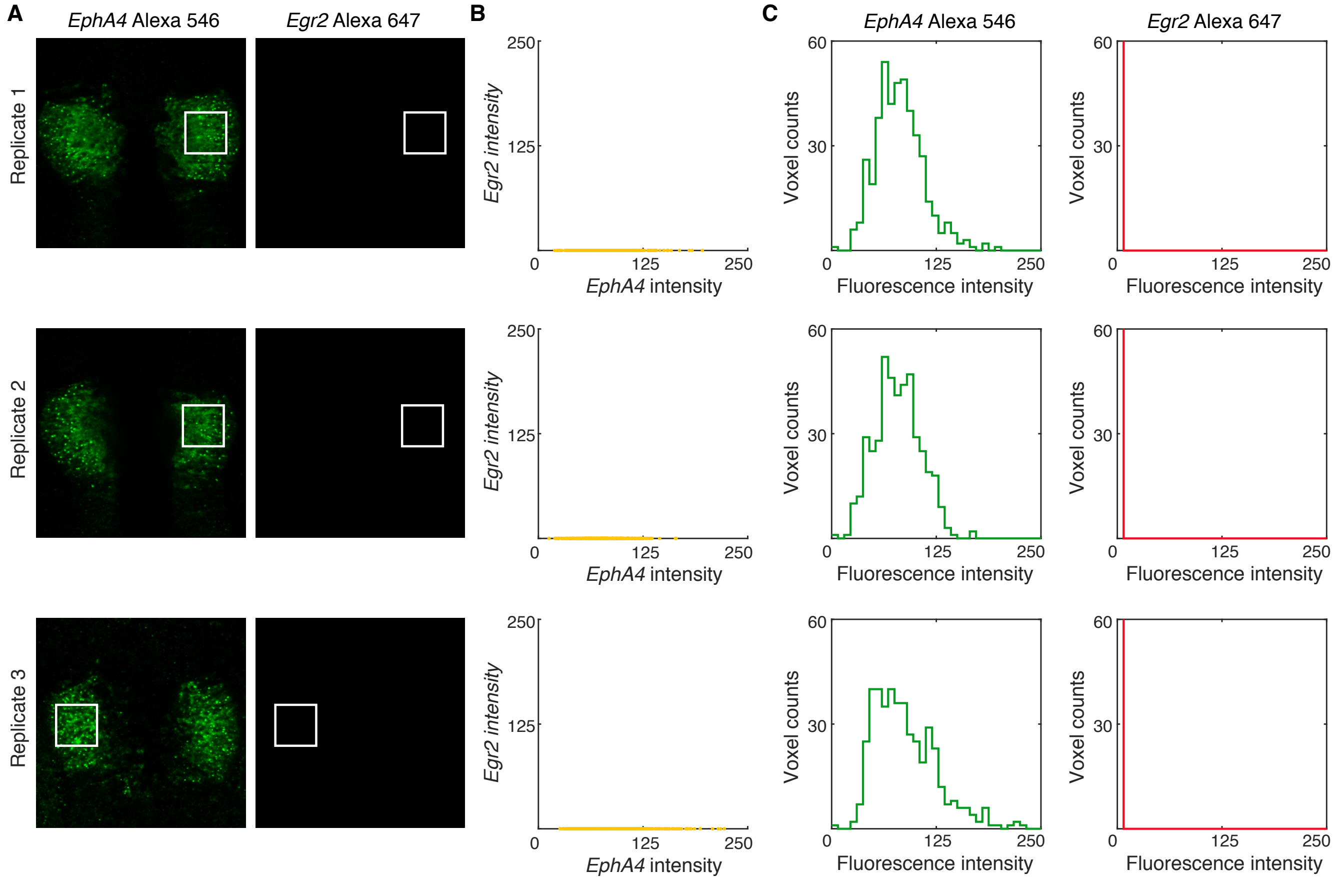

**Figure S15.** Characterizing signal plus background for *EphA4* in a 1-target experiment. (A) Individual channels from 2-channel confocal images depicting regions used to estimate signal plus background for *EphA4* and background for *Egr2*. *EphA4* channel: 20 probe pairs and amplifier B1-Alexa546. *Egr2* channel: no probes, no amplifier. For each of 3 replicate embryos, a representative optical section was selected based on the expression depth of the target mRNA. Pixel size: $0.4\times0.4$ $\mu$m. (B) Raw voxel intensity scatter plots for the selected regions of panel A representing signal plus background for *EphA4* and background for *Egr2*. (C) Raw voxel intensity histograms for the selected regions of panel A representing signal plus background for *EphA4* and background for *Egr2*. Same microscope settings used for all replicates in Figures S14–S17. Voxel size: $2.1\times2.1\times2.7$ $\mu$m. Whole-mount wildtype chicken embryos fixed stage HH 10.

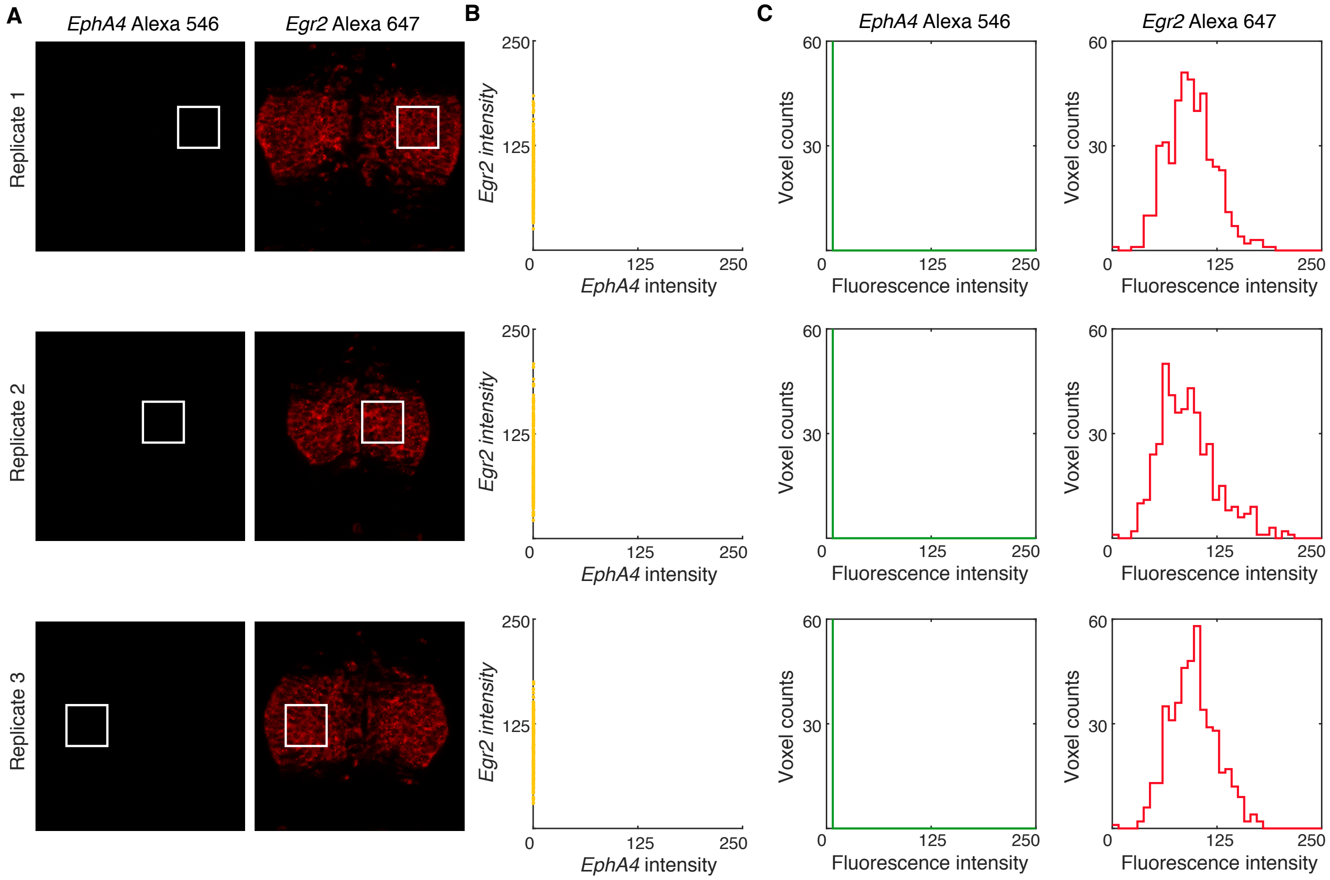

**Figure S16.** Characterizing signal plus background for *Egr2* in a 1-target experiment. (A) Individual channels from 2-channel confocal images depicting regions used to estimate background for *EphA4* and signal plus background for *Egr2*. *EphA4* channel: no probes, no amplifier. *Egr2* channel: 20 probe pairs and amplifier B3-Alexa647. For each of 3 replicate embryos, a representative optical section was selected based on the expression depth of the target mRNA. Pixel size: $0.4 \times 0.4$ $\mu$m. (B) Raw voxel intensity scatter plots for the selected regions of panel A representing background for *EphA4* and signal plus background for *Egr2*. (C) Raw voxel intensity histograms for the selected regions of panel A representing background for *EphA4* and signal plus background for *Egr2*. Same microscope settings used for all replicates in Figures S14–S17. Voxel size: $2.1 \times 2.1 \times 2.7$ $\mu$m. Whole-mount wildtype chicken embryos fixed stage HH 10.

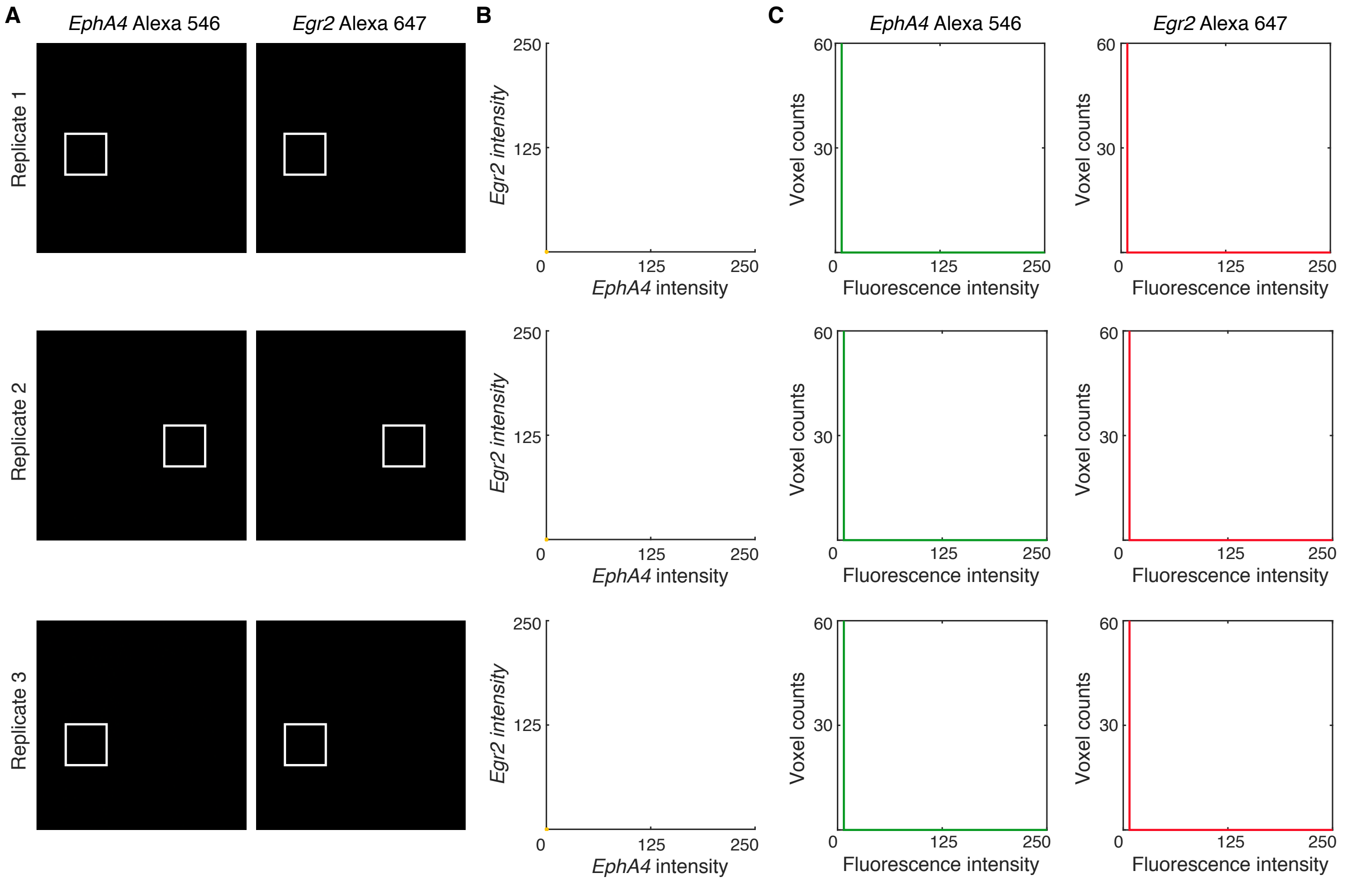

**Figure S17.** Characterizing background for *EphA4* and *Egr2*. Individual channels from 2-channel confocal images depicting regions used to estimate background using the standard HCR v3.0 in situ protocol (Section S2.1) omitting probes (BACK ≈ AF + NSA; see Section S1.6 for definitions). For each of 3 replicate embryos, a representative optical section was selected at approximately the depth where *EphA4* and *Egr2* are expressed. Same microscope settings used for all replicates in Figures S14–S17. Pixel size: $0.4\times0.4$ $\mu$m. (B) Raw voxel intensity scatter plots for the selected region of panel A. (C) Raw voxel intensity histograms for the scatter plots of panel B. Voxel size: $2.1\times2.1\times2.7$ $\mu$m. Whole-mount wildtype chicken embryos fixed stage HH 10.

##### S3.4.2 Redundant 2-channel detection of *Dmbx1*

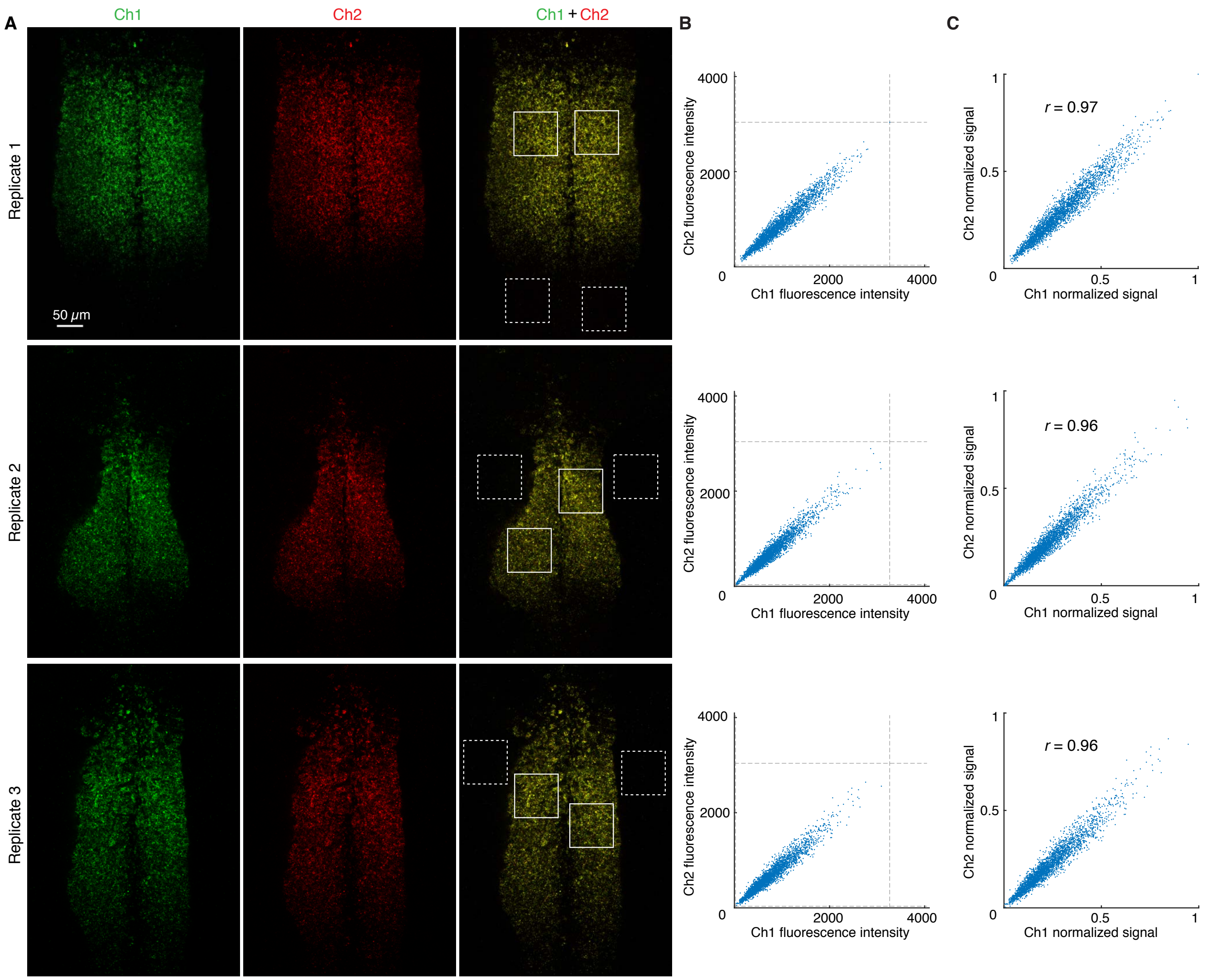

**Figure S18. Redundant 2-channel detection of *Dmbx1* (cf. Figure 5).** (A) Confocal images: individual channels and merge. Solid boundaries denote regions of high expression; dashed boundaries denote regions of no/low expression. Pixel size: 0.2×0.2 $\mu$m. Probe sets: 20 split-initiator probe pairs per channel. Amplifiers: B1-Alexa546 (Ch1) and B2-Alexa647 (Ch2). Whole-mount chicken embryos fixed stage HH 10. (B) Raw voxel intensity scatter plots representing signal plus background for voxels within solid boundaries of panel A. Voxel size: 2.1×2.1×2.7 $\mu$m. Dashed lines represent BOT and TOP values (Table S16) used to normalize data for panel C using methods of Section S1.6.5. (C) Normalized voxel intensity scatter plots representing estimated normalized signal (Pearson correlation coefficient, $r$).

##### S3.4.3 Redundant 2-channel detection of *EphA4*

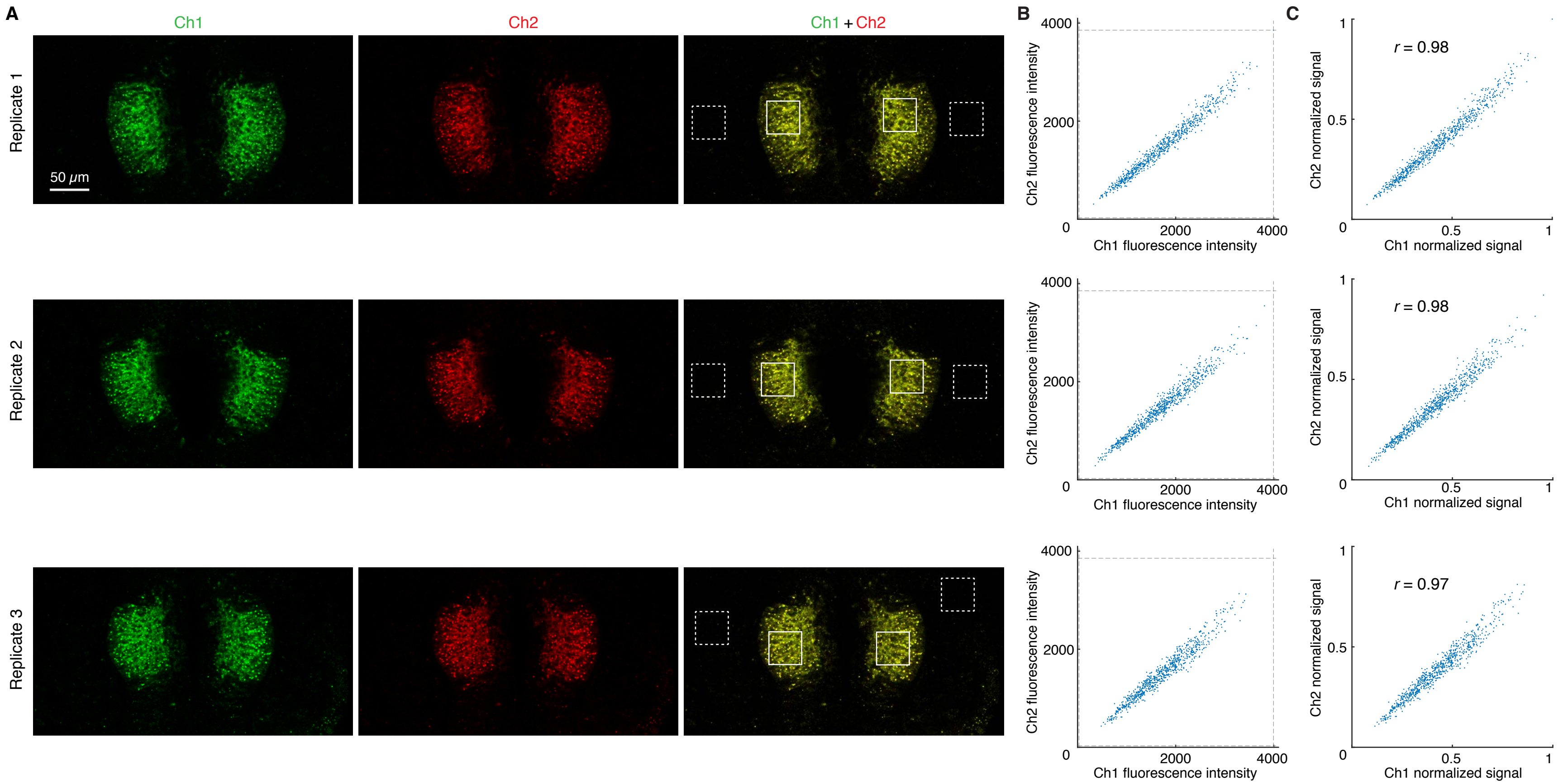

**Figure S19. Redundant 2-channel detection of *EphA4* (cf. Figure 5).** (A) Confocal images: individual channels and merge. Solid boundaries denote regions of high expression; dashed boundaries denote regions of no/low expression. Pixel size: $0.2 \times 0.2$ $\mu$m. Probe sets: 20 split-initiator probe pairs per channel. Amplifiers: B1-Alexa546 (Ch1) and B2-Alexa647 (Ch2). Whole-mount chicken embryos fixed stage HH 10. (B) Raw voxel intensity scatter plots representing signal plus background for voxels within solid boundaries of panel A. Voxel size: $2.1 \times 2.1 \times 2.7$ $\mu$m. Dashed lines represent BOT and TOP values (Table S16) used to normalize data for panel C using methods of Section S1.6.5. (C) Normalized voxel intensity scatter plots representing estimated normalized signal (Pearson correlation coefficient, $r$).

| Target mRNA | Channel | BACK | SIG+BACK | SIG | SIG/BACK | BOT | TOP |
|---|---|---|---|---|---|---|---|
| *Dmbx1* | Alexa546 | $24 \pm 1$ | $840 \pm 80$ | $810 \pm 80$ | $34 \pm 4$ | 24 | 3266 |
| *Dmbx1* | Alexa647 | $35 \pm 5$ | $770 \pm 70$ | $730 \pm 70$ | $21 \pm 3$ | 35 | 3040 |
| *EphA4* | Alexa546 | $29 \pm 1$ | $1720 \pm 10$ | $1690 \pm 10$ | $59 \pm 2$ | 29 | 3995 |
| *EphA4* | Alexa647 | $30 \pm 1$ | $1490 \pm 10$ | $1460 \pm 10$ | $49 \pm 2$ | 30 | 3855 |

**Table S16. Estimated signal-to-background for redundant 2-channel detection of *Dmbx1* and *EphA4*.** Mean $\pm$ standard error, $N = 3$ replicate embryos. Analysis based on rectangular regions depicted in Figure S18A and S19A using methods of Section S1.6.2. BOT and TOP values used to calculate normalized voxel intensities for scatter plots of Figures 5C, S18C, and S19C using methods of Section S1.6.5.

#### S3.5 In situ validation of automatic background suppression with split-initiator probes for mRNA flow cytometry with cultured human and bacterial cells

The methods of Sections S1.7.2 and S1.7.3 are used to measure:

- signal, background, and signal-to-background (Figures S20A–S22A and Tables S17A–S19A) .
- background components (AF, NSA, NSD; Figures S20A–S22A and Tables S17B–S19B).
- split-initiator HCR suppression (Figures S20B–S22B and Tables S17C–S19C).

Additional measurements of these quantities are provided in the 2-channel experiments of Figures S24–S28 and Tables S20–S24.

##### S3.5.1 Measurement of signal, background, signal-to-background, background components, and split-initiator HCR suppression for *d2eGFP* transgenic target in HEK cells

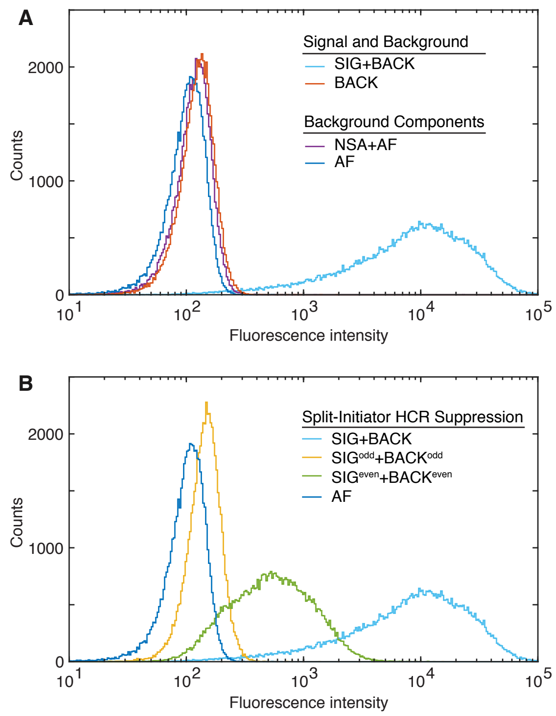

**Figure S20. Measurement of signal and background, background components, and split-initiator HCR suppression for *d2eGFP* transgenic target in HEK cells (cf. Figure 6A).** (A) Signal and background: use experiments of Types 1a and 1b in Table S7A (odd + even probes, hairpins) to measure SIG+BACK (GFP+ cells) and BACK (WT cells). Background components: use experiment of Type 2 in Table S7B (no probes, with hairpins) to measure NSA+AF (GFP+ cells); use experiment of Type 3 (no probes, no hairpins) to measure AF (GFP+ cells). (B) Split-initiator HCR suppression: use experiment of Types 4a in Table S5C (odd probes, hairpins) to measure SIG$^{odd}$+BACK$^{odd}$ (GFP+ cells); use experiment of Type 5a in Table S5C (even probes, hairpins) to measure SIG$^{even}$+BACK$^{even}$ (GFP+ cells). Distribution of single-cell fluorescence intensities. Protocol: in situ HCR v3.0 (Section S2.3). Probe set: 12 split-initiator probe pairs. Amplifier: B3-Alexa594. Sample: 55,000 HEK cells in suspension (GFP+ or WT).

| | Quantity | Channel B3-Alexa594 | Reagents Probes | Hairpins | Cell type |
|---|---|---|---|---|---|
| **A** | SIG+NSD+NSA+AF = SIG+BACK | 13 220 ± 60 | odd + even | ✓ | GFP+ |
| | NSD+NSA+AF = BACK | 128.5 ± 0.2 | odd + even | ✓ | WT |
| | SIG | 13 090 ± 60 | | | |
| | SIG/BACK | 101.8 ± 0.5 | | | |
| **B** | NSA+AF | 120.7 ± 0.5 | | ✓ | GFP+ |
| | AF | 104.7 ± 0.2 | | ✓ | GFP+ |
| | NSA | 16.0 ± 0.5 | | | |
| | NSD | 7.8 ± 0.5 | | | |
| **C** | $SIG^{odd}$+$BACK^{odd}$ | 149.1 ± 0.3 | odd | ✓ | GFP+ |
| | $SIG^{even}$+$BACK^{even}$ | 710 ± 3 | even | ✓ | GFP+ |
| | $SIG^{odd}$ | 28.4 ± 0.6 | | | |
| | $SIG^{even}$ | 589 ± 3 | | | |
| | SIG/$SIG^{odd}$ | 461 ± 9 | | | |
| | SIG/$SIG^{even}$ | 22.2 ± 0.1 | | | |

**Table S17. Estimated signal-to-background, background components, and split-initiator HCR suppression for *d2eGFP* transgenic target in HEK cells (cf. Figure 6A).** (A) Signal-to-background (SIG/BACK). (B) Background components (AF, NSA, NSD). (C) Split-initiator HCR suppression (SIG/$SIG^{odd}$, SIG/$SIG^{even}$). The signal estimates $SIG^{odd}$ and $SIG^{even}$ are calculated using the background approximation $BACK^{odd}$ = $BACK^{even}$ $\approx$ NSA+AF, which leads to lower bounds on SIG/$SIG^{odd}$ and SIG/$SIG^{even}$. Mean $\pm$ standard error, $N = 55,000$ cells. Analysis based on single-cell intensities of Figure S20 using methods of Section S1.7.2.

##### S3.5.2 Measurement of signal, background, signal-to-background, background components, and split-initiator HCR suppression for *GAPDH* endogenous target in HEK cells

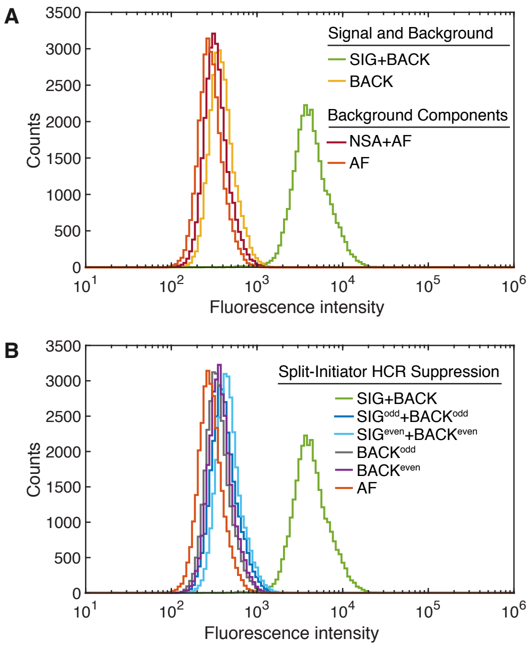

**Figure S21. Measurement of signal and background, background components, and split-initiator HCR suppression for *GAPDH* endogenous target in HEK cells.** (A) Signal and background: use experiments of Types 1a and 1b in Table S8A to measure SIG+BACK (even + odd probes, hairpins) and BACK (Tg(odd) + Tg(even) probes, hairpins). Background components: use experiment of Type 2 in Table S8B (no probes, with hairpins) to measure NSA+AF; use experiment of Type 3 (no probes, no hairpins) to measure AF. (B) Split-initiator HCR suppression: use experiment of Types 4a and 4b in Table S8C (odd probes, hairpins) to measure SIG$^{\mathrm{odd}}$+BACK$^{\mathrm{odd}}$ (odd probes, hairpins) and BACK$^{\mathrm{odd}}$ (Tg(odd) probes, hairpins); use experiments of Types 5a and 5b in Table S8C to measure SIG$^{\mathrm{even}}$+BACK$^{\mathrm{even}}$ (even probes, hairpins) and BACK$^{\mathrm{even}}$ (Tg(even) probes, hairpins). Distribution of single-cell fluorescence intensities. Protocol: in situ HCR v3.0 (Section S2.3). Probe set: 10 split-initiator probe pairs. Amplifier: B4-Alexa594. Sample: 30,000 HEK cells in suspension (WT).

| | Quantity | Channel B4-Alexa594 | | Reagents Probes | Hairpins | Cell type |
|---|---|---|---|---|---|---|
| **A** | SIG+NSD+NSA+AF = SIG+BACK | 4775 | ± 15 | odd + even | ✓ | WT |
| | NSD+NSA+AF = BACK | 414.0 | ± 0.9 | Tg(odd) + Tg(even) | ✓ | WT |
| | SIG | 4362 | ± 15 | | | |
| | SIG/BACK | 10.55 | ± 0.04 | | | |
| **B** | NSA+AF | 353.7 | ± 0.8 | | ✓ | WT |
| | AF | 304.0 | ± 0.7 | | ✓ | WT |
| | NSA | 50 | ± 1 | | | |
| | NSD | 60 | ± 1 | | | |
| **C** | $SIG^{odd}$+$NSD^{odd}$+NSA+AF = $SIG^{odd}$+$BACK^{odd}$ | 450 | ± 7 | odd | ✓ | WT |
| | $NSD^{odd}$+NSA+AF = $BACK^{odd}$ | 371.2 | ± 0.8 | Tg(odd) | ✓ | WT |
| | $SIG^{even}$+$NSD^{even}$+NSA+AF = $SIG^{even}$+$BACK^{even}$ | 499 | ± 1 | even | ✓ | WT |
| | $NSD^{even}$+NSA+AF = $BACK^{even}$ | 397.3 | ± 0.8 | Tg(even) | ✓ | WT |
| | $NSD^{odd}$ | 18 | ± 1 | | | |
| | $NSD^{even}$ | 44 | ± 1 | | | |
| | $SIG^{odd}$ | 79 | ± 7 | | | |
| | $SIG^{even}$ | 102 | ± 1 | | | |
| | SIG/$SIG^{odd}$ | 55 | ± 5 | | | |
| | SIG/$SIG^{even}$ | 42.9 | ± 0.6 | | | |

**Table S18. Estimated signal-to-background, background components, and split-initiator HCR suppression for *GAPDH* endogenous target in HEK cells.** (A) Signal-to-background (SIG/BACK). (B) Background components (AF, NSA, NSD). (C) Split-initiator HCR suppression (SIG/$SIG^{odd}$, SIG/$SIG^{even}$). Mean ± standard error, $N = 30,000$ cells. Analysis based on single-cell intensities of Figure S21 using methods of Section S1.7.3.

##### S3.5.3 Measurement of signal, background, signal-to-background, background components, and split-initiator HCR suppression for *eGFP* transgenic target in *E. coli*

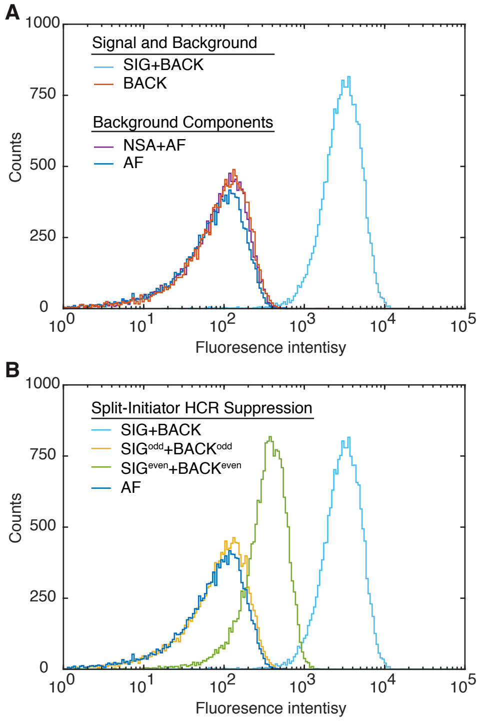

**Figure S22. Measurement of signal and background, background components, and split-initiator HCR suppression for *eGFP* transgenic target in *E. coli* (cf. Figure 6A).** (A) Signal and background: use experiments of Types 1a and 1b in Table S7A (odd + even probes, hairpins) to measure SIG+BACK (GFP+ cells) and BACK (WT cells). Background components: use experiment of Type 2 in Table S7B (no probes, with hairpins) to measure NSA+AF (GFP+ cells); use experiment of Type 3 (no probes, no hairpins) to measure AF (GFP+ cells). (B) Split-initiator HCR suppression: use experiment of Types 4a in Table S5C (odd probes, hairpins) to measure $\text{SIG}^{\text{odd}}$+$\text{BACK}^{\text{odd}}$ (GFP+ cells); use experiment of Type 5a in Table S5C (even probes, hairpins) to measure $\text{SIG}^{\text{even}}$+$\text{BACK}^{\text{even}}$ (GFP+ cells). Distribution of single-cell fluorescence intensities. Protocol: in situ HCR v3.0 (Section S2.4). Probe set: 12 split-initiator probe pairs. Amplifier: B3-Alexa594. Sample: 18,000 *E. coli* DH5$\alpha$ in suspension (GFP+ or WT).

| | Quantity | Channel B3-Alexa594 | Reagents Probes | Reagents Hairpins | Cell type |
|---|---|---|---|---|---|
| **A** | SIG+NSD+NSA+AF = SIG+BACK | $3330 \pm 10$ | odd + even | ✓ | GFP+ |
| | NSD+NSA+AF = BACK | $120 \pm 20$ | odd + even | ✓ | WT |
| | SIG | $3200 \pm 30$ | | | |
| | SIG/BACK | $26 \pm 5$ | | | |
| **B** | NSA+AF | $72.5 \pm 0.7$ | | ✓ | GFP+ |
| | AF | $55.7 \pm 0.7$ | | ✓ | GFP+ |
| | NSA | $17 \pm 1$ | | | |
| | NSD | $50 \pm 20$ | | | |
| **C** | $\mathrm{SIG}^{\mathrm{odd}}$+$\mathrm{BACK}^{\mathrm{odd}}$ | $71.3 \pm 0.7$ | odd | ✓ | GFP+ |
| | $\mathrm{SIG}^{\mathrm{even}}$+$\mathrm{BACK}^{\mathrm{even}}$ | $400 \pm 10$ | even | ✓ | GFP+ |
| | $\mathrm{SIG}^{\mathrm{odd}}$ | $<1$ | | | |
| | $\mathrm{SIG}^{\mathrm{even}}$ | $320 \pm 10$ | | | |
| | SIG/$\mathrm{SIG}^{\mathrm{odd}}$ | $>3000$ | | | |
| | SIG/$\mathrm{SIG}^{\mathrm{even}}$ | $9.9 \pm 0.5$ | | | |

**Table S19. Estimated signal-to-background, background components, and split-initiator HCR suppression for *eGFP* transgenic target in *E. coli* (cf. Figure 6A).** (A) Signal-to-background (SIG/BACK). (B) Background components (AF, NSA, NSD). (C) Split-initiator HCR suppression (SIG/$\mathrm{SIG}^{\mathrm{odd}}$, SIG/$\mathrm{SIG}^{\mathrm{even}}$). The signal estimates $\mathrm{SIG}^{\mathrm{odd}}$ and $\mathrm{SIG}^{\mathrm{even}}$ are calculated using the background approximation $\mathrm{BACK}^{\mathrm{odd}}$ = $\mathrm{BACK}^{\mathrm{even}}$ $\approx$ NSA+AF, which leads to lower bounds on SIG/$\mathrm{SIG}^{\mathrm{odd}}$ and SIG/$\mathrm{SIG}^{\mathrm{even}}$. Mean $\pm$ standard error, $N = 18,000$ cells. Analysis based on single-cell intensities of Figure S22 using methods of Section S1.7.2.

#### S3.6 qHCR flow cytometry: analog mRNA relative quantitation for high-throughput analyses of human and bacterial cells (cf. Figure 6)

##### S3.6.1 Testing for a crowding effect

In order to perform multiplexed quantitative flow cytometry using HCR, it is important that there is not a crowding effect in which amplification polymers tethered to one target molecule affect the signal intensity for a different target molecule. To test for a possible crowding effect, we analyzed two highly-expressed target mRNAs (*GAPDH* and *ACTB*) individually (1-target studies) and also simultaneously (2-target studies) within HEK cells. Figure S23 compares the signal intensity distributions for 1-target and 2-target studies, revealing similar intensity distributions whether targets were detected alone or together, suggesting that there is not a significant crowding effect (either antagonistic or synergistic).

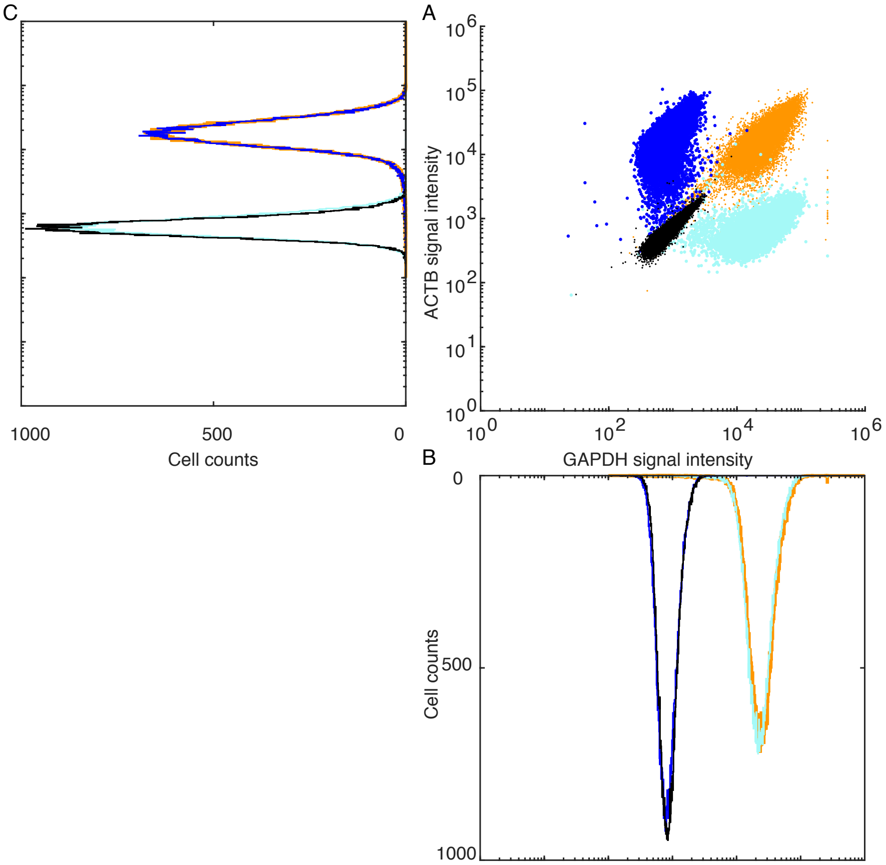

**Figure S23.** Comparison of signal intensity distributions for individual and multiplexed floHCR of *GAPDH* and *ACTB*. (A) Raw single-cell fluorescence intensity scatter plots: *GAPDH* channel vs *ACTB* channel. (B) Single-cell fluorescence intensity histogram for *GAPDH* channel. (C) Single-cell fluorescence intensity histogram for *ACTB* channel. Orange data: signal plus background for *GAPDH* and *ACTB* . Cyan data: signal plus background for *GAPDH* and autofluoresence for *ACTB* . Blue data: background for *ACTB* and signal plus autofluoresence for *GAPDH*. Black data: autofluoresence for *GAPDH* and *ACTB*. Sample: 65,000 HEK cells in suspension (WT).

##### S3.6.2 Redundant 2-channel detection of *GAPDH* endogenous target in HEK cells

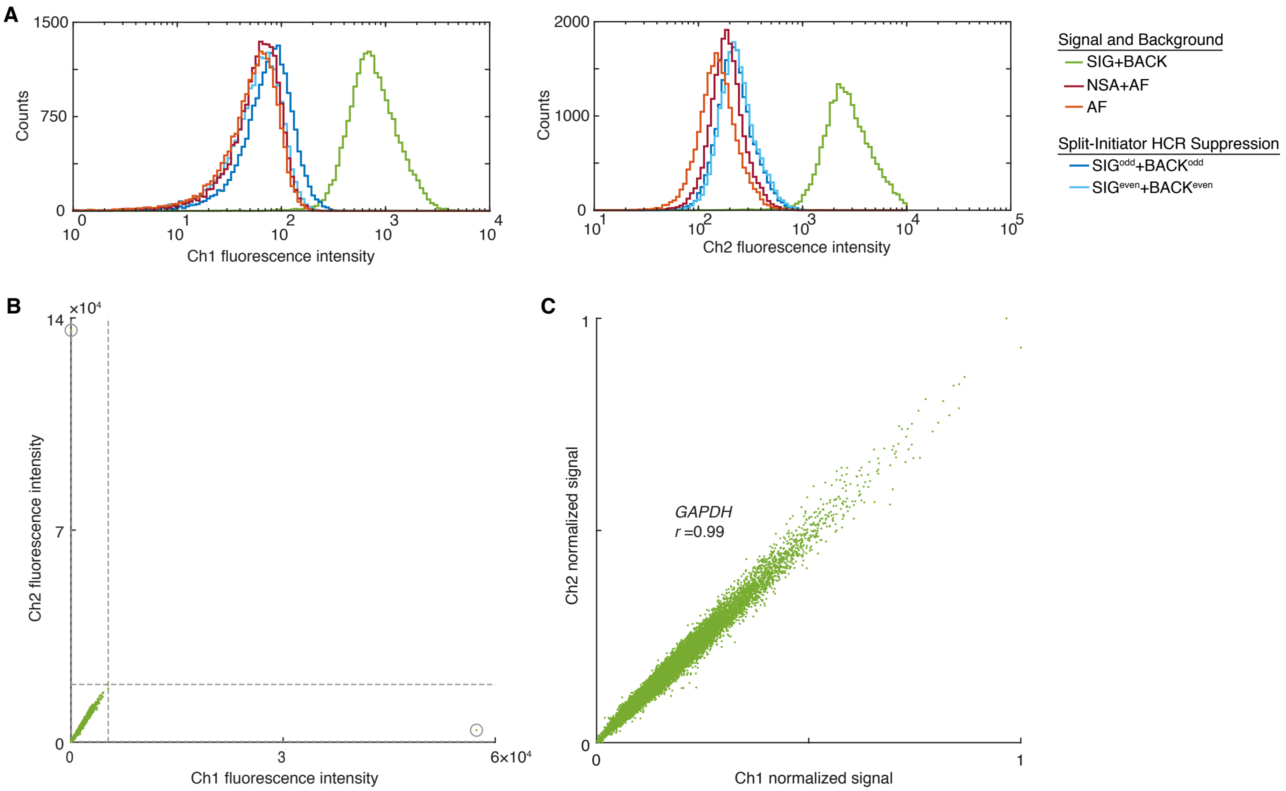

**Figure S24. Measurement of signal and background, background components, and split-initiator HCR suppression for redundant 2-channel detection of *GAPDH* endogenous target (cf. Figure 6B).** (A) Signal and background: use experiment of Type 1a in Table S8A to measure SIG+BACK (even + odd probes, hairpins). Background components: use experiment of Type 2 in Table S8B (no probes, with hairpins) to measure NSA+AF; use experiment of Type 3 (no probes, no hairpins) to measure AF. Split-initiator HCR suppression: use experiment of Type 4a in Table S8C (odd probes, hairpins) to measure $\text{SIG}^{\text{odd}}$+$\text{BACK}^{\text{odd}}$ (odd probes, hairpins); use experiment of Type 5a in Table S8C to measure $\text{SIG}^{\text{even}}$+$\text{BACK}^{\text{even}}$ (even probes, hairpins). Single-cell fluorescence intensity histograms for Ch1 and Ch2. (B) Raw single-cell fluorescence intensity scatter plots representing signal plus background. Dashed lines represent BOT and TOP values (Table S20) used to normalize data for panel C using methods of of Section S1.7.4 (outliers excluded from normalized scatter plots marked with circles). (C) Normalized single-cell fluorescence intensity scatter plots representing estimated normalized signal (Pearson correlation coefficient, $r$). Protocol: in situ HCR v3.0 (Section S2.3). Probe sets: 10 split-initiator probe pairs per channel. Amplifiers: B5-Alexa488 (Ch1) and B4-Alexa594 (Ch2). Sample: 20,000 HEK cells in suspension (WT).

| | Quantity | Channel | | Reagents | | Cell type |
|---|---|---|---|---|---|---|
| | | Ch1: B5-Alexa488 | Ch2: B4-Alexa594 | Probes | Hairpins | |
| **A** | SIG+BACK | 870 ± 5 | 3105 ± 14 | odd + even | ✓ | WT |
| | SIG | 808 ± 5 | 2904 ± 14 | | | |
| | SIG/BACK | 12.86 ± 0.08 | 14.43 ± 0.08 | | | |
| **B** | NSA+AF | 62.8 ± 0.2 | 201.2 ± 0.6 | | ✓ | WT |
| | AF | 57.4 ± 0.2 | 166.5 ± 0.5 | | ✓ | WT |
| | NSA | 5.4 ± 0.3 | 34.7 ± 0.8 | | | |
| **C** | $SIG^{odd}$+$BACK^{odd}$ | 82.8 ± 0.3 | 244.5 ± 0.8 | odd | ✓ | WT |
| | $SIG^{even}$+$BACK^{even}$ | 63.0 ± 0.2 | 256.6 ± 0.8 | even | ✓ | WT |
| | $SIG^{odd}$ | 20.0 ± 0.4 | 43 ± 1 | | | |
| | $SIG^{even}$ | 0.3 ± 0.3 | 55 ± 1 | | | |
| | SIG/$SIG^{odd}$ | 40.4 ± 0.8 | 67 ± 2 | | | |
| | SIG/$SIG^{even}$ | 3229 ± 4 | 52 ± 1 | | | |
| **D** | BOT | 62.8 | 201.2 | | | |
| | TOP | 5265.2 | 19 049.4 | | | |

**Table S20. Estimated signal-to-background, background components, and split-initiator HCR suppression for redundant 2-channel detection of *GAPDH* endogenous target (cf. Figure 6B).** (A) Signal-to-background (SIG/BACK). (B) Background components (AF, NSA). (C) Split-initiator HCR suppression (SIG/$SIG^{odd}$, SIG/$SIG^{even}$). The signal estimates SIG, $SIG^{odd}$ and $SIG^{even}$ are calculated using the background approximation BACK = $BACK^{odd}$ = $BACK^{even}$ $\approx$ NSA+AF. Mean $\pm$ standard error, $N = 20,000$ cells. Analysis based on single-cell intensities of Figure S24 using methods of Section S1.7.3. (D) BOT and TOP values used to calculate normalized single-cell intensities for scatter plots of Figures 6B and S24C using methods of Section S1.7.4.

##### S3.6.3 Redundant 2-channel detection of *PGK1* endogenous target in HEK cells

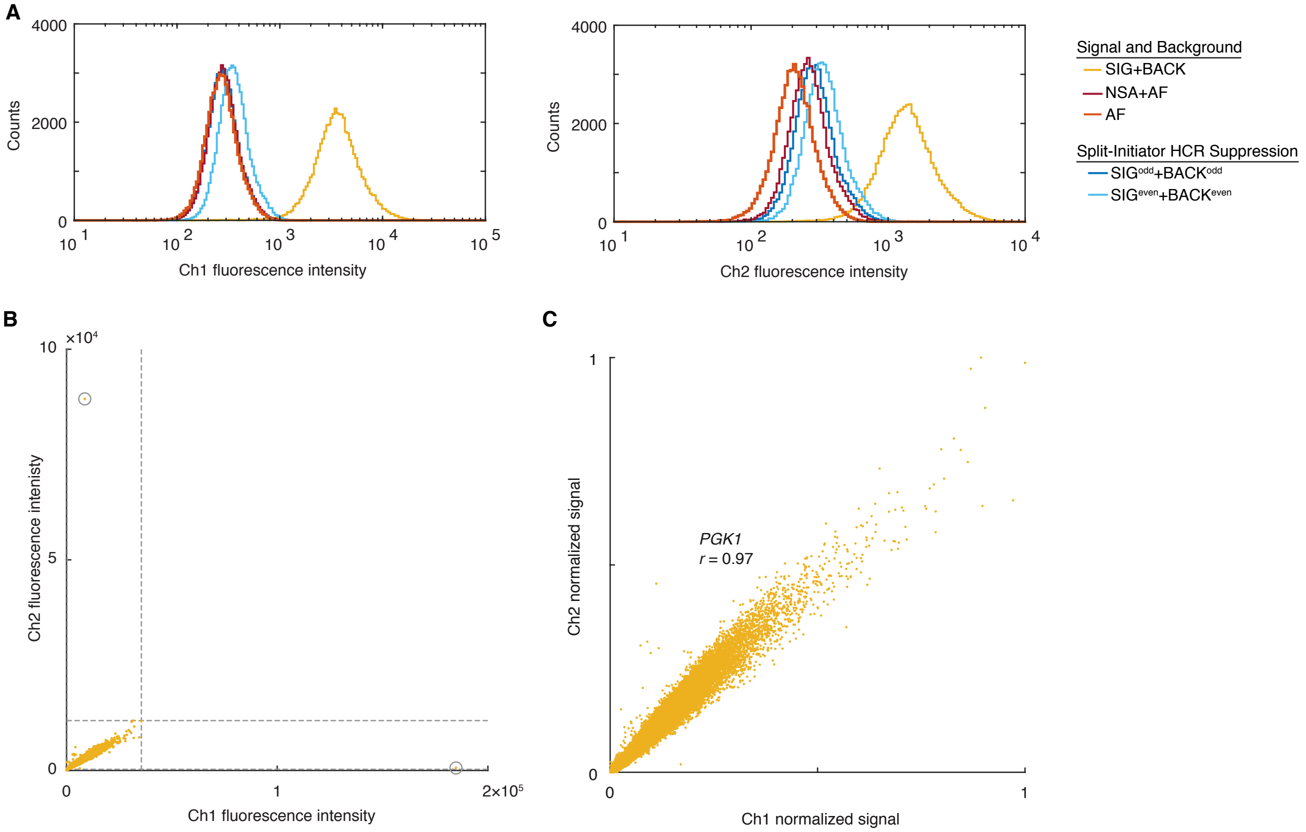

**Figure S25. Measurement of signal and background, background components, and split-initiator HCR suppression for redundant 2-channel detection of *PGK1* endogenous target.** (A) Signal and background: use experiment of Type 1a in Table S8A to measure SIG+BACK (even + odd probes, hairpins). Background components: use experiment of Type 2 in Table S8B (no probes, with hairpins) to measure NSA+AF; use experiment of Type 3 (no probes, no hairpins) to measure AF. Split-initiator HCR suppression: use experiment of Type 4a in Table S8C (odd probes, hairpins) to measure $\mathrm{SIG}^{\mathrm{odd}}+\mathrm{BACK}^{\mathrm{odd}}$ (odd probes, hairpins); use experiment of Type 5a in Table S8C to measure $\mathrm{SIG}^{\mathrm{even}}+\mathrm{BACK}^{\mathrm{even}}$ (even probes, hairpins). Single-cell fluorescence intensity histograms for Ch1 and Ch2. (B) Raw single-cell fluorescence intensity scatter plots representing signal plus background. Dashed lines represent BOT and TOP values (Table S21) used to normalize data for panel C using methods of of Section S1.7.4 (outliers excluded from normalized scatter plots marked with circles). (C) Normalized single-cell fluorescence intensity scatter plots representing estimated normalized signal (Pearson correlation coefficient, $r$). Protocol: in situ HCR v3.0 (Section S2.3). Probe sets: 18 split-initiator probe pairs per channel. Amplifiers: B1-Alexa488 (Ch1) and B2-Alexa594 (Ch2). Sample: 24,000 HEK cells in suspension (WT).

| | Quantity | Channel | | Reagents | | Cell type |
|---|---|---|---|---|---|---|
| | | Ch1: B1-Alexa488 | Ch2: B2-Alexa594 | Probes | Hairpins | |
| **A** | SIG+BACK | 4145 ± 11 | 1528 ± 4 | odd + even | ✓ | WT |
| | SIG | 3843 ± 11 | 1248 ± 4 | | | |
| | SIG/BACK | 12.72 ± 0.04 | 4.47 ± 0.02 | | | |
| **B** | NSA+AF | 302.1 ± 0.5 | 279.4 ± 0.5 | | ✓ | WT |
| | AF | 289.3 ± 0.5 | 220.1 ± 0.4 | | ✓ | WT |
| | NSA | 12.9 ± 0.7 | 59.3 ± 0.6 | | | |
| **C** | $SIG^{odd}+BACK^{odd}$ | 301.3 ± 0.5 | 309.5 ± 0.6 | odd | ✓ | WT |
| | $SIG^{even}+BACK^{even}$ | 380.5 ± 0.6 | 374 ± 7 | even | ✓ | WT |
| | $SIG^{odd}$ | < 0.7 | 30.1 ± 0.8 | | | |
| | $SIG^{even}$ | 78.4 ± 0.8 | 94 ± 7 | | | |
| | $SIG/SIG^{odd}$ | >5000 | 41 ± 1 | | | |
| | $SIG/SIG^{even}$ | 49.0 ± 0.5 | 13 ± 1 | | | |
| **D** | BOT | 302.1 | 279.4 | | | |
| | TOP | 35 538.8 | 11 847.7 | | | |

**Table S21. Estimated signal-to-background, background components, and split-initiator HCR suppression for redundant 2-channel detection of *PGK1* endogenous target.** (A) Signal-to-background (SIG/BACK). (B) Background components (AF, NSA). (C) Split-initiator HCR suppression ($SIG/SIG^{odd}$, $SIG/SIG^{even}$). The signal estimates SIG, $SIG^{odd}$ and $SIG^{even}$ are calculated using the background approximation BACK = $BACK^{odd}$ = $BACK^{even}$ $\approx$ NSA+AF. Mean ± standard error, $N = 54,000$ cells. Analysis based on single-cell intensities of Figure S25 using methods of Section S1.7.3. (D) BOT and TOP values used to calculate normalized single-cell intensities for scatter plots of Figure S25C using methods of Section S1.7.4.

##### S3.6.4 Redundant 2-channel detection of *fusA* endogenous target in *E. coli*

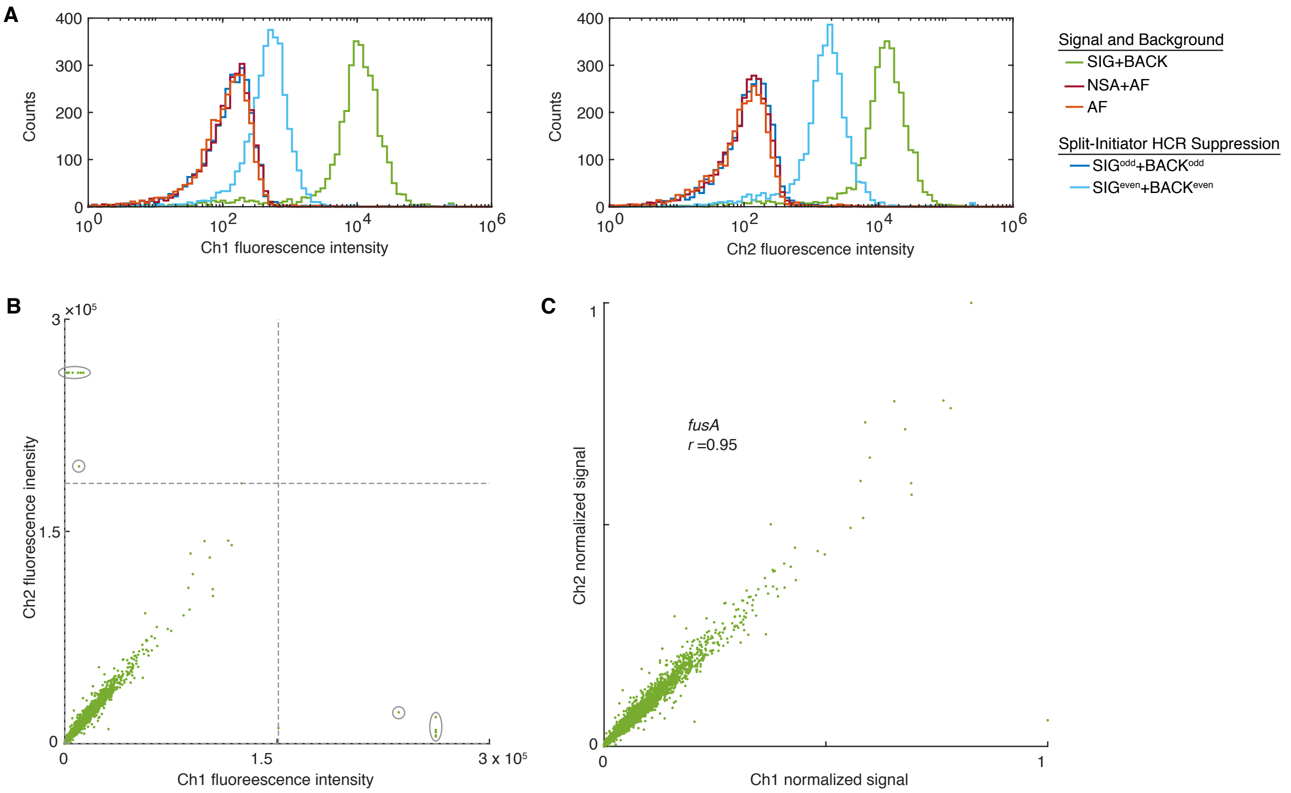

**Figure S26. Measurement of signal and background, background components, and split-initiator HCR suppression for redundant 2-channel detection of *fusA* endogenous target (cf. Figure 6B).** (A) Signal and background: use experiment of Type 1a in Table S8A to measure SIG+BACK (even + odd probes, hairpins). Background components: use experiment of Type 2 in Table S8B (no probes, with hairpins) to measure NSA+AF; use experiment of Type 3 (no probes, no hairpins) to measure AF. Split-initiator HCR suppression: use experiment of Type 4a in Table S8C (odd probes, hairpins) to measure SIG$^{odd}$+BACK$^{odd}$ (odd probes, hairpins); use experiment of Type 5a in Table S8C to measure SIG$^{even}$+BACK$^{even}$ (even probes, hairpins). Single-cell fluorescence intensity histograms for Ch1 and Ch2. (B) Raw single-cell fluorescence intensity scatter plots representing signal plus background. Dashed lines represent BOT and TOP values (Table S22) used to normalize data for panel C using methods of of Section S1.7.4 (outliers excluded from normalized scatter plots marked with ellipses). (C) Normalized single-cell fluorescence intensity scatter plots representing estimated normalized signal (Pearson correlation coefficient, $r$). Protocol: in situ HCR v3.0 (Section S2.4). Probe sets: 18 split-initiator probe pairs per channel. Amplifiers: B3-Alexa488 (Ch1) and B2-Alexa594 (Ch2). Sample: 3,400 *E. coli* in suspension (WT).

| | Quantity | Channel | | | | Reagents | | Cell type |
|---|---|---|---|---|---|---|---|---|
| | | Ch1: B3-Alexa488 | | Ch2: B2-Alexa594 | | Probes | Hairpins | |
| **A** | SIG+BACK | 13 100 | ± 300 | 17 500 | ± 300 | odd + even | ✓ | WT |
| | SIG | 13 000 | ± 300 | 15 700 | ± 300 | | | |
| | SIG/BACK | 99 | ± 3 | 135 | ± 7 | | | |
| **B** | NSA+AF | 130 | ± 3 | 116 | ± 5 | | ✓ | WT |
| | AF | 126 | ± 7 | 120 | ± 10 | | ✓ | WT |
| | NSA | $<7$ | | $<14$ | | | | |
| **C** | $\text{SIG}^{\text{odd}}$+$\text{BACK}^{\text{odd}}$ | 500 | ± 100 | 400 | ± 100 | odd | ✓ | WT |
| | $\text{SIG}^{\text{even}}$+$\text{BACK}^{\text{even}}$ | 1100 | ± 200 | 2600 | ± 200 | even | ✓ | WT |
| | $\text{SIG}^{\text{odd}}$ | 300 | ± 100 | 300 | ± 100 | | | |
| | $\text{SIG}^{\text{even}}$ | 900 | ± 200 | 2500 | ± 200 | | | |
| | $\text{SIG/SIG}^{\text{odd}}$ | 40 | ± 20 | 50 | ± 20 | | | |
| | $\text{SIG/SIG}^{\text{even}}$ | 14 | ± 3 | 6.2 | ± 0.6 | | | |
| **D** | BOT | 130.3 | | 116 | | | | |
| | TOP | 150 951.6 | | 183 947.3 | | | | |

**Table S22. Estimated signal-to-background, background components, and split-initiator HCR suppression for redundant 2-channel detection of *fusA* endogenous target (cf. Figure 6B).** (A) Signal-to-background (SIG/BACK). (B) Background components (AF, NSA). (C) Split-initiator HCR suppression ($\text{SIG/SIG}^{\text{odd}}$, $\text{SIG/SIG}^{\text{even}}$). The signal estimates SIG, $\text{SIG}^{\text{odd}}$ and $\text{SIG}^{\text{even}}$ are calculated using the background approximation BACK = $\text{BACK}^{\text{odd}}$ = $\text{BACK}^{\text{even}}$ $\approx$ NSA+AF. Mean ± standard error, $N = 3,400$ cells. Analysis based on single-cell intensities of Figure S26 using methods of Section S1.7.3. (D) BOT and TOP values used to calculate normalized single-cell intensities for scatter plots of Figures 6B and S26C using methods of Section S1.7.4.

##### S3.6.5 Multiplexed 2-channel detection of *GAPDH* and *PGK1* endogenous targets in HEK cells

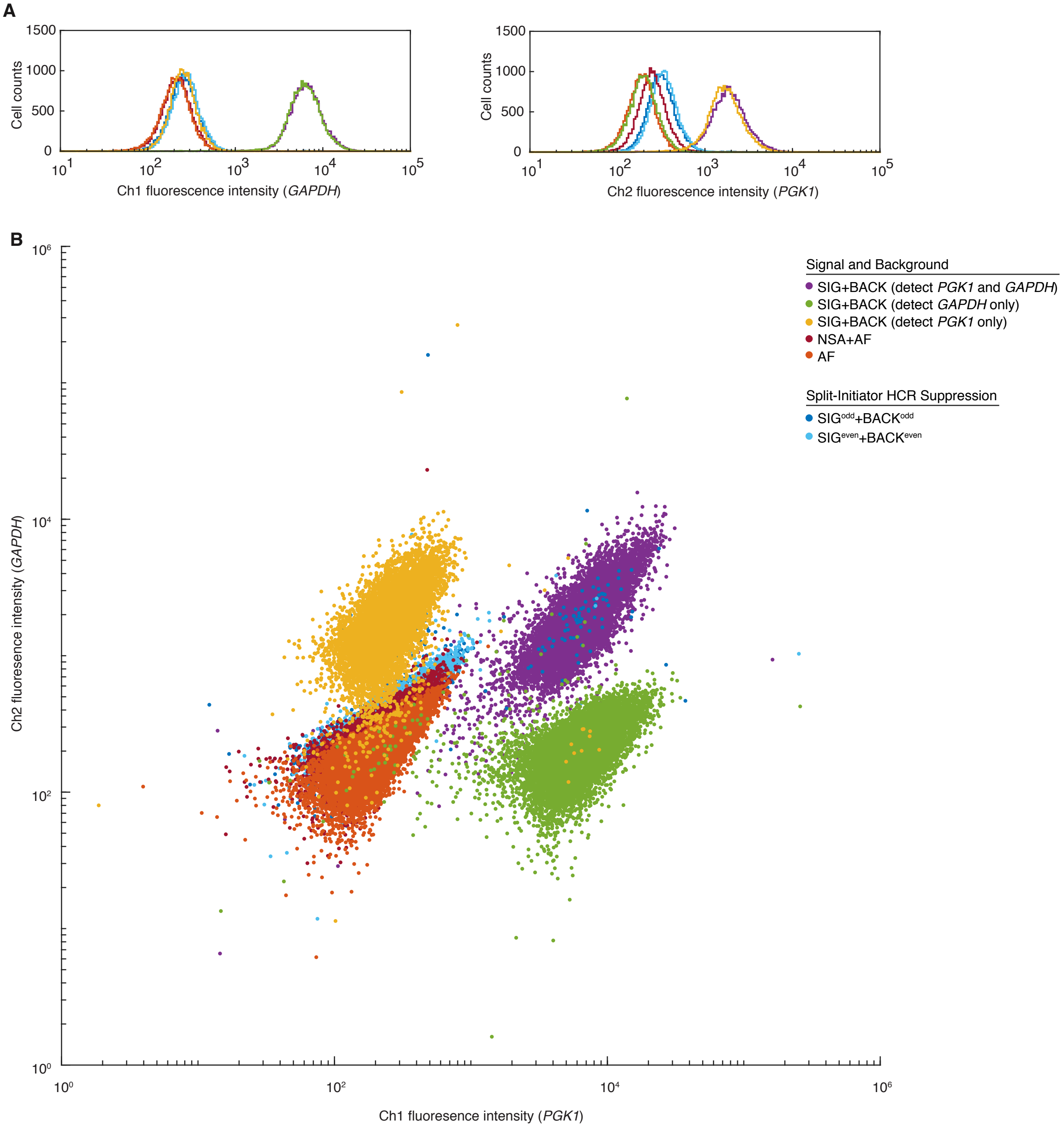

**Figure S27. Measurement of signal and background, background components, and split-initiator HCR suppression for multiplexed 2-channel detection of *GAPDH* and *PGK1* endogenous targets.** (A) Single-cell fluorescence intensity histograms for Ch1 and Ch2. (B) Raw single-cell fluorescence intensity scatter plots for Ch1 vs Ch2. Signal and background: use experiment of Type 1a in Table S8A to measure SIG+BACK (even + odd probes, hairpins). Background components: use experiment of Type 2 in Table S8B (no probes, with hairpins) to measure NSA+AF; use experiment of Type 3 (no probes, no hairpins) to measure AF. Split-initiator HCR suppression: use experiment of Type 4a in Table S8C (odd probes, hairpins) to measure $SIG^{odd}$+$BACK^{odd}$ (odd probes, hairpins); use experiment of Type 5a in Table S8C to measure $SIG^{even}$+$BACK^{even}$ (even probes, hairpins). Protocol: in situ HCR v3.0 (Section S2.3). Ch1: target mRNA *GAPDH*, probe set with 10 split-initiator probe pairs, amplifier B4-Alexa488. Ch2: target mRNA *PGK1*, probe set with 18 split-initiator probe pairs, amplifier B2-Alexa594. Sample: 18,000 HEK cells in suspension (WT).

| | | Channel | | Reagents | | |
|---|---|---|---|---|---|---|
| | Quantity | Ch1: B4-Alexa488 | Ch2: B2-Alexa594 | Probes | Hairpins | Cell type |
| **A** | SIG+BACK | $6980 \pm 30$ | $2073 \pm 8$ | odd + even | ✓ | WT |
| | SIG | $6760 \pm 30$ | $1806 \pm 8$ | | | |
| | SIG/BACK | $29.6 \pm 0.1$ | $6.77 \pm 0.05$ | | | |
| **B** | NSA+AF | $228.2 \pm 0.7$ | $266.6 \pm 1.5$ | | ✓ | WT |
| | AF | $219.7 \pm 0.6$ | $198.4 \pm 0.6$ | | ✓ | WT |
| | NSA | $8.5 \pm 0.9$ | $68 \pm 2$ | | | |
| **C** | $SIG^{odd}+BACK^{odd}$ | $301 \pm 5$ | $355 \pm 9$ | odd | ✓ | WT |
| | $SIG^{even}+BACK^{even}$ | $303 \pm 14$ | $362.8 \pm 1.2$ | even | ✓ | WT |
| | $SIG^{odd}$ | $73 \pm 5$ | $88 \pm 9$ | | | |
| | $SIG^{even}$ | $74 \pm 14$ | $96 \pm 2$ | | | |
| | $SIG/SIG^{odd}$ | $93 \pm 6$ | $21 \pm 2$ | | | |
| | $SIG/SIG^{even}$ | $91 \pm 17$ | $18.8 \pm 0.4$ | | | |

**Table S23. Estimated signal-to-background, background components, and split-initiator HCR suppression for multiplexed 2-channel detection of *GAPDH* and *PGK1* endogenous targets.** (A) Signal-to-background (SIG/BACK). (B) Background components (AF, NSA). (C) Split-initiator HCR suppression ($SIG/SIG^{odd}$, $SIG/SIG^{even}$). The signal estimates SIG, $SIG^{odd}$ and $SIG^{even}$ are calculated using the background approximation BACK = $BACK^{odd}$ = $BACK^{even}$ $\approx$ NSA+AF. Mean $\pm$ standard error, $N = 18,000$ cells. Analysis based on single-cell intensities of Figure S27 using methods of Section S1.7.3.

##### S3.6.6 Multiplexed 2-channel detection of *fusA* and *icd* endogenous targets in *E. coli*

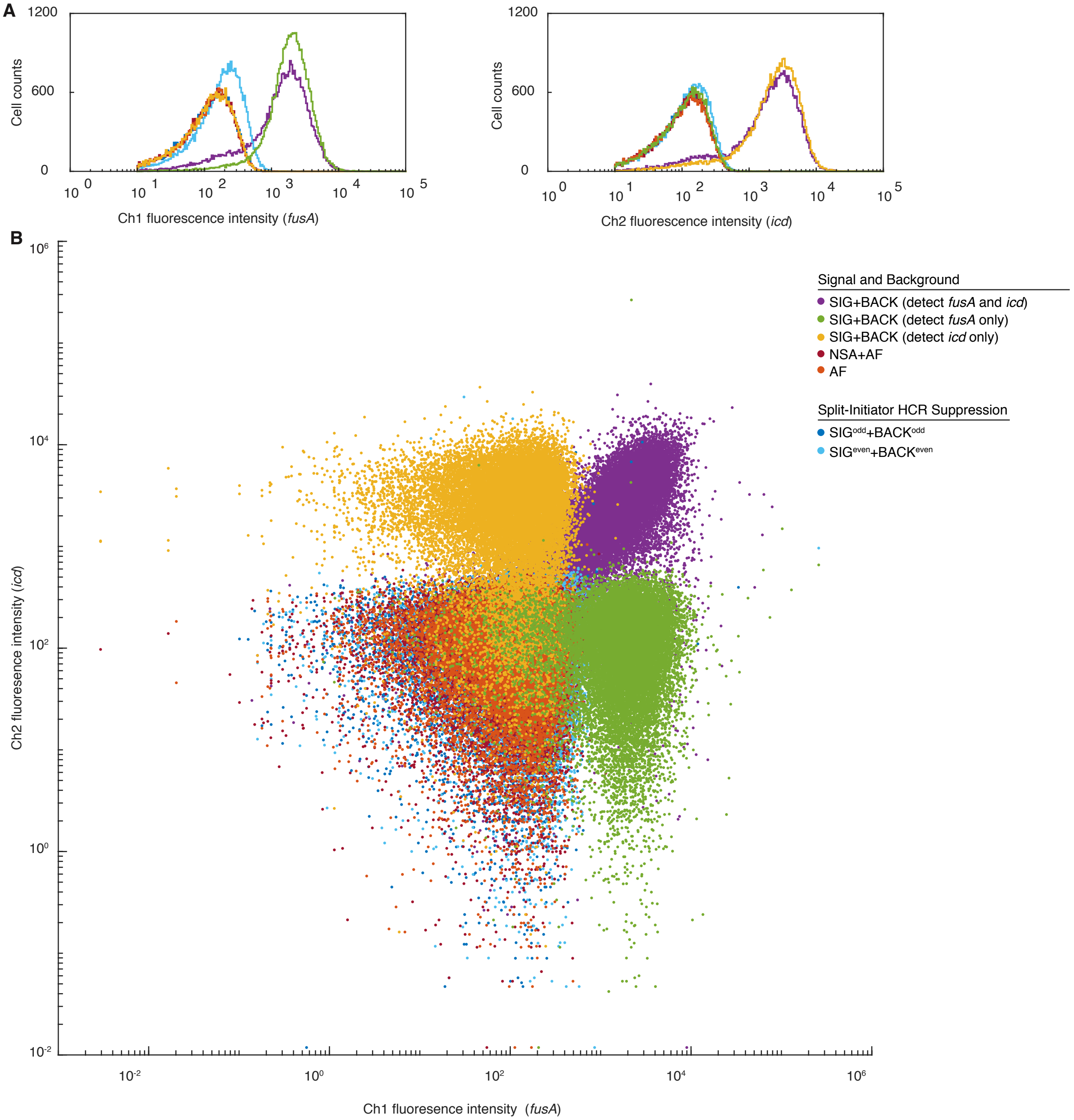

**Figure S28. Measurement of signal and background, background components, and split-initiator HCR suppression for multiplexed 2-channel detection of *fusA* and *icd* endogenous targets.** (A) Single-cell fluorescence intensity histograms for Ch1 and Ch2. (B) Raw single-cell fluorescence intensity scatter plots for Ch1 vs Ch2. Signal and background: use experiment of Type 1a in Table S8A to measure SIG+BACK (even + odd probes, hairpins). Background components: use experiment of Type 2 in Table S8B (no probes, with hairpins) to measure NSA+AF; use experiment of Type 3 (no probes, no hairpins) to measure AF. Split-initiator HCR suppression: use experiment of Type 4a in Table S8C (odd probes, hairpins) to measure $\text{SIG}^{\text{odd}}+\text{BACK}^{\text{odd}}$ (odd probes, hairpins); use experiment of Type 5a in Table S8C to measure $\text{SIG}^{\text{even}}+\text{BACK}^{\text{even}}$ (even probes, hairpins). Protocol: in situ HCR v3.0 (Section S2.4). Ch1: target mRNA *fusA*, probe set with 18 split-initiator probe pairs, amplifier B3-Alexa488. Ch2: target mRNA *icd*, probe set with 20 split-initiator probe pairs, amplifier B1-Alexa594. Sample: 35,000 *E. coli* DH5$\alpha$ in suspension.

| | Quantity | Channel: Ch1: B3-Alexa488 | | Channel: Ch2: B1-Alexa594 | | Reagents: Probes | Reagents: Hairpins | Cell type |
|---|---|---|---|---|---|---|---|---|
| **A** | SIG+BACK | 1756 | ± 9 | 2533 | ± 12 | odd + even | ✓ | WT |
| | SIG | 1673 | ± 9 | 2470 | ± 12 | | | |
| | SIG/BACK | 20.1 | ± 0.2 | 38.9 | ± 0.5 | | | |
| **B** | NSA+AF | 83.2 | ± 0.7 | 63.5 | ± 0.7 | | ✓ | WT |
| | AF | 82.3 | ± 0.7 | 60.1 | ± 0.7 | | ✓ | WT |
| | NSA | 1 | ± 1 | 3 | ± 1 | | | |
| **C** | $SIG^{odd}$+$BACK^{odd}$ | 86 | ± 1 | 67 | ± 1 | odd | ✓ | WT |
| | $SIG^{even}$+$BACK^{even}$ | 180 | ± 8 | 93 | ± 1 | even | ✓ | WT |
| | $SIG^{odd}$ | 3 | ± 1 | 3 | ± 1 | | | |
| | $SIG^{even}$ | 97 | ± 8 | 29 | ± 1 | | | |
| | SIG/$SIG^{odd}$ | 600 | ± 300 | 800 | ± 300 | | | |
| | SIG/$SIG^{even}$ | 17 | ± 8 | 85 | ± 1 | | | |

**Table S24. Estimated signal-to-background, background components, and split-initiator HCR suppression for multiplexed 2-channel detection of *fusA* and *icd* endogenous targets.** (A) Signal-to-background (SIG/BACK). (B) Background components (AF, NSA). (C) Split-initiator HCR suppression (SIG/$SIG^{odd}$, SIG/$SIG^{even}$). The signal estimates SIG, $SIG^{odd}$ and $SIG^{even}$ are calculated using the background approximation BACK = $BACK^{odd}$ = $BACK^{even}$ $\approx$ NSA+AF. Mean $\pm$ standard error, $N = 35,000$ cells. Analysis based on single-cell intensities of Figure S28 using methods of Section S1.7.3.

#### S3.7 dHCR imaging: digital mRNA absolute quantitation in an anatomical context (cf. Figure 7)

##### S3.7.1 Redundant 2-channel detection of single *BRAF* mRNAs in HEK cells using in situ HCR v3.0

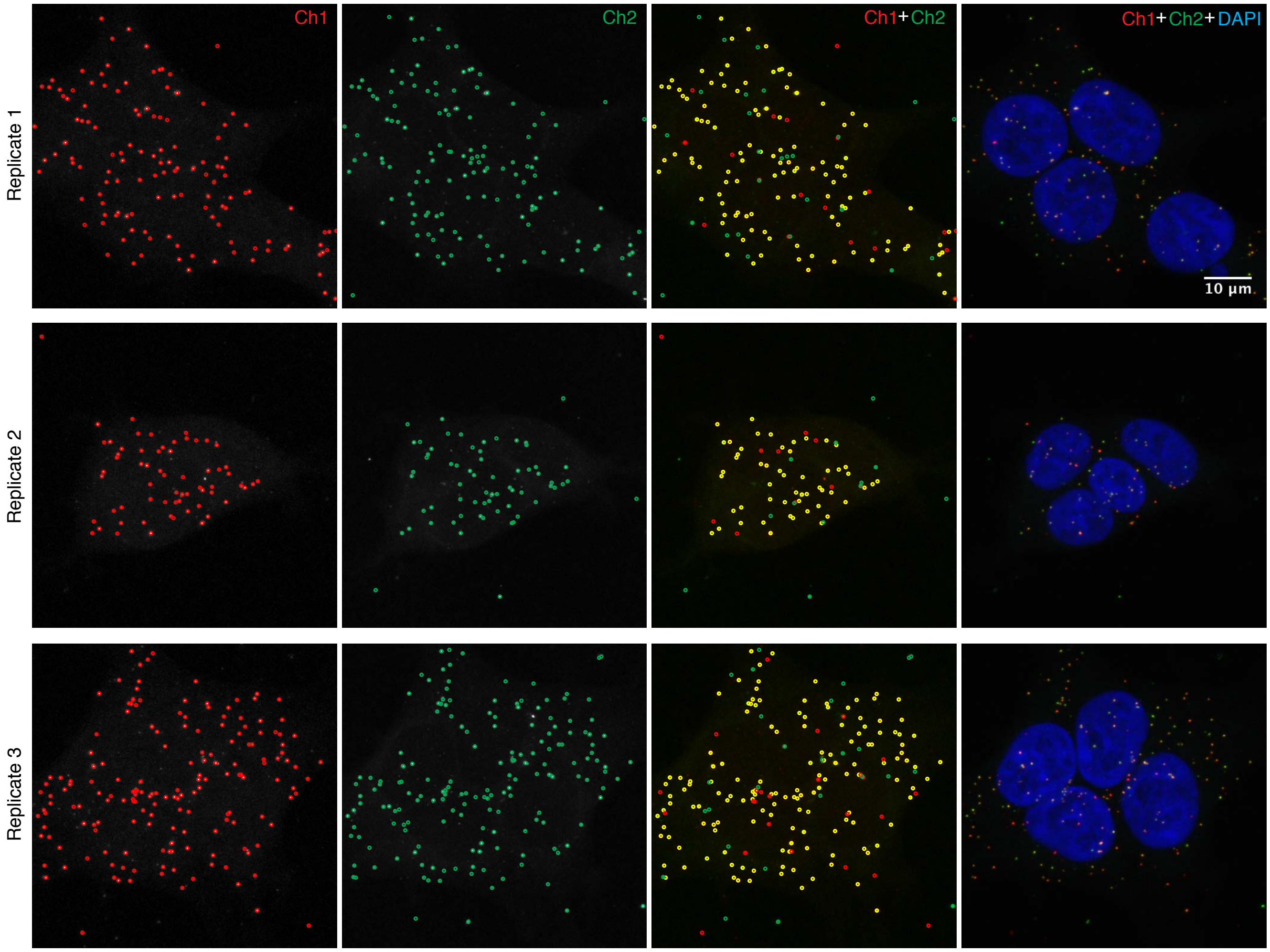

**Figure S29. Redundant 2-channel detection of single *BRAF* mRNAs in HEK cells using in situ HCR v3.0 (cf. Figure 7A).** Confocal images: individual channels and merge (without and with DAPI nuclear stain). Maximum intensity projection in the axial direction over 7.14 $\mu$m (17 focal planes). Pixel size: 0.062×0.062 $\mu$m. Probe sets: 23 split-initiator probe pairs per channel. Amplifiers: B3-Alexa647 (Ch1) and B4-Alexa546 (Ch2). Red circles: dots detected in Ch1. Green circles: dots detected in Ch2. Yellow circles: dots detected in both channels.

| | Dots | | Colocalized dots | Colocalization fractions | |
|---|---|---|---|---|---|
| | $N_1$ | $N_2$ | $N_{12}$ | $C_1$ | $C_2$ |
| Replicate 1 | 129 | 136 | 110 | 0.85 | 0.81 |
| Replicate 2 | 63 | 65 | 53 | 0.84 | 0.82 |
| Replicate 3 | 170 | 170 | 144 | 0.85 | 0.85 |
| Mean | | | | 0.85 ± 0.003 | 0.82 ± 0.01 |

**Table S25. Dot colocalization fractions for redundant 2-channel detection of single *BRAF* mRNAs in HEK cells using in situ HCR v3.0 (cf. Figure 7A).** Mean ± standard error, $N = 3$ replicate samples. Analysis based on the images of Figure S29 using the methods of Section S1.6.6 with the settings in Table S6.

##### S3.7.2 Redundant 2-channel detection of single *Dmbx1* mRNAs in whole-mount chicken embryos using in situ HCR v3.0

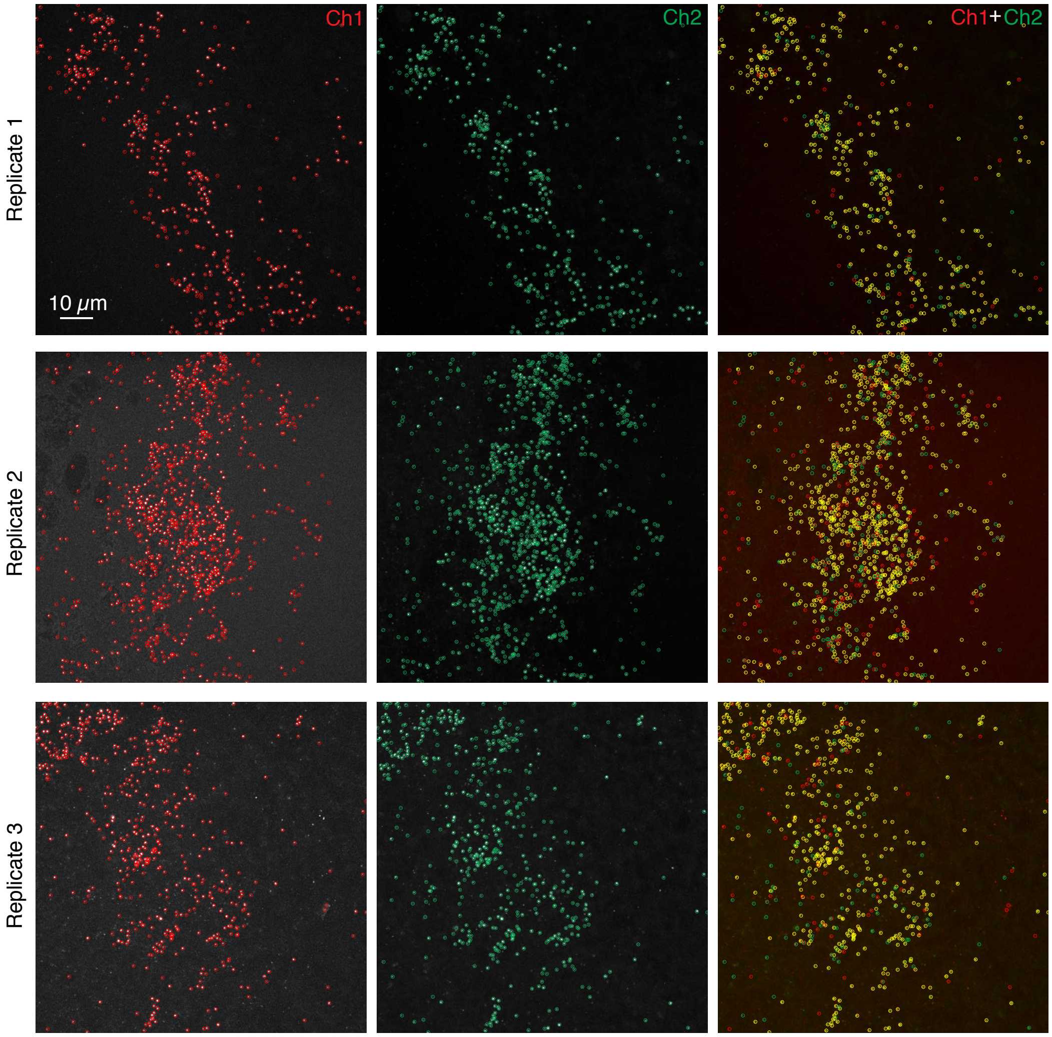

**Figure S30. Redundant 2-channel detection of single *Dmbx1* mRNAs in whole-mount chicken embryos using in situ HCR v3.0 (cf. Figure 7B).** Confocal images: individual channels and merge. Maximum intensity projection in the axial direction over 5.04-23.52 $\mu$m (12, 54, 56 focal planes for replicates 1, 2, 3 depending on sample thickness). Pixel size: $0.099 \times 0.099$ $\mu$m. Probe sets: 25 split-initiator probe pairs per channel. Amplifiers: B2-Alexa647 (Ch1) and B1-Alexa594 (Ch2). Embryos fixed stage HH 8. Red circles: dots detected in Ch1. Green circles: dots detected in Ch2. Yellow circles: dots detected in both channels.

| | Dots | | Colocalized dots | Colocalization fractions | |
|---|---|---|---|---|---|
| | $N_1$ | $N_2$ | $N_{12}$ | $C_1$ | $C_2$ |
| Replicate 1 | 403 | 417 | 364 | 0.90 | 0.87 |
| Replicate 2 | 992 | 990 | 794 | 0.80 | 0.80 |
| Replicate 3 | 526 | 539 | 448 | 0.85 | 0.83 |
| Mean | | | | $0.85 \pm 0.05$ | $0.84 \pm 0.04$ |

**Table S26. Dot colocalization fractions for redundant 2-channel detection of single *Dmbx1* mRNAs in whole-mount chicken embryos using in situ HCR v3.0 (cf. Figure 7B).** Mean $\pm$ standard error, $N = 3$ replicate embryos. Analysis based on the images of Figure S30 using the methods of Section S1.6.6 with the settings in Table S6.

##### S3.7.3 Redundant 2-channel detection of single *kdrl* mRNAs in whole-mount zebrafish embryos using in situ HCR v2.0 (Shah *et al.*, 2016)

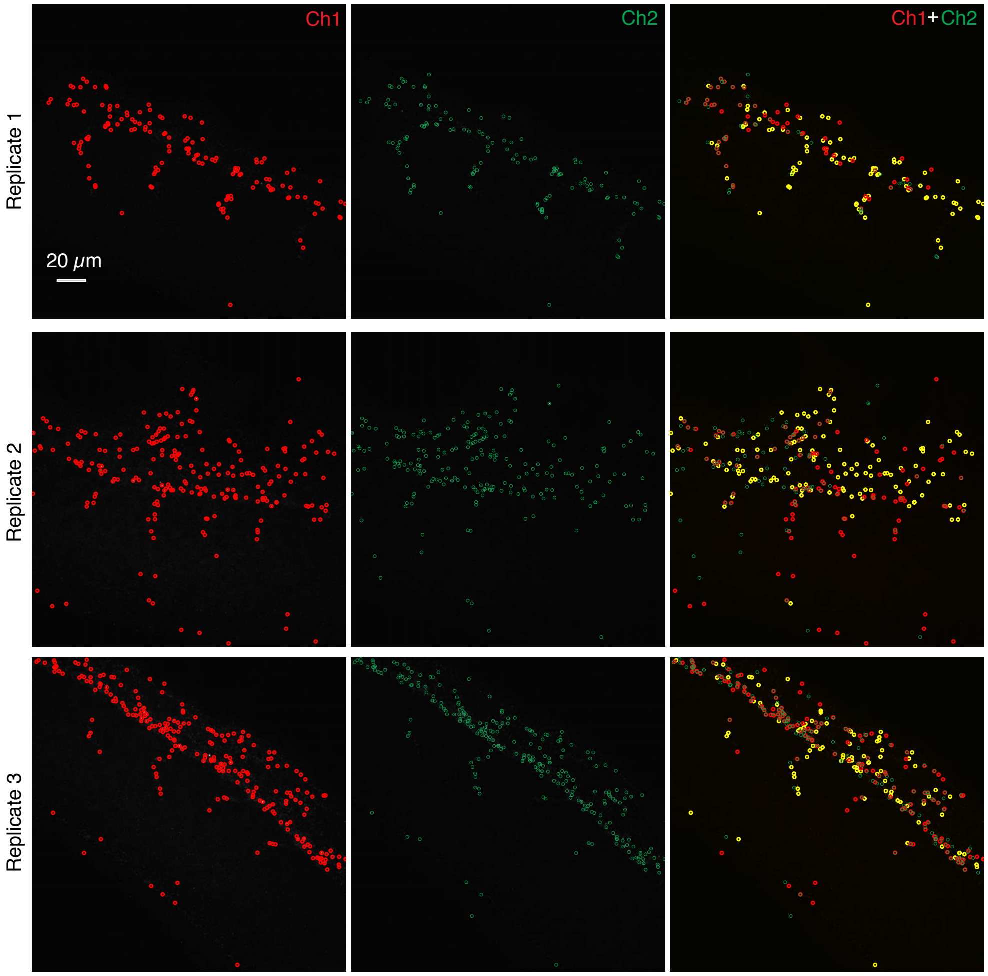

**Figure S31. Redundant 2-channel detection of single *kdrl* mRNAs in whole-mount zebrafish embryos using in situ HCR v2.0 (cf. Figure 7).** Spinning disk confocal images: individual channels and merge from Shah *et al.* (2016). Maximum intensity projection in the axial direction over 13 $\mu$m (39 focal planes). Pixel size: 0.217×0.217 $\mu$m. Probe sets: 39 standard probes per channel, each incorporating a 30-nt target-binding domain and a full HCR initiator. Amplifiers: B3-Alexa647 (Ch1) and B2-Alexa546 (Ch2). Embryos fixed 27 hpf. Red circles: dots detected in Ch1. Green circles: dots detected in Ch2. Yellow circles: dots detected in both channels.

| | Dots | | Colocalized dots | Colocalization fractions | |
|---|---|---|---|---|---|
| | $N_1$ | $N_2$ | $N_{12}$ | $C_1$ | $C_2$ |
| Replicate 1 | 139 | 132 | 79 | 0.57 | 0.60 |
| Replicate 2 | 220 | 215 | 113 | 0.51 | 0.52 |
| Replicate 3 | 243 | 245 | 91 | 0.37 | 0.37 |
| Mean | | | | 0.49 ± 0.06 | 0.50 ± 0.07 |

**Table S27. Dot colocalization fractions for redundant 2-channel detection of single *kdrl* mRNAs in whole-mount zebrafish embryos using in situ HCR v3.0 (cf. Figure 7).** Mean ± standard error, $N = 3$ replicate embryos. Analysis based on the images of Figure S31 using the methods of Section S1.6.6 with the settings in Table S6.

#### S4 Probe sequences

Target mRNA sequences were obtained from the National Center for Biotechnology Information (NCBI) (McEntyre & Ostell, 2002) with the exception of *d2eGFP* (pd2EGFP-1, Clontech, Cat. #6008-1). Spatial and temporal expression information for whole-mount chicken embryos were obtained from the Gallus Expression in Situ Hybridization Analysis (GEISHA) (Bell *et al.*, 2004; Darnell *et al.*, 2007). Within a given probe set, each DNA standard probe or split-initiator probe pair initiates the same DNA HCR amplifier. Sequences are listed $5'$ to $3'$. Probes are numbered consecutively moving along a target mRNA. For redundant detection experiments, two probe sets are used with each probe set taking alternating probe pairs from along the target (this leads to non-consecutive numbers within each probe set).

#### S4.1 Standard probes for Figures 3, S5, and S7

Organism: ***G. gallus domesticus***
Target mRNA: **SRY (sex determining region Y)-box 10 (*Sox10*)**
Probe set: **5, 10, or 20 probes (each carrying 2 HCR initiators)**
HCR amplifier: **B3-Alexa647**

| # | Initiator I1 | Spacer | Probe Sequence (50 nt) | Spacer | Initiator I2 |
|---|---|---|---|---|---|
| 1 | gTCCCTgCCTCTATATCTCCACTCAACTTTAACCCg | TACAA | CATggACCCgTCACTCCATgTCTTgAgTCTTCCTCATCTAgAAggCCAAT | TAAAA | AAAgTCTAATCCgTCCCTgCCTCTATATCTCCACTC |
| 2 | gTCCCTgCCTCTATATCTCCACTCAACTTTAACCCg | TACAA | CCAgCAgggATCAAgATTCATgCATgTgTgAATCTTAggCAggACTgCTg | TAAAA | AAAgTCTAATCCgTCCCTgCCTCTATATCTCCACTC |
| 3 | gTCCCTgCCTCTATATCTCCACTCAACTTTAACCCg | TACAA | CgggCTATgAAATgAgAAAggCTAAggCTgACAgTgCAgTTCCTgAATCC | TAAAA | AAAgTCTAATCCgTCCCTgCCTCTATATCTCCACTC |
| 4 | gTCCCTgCCTCTATATCTCCACTCAACTTTAACCCg | TACAA | TTCACgTTTTCAgCAgACACAgTCAAATgCTggAggAgCAAggACCTggT | TAAAA | AAAgTCTAATCCgTCCCTgCCTCTATATCTCCACTC |
| 5 | gTCCCTgCCTCTATATCTCCACTCAACTTTAACCCg | TACAA | ATTggAACCACATCTgggTgTTggCAAgTgCATggTAgCTTTCTTggTgC | TAAAA | AAAgTCTAATCCgTCCCTgCCTCTATATCTCCACTC |
| 6 | gTCCCTgCCTCTATATCTCCACTCAACTTTAACCCg | TACAA | AgAggCggggAgAAAAgCTATAgCgTgCAgCTgTgAAAATCAgCAAggAA | TAAAA | AAAgTCTAATCCgTCCCTgCCTCTATATCTCCACTC |
| 7 | gTCCCTgCCTCTATATCTCCACTCAACTTTAACCCg | TACAA | ATAAAATCCATgCAggAAggggTgTgggATTAAACAgATgggACAggggg | TAAAA | AAAgTCTAATCCgTCCCTgCCTCTATATCTCCACTC |
| 8 | gTCCCTgCCTCTATATCTCCACTCAACTTTAACCCg | TACAA | gATggCgATAATgTgATgAACAAACgAgCAgTgATgTACACCCCATCggC | TAAAA | AAAgTCTAATCCgTCCCTgCCTCTATATCTCCACTC |
| 9 | gTCCCTgCCTCTATATCTCCACTCAACTTTAACCCg | TACAA | ATCCACgAgAgTATCTTTCCATCCTgAgTgAAAgTAggAgggAggTgCTg | TAAAA | AAAgTCTAATCCgTCCCTgCCTCTATATCTCCACTC |
| 10 | gTCCCTgCCTCTATATCTCCACTCAACTTTAACCCg | TACAA | ACCCgTTAgAAggTCCCACAACACATCTCTCTgATCAgTTgTCAggAgTC | TAAAA | AAAgTCTAATCCgTCCCTgCCTCTATATCTCCACTC |
| 11 | gTCCCTgCCTCTATATCTCCACTCAACTTTAACCCg | TACAA | CCggCgAgAggCAgTggTggTCTTCAgAACCCACTgggCTCATCTCCACC | TAAAA | AAAgTCTAATCCgTCCCTgCCTCTATATCTCCACTC |
| 12 | gTCCCTgCCTCTATATCTCCACTCAACTTTAACCCg | TACAA | CCTCgCCCTgCgTggCCTTgCCATTTTTCCgCCTCCgTggCTggTACTTg | TAAAA | AAAgTCTAATCCgTCCCTgCCTCTATATCTCCACTC |
| 13 | gTCCCTgCCTCTATATCTCCACTCAACTTTAACCCg | TACAA | TTCTTgTAgTgAgCCTggATAgAggCAgCCCCgCCAgCCTCCCCCTCCAC | TAAAA | AAAgTCTAATCCgTCCCTgCCTCTATATCTCCACTC |
| 14 | gTCCCTgCCTCTATATCTCCACTCAACTTTAACCCg | TACAA | TgCCCATCggACATgggTgACCCTTCCCCAggATgCCTgTggTCCAggTg | TAAAA | AAAgTCTAATCCgTCCCTgCCTCTATATCTCCACTC |
| 15 | gTCCCTgCCTCTATATCTCCACTCAACTTTAACCCg | TACAA | TTAggggTggTgggAggAgTgggAgggCCgTggCTCTgACCTgAAgAgTg | TAAAA | AAAgTCTAATCCgTCCCTgCCTCTATATCTCCACTC |
| 16 | gTCCCTgCCTCTATATCTCCACTCAACTTTAACCCg | TACAA | CCCCAAgggAACgCCCTTCTCgCTTggAgTCAgCTTTgCCTgCCTgCAgC | TAAAA | AAAgTCTAATCCgTCCCTgCCTCTATATCTCCACTC |
| 17 | gTCCCTgCCTCTATATCTCCACTCAACTTTAACCCg | TACAA | ggCCTgggTggCCggCgTgTCCgTTgggTggCAggTATTggTCAAATTCg | TAAAA | AAAgTCTAATCCgTCCCTgCCTCTATATCTCCACTC |
| 18 | gTCCCTgCCTCTATATCTCCACTCAACTTTAACCCg | TACAA | TCCATgCTgCTTggAgATCCAggCTgAgTgTCCACTggCCgCAgCCAggg | TAAAA | AAAgTCTAATCCgTCCCTgCCTCTATATCTCCACTC |
| 19 | gTCCCTgCCTCTATATCTCCACTCAACTTTAACCCg | TACAA | AACCCTgTgAAggCTgCAgCTCCTgTCCCAgTgCCCCCCTgTTCTCCCTC | TAAAA | AAAgTCTAATCCgTCCCTgCCTCTATATCTCCACTC |
| 20 | gTCCCTgCCTCTATATCTCCACTCAACTTTAACCCg | TACAA | gggTCCAgTCATAgCCgCTCAgCACCTggCTgACCgCTTCACggATgCAg | TAAAA | AAAgTCTAATCCgTCCCTgCCTCTATATCTCCACTC |

#### S4.2 Split-initiator probes for Figures 3, S6, S8, S9, and S10

Organism: ***G. gallus domesticus***
Target mRNA: **SRY (sex determining region Y)-box 10 (*Sox10*)**
Probe set: **5, 10, or 20 split-initiator probe pairs (each probe carries half an HCR initiator)**
HCR amplifier: **B3-Alexa647**

| Odd # | 1st Half of Initiator I1 | Spacer | Probe Sequence (25 nt) |
|---|---|---|---|
| 1 | gTCCCTgCCTCTATATCT | TT | gTCTTCCTCATCTAgAAggCCAATA |
| 3 | gTCCCTgCCTCTATATCT | TT | TgTgAATCTTAggCAggACTgCTgC |
| 5 | gTCCCTgCCTCTATATCT | TT | gCTgACAgTgCAgTTCCTgAATCCT |
| 7 | gTCCCTgCCTCTATATCT | TT | TgCTggAggAgCAAggACCTggTCT |
| 9 | gTCCCTgCCTCTATATCT | TT | CAAgTgCATggTAgCTTTCTTggTg |
| 11 | gTCCCTgCCTCTATATCT | TT | AgCTgTgAAAATCAgCAAggAAgCA |
| 13 | gTCCCTgCCTCTATATCT | TT | gggATTAAACAgATgggACAggggg |
| 15 | gTCCCTgCCTCTATATCT | TT | AgCAgTgATgTACACCCCATCggCC |
| 17 | gTCCCTgCCTCTATATCT | TT | TgAgTgAAAgTAggAgggAggTgCT |
| 19 | gTCCCTgCCTCTATATCT | TT | CTCTCTgATCAgTTgTCAggAgTCA |
| 21 | gTCCCTgCCTCTATATCT | TT | gAACCCACTgggCTCATCTCCACCT |
| 23 | gTCCCTgCCTCTATATCT | TT | TTCCgCCTCCgTggCTggTACTTgT |
| 25 | gTCCCTgCCTCTATATCT | TT | AgCCCCgCCAgCCTCCCCCTCCACC |
| 27 | gTCCCTgCCTCTATATCT | TT | CCCAggATgCCTgTggTCCAggTgg |
| 29 | gTCCCTgCCTCTATATCT | TT | gCCgTggCTCTgACCTgAAgAgTgC |
| 31 | gTCCCTgCCTCTATATCT | TT | gAgTCAgCTTTgCCTgCCTgCAgCT |
| 33 | gTCCCTgCCTCTATATCT | TT | ggTggCAggTATTggTCAAATTCgT |
| 35 | gTCCCTgCCTCTATATCT | TT | AgTgTCCACTggCCgCAgCCAgggC |
| 37 | gTCCCTgCCTCTATATCT | TT | CCCAgTgCCCCCCTgTTCTCCCTCC |
| 39 | gTCCCTgCCTCTATATCT | TT | TggCTgACCgCTTCACggATgCAgA |

| Even # | Probe Sequence (25 nt) | Spacer | 2nd Half of Initiator I1 |
|---|---|---|---|
| 2 | CCATggACCCgTCACTCCATgTCTT | TT | CCACTCAACTTTAACCCg |
| 4 | TCCAgCAgggATCAAgATTCATgCA | TT | CCACTCAACTTTAACCCg |
| 6 | ACgggCTATgAAATgAgAAAggCTA | TT | CCACTCAACTTTAACCCg |
| 8 | TTCACgTTTTCAgCAgACACAgTCA | TT | CCACTCAACTTTAACCCg |
| 10 | AATATTggAACCACATCTgggTgTT | TT | CCACTCAACTTTAACCCg |
| 12 | gAggCggggAgAAAAgCTATAgCgT | TT | CCACTCAACTTTAACCCg |
| 14 | TTATAAAATCCATgCAggAAggggT | TT | CCACTCAACTTTAACCCg |
| 16 | AgATggCgATAATgTgATgAACAAA | TT | CCACTCAACTTTAACCCg |
| 18 | TATATCCACgAgAgTATCTTTCCAT | TT | CCACTCAACTTTAACCCg |
| 20 | TACCCgTTAgAAggTCCCACAACAC | TT | CCACTCAACTTTAACCCg |
| 22 | CCCggCgAgAggCAgTggTggTCTT | TT | CCACTCAACTTTAACCCg |
| 24 | CCCTCgCCCTgCgTggCCTTgCCAT | TT | CCACTCAACTTTAACCCg |
| 26 | ATTCTTgTAgTgAgCCTggATAgAg | TT | CCACTCAACTTTAACCCg |
| 28 | gTgCCCATCggACATgggTgACCCT | TT | CCACTCAACTTTAACCCg |
| 30 | CTTAggggTggTgggAggAgTgggA | TT | CCACTCAACTTTAACCCg |
| 32 | TCCCCAAgggAACgCCCTTCTCgCT | TT | CCACTCAACTTTAACCCg |
| 34 | TggCCTgggTggCCggCgTgTCCgT | TT | CCACTCAACTTTAACCCg |
| 36 | CTCCATgCTgCTTggAgATCCAggC | TT | CCACTCAACTTTAACCCg |
| 38 | AAACCCTgTgAAggCTgCAgCTCCT | TT | CCACTCAACTTTAACCCg |
| 40 | AgggTCCAgTCATAgCCgCTCAgCA | TT | CCACTCAACTTTAACCCg |

#### S4.3 Standard probes for Figures S9 and S10

Organism: ***G. gallus domesticus***
Target mRNA: **SRY (sex determining region Y)-box 10 (*Sox10*)**
Probe set: **20 probe pairs (odd probes carry full HCR initiator, even probes carry no initiator)**
HCR amplifier: **B3-Alexa647**

| Odd # | Initiator I1 | Spacer | Probe Sequence (25 nt) |
|---|---|---|---|
| 1 | gTCCCTgCCTCTATATCTCCACTCAACTTTAACCCg | TACAA | gTCTTCCTCATCTAgAAggCCAATA |
| 3 | gTCCCTgCCTCTATATCTCCACTCAACTTTAACCCg | TACAA | TgTgAATCTTAggCAggACTgCTgC |
| 5 | gTCCCTgCCTCTATATCTCCACTCAACTTTAACCCg | TACAA | gCTgACAgTgCAgTTCCTgAATCCT |
| 7 | gTCCCTgCCTCTATATCTCCACTCAACTTTAACCCg | TACAA | TgCTggAggAgCAAggACCTggTCT |
| 9 | gTCCCTgCCTCTATATCTCCACTCAACTTTAACCCg | TACAA | CAAgTgCATggTAgCTTTCTTggTg |
| 11 | gTCCCTgCCTCTATATCTCCACTCAACTTTAACCCg | TACAA | AgCTgTgAAAATCAgCAAggAAgCA |
| 13 | gTCCCTgCCTCTATATCTCCACTCAACTTTAACCCg | TACAA | gggATTAAACAgATgggACAggggg |
| 15 | gTCCCTgCCTCTATATCTCCACTCAACTTTAACCCg | TACAA | AgCAgTgATgTACACCCCATCggCC |
| 17 | gTCCCTgCCTCTATATCTCCACTCAACTTTAACCCg | TACAA | TgAgTgAAAgTAggAgggAggTgCT |
| 19 | gTCCCTgCCTCTATATCTCCACTCAACTTTAACCCg | TACAA | CTCTCTgATCAgTTgTCAggAgTCA |
| 21 | gTCCCTgCCTCTATATCTCCACTCAACTTTAACCCg | TACAA | gAACCCACTgggCTCATCTCCACCT |
| 23 | gTCCCTgCCTCTATATCTCCACTCAACTTTAACCCg | TACAA | TTCCgCCTCCgTggCTggTACTTgT |
| 25 | gTCCCTgCCTCTATATCTCCACTCAACTTTAACCCg | TACAA | AgCCCCgCCAgCCTCCCCCTCCACC |
| 27 | gTCCCTgCCTCTATATCTCCACTCAACTTTAACCCg | TACAA | CCCAggATgCCTgTggTCCAggTgg |
| 29 | gTCCCTgCCTCTATATCTCCACTCAACTTTAACCCg | TACAA | gCCgTggCTCTgACCTgAAgAgTgC |
| 31 | gTCCCTgCCTCTATATCTCCACTCAACTTTAACCCg | TACAA | gAgTCAgCTTTgCCTgCCTgCAgCT |
| 33 | gTCCCTgCCTCTATATCTCCACTCAACTTTAACCCg | TACAA | ggTggCAggTATTggTCAAATTCgT |
| 35 | gTCCCTgCCTCTATATCTCCACTCAACTTTAACCCg | TACAA | AgTgTCCACTggCCgCAgCCAgggC |
| 37 | gTCCCTgCCTCTATATCTCCACTCAACTTTAACCCg | TACAA | CCCAgTgCCCCCCTgTTCTCCCTCC |
| 39 | gTCCCTgCCTCTATATCTCCACTCAACTTTAACCCg | TACAA | TggCTgACCgCTTCACggATgCAgA |

| Even # | Probe Sequence (25 nt) |
|---|---|
| 2 | CCATggACCCgTCACTCCATgTCTT |
| 4 | TCCAgCAgggATCAAgATTCATgCA |
| 6 | ACgggCTATgAAATgAgAAAggCTA |
| 8 | TTCACgTTTTCAgCAgACACAgTCA |
| 10 | AATATTggAACCACATCTgggTgTT |
| 12 | gAggCggggAgAAAAgCTATAgCgT |
| 14 | TTATAAAATCCATgCAggAAggggT |
| 16 | AgATggCgATAATgTgATgAACAAA |
| 18 | TATATCCACgAgAgTATCTTTCCAT |
| 20 | TACCCgTTAgAAggTCCCACAACAC |
| 22 | CCCggCgAgAggCAgTggTggTCTT |
| 24 | CCCTCgCCCTgCgTggCCTTgCCAT |
| 26 | ATTCTTgTAgTgAgCCTggATAgAg |
| 28 | gTgCCCATCggACATgggTgACCCT |
| 30 | CTTAggggTggTgggAggAgTgggA |
| 32 | TCCCCAAgggAACgCCCTTCTCgCT |
| 34 | TggCCTgggTggCCggCgTgTCCgT |
| 36 | CTCCATgCTgCTTggAgATCCAggC |
| 38 | AAACCCTgTgAAggCTgCAgCTCCT |
| 40 | AgggTCCAgTCATAgCCgCTCAgCA |

Organism: ***G. gallus domesticus***

Target mRNA: **SRY (sex determining region Y)-box 10 (*Sox10*)**

Probe set: **20 probe pairs (odd probes carry no initiator, even probes carry full HCR initiator)**

HCR amplifier: **B3-Alexa647**

| Odd # | Probe Sequence (25 nt) | Even # | Probe Sequence (25 nt) | Spacer | Initiator I1 |
|---|---|---|---|---|---|
| 1 | gTCTTCCTCATCTAgAAggCCAATA | 2 | CCATggACCCgTCACTCCATgTCTT | TACTT | gTCCCTgCCTCTATATCTCCACTCAACTTTAACCCg |
| 3 | TgTgAATCTTAggCAggACTgCTgC | 4 | TCCAgCAgggATCAAgATTCATgCA | TACTT | gTCCCTgCCTCTATATCTCCACTCAACTTTAACCCg |
| 5 | gCTgACAgTgCAgTTCCTgAATCCT | 6 | ACgggCTATgAAATgAgAAAggCTA | TACTT | gTCCCTgCCTCTATATCTCCACTCAACTTTAACCCg |
| 7 | TgCTggAggAgCAAggACCTggTCT | 8 | TTCACgTTTTCAgCAgACACAgTCA | TACTT | gTCCCTgCCTCTATATCTCCACTCAACTTTAACCCg |
| 9 | CAAgTgCATggTAgCTTTCTTggTg | 10 | AATATTggAACCACATCTgggTgTT | TACTT | gTCCCTgCCTCTATATCTCCACTCAACTTTAACCCg |
| 11 | AgCTgTgAAAATCAgCAAggAAgCA | 12 | gAggCggggAgAAAAgCTATAgCgT | TACTT | gTCCCTgCCTCTATATCTCCACTCAACTTTAACCCg |
| 13 | gggATTAAACAgATgggACAggggg | 14 | TTATAAAATCCATgCAggAAggggT | TACTT | gTCCCTgCCTCTATATCTCCACTCAACTTTAACCCg |
| 15 | AgCAgTgATgTACACCCCATCggCC | 16 | AgATggCgATAATgTgATgAACAAA | TACTT | gTCCCTgCCTCTATATCTCCACTCAACTTTAACCCg |
| 17 | TgAgTgAAAgTAggAgggAggTgCT | 18 | TATATCCACgAgAgTATCTTTCCAT | TACTT | gTCCCTgCCTCTATATCTCCACTCAACTTTAACCCg |
| 19 | CTCTCTgATCAgTTgTCAggAgTCA | 20 | TACCCgTTAgAAggTCCCACAACAC | TACTT | gTCCCTgCCTCTATATCTCCACTCAACTTTAACCCg |
| 21 | gAACCCACTgggCTCATCTCCACCT | 22 | CCCggCgAgAggCAgTggTggTCTT | TACTT | gTCCCTgCCTCTATATCTCCACTCAACTTTAACCCg |
| 23 | TTCCgCCTCCgTggCTggTACTTgT | 24 | CCCTCgCCCTgCgTggCCTTgCCAT | TACTT | gTCCCTgCCTCTATATCTCCACTCAACTTTAACCCg |
| 25 | AgCCCCgCCAgCCTCCCCCTCCACC | 26 | ATTCTTgTAgTgAgCCTggATAgAg | TACTT | gTCCCTgCCTCTATATCTCCACTCAACTTTAACCCg |
| 27 | CCCAggATgCCTgTggTCCAggTgg | 28 | gTgCCCATCggACATgggTgACCCT | TACTT | gTCCCTgCCTCTATATCTCCACTCAACTTTAACCCg |
| 29 | gCCgTggCTCTgACCTgAAgAgTgC | 30 | CTTAggggTggTgggAggAgTgggA | TACTT | gTCCCTgCCTCTATATCTCCACTCAACTTTAACCCg |
| 31 | gAgTCAgCTTTgCCTgCCTgCAgCT | 32 | TCCCCAAgggAACgCCCTTCTCgCT | TACTT | gTCCCTgCCTCTATATCTCCACTCAACTTTAACCCg |
| 33 | ggTggCAggTATTggTCAAATTCgT | 34 | TggCCTgggTggCCggCgTgTCCgT | TACTT | gTCCCTgCCTCTATATCTCCACTCAACTTTAACCCg |
| 35 | AgTgTCCACTggCCgCAgCCAgggC | 36 | CTCCATgCTgCTTggAgATCCAggC | TACTT | gTCCCTgCCTCTATATCTCCACTCAACTTTAACCCg |
| 37 | CCCAgTgCCCCCCTgTTCTCCCTCC | 38 | AAACCCTgTgAAggCTgCAgCTCCT | TACTT | gTCCCTgCCTCTATATCTCCACTCAACTTTAACCCg |
| 39 | TggCTgACCgCTTCACggATgCAgA | 40 | AgggTCCAgTCATAgCCgCTCAgCA | TACTT | gTCCCTgCCTCTATATCTCCACTCAACTTTAACCCg |

#### S4.4 Split-initiator probes for Figure S11

Organism: ***G. gallus domesticus***
Target mRNA: **EPH receptor A4 (*EphA4*)**
Probe set: **20 split-initiator probe pairs (each probe carries half an HCR initiator)**
HCR amplifier: **B2-Alexa647**

| Odd # | 1st Half of Initiator I1 | Spacer | Probe Sequence (25 nt) |
|---|---|---|---|
| 1 | CCTCgTAAATCCTCATCA | AA | gCgCCgACggggACCCCggCCATgC |
| 5 | CCTCgTAAATCCTCATCA | AA | CCCAgCTCCCCCTgCACCgAgCggg |
| 9 | CCTCgTAAATCCTCATCA | AA | TgACTgggCTCCATCACATTgCAAA |
| 13 | CCTCgTAAATCCTCATCA | AA | TCCTTgTCgTTgTTTgATTCATAgT |
| 17 | CCTCgTAAATCCTCATCA | AA | AgTgggCACTTCTTATAgAAgACAC |
| 21 | CCTCgTAAATCCTCATCA | AA | ACgTCCTTCTCTTCCgAgTTgTTgA |
| 25 | CCTCgTAAATCCTCATCA | AA | ATTTTgCAAgCTTggCATTCACCAT |
| 29 | CCTCgTAAATCCTCATCA | AA | AAgAAgCCCCgATCACAggTgCAgg |
| 33 | CCTCgTAAATCCTCATCA | AA | gCgCTCCACTCCAAgTTCACTgACg |
| 37 | CCTCgTAAATCCTCATCA | AA | AAATgTACACCACTgCCACAggACC |
| 41 | CCTCgTAAATCCTCATCA | AA | TgCTTggACACTCCATTCACTgCCC |
| 45 | CCTCgTAAATCCTCATCA | AA | gCAACgCTgTgCCTCgTTATCTCTT |
| 49 | CCTCgTAAATCCTCATCA | AA | gCTgTCTTCACAATgCgATAgCTgC |
| 53 | CCTCgTAAATCCTCATCA | AA | AACTCAAACggCCCACTgAAgTCTC |
| 57 | CCTCgTAAATCCTCATCA | AA | gCTgCAATgAgAATgACCACgAgAA |
| 61 | CCTCgTAAATCCTCATCA | AA | TATgTAAAAggATCCACATATgTTC |
| 65 | CCTCgTAAATCCTCATCA | AA | CATTTAgTAACAACgCCTTCCAAgT |
| 69 | CCTCgTAAATCCTCATCA | AA | TCCCgATgCACATAgCTCATgTCAg |
| 73 | CCTCgTAAATCCTCATCA | AA | CCAAATTTAggTCTgTCgCTgCgTT |
| 77 | CCTCgTAAATCCTCATCA | AA | ggggAgCTgggATCCAgCAgggCTg |

| Even # | Probe Sequence (25 nt) | Spacer | 2nd Half of Initiator I1 |
|---|---|---|---|
| 2 | CAgACgCCgACgAggAgCgggAggA | AA | ATCATCCAgTAAACCgCC |
| 6 | CCTCCTTCCAgCgggCTCgCgATCC | AA | ATCATCCAgTAAACCgCC |
| 10 | ATCCAATCAgTTCgTAACCAATTAT | AA | ATCATCCAgTAAACCgCC |
| 14 | gCAAACTggCTCTCTCgAATAAAAC | AA | ATCATCCAgTAAACCgCC |
| 18 | ggAAACTgTgCCAggTTTCgAACTg | AA | ATCATCCAgTAAACCgCC |
| 22 | CCATCTgCCCCgCAgTACATTTTTg | AA | ATCATCCAgTAAACCgCC |
| 26 | TCTgTTgAgAgCgCCTTgTAgTATC | AA | ATCATCCAgTAAACCgCC |
| 30 | ATggATgCAgCATCATTTTCTgCTC | AA | ATCATCCAgTAAACCgCC |
| 34 | TCgTCCCgTCCTCCCTTgTTCTgTg | AA | ATCATCCAgTAAACCgCC |
| 38 | gTTTTCAgCCCgTTCTgCTgggggC | AA | ATCATCCAgTAAACCgCC |
| 42 | gACACAgCTTggTCCTggCTggggT | AA | ATCATCCAgTAAACCgCC |
| 46 | ggCCTgTCAggTTCCAgCCAggCCA | AA | ATCATCCAgTAAACCgCC |
| 50 | AAACCTTTgATgTCAgTATTCCTgg | AA | ATCATCCAgTAAACCgCC |
| 54 | ATgggggAAggAACTgTgTTAgTTg | AA | ATCATCCAgTAAACCgCC |
| 58 | TTgCTgCgCCTCCTgCTgATgACAA | AA | ATCATCCAgTAAACCgCC |
| 62 | TCCCTCACAgCTTgATTTggATCCT | AA | ATCATCCAgTAAACCgCC |
| 66 | TACTCAgTTATgATCATTACTggTT | AA | ATCATCCAgTAAACCgCC |
| 70 | TTgACCAgTATgTTTCgAgCAgCTA | AA | ATCATCCAgTAAACCgCC |
| 74 | AgTTTgTCCAgCATgTTgACAATCT | AA | ATCATCCAgTAAACCgCC |
| 78 | CTgACAgAAACAACCgCCgAgAACT | AA | ATCATCCAgTAAACCgCC |

#### S4.5 Split-initiator probes for Figures 4 and S12

Organism: ***G. gallus domesticus***
Target mRNA: **forkhead box D3 (*FoxD3*)**
Probe set: **12 split-initiator probe pairs (each probe carries half an HCR initiator)**
HCR amplifier: **B4-Alexa488**

| Odd # | 1st Half of Initiator I1 | Spacer | Probe Sequence (25 nt) |
|---|---|---|---|
| 1 | CCTCAACCTACCTCCAAC | AA | CgCCgCCCgATAgAgTCATCCCCgC |
| 3 | CCTCAACCTACCTCCAAC | AA | CgTCgATATCCACgTCCTCggCCgC |
| 5 | CCTCAACCTACCTCCAAC | AA | CCCgCAggTgCTCCTgCTggTgCCg |
| 7 | CCTCAACCTACCTCCAAC | AA | AgggCCCggCCAgCCCgTAggCgCC |
| 9 | CCTCAACCTACCTCCAAC | AA | gCAgAgCggCggggTgCgggTAggC |
| 11 | CCTCAACCTACCTCCAAC | AA | gCAgCgggCACgCgggCggCAgCAT |
| 13 | CCTCAACCTACCTCCAAC | AA | ggCTgggCCCgAgCTgCgCgTTgAA |
| 15 | CCTCAACCTACCTCCAAC | AA | gCTCggATTTCACgATggAgCCCgC |
| 17 | CCTCAACCTACCTCCAAC | AA | TggCggCggggCCgCCgATgATgTT |
| 19 | CCTCAACCTACCTCCAAC | AA | CCgACTgCACggTgACgggCggCCg |
| 21 | CCTCAACCTACCTCCAAC | AA | gCgCgATggCCgCggTggTCCTggC |
| 23 | CCTCAACCTACCTCCAAC | AA | CCgTTTCCCAgAgATACgTCCgggg |

| Even # | Probe Sequence (25 nt) | Spacer | 2nd Half of Initiator I1 |
|---|---|---|---|
| 2 | gCgCggTCTggCCggACATATCgCT | AT | TCTCACCATATTCgCTTC |
| 4 | CCggCgCgTCgTCTCCCTCgCCCAC | AT | TCTCACCATATTCgCTTC |
| 6 | AgCCCTgCATCATgAgCgCCgTCTg | AT | TCTCACCATATTCgCTTC |
| 8 | CggggggCAgCCCgTAggggCggCC | AT | TCTCACCATATTCgCTTC |
| 10 | gCCCgACgggCgggATgTAggggTA | AT | TCTCACCATATTCgCTTC |
| 12 | CTTTgCggCTCAgCTCgCCCgACgg | AT | TCTCACCATATTCgCTTC |
| 14 | CCCCCAAACTgCTgAgCTgCAgCTg | AT | TCTCACCATATTCgCTTC |
| 16 | CgATgCTgAACgAggggCggCTgCT | AT | TCTCACCATATTCgCTTC |
| 18 | ggAAAgTCTgCgCgCTgggCgCCgA | AT | TCTCACCATATTCgCTTC |
| 20 | ACgCCAgCggCTggTgCgCCACCAg | AT | TCTCACCATATTCgCTTC |
| 22 | TgATgTTggTAggCACgCTgAggAT | AT | TCTCACCATATTCgCTTC |
| 24 | AAATAAAAACCCCgAAAgCgACCTC | AT | TCTCACCATATTCgCTTC |

Organism: ***G. gallus domesticus***
Target mRNA: **diencephalon/mesencephalon homeobox 1 (*Dmbx1*)**
Probe set: **20 split-initiator probe pairs (each probe carries half an HCR initiator)**
HCR amplifier: **B1-Alexa514**

| Odd # | 1st Half of Initiator I1 | Spacer | Probe Sequence (25 nt) |
|---|---|---|---|
| 1 | gAggAgggCAgCAAACgg | AA | CgCTCAgTgAgTTCATggCATgCAg |
| 5 | gAggAgggCAgCAAACgg | AA | AAATgATgTCAgCCAgTCTCTCCgC |
| 9 | gAggAgggCAgCAAACgg | AA | AAATgATgTCAgCCAgTCTCTCCgC |
| 13 | gAggAgggCAgCAAACgg | AA | CAgTgTCAggTATTgTggACTgggT |
| 17 | gAggAgggCAgCAAACgg | AA | CCTCCTCTCTgTCAgTCTggTCCTC |
| 21 | gAggAgggCAgCAAACgg | AA | CgCTgATgggAgACTCTgATTTTgg |
| 25 | gAggAgggCAgCAAACgg | AA | CTgCCATgTgCTggCggAATTgCTC |
| 29 | gAggAgggCAgCAAACgg | AA | gACAgTgCAgggAgCTCAgAggCgC |
| 33 | gAggAgggCAgCAAACgg | AA | TCTCAATACTTgTTgTTTTACTgTT |
| 37 | gAggAgggCAgCAAACgg | AA | gACCTCCggTgCATCTTCTTATggg |
| 41 | gAggAgggCAgCAAACgg | AA | CggTgCTAgTAAgACATTAgTAAAT |
| 45 | gAggAgggCAgCAAACgg | AA | TCACAgCAgTCCAAAgggACAgTTC |
| 49 | gAggAgggCAgCAAACgg | AA | gTTTTCCCAAAgAATgCATCgACAA |
| 53 | gAggAgggCAgCAAACgg | AA | ggggAATAAAAgCAAAAgAggCCAC |
| 57 | gAggAgggCAgCAAACgg | AA | TTTgCTCTAAgCACCATTAAgACTC |
| 61 | gAggAgggCAgCAAACgg | AA | ggAAgTgCTTAAACAggAAATTCAC |
| 65 | gAggAgggCAgCAAACgg | AA | AATTTggCTTTCATTTTTCTCCCCA |
| 69 | gAggAgggCAgCAAACgg | AA | CTAgACCAAAATgCTCTCCAAAAAg |
| 73 | gAggAgggCAgCAAACgg | AA | TAAgAACAgCTTgCATTAATCgTgg |
| 77 | gAggAgggCAgCAAACgg | AA | CTTggCCTCCAgCATTgCAgCATTT |

| Even # | Probe Sequence (25 nt) | Spacer | 2nd Half of Initiator I1 |
|---|---|---|---|
| 2 | CTgCCTgCTggTgCAggTTgTACAT | TA | gAAgAgTCTTCCTTTACg |
| 6 | ggTgCTgggATCCATAACgTgCCTC | TA | gAAgAgTCTTCCTTTACg |
| 10 | gCTTCTgCAgCTgCTCCTTCTggAg | TA | gAAgAgTCTTCCTTTACg |
| 14 | CCTCACTTgggATgCTCTgAgTCTT | TA | gAAgAgTCTTCCTTTACg |
| 18 | TAgCCTCATCCAAggTgCTCTTAAA | TA | gAAgAgTCTTCCTTTACg |
| 22 | CACTgCTAgATgAAggAgTCACggT | TA | gAAgAgTCTTCCTTTACg |
| 26 | AggAgTAATggACCAAgTTgTTggT | TA | gAAgAgTCTTCCTTTACg |
| 30 | CgTgggAgAgAgATTggTAgTACgA | TA | gAAgAgTCTTCCTTTACg |
| 34 | CATgCTgCTTTgCCCggAgCCTTAg | TA | gAAgAgTCTTCCTTTACg |
| 38 | ggTTCCAggAgTgACATgTCTggTg | TA | gAAgAgTCTTCCTTTACg |
| 42 | AgCCAgTAgCAgTgTCTgATgCAAT | TA | gAAgAgTCTTCCTTTACg |
| 46 | gCTCTTTCTgAATgTTACAggCTTA | TA | gAAgAgTCTTCCTTTACg |
| 50 | TATgTACAAgACAAAgCAggACTCT | TA | gAAgAgTCTTCCTTTACg |
| 54 | gACTAgCTACCAAAACTgAgAgAgA | TA | gAAgAgTCTTCCTTTACg |
| 58 | gAgCAgTgAATTgCATAATggTTTT | TA | gAAgAgTCTTCCTTTACg |
| 62 | AgTAAAggAAAAAACACTTgCCTTT | TA | gAAgAgTCTTCCTTTACg |
| 66 | AACAATCAAgTCAAAAgTAACCATg | TA | gAAgAgTCTTCCTTTACg |
| 70 | AgTTTTTATTgTTCTCTATTgTCgA | TA | gAAgAgTCTTCCTTTACg |
| 74 | TAgAATTTggTgATCggAgCgTTTT | TA | gAAgAgTCTTCCTTTACg |
| 78 | AATAgAAAgCCCCCgATTATCACCC | TA | gAAgAgTCTTCCTTTACg |

Organism: ***G. gallus domesticus***

Target mRNA: **SRY (sex determining region Y)-box 10 (*Sox10*)**

Probe set: **20 split-initiator probe pairs (each probe carries half an HCR initiator)**

HCR amplifier: **B3-Alexa546**

| Odd # | 1st Half of Initiator I1 | Spacer | Probe Sequence (25 nt) |
|---|---|---|---|
| 1 | gTCCCTgCCTCTATATCT | TT | gTCTTCCTCATCTAgAAggCCAATA |
| 3 | gTCCCTgCCTCTATATCT | TT | TgTgAATCTTAggCAggACTgCTgC |
| 5 | gTCCCTgCCTCTATATCT | TT | gCTgACAgTgCAgTTCCTgAATCCT |
| 7 | gTCCCTgCCTCTATATCT | TT | TgCTggAggAgCAAggACCTggTCT |
| 9 | gTCCCTgCCTCTATATCT | TT | CAAgTgCATggTAgCTTTCTTggTg |
| 11 | gTCCCTgCCTCTATATCT | TT | AgCTgTgAAAATCAgCAAggAAgCA |
| 13 | gTCCCTgCCTCTATATCT | TT | gggATTAAACAgATgggACAggggg |
| 15 | gTCCCTgCCTCTATATCT | TT | AgCAgTgATgTACACCCCATCggCC |
| 17 | gTCCCTgCCTCTATATCT | TT | TgAgTgAAAgTAggAgggAggTgCT |
| 19 | gTCCCTgCCTCTATATCT | TT | CTCTCTgATCAgTTgTCAggAgTCA |
| 21 | gTCCCTgCCTCTATATCT | TT | gAACCCACTgggCTCATCTCCACCT |
| 23 | gTCCCTgCCTCTATATCT | TT | TTCCgCCTCCgTggCTggTACTTgT |
| 25 | gTCCCTgCCTCTATATCT | TT | AgCCCCgCCAgCCTCCCCCTCCACC |
| 27 | gTCCCTgCCTCTATATCT | TT | CCCAggATgCCTgTggTCCAggTgg |
| 29 | gTCCCTgCCTCTATATCT | TT | gCCgTggCTCTgACCTgAAgAgTgC |
| 31 | gTCCCTgCCTCTATATCT | TT | gAgTCAgCTTTgCCTgCCTgCAgCT |
| 33 | gTCCCTgCCTCTATATCT | TT | ggTggCAggTATTggTCAAATTCgT |
| 35 | gTCCCTgCCTCTATATCT | TT | AgTgTCCACTggCCgCAgCCAgggC |
| 37 | gTCCCTgCCTCTATATCT | TT | CCCAgTgCCCCCCTgTTCTCCCTCC |
| 39 | gTCCCTgCCTCTATATCT | TT | TggCTgACCgCTTCACggATgCAgA |

| Even # | Probe Sequence (25 nt) | Spacer | 2nd Half of Initiator I1 |
|---|---|---|---|
| 2 | CCATggACCCgTCACTCCATgTCTT | TT | CCACTCAACTTTAACCCg |
| 4 | TCCAgCAgggATCAAgATTCATgCA | TT | CCACTCAACTTTAACCCg |
| 6 | ACgggCTATgAAATgAgAAAggCTA | TT | CCACTCAACTTTAACCCg |
| 8 | TTCACgTTTTCAgCAgACACAgTCA | TT | CCACTCAACTTTAACCCg |
| 10 | AATATTggAACCACATCTgggTgTT | TT | CCACTCAACTTTAACCCg |
| 12 | gAggCggggAgAAAAgCTATAgCgT | TT | CCACTCAACTTTAACCCg |
| 14 | TTATAAAATCCATgCAggAAggggT | TT | CCACTCAACTTTAACCCg |
| 16 | AgATggCgATAATgTgATgAACAAA | TT | CCACTCAACTTTAACCCg |
| 18 | TATATCCACgAgAgTATCTTTCCAT | TT | CCACTCAACTTTAACCCg |
| 20 | TACCCgTTAgAAggTCCCACAACAC | TT | CCACTCAACTTTAACCCg |
| 22 | CCCggCgAgAggCAgTggTggTCTT | TT | CCACTCAACTTTAACCCg |
| 24 | CCCTCgCCCTgCgTggCCTTgCCAT | TT | CCACTCAACTTTAACCCg |
| 26 | ATTCTTgTAgTgAgCCTggATAgAg | TT | CCACTCAACTTTAACCCg |
| 28 | gTgCCCATCggACATgggTgACCCT | TT | CCACTCAACTTTAACCCg |
| 30 | CTTAggggTggTgggAggAgTgggA | TT | CCACTCAACTTTAACCCg |
| 32 | TCCCCAAgggAACgCCCTTCTCgCT | TT | CCACTCAACTTTAACCCg |
| 34 | TggCCTgggTggCCggCgTgTCCgT | TT | CCACTCAACTTTAACCCg |
| 36 | CTCCATgCTgCTTggAgATCCAggC | TT | CCACTCAACTTTAACCCg |
| 38 | AAACCCTgTgAAggCTgCAgCTCCT | TT | CCACTCAACTTTAACCCg |
| 40 | AgggTCCAgTCATAgCCgCTCAgCA | TT | CCACTCAACTTTAACCCg |

Organism: ***G. gallus domesticus***
Target mRNA: **EPH receptor A4 (*EphA4*)**
Probe set: **20 split-initiator probe pairs (each probe carries half an HCR initiator)**
HCR amplifier: **B2-Alexa647**

| Odd # | 1st Half of Initiator I1 | Spacer | Probe Sequence (25 nt) |
|---|---|---|---|
| 1 | CCTCgTAAATCCTCATCA | AA | gCgCCgACggggACCCCggCCATgC |
| 5 | CCTCgTAAATCCTCATCA | AA | CCCAgCTCCCCCTgCACCgAgCggg |
| 9 | CCTCgTAAATCCTCATCA | AA | TgACTgggCTCCATCACATTgCAAA |
| 13 | CCTCgTAAATCCTCATCA | AA | TCCTTgTCgTTgTTTgATTCATAgT |
| 17 | CCTCgTAAATCCTCATCA | AA | AgTgggCACTTCTTATAgAAgACAC |
| 21 | CCTCgTAAATCCTCATCA | AA | ACgTCCTTCTCTTCCgAgTTgTTgA |
| 25 | CCTCgTAAATCCTCATCA | AA | ATTTTgCAAgCTTggCATTCACCAT |
| 29 | CCTCgTAAATCCTCATCA | AA | AAgAAgCCCCgATCACAggTgCAgg |
| 33 | CCTCgTAAATCCTCATCA | AA | gCgCTCCACTCCAAgTTCACTgACg |
| 37 | CCTCgTAAATCCTCATCA | AA | AAATgTACACCACTgCCACAggACC |
| 41 | CCTCgTAAATCCTCATCA | AA | TgCTTggACACTCCATTCACTgCCC |
| 45 | CCTCgTAAATCCTCATCA | AA | gCAACgCTgTgCCTCgTTATCTCTT |
| 49 | CCTCgTAAATCCTCATCA | AA | gCTgTCTTCACAATgCgATAgCTgC |
| 53 | CCTCgTAAATCCTCATCA | AA | AACTCAAACggCCCACTgAAgTCTC |
| 57 | CCTCgTAAATCCTCATCA | AA | gCTgCAATgAgAATgACCACgAgAA |
| 61 | CCTCgTAAATCCTCATCA | AA | TATgTAAAAggATCCACATATgTTC |
| 65 | CCTCgTAAATCCTCATCA | AA | CATTTAgTAACAACgCCTTCCAAgT |
| 69 | CCTCgTAAATCCTCATCA | AA | TCCCgATgCACATAgCTCATgTCAg |
| 73 | CCTCgTAAATCCTCATCA | AA | CCAAATTTAggTCTgTCgCTgCgTT |
| 77 | CCTCgTAAATCCTCATCA | AA | ggggAgCTgggATCCAgCAgggCTg |

| Even # | Probe Sequence (25 nt) | Spacer | 2nd Half of Initiator I1 |
|---|---|---|---|
| 2 | CAgACgCCgACgAggAgCgggAggA | AA | ATCATCCAgTAAACCgCC |
| 6 | CCTCCTTCCAgCgggCTCgCgATCC | AA | ATCATCCAgTAAACCgCC |
| 10 | ATCCAATCAgTTCgTAACCAATTAT | AA | ATCATCCAgTAAACCgCC |
| 14 | gCAAACTggCTCTCTCgAATAAAAC | AA | ATCATCCAgTAAACCgCC |
| 18 | ggAAACTgTgCCAggTTTCgAACTg | AA | ATCATCCAgTAAACCgCC |
| 22 | CCATCTgCCCCgCAgTACATTTTTg | AA | ATCATCCAgTAAACCgCC |
| 26 | TCTgTTgAgAgCgCCTTgTAgTATC | AA | ATCATCCAgTAAACCgCC |
| 30 | ATggATgCAgCATCATTTTCTgCTC | AA | ATCATCCAgTAAACCgCC |
| 34 | TCgTCCCgTCCTCCCTTgTTCTgTg | AA | ATCATCCAgTAAACCgCC |
| 38 | gTTTTCAgCCCgTTCTgCTgggggC | AA | ATCATCCAgTAAACCgCC |
| 42 | gACACAgCTTggTCCTggCTggggT | AA | ATCATCCAgTAAACCgCC |
| 46 | ggCCTgTCAggTTCCAgCCAggCCA | AA | ATCATCCAgTAAACCgCC |
| 50 | AAACCTTTgATgTCAgTATTCCTgg | AA | ATCATCCAgTAAACCgCC |
| 54 | ATgggggAAggAACTgTgTTAgTTg | AA | ATCATCCAgTAAACCgCC |
| 58 | TTgCTgCgCCTCCTgCTgATgACAA | AA | ATCATCCAgTAAACCgCC |
| 62 | TCCCTCACAgCTTgATTTggATCCT | AA | ATCATCCAgTAAACCgCC |
| 66 | TACTCAgTTATgATCATTACTggTT | AA | ATCATCCAgTAAACCgCC |
| 70 | TTgACCAgTATgTTTCgAgCAgCTA | AA | ATCATCCAgTAAACCgCC |
| 74 | AgTTTgTCCAgCATgTTgACAATCT | AA | ATCATCCAgTAAACCgCC |
| 78 | CTgACAgAAACAACCgCCgAgAACT | AA | ATCATCCAgTAAACCgCC |

#### S4.6 Split-initiator probes for Figures S13–S17

Organism: ***G. gallus domesticus***
Target mRNA: **EPH receptor A4 (*EphA4*)**
Probe set: **20 split-initiator probe pairs (each probe carries half an HCR initiator)**
HCR amplifier: **B1-Alexa546**

| Odd # | 1st Half of Initiator I1 | Spacer | Probe Sequence (25 nt) |
|---|---|---|---|
| 3 | gAggAgggCAgCAAACgg | AA | TACACgCgggAgCCggTgACggCCC |
| 7 | gAggAgggCAgCAAACgg | AA | TCATCCATTATgCTCACTTCCTCCC |
| 11 | gAggAgggCAgCAAACgg | AA | TATACCCTCTgAgCCCCCTCgCggg |
| 15 | gAggAgggCAgCAAACgg | AA | ACCTCTgTATTCAgCTTCATgATCC |
| 19 | gAggAgggCAgCAAACgg | AA | gATgTATCAgCCCCAgTAATggTgT |
| 23 | gAggAgggCAgCAAACgg | AA | CAgTTgCCAATgggTACCAgCCATT |
| 27 | gAggAgggCAgCAAACgg | AA | TgAggCgggCATTTggCACATgCAA |
| 31 | gAggAgggCAgCAAACgg | AA | ggTgCggATggAgggCgAgTgCAgg |
| 35 | gAggAgggCAgCAAACgg | AA | CgCTTgCACACCACgTTgTAggAgA |
| 39 | gAggAgggCAgCAAACgg | AA | AggAggTCAgTgATggAAACCTTCg |
| 43 | gAggAgggCAgCAAACgg | AA | gCTgCTTggTTAgTTgTCACAgTgA |
| 47 | gAggAgggCAgCAAACgg | AA | ACTTCATACTCCAggATgACTCCAT |
| 51 | gAggAgggCAgCAAACgg | AA | TgAAATACATATgAAgTCAgggggT |
| 55 | gAggAgggCAgCAAACgg | AA | ACTgTgggATTggTACCATCgCCAA |
| 59 | gAggAgggCAgCAAACgg | AA | TCTgCCTCTTgCTTAgCTTTACTgT |
| 63 | gAggAgggCAgCAAACgg | AA | TTCAgAgTCTTgATAgCCACACAgA |
| 67 | gAggAgggCAgCAAACgg | AA | CgAAgCATCCCCACCAACTggATTA |
| 71 | gAggAgggCAgCAAACgg | AA | TggAgAgCAATggggCAgTCCATTg |
| 75 | gAggAgggCAgCAAACgg | AA | CTCTTCAggCTgTTAgggTTgCggA |
| 79 | gAggAgggCAgCAAACgg | AA | TCCATTTTAATggCTTggAgCCAgT |

| Even # | Probe Sequence (25 nt) | Spacer | 2nd Half of Initiator I1 |
|---|---|---|---|
| 4 | TCCAgCAgggTCACTTCgTTggCgg | TA | gAAgAgTCTTCCTTTACg |
| 8 | TggTAggTgCggATCggAgTgTTCT | TA | gAAgAgTCTTCCTTTACg |
| 12 | TCTCTCAgCgTgAACTTgATTTCAA | TA | gAAgAgTCTTCCTTTACg |
| 16 | TTCTTgCTgAgAggCCCCACgTCCC | TA | gAAgAgTCTTCCTTTACg |
| 20 | CAggAgCCACgAACCTCCACCAgAg | TA | gAAgAgTCTTCCTTTACg |
| 24 | CgTTCTTCATAgCCAgCATTgCACA | TA | gAAgAgTCTTCCTTTACg |
| 28 | gTAgAgCCTTCCCAgATggAgTAgC | TA | gAAgAgTCTTCCTTTACg |
| 32 | TCgTTgACgTTggAAATCAggTTCT | TA | gAAgAgTCTTCCTTTACg |
| 36 | CAgTggCTgggCTCCCCTgCCCCgC | TA | gAAgAgTCTTCCTTTACg |
| 40 | ACCTCAAAggTgTAgTTggTgTgTg | TA | gAAgAgTCTTCCTTTACg |
| 44 | gCCTggATCAATgCAATTggggATg | TA | gAAgAgTCTTCCTTTACg |
| 48 | TCgTTTTggTCCTTTTCATAgTACT | TA | gAAgAgTCTTCCTTTACg |
| 52 | TATCCTgCTgCTgTCCTggCCCgCA | TA | gAAgAgTCTTCCTTTACg |
| 56 | ACACTgCCAgCCACTgAAACAAgCA | TA | gAAgAgTCTTCCTTTACg |
| 60 | ACACCTTggTTCAAATgTTTCTCCT | TA | gAAgAgTCTTCCTTTACg |
| 64 | CTCCgTTgTTTgTCAgTgTAACCAg | TA | gAAgAgTCTTCCTTTACg |
| 68 | AgATACTTCATTCCTgAgCCgATgC | TA | gAAgAgTCTTCCTTTACg |
| 72 | TTCTgCCAgCAgTCTAACATCAgCT | TA | gAAgAgTCTTCCTTTACg |
| 76 | CTgggTCTggAgCTCTCgCTgCCTg | TA | gAAgAgTCTTCCTTTACg |
| 80 | gCAgCTgTgAAgTTATCCTTgTATC | TA | gAAgAgTCTTCCTTTACg |

Organism: ***G. gallus domesticus***
Target mRNA: **early growth response 2 (*Egr2*)**
Probe set: **20 split-initiator probe pairs (each probe carries half an HCR initiator)**
HCR amplifier: **B3-Alexa647**

| Odd # | 1st Half of Initiator I1 | Spacer | Probe Sequence (25 nt) | Even # | Probe Sequence (25 nt) | Spacer | 2nd Half of Initiator I1 |
|---|---|---|---|---|---|---|---|
| 1 | gTCCCTgCCTCTATATCT | TT | gCAgTgCTgggCTgCTTgCAgCTgT | 2 | CgTgCCAgggCTgTgCCAgggCTCA | TT | CCACTCAACTTTAACCCg |
| 3 | gTCCCTgCCTCTATATCT | TT | gTCTggCggTAACTATTTATggggg | 4 | CCCgCCgCAgCTCgCgCTggAggAC | TT | CCACTCAACTTTAACCCg |
| 5 | gTCCCTgCCTCTATATCT | TT | CACgCCgCTCATCTggTCgAACggC | 6 | gTCCACgCCgAgCATgCCgTCTgCg | TT | CCACTCAACTTTAACCCg |
| 7 | gTCCCTgCCTCTATATCT | TT | AACCAACCCCAACCAgTgCgTACAA | 8 | CTCTTCggTCACCgTAAAgACAAAA | TT | CCACTCAACTTTAACCCg |
| 9 | gTCCCTgCCTCTATATCT | TT | CCACCCCCCCCCCCCgCAgACgCAA | 10 | gTgCgCTCgTCCAgCCCgggCCCTT | TT | CCACTCAACTTTAACCCg |
| 11 | gTCCCTgCCTCTATATCT | TT | ggTgTCAgCgTggAATAATTAAgAg | 12 | CCCCCggCAggCACCTACggAAATA | TT | CCACTCAACTTTAACCCg |
| 13 | gTCCCTgCCTCTATATCT | TT | TgCACTTgAgTAgCTgAgAgCCTgA | 14 | ggAggCCgCgAgCAgAgCCTTggCT | TT | CCACTCAACTTTAACCCg |
| 15 | gTCCCTgCCTCTATATCT | TT | TTTTgCCTggCAgCCCAAATggTgC | 16 | TgTgACAAgTgTggTAACgCgACTT | TT | CCACTCAACTTTAACCCg |
| 17 | gTCCCTgCCTCTATATCT | TT | CgACCAgTCATTACTTTCCTCCgCA | 18 | CAgAATAAATACgggATATCTCACC | TT | CCACTCAACTTTAACCCg |
| 19 | gTCCCTgCCTCTATATCT | TT | ATATAAggACTgAggAACggggCCC | 20 | CCCTgAATgCCCgggACgTCACTgC | TT | CCACTCAACTTTAACCCg |
| 21 | gTCCCTgCCTCTATATCT | TT | CgCgCCCTTgCgCTCCTTCTggCgC | 22 | TCCgggCTgCggggCggCggCggTC | TT | CCACTCAACTTTAACCCg |
| 23 | gTCCCTgCCTCTATATCT | TT | ACCCCCCACATgCAgCCgggTACgg | 24 | gCTgTCgTCggAgCggATCggTAAg | TT | CCACTCAACTTTAACCCg |
| 25 | gTCCCTgCCTCTATATCT | TT | ggCAAACTTTCCggTCTCggCgTCg | 26 | ACAgCCATACTACAACCAgggAggg | TT | CCACTCAACTTTAACCCg |
| 27 | gTCCCTgCCTCTATATCT | TT | CgCTAAgATgAggggAggCgAAAgC | 28 | TgTgggACCAggTggCAAAgCTgCC | TT | CCACTCAACTTTAACCCg |
| 29 | gTCCCTgCCTCTATATCT | TT | CgCTgTCCCTTCgggAgCCTgggAA | 30 | CCATgTgCCACCTCTCCggCAgACg | TT | CCACTCAACTTTAACCCg |
| 31 | gTCCCTgCCTCTATATCT | TT | gAggTTgTgCTCCgCggCCgAgACA | 32 | TgCCgACTgAggACTCCACgACTCT | TT | CCACTCAACTTTAACCCg |
| 33 | gTCCCTgCCTCTATATCT | TT | gTCgCggCCAggCCgATgTgCgggg | 34 | AAAAAAAAAAAAggAgAAAAAAgCA | TT | CCACTCAACTTTAACCCg |
| 35 | gTCCCTgCCTCTATATCT | TT | AgCCATggTTATCCAAggCTgTggC | 36 | TgATgCACgACgCTCCggCTgTgAC | TT | CCACTCAACTTTAACCCg |
| 37 | gTCCCTgCCTCTATATCT | TT | CTCACCACgACggCACCgCATgCAT | 38 | TCACACCACACAAggCACCAAggAC | TT | CCACTCAACTTTAACCCg |
| 39 | gTCCCTgCCTCTATATCT | TT | gAgCgCCgACCCTgCCgTCgAAggg | 40 | TTATTCgTCgTCgCTTATAAACgCC | TT | CCACTCAACTTTAACCCg |

#### S4.7 Split-initiator probes Figures 5, S18, and S19

Organism: ***G. gallus domesticus***
Target mRNA: **diencephalon/mesencephalon homeobox 1 (*Dmbx1*)**
Probe set: **20 split-initiator probe pairs (each probe carries half an HCR initiator)**
HCR amplifier: **B1-Alexa546**

| Odd # | 1st Half of Initiator I1 | Spacer | Probe Sequence (25 nt) | Even # | Probe Sequence (25 nt) | Spacer | 2nd Half of Initiator I1 |
|---|---|---|---|---|---|---|---|
| 1 | gAggAgggCAgCAAACgg | AA | CgCTCAgTgAgTTCATggCATgCAg | 2 | CTgCCTgCTggTgCAggTTgTACAT | TA | gAAgAgTCTTCCTTTACg |
| 5 | gAggAgggCAgCAAACgg | AA | AAATgATgTCAgCCAgTCTCTCCgC | 6 | ggTgCTgggATCCATAACgTgCCTC | TA | gAAgAgTCTTCCTTTACg |
| 9 | gAggAgggCAgCAAACgg | AA | AAATgATgTCAgCCAgTCTCTCCgC | 10 | gCTTCTgCAgCTgCTCCTTCTggAg | TA | gAAgAgTCTTCCTTTACg |
| 13 | gAggAgggCAgCAAACgg | AA | CAgTgTCAggTATTgTggACTgggT | 14 | CCTCACTTgggATgCTCTgAgTCTT | TA | gAAgAgTCTTCCTTTACg |
| 17 | gAggAgggCAgCAAACgg | AA | CCTCCTCTCTgTCAgTCTggTCCTC | 18 | TAgCCTCATCCAAggTgCTCTTAAA | TA | gAAgAgTCTTCCTTTACg |
| 21 | gAggAgggCAgCAAACgg | AA | CgCTgATgggAgACTCTgATTTTgg | 22 | CACTgCTAgATgAAggAgTCACggT | TA | gAAgAgTCTTCCTTTACg |
| 25 | gAggAgggCAgCAAACgg | AA | CTgCCATgTgCTggCggAATTgCTC | 26 | AggAgTAATggACCAAgTTgTTggT | TA | gAAgAgTCTTCCTTTACg |
| 29 | gAggAgggCAgCAAACgg | AA | gACAgTgCAgggAgCTCAgAggCgC | 30 | CgTgggAgAgAgATTggTAgTACgA | TA | gAAgAgTCTTCCTTTACg |
| 33 | gAggAgggCAgCAAACgg | AA | TCTCAATACTTgTTgTTTTACTgTT | 34 | CATgCTgCTTTgCCCggAgCCTTAg | TA | gAAgAgTCTTCCTTTACg |
| 37 | gAggAgggCAgCAAACgg | AA | gACCTCCggTgCATCTTCTTATggg | 38 | ggTTCCAggAgTgACATgTCTggTg | TA | gAAgAgTCTTCCTTTACg |
| 41 | gAggAgggCAgCAAACgg | AA | CggTgCTAgTAAgACATTAgTAAAT | 42 | AgCCAgTAgCAgTgTCTgATgCAAT | TA | gAAgAgTCTTCCTTTACg |
| 45 | gAggAgggCAgCAAACgg | AA | TCACAgCAgTCCAAAgggACAgTTC | 46 | gCTCTTTCTgAATgTTACAggCTTA | TA | gAAgAgTCTTCCTTTACg |
| 49 | gAggAgggCAgCAAACgg | AA | gTTTTCCCAAAgAATgCATCgACAA | 50 | TATgTACAAgACAAAgCAggACTCT | TA | gAAgAgTCTTCCTTTACg |
| 53 | gAggAgggCAgCAAACgg | AA | ggggAATAAAAgCAAAAgAggCCAC | 54 | gACTAgCTACCAAAACTgAgAgAgA | TA | gAAgAgTCTTCCTTTACg |
| 57 | gAggAgggCAgCAAACgg | AA | TTTgCTCTAAgCACCATTAAgACTC | 58 | gAgCAgTgAATTgCATAATggTTTT | TA | gAAgAgTCTTCCTTTACg |
| 61 | gAggAgggCAgCAAACgg | AA | ggAAgTgCTTAAACAggAAATTCAC | 62 | AgTAAAggAAAAAACACTTgCCTTT | TA | gAAgAgTCTTCCTTTACg |
| 65 | gAggAgggCAgCAAACgg | AA | AATTTggCTTTCATTTTTCTCCCCA | 66 | AACAATCAAgTCAAAAgTAACCATg | TA | gAAgAgTCTTCCTTTACg |
| 69 | gAggAgggCAgCAAACgg | AA | CTAgACCAAAATgCTCTCCAAAAAg | 70 | AgTTTTTATTgTTCTCTATTgTCgA | TA | gAAgAgTCTTCCTTTACg |
| 73 | gAggAgggCAgCAAACgg | AA | TAAgAACAgCTTgCATTAATCgTgg | 74 | TAgAATTTggTgATCggAgCgTTTT | TA | gAAgAgTCTTCCTTTACg |
| 77 | gAggAgggCAgCAAACgg | AA | CTTggCCTCCAgCATTgCAgCATTT | 78 | AATAgAAAgCCCCCgATTATCACCC | TA | gAAgAgTCTTCCTTTACg |

Organism: ***G. gallus domesticus***
Target mRNA: **diencephalon/mesencephalon homeobox 1 (*Dmbx1*)**
Probe set: **20 split-initiator probe pairs (each probe carries half an HCR initiator)**
HCR amplifier: **B2-Alexa647**

| Odd # | 1st Half of Initiator I1 | Spacer | Probe Sequence (25 nt) |
|---|---|---|---|
| 3 | CCTCgTAAATCCTCATCA | AA | AgTCgggCgCgTgCTgTgCTTgCTg |
| 7 | CCTCgTAAATCCTCATCA | AA | AggCCgTgCgACTTCgCCTTTgTTT |
| 11 | CCTCgTAAATCCTCATCA | AA | CACTgTggCTgCTTTCACAgTCCTT |
| 15 | CCTCgTAAATCCTCATCA | AA | ACAgggTTAggTTgAggTCTgTgCT |
| 19 | CCTCgTAAATCCTCATCA | AA | CgTCCACACCAggACTCTTgTCCAC |
| 23 | CCTCgTAAATCCTCATCA | AA | AggAgTAggAgTgAgTTTgAgCCAg |
| 27 | CCTCgTAAATCCTCATCA | AA | TggCAgACggTgTCCCCATTTCgAA |
| 31 | CCTCgTAAATCCTCATCA | AA | ggATgggTgAggACCAgACCTgCTg |
| 35 | CCTCgTAAATCCTCATCA | AA | gTAgggTgTCAAgCCCAAgggATgC |
| 39 | CCTCgTAAATCCTCATCA | AA | ACTCCAggAAgAgATgAgggTggAA |
| 43 | CCTCgTAAATCCTCATCA | AA | TCCTCCTCAAATATTTAAAgAAgAC |
| 47 | CCTCgTAAATCCTCATCA | AA | ACATTATCgCAgggATgAggTgAgg |
| 51 | CCTCgTAAATCCTCATCA | AA | gCggTggATgCTTTCAACATTgTAA |
| 55 | CCTCgTAAATCCTCATCA | AA | gggAgCgTggCTgATTTgTgACTTT |
| 59 | CCTCgTAAATCCTCATCA | AA | TCAgCTTTAgCAgAgAAgAgAgAAg |
| 63 | CCTCgTAAATCCTCATCA | AA | TCTgTCTgTgAACAAgTgCTATTAg |
| 67 | CCTCgTAAATCCTCATCA | AA | TTACACTTCACTgAAgACCAAAgAg |
| 71 | CCTCgTAAATCCTCATCA | AA | gTATgAACACAgTgggAgTTCATAC |
| 75 | CCTCgTAAATCCTCATCA | AA | AgggTCTgAAgCTgCACAgCTTgAg |
| 79 | CCTCgTAAATCCTCATCA | AA | CAAgCCAATCTACTCCTCgCTgCAg |

| Even # | Probe Sequence (25 nt) | Spacer | 2nd Half of Initiator I1 |
|---|---|---|---|
| 4 | gggTgAgggCgTgCACTgAgggCCg | AA | ATCATCCAgTAAACCgCC |
| 8 | CCAgTgCCTCCAgCTgTTgggCAgT | AA | ATCATCCAgTAAACCgCC |
| 12 | CCAgCACAggTggTTCCgTCTTCCC | AA | ATCATCCAgTAAACCgCC |
| 16 | gCgCTgATTCACTggCTgACTgCTC | AA | ATCATCCAgTAAACCgCC |
| 20 | TCgCTCTCTTgCAgTTCAAAgCCTT | AA | ATCATCCAgTAAACCgCC |
| 24 | gCAggCggAAgAggCTCAggggTgA | AA | ATCATCCAgTAAACCgCC |
| 28 | TgTTgACATTCATgCCCAAgTAggg | AA | ATCATCCAgTAAACCgCC |
| 32 | ggCTTggAAgAgAgCTggAggCCTg | AA | ATCATCCAgTAAACCgCC |
| 36 | ACggTTgCCCATCTggCAgTCAgTT | AA | ATCATCCAgTAAACCgCC |
| 40 | AAAgTTTTCCCTgATAgggAgCACC | AA | ATCATCCAgTAAACCgCC |
| 44 | CTgTCTAAACACACATCCTCTCCCT | AA | ATCATCCAgTAAACCgCC |
| 48 | AAAAAgggTgTATATAACACggTTg | AA | ATCATCCAgTAAACCgCC |
| 52 | TTCTgTAACACTgACAgTAACACAC | AA | ATCATCCAgTAAACCgCC |
| 56 | AACCCAAgAAgAgCAACTAgCTgTg | AA | ATCATCCAgTAAACCgCC |
| 60 | TTgCATCATTTCCTggCCgTTATAA | AA | ATCATCCAgTAAACCgCC |
| 64 | CAgCAgCATTTggCCAgCATTTTgT | AA | ATCATCCAgTAAACCgCC |
| 68 | AACCCCATAATTTgTAAATgggggA | AA | ATCATCCAgTAAACCgCC |
| 72 | TTgTCAAggAACCATATAATTCATg | AA | ATCATCCAgTAAACCgCC |
| 76 | CACTTgTTACATTCTCACTTgCTAA | AA | ATCATCCAgTAAACCgCC |
| 80 | ggTTgCTTggggACATggTACTTTT | AA | ATCATCCAgTAAACCgCC |

Organism: ***G. gallus domesticus***
Target mRNA: **EPH receptor A4 (*EphA4*)**
Probe set: **20 split-initiator probe pairs (each probe carries half an HCR initiator)**
HCR amplifier: **B1-Alexa546**

| Odd # | 1st Half of Initiator I1 | Spacer | Probe Sequence (25 nt) |
|---|---|---|---|
| 3 | gAggAgggCAgCAAACgg | AA | TACACgCgggAgCCggTgACggCCC |
| 7 | gAggAgggCAgCAAACgg | AA | TCATCCATTATgCTCACTTCCTCCC |
| 11 | gAggAgggCAgCAAACgg | AA | TATACCCTCTgAgCCCCCTCgCggg |
| 15 | gAggAgggCAgCAAACgg | AA | ACCTCTgTATTCAgCTTCATgATCC |
| 19 | gAggAgggCAgCAAACgg | AA | gATgTATCAgCCCCAgTAATggTgT |
| 23 | gAggAgggCAgCAAACgg | AA | CAgTTgCCAATgggTACCAgCCATT |
| 27 | gAggAgggCAgCAAACgg | AA | TgAggCgggCATTTggCACATgCAA |
| 31 | gAggAgggCAgCAAACgg | AA | ggTgCggATggAgggCgAgTgCAgg |
| 35 | gAggAgggCAgCAAACgg | AA | CgCTTgCACACCACgTTgTAggAgA |
| 39 | gAggAgggCAgCAAACgg | AA | AggAggTCAgTgATggAAACCTTCg |
| 43 | gAggAgggCAgCAAACgg | AA | gCTgCTTggTTAgTTgTCACAgTgA |
| 47 | gAggAgggCAgCAAACgg | AA | ACTTCATACTCCAggATgACTCCAT |
| 51 | gAggAgggCAgCAAACgg | AA | TgAAATACATATgAAgTCAgggggT |
| 55 | gAggAgggCAgCAAACgg | AA | ACTgTgggATTggTACCATCgCCAA |
| 59 | gAggAgggCAgCAAACgg | AA | TCTgCCTCTTgCTTAgCTTTACTgT |
| 63 | gAggAgggCAgCAAACgg | AA | TTCAgAgTCTTgATAgCCACACAgA |
| 67 | gAggAgggCAgCAAACgg | AA | CgAAgCATCCCCACCAACTggATTA |
| 71 | gAggAgggCAgCAAACgg | AA | TggAgAgCAATggggCAgTCCATTg |
| 75 | gAggAgggCAgCAAACgg | AA | CTCTTCAggCTgTTAgggTTgCggA |
| 79 | gAggAgggCAgCAAACgg | AA | TCCATTTTAATggCTTggAgCCAgT |

| Even # | Probe Sequence (25 nt) | Spacer | 2nd Half of Initiator I1 |
|---|---|---|---|
| 4 | TCCAgCAgggTCACTTCgTTggCgg | TA | gAAgAgTCTTCCTTTACg |
| 8 | TggTAggTgCggATCggAgTgTTCT | TA | gAAgAgTCTTCCTTTACg |
| 12 | TCTCTCAgCgTgAACTTgATTTCAA | TA | gAAgAgTCTTCCTTTACg |
| 16 | TTCTTgCTgAgAggCCCCACgTCCC | TA | gAAgAgTCTTCCTTTACg |
| 20 | CAggAgCCACgAACCTCCACCAgAg | TA | gAAgAgTCTTCCTTTACg |
| 24 | CgTTCTTCATAgCCAgCATTgCACA | TA | gAAgAgTCTTCCTTTACg |
| 28 | gTAgAgCCTTCCCAgATggAgTAgC | TA | gAAgAgTCTTCCTTTACg |
| 32 | TCgTTgACgTTggAAATCAggTTCT | TA | gAAgAgTCTTCCTTTACg |
| 36 | CAgTggCTgggCTCCCCTgCCCCgC | TA | gAAgAgTCTTCCTTTACg |
| 40 | ACCTCAAAggTgTAgTTggTgTgTg | TA | gAAgAgTCTTCCTTTACg |
| 44 | gCCTggATCAATgCAATTggggATg | TA | gAAgAgTCTTCCTTTACg |
| 48 | TCgTTTTggTCCTTTTCATAgTACT | TA | gAAgAgTCTTCCTTTACg |
| 52 | TATCCTgCTgCTgTCCTggCCCgCA | TA | gAAgAgTCTTCCTTTACg |
| 56 | ACACTgCCAgCCACTgAAACAAgCA | TA | gAAgAgTCTTCCTTTACg |
| 60 | ACACCTTggTTCAAATgTTTCTCCT | TA | gAAgAgTCTTCCTTTACg |
| 64 | CTCCgTTgTTTgTCAgTgTAACCAg | TA | gAAgAgTCTTCCTTTACg |
| 68 | AgATACTTCATTCCTgAgCCgATgC | TA | gAAgAgTCTTCCTTTACg |
| 72 | TTCTgCCAgCAgTCTAACATCAgCT | TA | gAAgAgTCTTCCTTTACg |
| 76 | CTgggTCTggAgCTCTCgCTgCCTg | TA | gAAgAgTCTTCCTTTACg |
| 80 | gCAgCTgTgAAgTTATCCTTgTATC | TA | gAAgAgTCTTCCTTTACg |

Organism: ***G. gallus domesticus***
Target mRNA: **EPH receptor A4 (*EphA4*)**
Probe set: **20 split-initiator probe pairs (each probe carries half an HCR initiator)**
HCR amplifier: **B2-Alexa647**

| Odd # | 1st Half of Initiator I1 | Spacer | Probe Sequence (25 nt) | Even # | Probe Sequence (25 nt) | Spacer | 2nd Half of Initiator I1 |
|---|---|---|---|---|---|---|---|
| 1 | CCTCgTAAATCCTCATCA | AA | gCgCCgACggggACCCCggCCATgC | 2 | CAgACgCCgACgAggAgCgggAggA | AA | ATCATCCAgTAAACCgCC |
| 5 | CCTCgTAAATCCTCATCA | AA | CCCAgCTCCCCCTgCACCgAgCggg | 6 | CCTCCTTCCAgCgggCTCgCgATCC | AA | ATCATCCAgTAAACCgCC |
| 9 | CCTCgTAAATCCTCATCA | AA | TgACTgggCTCCATCACATTgCAAA | 10 | ATCCAATCAgTTCgTAACCAATTAT | AA | ATCATCCAgTAAACCgCC |
| 13 | CCTCgTAAATCCTCATCA | AA | TCCTTgTCgTTgTTTgATTCATAgT | 14 | gCAAACTggCTCTCTCgAATAAAAC | AA | ATCATCCAgTAAACCgCC |
| 17 | CCTCgTAAATCCTCATCA | AA | AgTgggCACTTCTTATAgAAgACAC | 18 | ggAAACTgTgCCAggTTTCgAACTg | AA | ATCATCCAgTAAACCgCC |
| 21 | CCTCgTAAATCCTCATCA | AA | ACgTCCTTCTCTTCCgAgTTgTTgA | 22 | CCATCTgCCCCgCAgTACATTTTTg | AA | ATCATCCAgTAAACCgCC |
| 25 | CCTCgTAAATCCTCATCA | AA | ATTTTgCAAgCTTggCATTCACCAT | 26 | TCTgTTgAgAgCgCCTTgTAgTATC | AA | ATCATCCAgTAAACCgCC |
| 29 | CCTCgTAAATCCTCATCA | AA | AAgAAgCCCCgATCACAggTgCAgg | 30 | ATggATgCAgCATCATTTTCTgCTC | AA | ATCATCCAgTAAACCgCC |
| 33 | CCTCgTAAATCCTCATCA | AA | gCgCTCCACTCCAAgTTCACTgACg | 34 | TCgTCCCgTCCTCCCTTgTTCTgTg | AA | ATCATCCAgTAAACCgCC |
| 37 | CCTCgTAAATCCTCATCA | AA | AAATgTACACCACTgCCACAggACC | 38 | gTTTTCAgCCCgTTCTgCTgggggC | AA | ATCATCCAgTAAACCgCC |
| 41 | CCTCgTAAATCCTCATCA | AA | TgCTTggACACTCCATTCACTgCCC | 42 | gACACAgCTTggTCCTggCTggggT | AA | ATCATCCAgTAAACCgCC |
| 45 | CCTCgTAAATCCTCATCA | AA | gCAACgCTgTgCCTCgTTATCTCTT | 46 | ggCCTgTCAggTTCCAgCCAggCCA | AA | ATCATCCAgTAAACCgCC |
| 49 | CCTCgTAAATCCTCATCA | AA | gCTgTCTTCACAATgCgATAgCTgC | 50 | AAACCTTTgATgTCAgTATTCCTgg | AA | ATCATCCAgTAAACCgCC |
| 53 | CCTCgTAAATCCTCATCA | AA | AACTCAAACggCCCACTgAAgTCTC | 54 | ATgggggAAggAACTgTgTTAgTTg | AA | ATCATCCAgTAAACCgCC |
| 57 | CCTCgTAAATCCTCATCA | AA | gCTgCAATgAgAATgACCACgAgAA | 58 | TTgCTgCgCCTCCTgCTgATgACAA | AA | ATCATCCAgTAAACCgCC |
| 61 | CCTCgTAAATCCTCATCA | AA | TATgTAAAAggATCCACATATgTTC | 62 | TCCCTCACAgCTTgATTTggATCCT | AA | ATCATCCAgTAAACCgCC |
| 65 | CCTCgTAAATCCTCATCA | AA | CATTTAgTAACAACgCCTTCCAAgT | 66 | TACTCAgTTATgATCATTACTggTT | AA | ATCATCCAgTAAACCgCC |
| 69 | CCTCgTAAATCCTCATCA | AA | TCCCgATgCACATAgCTCATgTCAg | 70 | TTgACCAgTATgTTTCgAgCAgCTA | AA | ATCATCCAgTAAACCgCC |
| 73 | CCTCgTAAATCCTCATCA | AA | CCAAATTTAggTCTgTCgCTgCgTT | 74 | AgTTTgTCCAgCATgTTgACAATCT | AA | ATCATCCAgTAAACCgCC |
| 77 | CCTCgTAAATCCTCATCA | AA | ggggAgCTgggATCCAgCAgggCTg | 78 | CTgACAgAAACAACCgCCgAgAACT | AA | ATCATCCAgTAAACCgCC |

#### S4.8 Split-initiator probes for Figure 6, S20–S28

Organism: ***H. sapiens sapiens***
Target mRNA: **destabilized enhanced green fluorescent protein (*Tg(d2eGFP)*)**
Probe set: **12 split-initiator probe pairs (each probe carries half an HCR initiator)**
HCR amplifier: **B3-Alexa594 (Figures 6A and S20)**

| Odd # | 1st Half of Initiator I1 | Spacer | Probe Sequence (25 nt) |
|---|---|---|---|
| 1 | gTCCCTgCCTCTATATCT | TT | TTgTggCCgTTTACgTCgCCgTCCA |
| 3 | gTCCCTgCCTCTATATCT | TT | AgggTCAgCTTgCCgTAggTggCAT |
| 5 | gTCCCTgCCTCTATATCT | TT | gTCACgAgggTgggCCAgggCACgg |
| 7 | gTCCCTgCCTCTATATCT | TT | TgCTTCATgTggTCggggTAgCggC |
| 9 | gTCCCTgCCTCTATATCT | TT | ATggTgCgCTCCTggACgTAgCCTT |
| 11 | gTCCCTgCCTCTATATCT | TT | CCCTCgAACTTCACCTCggCgCggg |
| 13 | gTCCCTgCCTCTATATCT | TT | CCgTCCTCCTTgAAgTCgATgCCCT |
| 15 | gTCCCTgCCTCTATATCT | TT | ATATAgACgTTgTggCTgTTgTAgT |
| 17 | gTCCCTgCCTCTATATCT | TT | TTgTggCggATCTTgAAgTTCACCT |
| 19 | gTCCCTgCCTCTATATCT | TT | ATggggTgTTCTgCTggTAgTggT |
| 21 | gTCCCTgCCTCTATATCT | TT | gCggACTgggTgCTCAggTAgTggT |
| 23 | gTCCCTgCCTCTATATCT | TT | ACgAACTCCAgCAggACCATgTgAT |

| Even # | Probe Sequence (25 nt) | Spacer | 2nd Half of Initiator I1 |
|---|---|---|---|
| 2 | CCCTCgCCCTCgCCggACACgCTgA | TT | CCACTCAACTTTAACCCg |
| 4 | AgCTTgCCggTggTgCAgATgAACT | TT | CCACTCAACTTTAACCCg |
| 6 | AAgCACTgCACgCCgTAggTCAggg | TT | CCACTCAACTTTAACCCg |
| 8 | ggCATggCggACTTgAAgAAgTCgT | TT | CCACTCAACTTTAACCCg |
| 10 | TTgTAgTTgCCgTCgTCCTTgAAgA | TT | CCACTCAACTTTAACCCg |
| 12 | AgCTCgATgCggTTCACCAgggTgT | TT | CCACTCAACTTTAACCCg |
| 14 | TACTCCAgCTTgTgCCCCAggATgT | TT | CCACTCAACTTTAACCCg |
| 16 | ATgCCgTTCTTCTgCTTgTCggCCA | TT | CCACTCAACTTTAACCCg |
| 18 | gCgAgCTgCACgCTgCCgTCCTCgA | TT | CCACTCAACTTTAACCCg |
| 20 | TCgggCAgCAgCACggggCCgTCgC | TT | CCACTCAACTTTAACCCg |
| 22 | CgCTTCTCgTTggggTCTTTgCTCA | TT | CCACTCAACTTTAACCCg |
| 24 | ATgCCgAgAgTgATCCCggCggCgg | TT | CCACTCAACTTTAACCCg |

Organism: ***H. sapiens sapiens***
Target mRNA: **glyceraldehyde-3-phosphate dehydrogenase (*GAPDH*)**
Probe set: **10 split-initiator probe pairs (each probe carries half an HCR initiator)**
HCR amplifier: **B5-Alexa488 (Figures 6B, S24, and S23)**

| Odd # | 1st Half of Initiator I1 | Spacer | Probe Sequence (25 nt) |
|---|---|---|---|
| 3 | CTCACTCCCAATCTCTAT | AA | gggTCATTgATggCAACAATATCCA |
| 7 | CTCACTCCCAATCTCTAT | AA | TTTCCATTgATgACAAgCTTCCCgT |
| 11 | CTCACTCCCAATCTCTAT | AA | gCCTTCTCCATggTggTgAAgACgC |
| 15 | CTCACTCCCAATCTCTAT | AA | AggCTgTTgTCATACTTCTCATggT |
| 19 | CTCACTCCCAATCTCTAT | AA | gCATggACTgTggTCATgAgTCCTT |
| 23 | CTCACTCCCAATCTCTAT | AA | gCCTTggCAgCgCCAgTAgAggCAg |
| 27 | CTCACTCCCAATCTCTAT | AA | ggTTTTTCTAgACggCAggTCAggT |
| 31 | CTCACTCCCAATCTCTAT | AA | CTgTTgAAgTCAgAggAgACCACCT |
| 35 | CTCACTCCCAATCTCTAT | AA | CTggTggTCCAggggTCTTACTCCT |
| 39 | CTCACTCCCAATCTCTAT | AA | CTACATggCAACTgTgAggAggggA |

| Even # | Probe Sequence (25 nt) | Spacer | 2nd Half of Initiator I1 |
|---|---|---|---|
| 4 | TAAACCATgTAgTTgAggTCAATgA | AA | CTACCCTACAAATCCAAT |
| 8 | TCTCgCTCCTggAAgATggTgATgg | AA | CTACCCTACAAATCCAAT |
| 12 | TTggCTCCCCCCTgCAAATgAgCCC | AA | CTACCCTACAAATCCAAT |
| 16 | gTgCAggAggCATTgCTgATgATCT | AA | CTACCCTACAAATCCAAT |
| 20 | TCCACAgTCTTCTgggTggCAgTgA | AA | CTACCCTACAAATCCAAT |
| 24 | TTCAgCTCAgggATgACCTTgCCCA | AA | CTACCCTACAAATCCAAT |
| 28 | ACCTTCTTgATgTCATCATATTTgg | AA | CTACCCTACAAATCCAAT |
| 32 | gCgTCAAAggTggAggAgTgggTgT | AA | CTACCCTACAAATCCAAT |
| 36 | TCTCTTCCTCTTgTgCTCTTgCTgg | AA | CTACCCTACAAATCCAAT |
| 40 | CCTAggCCCCTCCCCTCTTCAAggg | AA | CTACCCTACAAATCCAAT |

Organism: ***H. sapiens sapiens***
Target mRNA: **glyceraldehyde-3-phosphate dehydrogenase (*GAPDH*)**
Probe set: **10 split-initiator probe pairs (each probe carries half an HCR initiator)**
HCR amplifier: **B4-Alexa594 (Figures 6B and S24), B4-Alexa488 (Figures 6C and S27)**

| Odd # | 1st Half of Initiator I1 | Spacer | Probe Sequence (25 nt) |
|---|---|---|---|
| 1 | CCTCAACCTACCTCCAAC | AA | ACCAggCgCCCAATACgACCAAATC |
| 5 | CCTCAACCTACCTCCAAC | AA | CCATgggTggAATCATATTggAACA |
| 9 | CCTCAACCTACCTCCAAC | AA | gCATCgCCCCACTTgATTTTggAgg |
| 13 | CCTCAACCTACCTCCAAC | AA | gCAgAgggggCAgAgATgATgACCC |
| 17 | CCTCAACCTACCTCCAAC | AA | TTggCCAggggTgCTAAgCAgTTgg |
| 21 | CCTCAACCTACCTCCAAC | AA | TCACgCCACAgTTTCCCggAggggC |
| 25 | CCTCAACCTACCTCCAAC | AA | CggAAggCCATgCCAgTgAgCTTCC |
| 29 | CCTCAACCTACCTCCAAC | AA | AgggggCCCTCCgACgCCTgCTTCA |
| 33 | CCTCAACCTACCTCCAAC | AA | TggTCgTTgAgggCAATgCCAgCCC |
| 37 | CCTCAACCTACCTCCAAC | AA | CAgggACTCCCCAgCAgTgAgggTC |

| Even # | Probe Sequence (25 nt) | Spacer | 2nd Half of Initiator I1 |
|---|---|---|---|
| 2 | TTACCAgAgTTAAAAgCAgCCCTgg | AT | TCTCACCATATTCgCTTC |
| 6 | TCAgCCTTgACggTgCCATggAATT | AT | TCTCACCATATTCgCTTC |
| 10 | gTggACTCCACgACgTACTCAgCgC | AT | TCTCACCATATTCgCTTC |
| 14 | ACACCCATgACgAACATgggggCAT | AT | TCTCACCATATTCgCTTC |
| 18 | ACgATACCAAAgTTgTCATggATgA | AT | TCTCACCATATTCgCTTC |
| 22 | ATgATgTTCTggAgAgCCCCgCggC | AT | TCTCACCATATTCgCTTC |
| 26 | ACCACTgACACgTTggCAgTggggA | AT | TCTCACCATATTCgCTTC |
| 30 | TgCTCAgTgTAgCCCAggATgCCCT | AT | TCTCACCATATTCgCTTC |
| 34 | TCATACCAggAAATgAgCTTgACAA | AT | TCTCACCATATTCgCTTC |
| 38 | TTCAgTgTggTgggggACTgAgTgT | AT | TCTCACCATATTCgCTTC |

Organism: ***H. sapiens sapiens***

Target mRNA: **actin beta (*ACTB*)**

Probe set: **10 split-initiator probe pairs (each probe carries half an HCR initiator)**

HCR amplifier: **B2-Alexa594 (Figure S23)**

| Odd # | 1st Half of Initiator I1 | Spacer | Probe Sequence (25 nt) |
|---|---|---|---|
| 1 | CCTCgTAAATCCTCATCA | AA | gCggggCggACgCggTCTCggCggT |
| 5 | CCTCgTAAATCCTCATCA | AA | gCTggCggCgggTgTggACgggCgg |
| 9 | CCTCgTAAATCCTCATCA | AA | TgCgCAAgTTAggTTTTgTCAAgAA |
| 13 | CCTCgTAAATCCTCATCA | AA | ACCAAAACAAAACAAAAAAAACAAA |
| 17 | CCTCgTAAATCCTCATCA | AA | CACCTTCACCgTTCCAgTTTTTAAA |
| 21 | CCTCgTAAATCCTCATCA | AA | AgTCCTCggCCACATTgTgAACTTT |
| 25 | CCTCgTAAATCCTCATCA | AA | ACAACgCATCTCATATTTggAATgA |
| 29 | CCTCgTAAATCCTCATCA | AA | ATTCTCCTTAgAgAgAAgTggggTg |
| 33 | CCTCgTAAATCCTCATCA | AA | ACACgAAAgCAATgCTATCACCTCC |
| 37 | CCTCgTAAATCCTCATCA | AA | CAAAATAAAAAAgTATTAAggCgAA |

| Even # | Probe Sequence (25 nt) | Spacer | 2nd Half of Initiator I1 |
|---|---|---|---|
| 2 | gATCggCAAAggCgAggCTCTgTgC | AA | ATCATCCAgTAAACCgCC |
| 6 | gCgCggCgATATCATCATCCATggT | AA | ATCATCCAgTAAACCgCC |
| 10 | AAgCCATgCCAATCTCATCTTgTTT | AA | ATCATCCAgTAAACCgCC |
| 14 | CTgAgTCAAgCCAAAAAAAAAAAAA | AA | ATCATCCAgTAAACCgCC |
| 18 | gggATgCTCgCTCCAACCgACTgCT | AA | ATCATCCAgTAAACCgCC |
| 22 | ATTAAAAAAACAACAATgTgCAATC | AA | ATCATCCAgTAAACCgCC |
| 26 | TTTTAggATggCAAgggACTTCCTg | AA | ATCATCCAgTAAACCgCC |
| 30 | TgTgTggACTTgggAgAggACTggg | AA | ATCATCCAgTAAACCgCC |
| 34 | TTAAAAAAATTTTgCATTACATAAT | AA | ATCATCCAgTAAACCgCC |
| 38 | CACgAAggCTCATCATTCAAAATAA | AA | ATCATCCAgTAAACCgCC |

Organism: ***H. sapiens sapiens***
Target mRNA: **phosphoglycerate kinase 1(*PGK1*)**
Probe set: **18 split-initiator probe pairs (each probe carries half an HCR initiator)**
HCR amplifier: **B1-Alexa488 (Figures 6B and S25)**

| Odd # | 1st Half of Initiator I1 | Spacer | Probe Sequence (25 nt) |
|---|---|---|---|
| 1 | gAggAgggCAgCAAACgg | AA | CCgCCCCTTCCCggCCgCTgCTCTC |
| 5 | gAggAgggCAgCAAACgg | AA | CAACgAgggAgCCgACTgCCgACgT |
| 9 | gAggAgggCAgCAAACgg | AA | gACTCTCATAACgACCCgCTTCCCT |
| 13 | gAggAgggCAgCAAACgg | AA | CTTCCCTTCTTCCTCCACATgAAAg |
| 17 | gAggAgggCAgCAAACgg | AA | gCCAAAAgCATCATTgACATAgACA |
| 21 | gAggAgggCAgCAAACgg | AA | gCTCTCCAAggCCTTTgCAAAgTAg |
| 25 | gAggAgggCAgCAAACgg | AA | CTTCTCAgCTTTggACATTAggTCT |
| 29 | gAggAgggCAgCAAACgg | AA | CCAgCCAgCAggTATgCCAgAAgCC |
| 33 | gAggAgggCAgCAAACgg | AA | AgCTTCCCATTCAAATACCCCCACA |
| 37 | gAggAgggCAgCAAACgg | AA | gTACTAAATATTgCTgAgAgCATCC |
| 41 | gAggAgggCAgCAAACgg | AA | ggCCACTAgCTgAATCTTgACATgg |
| 45 | gAggAgggCAgCAAACgg | AA | CTAAAAAATTCAAATgggATCTTgA |
| 49 | gAggAgggCAgCAAACgg | AA | TAATCATAATAACCTACATCAAAAg |
| 53 | gAggAgggCAgCAAACgg | AA | TCAATggACACTTTTATTgTTTACT |
| 57 | gAggAgggCAgCAAACgg | AA | CTgCCCCACTTCTTgCATTCAgCAA |
| 61 | gAggAgggCAgCAAACgg | AA | CTgATAAAAATAAAAgTTAgAATAA |
| 65 | gAggAgggCAgCAAACgg | AA | CAAgAgTTgAAAgTggTCACCTCTg |
| 69 | gAggAgggCAgCAAACgg | AA | gAgCCTTCCTCCATggTATgAAATA |

| Even # | Probe Sequence (25 nt) | Spacer | 2nd Half of Initiator I1 |
|---|---|---|---|
| 2 | CTACCgCCCCACACCCCgCCTCCCg | TA | gAAgAgTCTTCCTTTACg |
| 6 | gCTggggAgAgAggTCggTgATTCg | TA | gAAgAgTCTTCCTTTACg |
| 10 | gTTgTTCTTCATAggAACATTgAAg | TA | gAAgAgTCTTCCTTTACg |
| 14 | AACCTTgTTCCCAgAAgCATCTTTT | TA | gAAgAgTCTTCCTTTACg |
| 18 | CATggAgCTgTgggCTCTgTgAgCA | TA | gAAgAgTCTTCCTTTACg |
| 22 | CAggATggCCAggAAgggTCgCTCT | TA | gAAgAgTCTTCCTTTACg |
| 26 | AACAggCAAggTAATCTTCACACCA | TA | gAAgAgTCTTCCTTTACg |
| 30 | gCTTTCAggACCACAgTCCAAgCCC | TA | gAAgAgTCTTCCTTTACg |
| 34 | CATgAgAgCTTTggTTCCCCgggCA | TA | gAAgAgTCTTCCTTTACg |
| 38 | TgTgCACAggAACTAAAAggCAggA | TA | gAAgAgTCTTCCTTTACg |
| 42 | TTAAgggTTCCTggCACTgCATCTC | TA | gAAgAgTCTTCCTTTACg |
| 46 | ACTCTAgAATgCACAATggTTTAgT | TA | gAAgAgTCTTCCTTTACg |
| 50 | TgCTgAgTAgTgAAACAgTgACAAA | TA | gAAgAgTCTTCCTTTACg |
| 54 | gACAggAAAAAAAAAAAAATCACgg | TA | gAAgAgTCTTCCTTTACg |
| 58 | TCTAATTgTCCCATCTCTCCACTgC | TA | gAAgAgTCTTCCTTTACg |
| 62 | gACTTTTTAAATTATgATCATgTgT | TA | gAAgAgTCTTCCTTTACg |
| 66 | AACATggAggTATATACCTgAAAAA | TA | gAAgAgTCTTCCTTTACg |
| 70 | TgAAgAAgTggAAATATATgTggAA | TA | gAAgAgTCTTCCTTTACg |

Organism: ***H. sapiens sapiens***

Target mRNA: **phosphoglycerate kinase 1 (*PGK1*)**

Probe set: **18 split-initiator probe pairs (each probe carries half an HCR initiator)**

HCR amplifier: **B2-Alexa594 (Figures 6B, 6C, S25, and S27)**

| Odd # | 1st Half of Initiator I1 | Spacer | Probe Sequence (25 nt) |
|---|---|---|---|
| 3 | CCTCgTAAATCCTCATCA | AA | ACCgCgCgggCAggAACAgggCCCA |
| 7 | CCTCgTAAATCCTCATCA | AA | gTTAgAAAgCgACATTTTggAAATA |
| 11 | CCTCgTAAATCCTCATCA | AA | CTTAATCCTCTggTTgTTTgTTATC |
| 15 | CCTCgTAAATCCTCATCA | AA | ggAgTACTTgTCAggCATgggCACA |
| 19 | CCTCgTAAATCCTCATCA | AA | AgCCTTCTgTggCAgATTgACTCCT |
| 23 | CCTCgTAAATCCTCATCA | AA | ATCAAACTTgTCAgCAgTgACAAAg |
| 27 | CCTCgTAAATCCTCATCA | AA | AgTgACAgCCTCAgCATACTTCTTg |
| 31 | CCTCgTAAATCCTCATCA | AA | CCTAgAAgTggCTTTCACCACCTCA |
| 35 | CCTCgTAAATCCTCATCA | AA | CTCCAAACTggCACCACCCCCAgTg |
| 39 | CCTCgTAAATCCTCATCA | AA | CAgAAAATgCTAAgTTgACTTAggg |
| 43 | CCTCgTAAATCCTCATCA | AA | AgATgAgCTgAgATgCTgTgCAACT |
| 47 | CCTCgTAAATCCTCATCA | AA | TTTAACAggCAAAATATAAATATAT |
| 51 | CCTCgTAAATCCTCATCA | AA | TACAAATggAATTTCATCTTgTTTC |
| 55 | CCTCgTAAATCCTCATCA | AA | CTATTCTCACCCTTCCTAACAAAgT |
| 59 | CCTCgTAAATCCTCATCA | AA | AggCCCTTgATAAAgAATggACATT |
| 63 | CCTCgTAAATCCTCATCA | AA | ggCTggggCTTTTTTgTTATAAgCC |
| 67 | CCTCgTAAATCCTCATCA | AA | gTgAACAATATAAgCATATTACTTA |
| 71 | CCTCgTAAATCCTCATCA | AA | AATTgTgACAAAACTATACCgAgAg |

| Even # | Probe Sequence (25 nt) | Spacer | 2nd Half of Initiator I1 |
|---|---|---|---|
| 4 | gCTCCggAggCTTgCAgAATgCggA | AA | ATCATCCAgTAAACCgCC |
| 8 | AACgTCCAgCTTgTCCAgCgTCAgC | AA | ATCATCCAgTAAACCgCC |
| 12 | gCAgAATTTgATgCTTgggACAgCA | AA | ATCATCCAgTAAACCgCC |
| 16 | TTTgAgTTCTACAgCAACTggCTCT | AA | ATCATCCAgTAAACCgCC |
| 20 | CAgCTCCTTCTTCATCAAAAACCCA | AA | ATCATCCAgTAAACCgCC |
| 24 | AgTggCTTggCCAgTCTTggCATTC | AA | ATCATCCAgTAAACCgCC |
| 28 | ACCATTCCACACAATCTgCTTAgCC | AA | ATCATCCAgTAAACCgCC |
| 32 | TCCACCACCTATgATggTgATgCAg | AA | ATCATCCAgTAAACCgCC |
| 36 | CCCAggAAggACTTTACCTTCCAgg | AA | ATCATCCAgTAAACCgCC |
| 40 | ggTTTTAgCTAATgCCAAgTggAgA | AA | ATCATCCAgTAAACCgCC |
| 44 | AATgTATgCAAATCCAgggTgCAgT | AA | ATCATCCAgTAAACCgCC |
| 48 | AACTAAgCTAACACTgCTCACTTTC | AA | ATCATCCAgTAAACCgCC |
| 52 | ATggATCATCAATTTTgTCTCACTA | AA | ATCATCCAgTAAACCgCC |
| 56 | TAgACATCTgATCCgTTCCTCAAgA | AA | ATCATCCAgTAAACCgCC |
| 60 | gCACTAgCACAATgTCTgCCATAAA | AA | ATCATCCAgTAAACCgCC |
| 64 | gAgTgggAATCTTgAATgggAggAA | AA | ATCATCCAgTAAACCgCC |
| 68 | TTTCTTTAAAAAATAAAAAAAAAAg | AA | ATCATCCAgTAAACCgCC |
| 72 | gTTATgTAgACTTTTgATCTAATCT | AA | ATCATCCAgTAAACCgCC |

Organism: ***E. coli***
Target mRNA: **enhanced green fluorescent protein (*Tg(EGFP)*)**
Probe set: **12 split-initiator probe pairs (each probe carries half an HCR initiator)**
HCR amplifier: **B3-Alexa594 (Figures 6A and S22)**

| Odd # | 1st Half of Initiator I1 | Spacer | Probe Sequence (25 nt) |
|---|---|---|---|
| 1 | gTCCCTgCCTCTATATCT | TT | TgAAAAgTTCTTCTCCTTTACgCAT |
| 3 | gTCCCTgCCTCTATATCT | TT | ATTTgTgCCCATTAACATCACCATC |
| 5 | gTCCCTgCCTCTATATCT | TT | TAAgggTAAgTTTTCCgTATgTTgC |
| 7 | gTCCCTgCCTCTATATCT | TT | TAgTgACAAgTgTTggCCATggAAC |
| 9 | gTCCCTgCCTCTATATCT | TT | gCTgTTTCATATgATCTgggTATCT |
| 11 | gTCCCTgCCTCTATATCT | TT | ATATAgTTCTTTCCTgTACATAACC |
| 13 | gTCCCTgCCTCTATATCT | TT | CACCTTCAAACTTgACTTCAgCACg |
| 15 | gTCCCTgCCTCTATATCT | TT | TTCCATCTTCTTTAAAATCAATACC |
| 17 | gTCCCTgCCTCTATATCT | TT | TgATgTATACATTgTgTgAgTTATA |
| 19 | gTCCCTgCCTCTATATCT | TT | TgTTgTgTCTAATTTTgAAgTTAAC |
| 21 | gTCCCTgCCTCTATATCT | TT | CAATTggAgTATTTTgTTgATAATg |
| 23 | gTCCCTgCCTCTATATCT | TT | gggCAgATTgTgTggACAggTAATg |

| Even # | Probe Sequence (25 nt) | Spacer | 2nd Half of Initiator I1 |
|---|---|---|---|
| 2 | ATTCAACAAgAATTgggACAACTCC | TT | CCACTCAACTTTAACCCg |
| 4 | CACCTTCACCCTCTCCACTgACAgA | TT | CCACTCAACTTTAACCCg |
| 6 | gTAgTTTTCCAgTAgTgCAAATAAA | TT | CCACTCAACTTTAACCCg |
| 8 | CAAAgCATTgAACACCATAACCgAA | TT | CCACTCAACTTTAACCCg |
| 10 | CgggCATggCACTCTTgAAAAAgTC | TT | CCACTCAACTTTAACCCg |
| 12 | TCTTgTAgTTCCCgTCATCTTTgAA | TT | CCACTCAACTTTAACCCg |
| 14 | TTAACTCgATTCTATTAACAAgggT | TT | CCACTCAACTTTAACCCg |
| 16 | TgTATTCCAATTTgTgTCCAAgAAT | TT | CCACTCAACTTTAACCCg |
| 18 | TgATTCCATTCTTTTgTTTgTCTgC | TT | CCACTCAACTTTAACCCg |
| 20 | CTgCTAgTTgAACgCTTCCATCTTC | TT | CCACTCAACTTTAACCCg |
| 22 | TgTCTggTAAAAggACAgggCCATC | TT | CCACTCAACTTTAACCCg |
| 24 | CTCTCTTTTCgTTgggATCTTTCgA | TT | CCACTCAACTTTAACCCg |

Organism: ***E. coli***
Target mRNA: **GTP-binding protein chain elongation factor EF-G *(fusA)***
Probe set: **18 split-initiator probe pairs (each probe carries half an HCR initiator)**
HCR amplifier: **B2-Alexa594 (Figures 6B and S26 )**

| Odd # | 1st Half of Initiator I1 | Spacer | Probe Sequence (25 nt) |
|---|---|---|---|
| 1 | CCTCgTAAATCCTCATCA | AA | gTgCgATgggTgTTgTACgAgCCAT |
| 5 | CCTCgTAAATCCTCATCA | AA | CATgAACTTCACCgATTTTATggTT |
| 9 | CCTCgTAAATCCTCATCA | AA | gCTCATACTgCTTAgCCATACCAgA |
| 13 | CCTCgTAAATCCTCATCA | AA | CAACTgCgCAgTAAACCATTACCgC |
| 17 | CCTCgTAAATCCTCATCA | AA | TCAggAAgTTCgCACCCATgCggTC |
| 21 | CCTCgTAAATCCTCATCA | AA | TCATTTTCACCAggTCAACAACACC |
| 25 | CCTCgTAAATCCTCATCA | AA | ATTCgATCAggTTCTggTgCCATTC |
| 29 | CCTCgTAAATCCTCATCA | AA | TTTCgTTgTTCAgAACgCgCTgACg |
| 33 | CCTCgTAAATCCTCATCA | AA | CgTTgATCgCAggTACgTCAACCgg |
| 37 | CCTCgTAAATCCTCATCA | AA | ggTTACCAACAAACgggTCggTAgC |
| 41 | CCTCgTAAATCCTCATCA | AA | CgATAgCAgCAgCgATgTCgCCCgC |
| 45 | CCTCgTAAATCCTCATCA | AA | TCggTTCAACTgCgATggAgATTAC |
| 49 | CCTCgTAAATCCTCATCA | AA | ATTCACgCTTCATACggTCAACgAT |
| 53 | CCTCgTAAATCCTCATCA | AA | CACgACCACCAgACTgTTTCgCgTg |
| 57 | CCTCgTAAATCCTCATCA | AA | ATTCgCCAgggATTACACCACCTTT |
| 61 | CCTCgTAAATCCTCATCA | AA | CATggTAAgAACCgAAgTgCAgACg |
| 65 | CCTCgTAAATCCTCATCA | AA | CTTCAACCTTCATgATCggCTCAAg |
| 69 | CCTCgTAAATCCTCATCA | AA | CAgCgTggATCTTAACgCCAgTAAC |

| Even # | Probe Sequence (25 nt) | Spacer | 2nd Half of Initiator I1 |
|---|---|---|---|
| 2 | gCgCACTgATACCgATgTTACggTA | AA | ATCATCCAgTAAACCgCC |
| 6 | CCATCCAgTCCATggTTgCAgCgCC | AA | ATCATCCAgTAAACCgCC |
| 10 | gggTgTCgATgATgTTgATgCgATg | AA | ATCATCCAgTAAACCgCC |
| 14 | CggTTTCAgACTgCggCTgAACACC | AA | ATCATCCAgTAAACCgCC |
| 18 | gACgggTTTTgATCTggTTAACAAC | AA | ATCATCCAgTAAACCgCC |
| 22 | ggTCAgCgTCgTTCCAgTTgATAgC | AA | ATCATCCAgTAAACCgCC |
| 26 | TCAgCTCTTCAgAAgCTTCAgCTgC | AA | ATCATCCAgTAAACCgCC |
| 30 | ACgCAgAACCACAggTTACCAggAT | AA | ATCATCCAgTAAACCgCC |
| 34 | gAgTgTCTTTACCgTCgTCCAggAT | AA | ATCATCCAgTAAACCgCC |
| 38 | CACCggAgTAAACACggAAgAAggT | AA | ATCATCCAgTAAACCgCC |
| 42 | TgTCACCAgTggTTACgTCTTTCAg | AA | ATCATCCAgTAAACCgCC |
| 46 | CCATTTTTTCCTggTCAgCTTTggT | AA | ATCATCCAgTAAACCgCC |
| 50 | gTTTACCTACgTTCgCTTCAACgTT | AA | ATCATCCAgTAAACCgCC |
| 54 | TgTCgATAACAACATgACCATACTg | AA | ATCATCCAgTAAACCgCC |
| 58 | ggATACCTTTATCAACggCCgggAT | AA | ATCATCCAgTAAACCgCC |
| 62 | TAAACgCCAgTTCAgAggAgTCAAC | AA | ATCATCCAgTAAACCgCC |
| 66 | CACCggTgTTCTCTTCCggAgTTTC | AA | ATCATCCAgTAAACCgCC |
| 70 | ATCCgAACATTTCAgACAgCggTAC | AA | ATCATCCAgTAAACCgCC |

Organism: ***E.coli***
Target mRNA: **GTP-binding protein chain elongation factor EF-G *fusA***
Probe set: **18 split-initiator probe pairs (each probe carries half an HCR initiator)**
HCR amplifier: **B3-Alexa488 (Figures 6B, S26, and S28)**

| Odd # | 1st Half of Initiator I1 | Spacer | Probe Sequence (25 nt) |
|---|---|---|---|
| 3 | gTCCCTgCCTCTATATCT | TT | TAgTAgTggTTTTACCggCgTCgAT |
| 7 | gTCCCTgCCTCTATATCT | TT | TggTAATACCACgTTCCTgCTCCTg |
| 11 | gTCCCTgCCTCTATATCT | TT | CTTCgATTgTgAAgTCAACgTgCCC |
| 15 | gTCCCTgCCTCTATATCT | TT | CTTTATATTTgTTTgCCTgACgCCA |
| 19 | gTCCCTgCCTCTATATCT | TT | gCTgCAgCggAACCgggTTCgCgCC |
| 23 | gTCCCTgCCTCTATATCT | TT | TATCTTCgTATTCgAAggTTACgCC |
| 27 | gTCCCTgCCTCTATATCT | TT | gTTCTTCACCACCCAggTATTTTTC |
| 31 | gTCCCTgCCTCTATATCT | TT | gCATCgCCTgAACACCTTTgTTCTT |
| 35 | gTCCCTgCCTCTATATCT | TT | CgTCATCACTTgCgTgACgTTCAgC |
| 39 | gTCCCTgCCTCTATATCT | TT | TCAgTACggTATCACCAgAgTTAAC |
| 43 | gTCCCTgCCTCTATATCT | TT | TgATCggCgCATCCgggTCACACAg |
| 47 | gTCCCTgCCTCTATATCT | TT | CTTTAgCCAgACggCCCAgAgCCAg |
| 51 | gTCCCTgCCTCTATATCT | TT | ggATAgTTTCACggTAAgCAACCTg |
| 55 | gTCCCTgCCTCTATATCT | TT | ggTTTgAACCCggCTCCAgCgggTA |
| 59 | gTCCCTgCCTCTATATCT | TT | CCAgCggACCTgCTTTCAgCTgTTC |
| 63 | gTCCCTgCCTCTATATCT | TT | CTTTAAAggCgATAgAAgCAgCCAg |
| 67 | gTCCCTgCCTCTATATCT | TT | gACgACggCTCAAgTCACCgATAAC |
| 71 | gTCCCTgCCTCTATATCT | TT | TggTCAgAgAACgCAgCTgAgTTgC |

| Even # | Probe Sequence (25 nt) | Spacer | 2nd Half of Initiator I1 |
|---|---|---|---|
| 4 | CACCggTgTAgAACAgAATACgTTC | TT | CCACTCAACTTTAACCCg |
| 8 | AgAATgCAgTAgTCgCAgCggAAgT | TT | CCACTCAACTTTAACCCg |
| 12 | CATCgAgAACACgCATggAACgTTC | TT | CCACTCAACTTTAACCCg |
| 16 | TTTTgTTAACgAACgCAATgCgCgg | TT | CCACTCAACTTTAACCCg |
| 20 | TgAAATgTTCTTCAgCACCAATCgC | TT | CCACTCAACTTTAACCCg |
| 24 | TAgCCAgTTCAACCATgTCTgCCgg | TT | CCACTCAACTTTAACCCg |
| 28 | gAgCACCTTTgATTTCTgCTTCAgT | TT | CCACTCAACTTTAACCCg |
| 32 | ATggCAggTAATCAATTACCgCATC | TT | CCACTCAACTTTAACCCg |
| 36 | TTTTgAACgCCAgTgCAgAgAACgg | TT | CCACTCAACTTTAACCCg |
| 40 | AACgCTCACgTgCAgCTTTCACggA | TT | CCACTCAACTTTAACCCg |
| 44 | gCTCAgggAATTCCATACgTTCCAg | TT | CCACTCAACTTTAACCCg |
| 48 | CAgTCCATACACggAAAgACgggTC | TT | CCACTCAACTTTAACCCg |
| 52 | TACCTTCAACATCggTAACTTTCTg | TT | CCACTCAACTTTAACCCg |
| 56 | TgTCgTTgATgAACTCgTAgCCTTT | TT | CCACTCAACTTTAACCCg |
| 60 | TACCCATgTCTACTACCgggTAgCC | TT | CCACTCAACTTTAACCCg |
| 64 | gAACTggTTTCgCTTTCTTAAAgCC | TT | CCACTCAACTTTAACCCg |
| 68 | CAgATTCCTgACCTTTgAgCATACC | TT | CCACTCAACTTTAACCCg |
| 72 | ATTCCATAgTgTATgATgCACgACC | TT | CCACTCAACTTTAACCCg |

Organism: ***E. coli***
Target mRNA: **isocitrate dehydrogenase (*icd*)**
Probe set: **20 split-initiator probe pairs (each probe carries half an HCR initiator)**
HCR amplifier: **B1-Alexa594 (Figure S28)**

| Odd # | 1st Half of Initiator I1 | Spacer | Probe Sequence (25 nt) |
|---|---|---|---|
| 1 | gAggAgggCAgCAAACgg | AA | CCggAACAACTACTTTACTTTCCAT |
| 3 | gAggAgggCAgCAAACgg | AA | gATTTTCAggAACgTTgAgTTTgCC |
| 5 | gAggAgggCAgCAAACgg | AA | TggCTggggTTACATCTACACCgAT |
| 7 | gAggAgggCAgCAAACgg | AA | TTTTACgCTCgCCTTTATAggCTTT |
| 9 | gAggAgggCAgCAAACgg | AA | ACCTgACCATAAACCTgTgTggATTT |
| 11 | gAggAgggCAgCAAACgg | AA | TggCAACgCgATATTCACgAATCAg |
| 13 | gAggAgggCAgCAAACgg | AA | CAACgTTCAgAgAgCgAATACCgCC |
| 15 | gAggAgggCAgCAAACgg | AA | gATAgTAACgTACCggACgCAggCA |
| 17 | gAggAgggCAgCAAACgg | AA | ggAAgATAACCATATCggTCAgTTC |
| 19 | gAggAgggCAgCAAACgg | AA | CggCAgAgTCTgCTTTCCATTCgAT |
| 21 | gAggAgggCAgCAAACgg | AA | gAATTTTCTTCACCCCCATCTCTTC |
| 23 | gAggAgggCAgCAAACgg | AA | TggTgCCTTCTTCCgAACACggCTT |
| 25 | gAggAgggCAgCAAACgg | AA | CAgAgTCACgATCgTTAgCAATTgC |
| 27 | gAggAgggCAgCAAACgg | AA | CTTTAAACgCTCCTTCggTgAACTT |
| 29 | gAggAgggCAgCAAACgg | AA | CACCgTCgATCAgTTCACCgCCAAA |
| 31 | gAggAgggCAgCAAACgg | AA | CTTTAATgACgATCTCTTTgCCAgT |
| 33 | gAggAgggCAgCAAACgg | AA | CATATTCAgCCggACgCAgCAggAT |
| 35 | gAggAgggCAgCAAACgg | AA | CTgCCAgggCgTCAgAAATgTAgTC |
| 37 | gAggAgggCAgCAAACgg | AA | CgCATTCgTCACCgATgTTTgCACC |
| 39 | gAggAgggCAgCAAACgg | AA | CTTTgTCCTgACCggCATATTTCgg |

| Even # | Probe Sequence (25 nt) | Spacer | 2nd Half of Initiator I1 |
|---|---|---|---|
| 2 | TTTgCAgggTgATCTTCTTgCCTTg | TA | gAAgAgTCTTCCTTTACg |
| 4 | CATCACCTTCAATgTAAgggATAAT | TA | gAAgAgTCTTCCTTTACg |
| 6 | CgACTgCAgCgTCgACCACTTTCAg | TA | gAAgAgTCTTCCTTTACg |
| 8 | CACCggTgTAAATTTCCATCCAggA | TA | gAAgAgTCTTCCTTTACg |
| 10 | CAAgAgTTTCAgCAggCAgCCAgAC | TA | gAAgAgTCTTCCTTTACg |
| 12 | CAACCggAgTggTCAgCggACCTTT | TA | gAAgAgTCTTCCTTTACg |
| 14 | TgTAgAgATCCAgTTCCTggCgCAg | TA | gAAgAgTCTTCCTTTACg |
| 16 | ggTgTTTAACCgggCTTggAgTgCC | TA | gAAgAgTCTTCCTTTACg |
| 18 | CCgCATAAATgTCTTCCgAgTTTTC | TA | gAAgAgTCTTCCTTTACg |
| 20 | gCAggAATTTAATCACTTTCTCggC | TA | gAAgAgTCTTCCTTTACg |
| 22 | TACCgATACCACAATgTTCCgggAA | TA | gAAgAgTCTTCCTTTACg |
| 24 | ATTCgATCgCTgCACgAACCAgACg | TA | gAAgAgTCTTCCTTTACg |
| 26 | TgATgTTgCCTTTgTgCACCAgAgT | TA | gAAgAgTCTTCCTTTACg |
| 28 | CTTCACgCgCCAgCTggTAgCCCCA | TA | gAAgAgTCTTCCTTTACg |
| 30 | TCgggTTTTTAACTTTCAgCCACgg | TA | gAAgAgTCTTCCTTTACg |
| 32 | gTTgCAggAATgCATCAgCAATCAC | TA | gAAgAgTCTTCCTTTACg |
| 34 | CgTTCAggTTCATACAggCgATAAC | TA | gAAgAgTCTTCCTTTACg |
| 36 | gggCgATACCgATACCgCCAACCTg | TA | gAAgAgTCTTCCTTTACg |
| 38 | CAgTACCgTgggTggCTTCAAACAg | TA | gAAgAgTCTTCCTTTACg |
| 40 | CggAgAgAATAATAgAgCCAggATT | TA | gAAgAgTCTTCCTTTACg |

#### S4.9 Split-initiator probes for Figures 7, S29, and S30

Organism: ***H. sapiens sapiens***
Target mRNA: **B-Raf proto-oncogene, serine/threonine kinase (*BRAF*)**
Probe set: **23 split-initiator probe pairs (each probe carries half an HCR initiator)**
HCR amplifier: **B3-Alexa647**

| Odd # | 1st Half of Initiator I1 | Spacer | Probe Sequence (25 nt) |
|---|---|---|---|
| 1 | gTCCCTgCCTCTATATCT | TT | ACCgCTCAgCgCCgCCATCTTATAA |
| 5 | gTCCCTgCCTCTATATCT | TT | ggCAgggTCCgCAgCCgAAgAggCC |
| 9 | gTCCCTgCCTCTATATCT | TT | ATATATTgATggTggATTATgCTCC |
| 13 | gTCCCTgCCTCTATATCT | TT | AgAgCTAgAAACAgAAAAATCAgTT |
| 17 | gTCCCTgCCTCTATATCT | TT | CTTggggTTgCTCCgTgCCACATCT |
| 21 | gTCCCTgCCTCTATATCT | TT | CTCTCCATCCTgAATTCTgTAAACA |
| 25 | gTCCCTgCCTCTATATCT | TT | TTTTCgTACAAAgTTgTgTgTTgTA |
| 29 | gTCCCTgCCTCTATATCT | TT | AACTTCTgTACTACAACgCTggTgA |
| 33 | gTCCCTgCCTCTATATCT | TT | TAgggCAgTCTCTgCTAAggACgCC |
| 37 | gTCCCTgCCTCTATATCT | TT | TggTCggAAgggCTgTggAATTggA |
| 41 | gTCCCTgCCTCTATATCT | TT | TCTAATCAAgTCATCAATATTgACA |
| 45 | gTCCCTgCCTCTATATCT | TT | TggAgATTTCTgTAAggCTTTCACg |
| 49 | gTCCCTgCCTCTATATCT | TT | CTgCCCATCAggAATCTCCCAATCA |
| 53 | gTCCCTgCCTCTATATCT | TT | AggTgTAggTgCTgTCACATTCAAC |
| 57 | gTCCCTgCCTCTATATCT | TT | ggTCTCAATgATATggAgATggTgA |
| 61 | gTCCCTgCCTCTATATCT | TT | CACTgTAgCTAgACCAAAATCACCT |
| 65 | gTCCCTgCCTCTATATCT | TT | CTgAAAgCTgTATggATTTTTATCT |
| 69 | gTCCCTgCCTCTATATCT | TT | ACTAAAATCCTCTgTTTggAAACCA |
| 73 | gTCCCTgCCTCTATATCT | TT | TTTTgTTgCTACTCTCCTgAACTCT |
| 77 | gTCCCTgCCTCTATATCT | TT | ATTATATCTAgTCTTTAACCACACA |
| 81 | gTCCCTgCCTCTATATCT | TT | AAgTAAAgCCTCTAgAAgAggCTCT |
| 85 | gTCCCTgCCTCTATATCT | TT | TCTTCTggAgTCCCTAgTggACATg |
| 89 | gTCCCTgCCTCTATATCT | TT | ATTAAATTCTACTgACTTCCTAAAT |

| Even # | Probe Sequence (25 nt) | Spacer | 2nd Half of Initiator I1 |
|---|---|---|---|
| 2 | gCCCggCTCCgCgCCgCCACCACCg | TT | CCACTCAACTTTAACCCg |
| 6 | TTTgATATTCCACACCTCCTCCggA | TT | CCACTCAACTTTAACCCg |
| 10 | gCTggTgTATTCTTCATAggCCTCC | TT | CCACTCAACTTTAACCCg |
| 14 | AgAAgATgTAACggTATCCATTgAT | TT | CCACTCAACTTTAACCCg |
| 18 | gACTCTAACgATAggTTTTTgTggT | TT | CCACTCAACTTTAACCCg |
| 22 | ATCAgTgTCCCAACCAATTggTTTC | TT | CCACTCAACTTTAACCCg |
| 26 | gTCACAAAATgCTAAggTgAAAAAC | TT | CCACTCAACTTTAACCCg |
| 30 | TTggTCATAATTAACACACATCAgT | TT | CCACTCAACTTTAACCCg |
| 34 | gggTgCggAAggggATgATCCAgAT | TT | CCACTCAACTTTAACCCg |
| 38 | AAATTgATTTCgATgATCTTCATCT | TT | CCACTCAACTTTAACCCg |
| 42 | TCCTCCATCACCACgAAATCCTTgg | TT | CCACTCAACTTTAACCCg |
| 46 | AgATgACTTCCTTTCTCgCTgAggT | TT | CCACTCAACTTTAACCCg |
| 50 | AgATCCAATTCTTTgTCCCACTgTA | TT | CCACTCAACTTTAACCCg |
| 54 | TTCATTTTTgAAggCTTgTAACTgC | TT | CCACTCAACTTTAACCCg |
| 58 | ATCTATAAgTTTgATCATCTCAAAT | TT | CCACTCAACTTTAACCCg |
| 62 | CTgATgggACCCACTCCATCgAgAT | TT | CCACTCAACTTTAACCCg |
| 66 | AACAATTCCAAATgCATATACATCT | TT | CCACTCAACTTTAACCCg |
| 70 | TgTTTTTggAgAAgCACAAgCATAT | TT | CCACTCAACTTTAACCCg |
| 74 | AAgCAAACATATgTTCATTTATTTT | TT | CCACTCAACTTTAACCCg |
| 78 | TAAgTATAAATTTTAgTTTggggAA | TT | CCACTCAACTTTAACCCg |
| 82 | AAgTgAATgATACAAACCCCggAACA | TT | CCACTCAACTTTAACCCg |
| 86 | CTgCAAACACAggCATAggTAgggT | TT | CCACTCAACTTTAACCCg |
| 90 | AAAATTATTAAgAATAATAATAgAA | TT | CCACTCAACTTTAACCCg |

Organism: ***H. sapiens sapiens***
Target mRNA: **B-Raf proto-oncogene, serine/threonine kinase (*BRAF*)**
Probe set: **23 split-initiator probe pairs (each probe carries half an HCR initiator)**
HCR amplifier: **B4-Alexa546**

| Odd # | 1st Half of Initiator I1 | Spacer | Probe Sequence (25 nt) |
|---|---|---|---|
| 3 | CCTCAACCTACCTCCAAC | AA | CTCCATgTCCCCgTTgAACAgAgCC |
| 7 | CCTCAACCTACCTCCAAC | AA | ATgTTCCTgTgTCAACTTAATCATT |
| 11 | CCTCAACCTACCTCCAAC | AA | TTCTCTTTgTTggAgTgCATCTAgC |
| 15 | CCTCAACCTACCTCCAAC | AA | AggTAgCACTgAAAggCTAgAAgAg |
| 19 | CCTCAACCTACCTCCAAC | AA | CACTgTCCTCTgTTTgTTgggCAgg |
| 23 | CCTCAACCTACCTCCAAC | AA | CAATTCTTCTCCAgTAAgCCAggAA |
| 27 | CCTCAACCTACCTCCAAC | AA | ACCCTggAAAAgCAgCTTTCgACAA |
| 31 | CCTCAACCTACCTCCAAC | AA | gAACTTggAgACAAACAgCAAATCA |
| 35 | CCTCAACCTACCTCCAAC | AA | AATTTggggCCCAATAgAgTCCgAg |
| 39 | CCTCAACCTACCTCCAAC | AA | AgCTgATgAggATCggTCTCgTTgC |
| 43 | CCTCAACCTACCTCCAAC | AA | gggggTAgCAgACAAACCTgTggTT |
| 47 | CCTCAACCTACCTCCAAC | AA | CATTCgATTCCTgTCTTCTgAggAT |
| 51 | CCTCAACCTACCTCCAAC | AA | TCCCTTgTAgACTgTTCCAAATgAT |
| 55 | CCTCAACCTACCTCCAAC | AA | ATgTCgTgTTTTCCTgAgTACTCCT |
| 59 | CCTCAACCTACCTCCAAC | AA | TATATTATTACTCTTgAggTCTCTg |
| 63 | CCTCAACCTACCTCCAAC | AA | CAAAATggATCCAgACAACTgTTCA |
| 67 | CCTCAACCTACCTCCAAC | AA | CTTCATggCTTTTggACAgTTACTC |
| 71 | CCTCAACCTACCTCCAAC | AA | CgCACCATATCCCCCTgCCTggATg |
| 75 | CCTCAACCTACCTCCAAC | AA | gAgAgTATTTTATTCAATTTAACAT |
| 79 | CCTCAACCTACCTCCAAC | AA | AACCCTTggATgTTAAAAATCCAAT |
| 83 | CCTCAACCTACCTCCAAC | AA | AACTgAAgTTTACTACTTAAAATAA |
| 87 | CCTCAACCTACCTCCAAC | AA | CAggCTAACCgACTgCCAACTTCTC |
| 91 | CCTCAACCTACCTCCAAC | AA | ATTgTTATAAAAAgAAATAgTTATA |

| Even # | Probe Sequence (25 nt) | Spacer | 2nd Half of Initiator I1 |
|---|---|---|---|
| 4 | ggCgCCggCgCCggCgCCggCCTCg | AT | TCTCACCATATTCgCTTC |
| 8 | ACCAAATTTgTCCAATAgggCCTCT | AT | TCTCACCATATTCgCTTC |
| 12 | gTTCCCCAgAgATTCCAATAACTgT | AT | TCTCACCATATTCgCTTC |
| 16 | gggATTTTgAAAAACTgAAAgAgAT | AT | TCTCACCATATTCgCTTC |
| 20 | gACTgTAACTCCACACCTTgCAggT | AT | TCTCACCATATTCgCTTC |
| 24 | TggAACATTCTCCAACACTTCCACA | AT | TCTCACCATATTCgCTTC |
| 28 | TTTATAACCACATgTTTgACAgCgg | AT | TCTCACCATATTCgCTTC |
| 32 | TTCCTgTggTATTgggTggTgTTCA | AT | TCTCACCATATTCgCTTC |
| 36 | ggATTTTgAAggAgACggACTggTg | AT | TCTCACCATATTCgCTTC |
| 40 | TTCTATTgTgTTTATATgCACATTg | AT | TCTCACCATATTCgCTTC |
| 44 | AgTTAgTgAgCCAggTAATgAggCA | AT | TCTCACCATATTCgCTTC |
| 48 | ACTCgAgTCCCgTCTACCAAgTgTT | AT | TCTCACCATATTCgCTTC |
| 52 | TTTCACTgCCACATCACCATgCCAC | AT | TCTCACCATATTCgCTTC |
| 56 | ATAgCCCATgAAgAgTAggATATTC | AT | TCTCACCATATTCgCTTC |
| 60 | TTTTACTgTgAggTCTTCATgAAgA | AT | TCTCACCATATTCgCTTC |
| 64 | CATTCTgATgACTTCTggTgCCATC | AT | TCTCACCATATTCgCTTC |
| 68 | CTTTTTgAggCACTCTgCCATTAAT | AT | TCTCACCATATTCgCTTC |
| 72 | CACTCATTTgTTTCAgTggACAggA | AT | TCTCACCATATTCgCTTC |
| 76 | TgTTCTTTggTTCACCTTAAAAAAA | AT | TCTCACCATATTCgCTTC |
| 80 | CAATTTTTAgCAATgTCTATgTATT | AT | TCTCACCATATTCgCTTC |
| 84 | ATAgCTggCAACAAAAgTTgCATgA | AT | TCTCACCATATTCgCTTC |
| 88 | gATCTgTTCAgTTTgCCTTATCTAA | AT | TCTCACCATATTCgCTTC |
| 92 | gAAATAAAAgACATCCACATTTTCC | AT | TCTCACCATATTCgCTTC |

Organism: ***G. gallus domesticus***
Target mRNA: **diencephalon/mesencephalon homeobox 1 (*Dmbx1*)**
Probe set: **25 split-initiator probe pairs (each probe carries half an HCR initiator)**
HCR amplifier: **B1-Alexa594**

| Odd # | 1st Half of Initiator I1 | Spacer | Probe Sequence (25 nt) |
|---|---|---|---|
| 1 | gAggAgggCAgCAAACgg | AA | CgCTCAgTgAgTTCATggCATgCAg |
| 5 | gAggAgggCAgCAAACgg | AA | AAATgATgTCAgCCAgTCTCTCCgC |
| 9 | gAggAgggCAgCAAACgg | AA | AAATgATgTCAgCCAgTCTCTCCgC |
| 13 | gAggAgggCAgCAAACgg | AA | CAgTgTCAggTATTgTggACTgggT |
| 17 | gAggAgggCAgCAAACgg | AA | CCTCCTCTCTgTCAgTCTggTCCTC |
| 21 | gAggAgggCAgCAAACgg | AA | CgCTgATgggAgACTCTgATTTTgg |
| 25 | gAggAgggCAgCAAACgg | AA | CTgCCATgTgCTggCggAATTgCTC |
| 29 | gAggAgggCAgCAAACgg | AA | gACAgTgCAgggAgCTCAgAggCgC |
| 33 | gAggAgggCAgCAAACgg | AA | TCTCAATACTTgTTgTTTTACTgTT |
| 37 | gAggAgggCAgCAAACgg | AA | gACCTCCggTgCATCTTCTTATggg |
| 41 | gAggAgggCAgCAAACgg | AA | CggTgCTAgTAAgACATTAgTAAAT |
| 45 | gAggAgggCAgCAAACgg | AA | TCACAgCAgTCCAAAgggACAgTTC |
| 49 | gAggAgggCAgCAAACgg | AA | gTTTTCCCAAAgAATgCATCgACAA |
| 53 | gAggAgggCAgCAAACgg | AA | ggggAATAAAAgCAAAAgAggCCAC |
| 57 | gAggAgggCAgCAAACgg | AA | TTTgCTCTAAgCACCATTAAgACTC |
| 61 | gAggAgggCAgCAAACgg | AA | ggAAgTgCTTAAACAggAAATTCAC |
| 65 | gAggAgggCAgCAAACgg | AA | AATTTggCTTTCATTTTTCTCCCCA |
| 69 | gAggAgggCAgCAAACgg | AA | CTAgACCAAAATgCTCTCCAAAAAg |
| 73 | gAggAgggCAgCAAACgg | AA | TAAgAACAgCTTgCATTAATCgTgg |
| 77 | gAggAgggCAgCAAACgg | AA | CTTggCCTCCAgCATTgCAgCATTT |
| 81 | gAggAgggCAgCAAACgg | AA | gAAATCACTTTgCAgTTggTgAgTT |
| 85 | gAggAgggCAgCAAACgg | AA | AAgTCCTTTgggTTggTAgAAAAgT |
| 89 | gAggAgggCAgCAAACgg | AA | gCATTTTTggTCCTAggCAATACTA |
| 93 | gAggAgggCAgCAAACgg | AA | gACCAgATgATggCCTgCAgTgAAT |
| 97 | gAggAgggCAgCAAACgg | AA | AggTAACTCAACCCACggCTTCTgC |

| Even # | Probe Sequence (25 nt) | Spacer | 2nd Half of Initiator I1 |
|---|---|---|---|
| 2 | CTgCCTgCTggTgCAggTTgTACAT | TA | gAAgAgTCTTCCTTTACg |
| 6 | ggTgCTgggATCCATAACgTgCCTC | TA | gAAgAgTCTTCCTTTACg |
| 10 | gCTTCTgCAgCTgCTCCTTCTggAg | TA | gAAgAgTCTTCCTTTACg |
| 14 | CCTCACTTgggATgCTCTgAgTCTT | TA | gAAgAgTCTTCCTTTACg |
| 18 | TAgCCTCATCCAAggTgCTCTTAAA | TA | gAAgAgTCTTCCTTTACg |
| 22 | CACTgCTAgATgAAggAgTCACggT | TA | gAAgAgTCTTCCTTTACg |
| 26 | AggAgTAATggACCAAgTTgTTggT | TA | gAAgAgTCTTCCTTTACg |
| 30 | CgTgggAgAgAgATTggTAgTACgA | TA | gAAgAgTCTTCCTTTACg |
| 34 | CATgCTgCTTTgCCCggAgCCTTAg | TA | gAAgAgTCTTCCTTTACg |
| 38 | ggTTCCAggAgTgACATgTCTggTg | TA | gAAgAgTCTTCCTTTACg |
| 42 | AgCCAgTAgCAgTgTCTgATgCAAT | TA | gAAgAgTCTTCCTTTACg |
| 46 | gCTCTTTCTgAATgTTACAggCTTA | TA | gAAgAgTCTTCCTTTACg |
| 50 | TATgTACAAgACAAAgCAggACTCT | TA | gAAgAgTCTTCCTTTACg |
| 54 | gACTAgCTACCAAAACTgAgAgAgA | TA | gAAgAgTCTTCCTTTACg |
| 58 | gAgCAgTgAATTgCATAATggTTTT | TA | gAAgAgTCTTCCTTTACg |
| 62 | AgTAAAggAAAAAACACTTgCCTTT | TA | gAAgAgTCTTCCTTTACg |
| 66 | AACAATCAAgTCAAAAgTAACCATg | TA | gAAgAgTCTTCCTTTACg |
| 70 | AgTTTTTATTgTTCTCTATTgTCgA | TA | gAAgAgTCTTCCTTTACg |
| 74 | TAgAATTTggTgATCggAgCgTTTT | TA | gAAgAgTCTTCCTTTACg |
| 78 | AATAgAAAgCCCCCgATTATCACCC | TA | gAAgAgTCTTCCTTTACg |
| 82 | AgAgAAATgAggCCAAAACTgTggA | TA | gAAgAgTCTTCCTTTACg |
| 86 | TATTTggTTTggAgAAgATAAATAA | TA | gAAgAgTCTTCCTTTACg |
| 90 | ACCTAgCACCTgCCACAgAgCCAgT | TA | gAAgAgTCTTCCTTTACg |
| 94 | CTCCATTTCTTCTTTAAATCgAgCA | TA | gAAgAgTCTTCCTTTACg |
| 98 | AACACCCTTTCCCCCTCgTgTTTAA | TA | gAAgAgTCTTCCTTTACg |

Organism: ***G. gallus domesticus***
Target mRNA: **diencephalon/mesencephalon homeobox 1 (*Dmbx1*)**
Probe set: **25 split-initiator probe pairs (each probe carries half an HCR initiator)**
HCR amplifier: **B2-Alexa647**

| Odd # | 1st Half of Initiator I1 | Spacer | Probe Sequence (25 nt) |
|---|---|---|---|
| 3 | CCTCgTAAATCCTCATCA | AA | AgTCgggCgCgTgCTgTgCTTgCTg |
| 7 | CCTCgTAAATCCTCATCA | AA | AggCCgTgCgACTTCgCCTTTgTTT |
| 11 | CCTCgTAAATCCTCATCA | AA | CACTgTggCTgCTTTCACAgTCCTT |
| 15 | CCTCgTAAATCCTCATCA | AA | ACAgggTTAggTTgAggTCTgTgCT |
| 19 | CCTCgTAAATCCTCATCA | AA | CgTCCACACCAggACTCTTgTCCAC |
| 23 | CCTCgTAAATCCTCATCA | AA | AggAgTAggAgTgAgTTTgAgCCAg |
| 27 | CCTCgTAAATCCTCATCA | AA | TggCAgACggTgTCCCCATTTCgAA |
| 31 | CCTCgTAAATCCTCATCA | AA | ggATgggTgAggACCAgACCTgCTg |
| 35 | CCTCgTAAATCCTCATCA | AA | gTAgggTgTCAAgCCCAAgggATgC |
| 39 | CCTCgTAAATCCTCATCA | AA | ACTCCAggAAgAgATgAgggTggAA |
| 43 | CCTCgTAAATCCTCATCA | AA | TCCTCCTCAAATATTTAAAgAAgAC |
| 47 | CCTCgTAAATCCTCATCA | AA | ACATTATCgCAgggATgAggTgAgg |
| 51 | CCTCgTAAATCCTCATCA | AA | gCggTggATgCTTTCAACATTgTAA |
| 55 | CCTCgTAAATCCTCATCA | AA | gggAgCgTggCTgATTTgTgACTTT |
| 59 | CCTCgTAAATCCTCATCA | AA | TCAgCTTTAgCAgAgAAgAgAgAAg |
| 63 | CCTCgTAAATCCTCATCA | AA | TCTgTCTgTgAACAAgTgCTATTAg |
| 67 | CCTCgTAAATCCTCATCA | AA | TTACACTTCACTgAAgACCAAAgAg |
| 71 | CCTCgTAAATCCTCATCA | AA | gTATgAACACAgTgggAgTTCATAC |
| 75 | CCTCgTAAATCCTCATCA | AA | AgggTCTgAAgCTgCACAgCTTgAg |
| 79 | CCTCgTAAATCCTCATCA | AA | CAAgCCAATCTACTCCTCgCTgCAg |
| 83 | CCTCgTAAATCCTCATCA | AA | TggATTACTAAAATgAAgggTCATT |
| 87 | CCTCgTAAATCCTCATCA | AA | TTCTTggTACggTgAgTTCAAAggA |
| 91 | CCTCgTAAATCCTCATCA | AA | gACCCgCCTgACACCCTTTggATTC |
| 95 | CCTCgTAAATCCTCATCA | AA | gTTATgTAggCTATgCACACgTTgC |
| 99 | CCTCgTAAATCCTCATCA | AA | TTTTTAAgATgCATTATTgCAgTTg |

| Even # | Probe Sequence (25 nt) | Spacer | 2nd Half of Initiator I1 |
|---|---|---|---|
| 4 | gggTgAgggCgTgCACTgAgggCCg | AA | ATCATCCAgTAAACCgCC |
| 8 | CCAgTgCCTCCAgCTgTTgggCAgT | AA | ATCATCCAgTAAACCgCC |
| 12 | CCAgCACAggTggTTCCgTCTTCCC | AA | ATCATCCAgTAAACCgCC |
| 16 | gCgCTgATTCACTggCTgACTgCTC | AA | ATCATCCAgTAAACCgCC |
| 20 | TCgCTCTCTTgCAgTTCAAAgCCTT | AA | ATCATCCAgTAAACCgCC |
| 24 | gCAggCggAAgAggCTCAggggTgA | AA | ATCATCCAgTAAACCgCC |
| 28 | TgTTgACATTCATgCCCAAgTAggg | AA | ATCATCCAgTAAACCgCC |
| 32 | ggCTTggAAgAgAgCTggAggCCTg | AA | ATCATCCAgTAAACCgCC |
| 36 | ACggTTgCCCATCTggCAgTCAgTT | AA | ATCATCCAgTAAACCgCC |
| 40 | AAAgTTTTCCCTgATAgggAgCACC | AA | ATCATCCAgTAAACCgCC |
| 44 | CTgTCTAAACACACATCCTCTCCCT | AA | ATCATCCAgTAAACCgCC |
| 48 | AAAAAgggTgTATATAACACggTTg | AA | ATCATCCAgTAAACCgCC |
| 52 | TTCTgTAACACTgACAgTAACACAC | AA | ATCATCCAgTAAACCgCC |
| 56 | AACCCAAgAAgAgCAACTAgCTgTg | AA | ATCATCCAgTAAACCgCC |
| 60 | TTgCATCATTTCCTggCCgTTATAA | AA | ATCATCCAgTAAACCgCC |
| 64 | CAgCAgCATTTggCCAgCATTTTgT | AA | ATCATCCAgTAAACCgCC |
| 68 | AACCCCATAATTTgTAAATgggggA | AA | ATCATCCAgTAAACCgCC |
| 72 | TTgTCAAggAACCATATAATTCATg | AA | ATCATCCAgTAAACCgCC |
| 76 | CACTTgTTACATTCTCACTTgCTAA | AA | ATCATCCAgTAAACCgCC |
| 80 | ggTTgCTTggggACATggTACTTTT | AA | ATCATCCAgTAAACCgCC |
| 84 | CTTCTCAAgAAggAAAAACACTCTg | AA | ATCATCCAgTAAACCgCC |
| 88 | TTAgATCTgggTTTCCTCCCTCCCT | AA | ATCATCCAgTAAACCgCC |
| 92 | gCCCAgCTCTgCTgCgTgTTAgTgg | AA | ATCATCCAgTAAACCgCC |
| 96 | TggTATgAAgTAAgATgggAgCAAg | AA | ATCATCCAgTAAACCgCC |
| 100 | TgCCTCAgTTTAAgggATTTAgATg | AA | ATCATCCAgTAAACCgCC |
